# Shark tooth regeneration: RNAseq reveals genes for unlimited dental renewal

**DOI:** 10.1101/2025.08.26.672487

**Authors:** Alexandre P. Thiery, Kyle J. Martin, Katherine James, Rory L. Cooper, Ariane S. I. Standing, Wesley A. Dillard, Cameron Howitt, Ella F. Nicklin, Karly E. Cohen, Steven R. Byrum, Zerina Johanson, Gareth J. Fraser

## Abstract

Sharks are masters of tooth regeneration with a rapid and unlimited tooth supply. We present a comprehensive transcriptomic analysis across five distinct compartments from the embryonic shark mouth, capturing the full sequence of shark tooth development. Differential gene expression and gene regulatory network analyses reveal novel candidate markers upregulated within epithelial stem cells of the dental lamina. We show a potential role for this conserved regenerative network in the regulation of tooth production in sharks. Notably, we find the proto-oncogene *mycn* co-expressed with a definitive dental stem cell marker, *sox2*. Perturbation of *mycn* disrupts stem cell proliferation, underscoring its role in regulating dental stem cells in sharks. Our data showcase the power of transcriptome-based developmental approaches in the identification of predictive gene networks governing unlimited dental regeneration. The shark dental lamina provides an accessible system for the study of regeneration, offering exciting opportunities for investigating translational mechanisms of natural tooth renewal.

**Teaser:** Sequencing dental stem cells in sharks reveals new genes necessary for lifelong tooth development.

## Introduction

The diversity of dental regeneration in vertebrates is remarkable, with most toothed vertebrates capable of producing multiple generations. Sharks (and other elasmobranchs) have a unique system that provides unlimited and rapid tooth production throughout their lives. Therefore, sharks and their ilk offer an unrivalled set of models for the study of tooth development and regeneration (*1*, *2*).

Teeth in almost all vertebrates arise from an epithelial unit called the dental lamina (DL(*3*)); a crucial structure for the initiation, development and continued replacement of teeth (*3*, *4*). During embryogenesis, prior to the development of the first teeth, a field of dental competence known as the odontogenic band (OB) is established (*5–7*), which ultimately emerges, after proliferation and invaginated growth, as the established dental lamina (*5*, *8*, *9*). Regeneratively restricted dentitions (i.e., monophyodonty and diphyodonty in mammals) are coupled with a loss or break-down of the DL (*3*, *10*, *11*). There is overwhelming evidence that polyphyodonty (i.e., continuous tooth replacement) depends on the presence and maintenance of an epithelial stem cell niche found within the DL (*3*, *4*, *8*, *12–16*). At the deepest and lingual-most extent of the shark DL, lies the successional lamina (SL); the end point of the DL. The SL is the site of new tooth initiation, where all tooth generations are formed. The dental and successional lamina are characterized by sex-determining region Y-related box 2 (Sox2) positive dental progenitors initially associated with a tastebud region on the marginal oral surface (*8*, *15*). We have previously shown that cyclical upregulation of Wnt/ß-catenin signaling and enhanced proliferation of the SL are coupled with the onset of dental initiation within this Sox2+ progenitor niche (*16*). For instance, the Sox2 transcription factor is a primary marker of dental epithelial stem cells in vertebrate models (*15*–*18*), with Wnt/ß-catenin signaling playing a critical role in initiating dental development (*15*, *16*). Sox2 negatively regulates Wnt/ß-catenin signaling in osteoblasts (*19*), whilst the expression of constitutively-activated ß-catenin specifically within Sox2+ cells is sufficient to induce dental initiation in mice (*11*, *20*).

Research into successional dental regeneration in a diverse range of polyphyodont vertebrates is starting to uncover even more mechanisms through which lifelong dental regeneration is regulated (*3*, *4*, *8*, *12–16*). Importantly, in sharks (and most polyphyodont vertebrates), the embryonic dental lamina is maintained in its original embryonic state throughout life for a singular function – to make teeth and repeat. Although a continuous and permanent dental lamina may not be the basal condition for the gnathostome clade (*21*), the evolution of this character in elasmobranchs (and deeper clades of Chondrichthyans) has led to an incredible capacity for lifelong and rapid tooth regeneration, observed in extant elasmobranchs (*3*, *16*). Within sharks, many teeth develop ahead of function, aligned in discrete family units, with functional teeth at the jaw margins, followed by a developmental series of teeth embedded and hidden within the dental lamina (from the lingual extent of the DL to the functional teeth; Figure 1).

**Figure 1.**
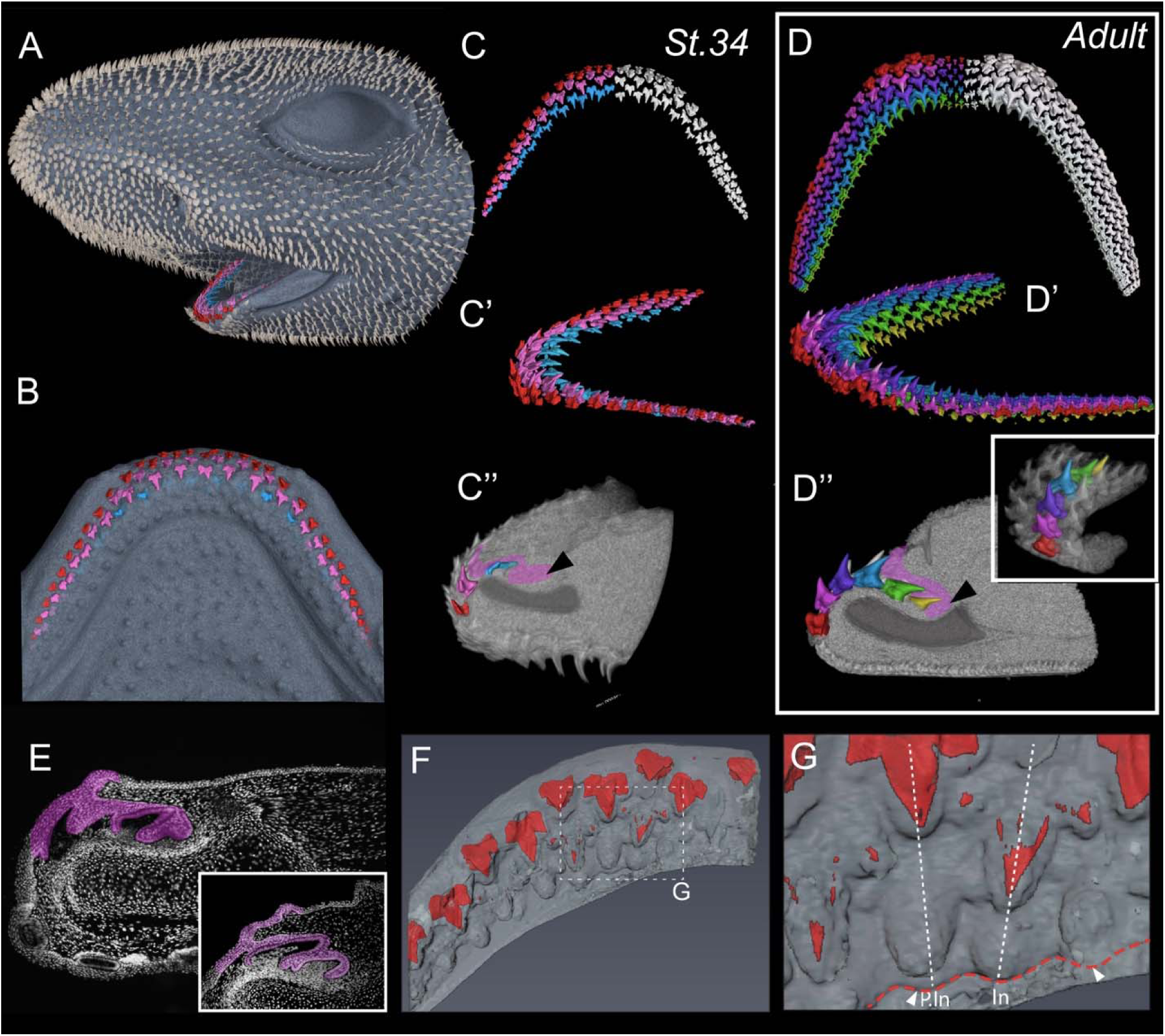
Micro-computed tomography (micro-CT) of the small-spotted catshark (*Scyliorhinus canicula*) dentition. Micro-computed tomography imaging of a Stage 34 catshark head shows mineralized denticles on the skin surface and tooth rows in the mouth (A). CT segmentation of the hatchling lower jaw reveals the tooth rows embedded within the jaw (B) and with only teeth segmented (C and C’). The first tooth row to develop (red) is followed by the second generation (pink), and then by the newest developing tooth generation (pink) (C). (C”) virtual CT section in sagittal view of the hatching lower dentition within the jaw; pink region demarcates the region of the dental lamina (black arrowhead marks the successional lamina). (D, D’, D”) equivalent CT images of the adult catshark dentition with increased number of tooth rows (6) red row is the oldest and the yellow row is the newest (inset image shows the tooth replacement whorl). (E) DAPI-counterstained histological sagittal section through a hatching catshark lower jaw at the 3-4 tooth stage and (inset) the two tooth stage (false-colored dental lamina in magenta).(F) CT-rendering of a dorsal view of the lower jaw dentition showing teeth (red) virtually exposed to show the progress of mineralization towards the jaw margin, with newer tooth rows covered with epithelium of the dental lamina. (G) Dotted region from F showing the tooth families at various stages of maturation; pre-initiation (P.In) phase of the successional lamina shows no sign of the new tooth placode (3rd row), compared to neighboring teeth undergoing the initiation phase (In).

Our study investigates the full gene repertoire of the dental lamina to uncover novel candidates associated with regulation of shark tooth regeneration. The small-spotted catshark (*Scyliorhinus canicula*) has multiple stages of tooth development at any given stage of life. Thus, the shark dental lamina offers a unique opportunity to study the complete developmental transcriptional output of cyclical and repeated tooth development. We performed RNAseq on five sub-regions of the dental lamina and oral epithelium in the shark (*Scyliorhinus canicula*; Figure 1): (i) the basihyal taste buds (BHTB; a non-dental tissue-outgroup*); (ii) the taste-tooth junction at the marginal aspect of the oral jaws (TTJ); (iii) the successional lamina (SL); (iv) early developing teeth within the DL (ET); and (v) late-stage developing teeth within the DL toward eruption (LT). *We classify the basihyal taste bud region as a non-dental outgroup due to its close relationship in terms of developmental proximity to the dental lamina and emergence of teeth, although in *Scyliorhinus canicula* this region never develops odontodes (teeth or tooth-like appendages, e.g., internal denticles (*22*)). By comparing gene expression from these distinct compartments, we identify differentially expressed genes (DEGs), including those within the progenitor cell population associated with the successional lamina. Therefore, the transcriptomic output from the SL compartment, specifically, is directly related to the regenerative production of unlimited teeth, conserved across all sharks and likely among vertebrates capable of producing multiple tooth generations.

## Results

### Stages of shark tooth development and preparation for de novo dental transcriptome assemblies

At the time of hatching from the egg case (stage 34; Figure 1A), the catshark (*Scyliorhinus canicula*) has between 3 and 5 generations of teeth at various stages of morphogenesis (Figure 1A-C”, E-G)); from erupted teeth at the jaw margin (Figure 1; red teeth) through a descending developmental series of stages, with newly initiating teeth emerging from the successional lamina (SL) at the distal ‘free-end’ of the dental lamina (DL; Figure 1C”, E-G). The adult catshark jaw exhibits 6-7 contemporaneous generations of teeth (Figure 1D – D’’), consistently presenting a complete developmental series from initiation to function at the jaw margin. All sharks, regardless of age, possess a developmental series of teeth in the jaws, from initiation to functional positions, showing that the dental lamina constantly produces new teeth throughout life, from embryogenesis to mature adult and beyond.

We selected the hatching stage of shark development (St. 34;(*23*)) for ease of complete tissue collection necessary for the subsequent transcriptome sequencing. At this stage, we were able to collect the complete series of developmental tooth stages including the dental lamina (Figure 1A-C”). Each oral tooth row is connected within a continuous jaw length DL (Fig 1E, F). Adjacent tooth families are staggered in their development, preventing adjacent teeth from overlapping (Figure 1). Segmented micro-computed tomography (micro-CT) scans reveal asynchronous timing of dental initiation, with visible undulations associated with each tooth family that can be seen at the aboral tip of the successional lamina (SL) (Fig 1F, G). Virtual sagittal sections at equivalent sites along the jaw provide further insight into the relationship between tooth developmental stage and the SL (Fig 1E). During morphogenesis, the SL retracts toward the developing tooth (Fig 1G). At this stage, the SL is in pre-initiation (*P.In*; Figure 1G), a phase marked by an absence of proliferation and Sox2+ dental progenitor cells (*16*).

Simultaneously, the adjacent tooth families undergo initiation (*In*; Figure 1G). During this stage there is a visible outgrowth of the SL both in 3D micro-CT segmentation (Fig 1F/G) and in sagittal cross section (Fig 1E, shown in magenta). Initiation of dental regeneration corresponds to a significant increase in activated β-catenin localization and cellular proliferation throughout the SL, including Sox2+ dental progenitors, suggesting a role for canonical Wnt signaling in the regulation and cyclicity of this process (*16*). A detailed understanding of shark tooth histological morphology was essential for the preparation and isolation of individual cellular compartments for the following transcriptome analyses.

### Pseudo-time series RNAseq analysis of the catshark dental lamina

To identify novel candidate markers involved in regulating the onset of dental regeneration within the DL, we micro-dissected (see Method scheme, Figure S1) and carried out RNA sequencing of dental sub-compartments associated with each stage of odontogenesis in hatchling catsharks (*Scyliorhinus canicula*; Figure 2, A-E). In total, five sub-regions were dissected (Figure 2A-F; color matched throughout the figures): (*i*) successional lamina (SL; red); (*ii*) early developing teeth (ET; orange); (*iii*) late developing teeth (LT; yellow); (*iv*) the taste-tooth junction (TTJ; magenta) and (*v*) the basihyal taste bud territory (BHTB; blue; abbreviated to TB in Figure 2). The TTJ houses a bifunctional Sox2+ progenitor cell population, which contributes to both the developing teeth and oral taste buds (BHTB; (*16*). Cells from the TTJ actively migrate to the successional lamina (SL), which houses dental progenitors and is the site of new tooth initiation and regeneration (*5*, *16*, *24*). The taste bud region associated with the basihyal (BHTB) was sampled as a ‘*non-dental*’ outgroup, compared to the TTJ (Figure 2C, magenta), which lies at the dental competent junction. Developmentally downstream of the SL (Figure 2F), we sampled early stage developing teeth (ET) at the bud and cap stage. Late-stage teeth (LT; Figure 2E and F) is an odontogenic phase involving matrix secretion and mineralization prior to eruption. This sequence of shark tooth development is referred to as a conveyor-belt (*5*) and creates a unique dental system of multiple tooth rows, forming ahead of function and lining the jaw, fed by a progenitor niche at the SL (the site of regeneration; Figure 1 and 2A-F). The lifelong maintenance of progenitor cells within the SL permits the repeated, conveyor-belt system of tooth production in all elasmobranchs.

**Figure 2.**
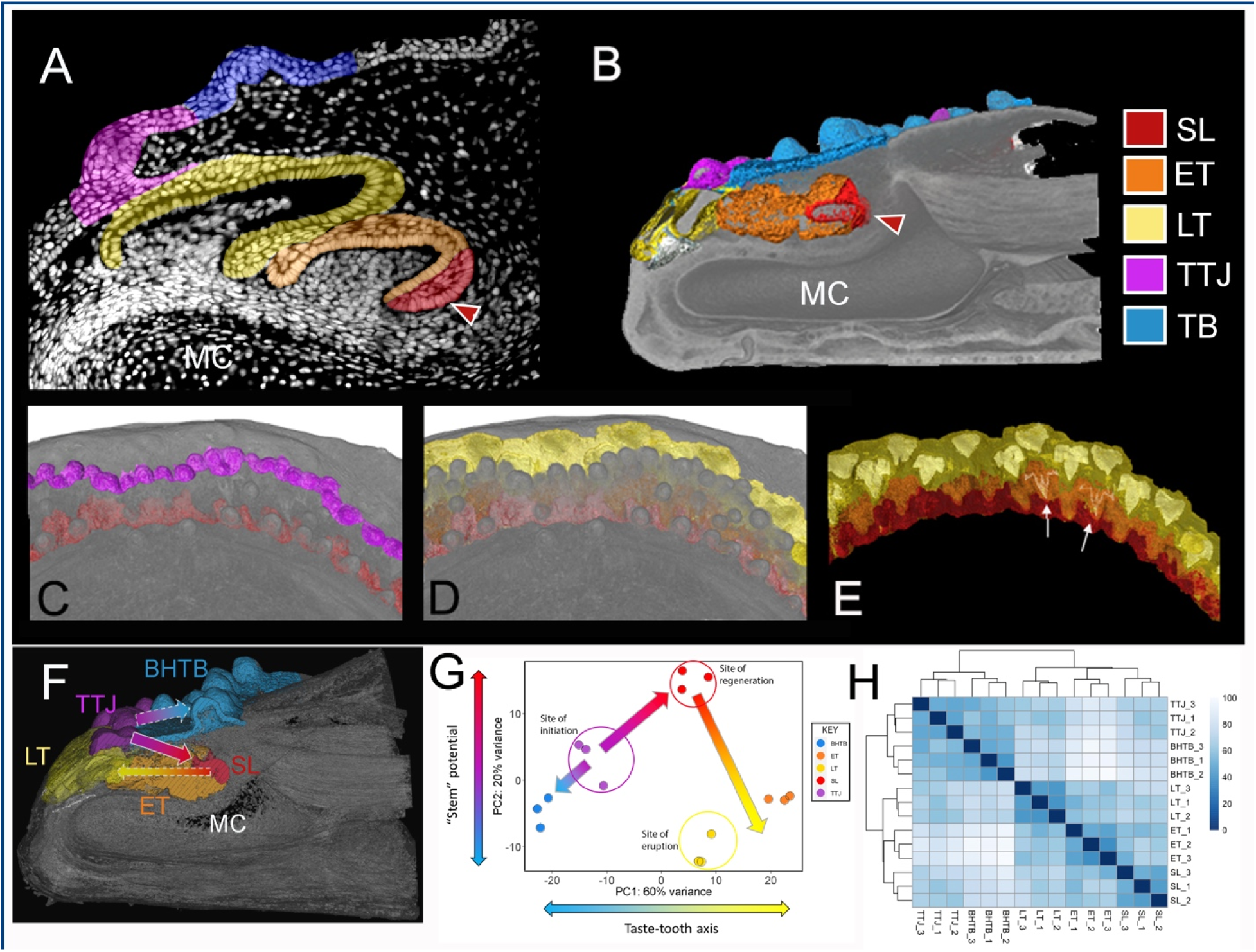
Dental lamina subdivisions: sub-compartments micro-dissected for RNAseq analysis. A) DAPI counter-stained (greyscale) sagittal cross section through the embryonic catshark (*S. canicula*) lower jaw depicting the color-coded (throughout the manuscript) sub-regions dissected for RNAseq: Key = successional lamina (SL) red; Early Teeth (ET) orange; Late Teeth (LT) yellow; Tooth-Taste Junction (TTJ) magenta; Basi-Hyal Taste Buds (BHTB) blue. B) sagittal virtual section (micro-CT) showing each of the same 5 color-coded compartments of the stage 34 (hatching) catshark lower jaw (equivalent stage of dissection for the tissues used for RNAseq). MC, Meckel’s cartilage. (C-E) μCT dorsal images of the catshark lower jaw (St. 34) showing the segmented rendering of separate compartments: Red, SL where new teeth initiate; magenta at the level of the TTJ, the initial stem niche for tooth initiation in sharks; yellow and orange separate the late and early tooth stages (respectively). Pseudo-time is represented by arrows in (F), where the TTJ acts as a bifunctional progenitor niche from which cells migrate to regions of taste bud development (BHTH; blue) and to the successional lamina (SL; red), before teeth are made and enter the cycle of early tooth development (ET; orange) and then lastly the late tooth development phase where teeth mineralize and move into functional position before exfoliation (LT; yellow). (G) pseudo-time is represented again in a principal component analysis (PCA) of the RNAseq replicates showing the taste-tooth axis (PC1) with 60% of the variation among the compartments. PC2 shows the relative “stem potential” for each compartment represented by the Y-axis of variation (PC2: 20%) with the SL (Red) replicates at the site of tooth regeneration where stemness is high, compared with the initial site of competence (TTJ; magenta), and the other compartments toward the terminal phase of the cycle that end with eruption of the functional teeth (LT; yellow). (H) Quality control plot of the RNAseq data showing sample-sample clustering. Samples cluster clearly by tissue type not sample, revealing that expression differences between tissue samples is mostly the result of tissue type and not variation in the biological replicates.

A *de novo* transcriptome was assembled using Trinity and annotated with Trinotate, to which sample reads were aligned. Reads were normalized and differential expression was computed with DESeq2. A principal component analysis (PCA) was performed on normalized read counts confirming a high degree of concordance between pooled sample replicates from each tissue type (Figure 2G). This ordering of samples along principal component 1 (PC1) primarily reflects the spatial arrangement of their respective tissues during dental development (Fig 2G), explaining 60% of the variation in gene expression between the extreme end points, i.e., from the taste territories (BHTB and TTJ) to developing teeth (ET/LT; orange and yellow, respectively; Figure 2G), at the end of the dental conveyor belt. In contrast, PC2 correlates with cell differentiation; the three most differentiated tissues (ET, LT and BHTB) are found at the bottom of the axis, and the two ‘stem/progenitor’ niches (TTJ and SL (*16*)) cluster at the top, and are closely linked via cell migration from the TTJ niche into the deeply invaginated SL (*16*). Sample distance matrices on log-normalized read counts show that tissue replicates (3 replicates per tissue compartment) were highly correlated, highlighting the reliability of the compartmental dissections (Figure 2H; Figure S1). Furthermore, clustering of our sample distance matrices (Figure 2H) also reveal transcriptional relationships between tissues, e.g., SL, ET and LT samples cluster together, reflecting the developmental relationship between these tissues; cells from the SL contribute directly to the epithelial compartment of the ET and subsequently to the LT morphogenesis, all embedded within the invaginated dental lamina (*3*). In contrast, TTJ and BHTB samples cluster closely within a separate taste node, as both regions contain functional taste papillae (Figure 2).

### Differential expression analysis (DEA) of the shark dental lamina compartments

To explore transcriptional differences between tissue compartments and identify the unique transcriptional signature of each, we ran Differential Expression Analysis (DEA) using a likelihood ratio test (LRT; Figure 3). This approach works by fitting and comparing two models of gene expression to identify genes which vary as a function of tissue, preventing the need for multiple pairwise comparisons which are difficult to interpret. First, we use a full model whereby expression is modelled as a function of tissue, with the additional co-variate of the biological sample from which the tissue was collected. For the second, or *reduced* model, the grouping variable (tissue) was removed. Once differentially expressed genes were identified (FDR < 0.0001), they were then clustered in order to visualize expression across the five dental compartments.

**Figure 3.**
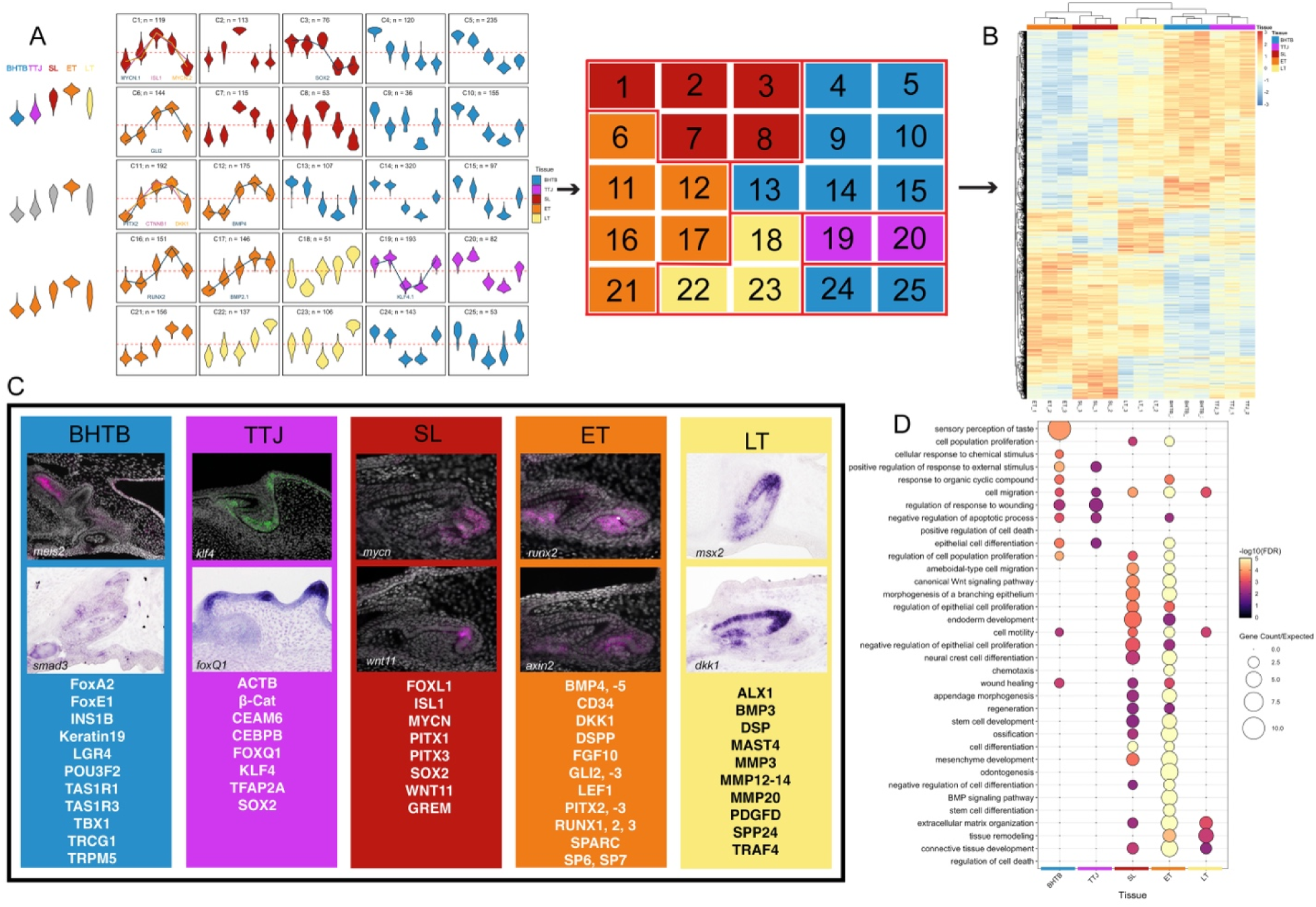
Self-Organizing maps of the DEGs that vary as a function of tissue distributed within 25 gene clusters and functional enrichment across all compartments of the dental conveyor belt in sharks. (A) SOM violin plots of the gene expression range for each cluster (1–25), color-coded based on the compartment (see key). The number of genes per cluster are represented on each cluster plot. Tissue identities were established for each cluster based on the tissue which exhibited the highest median expression. Expression of key genes (present within the SL GRN; Figure 5) is overlaid in a selection of super-clusters (Cluster 1: MYCN.1, –.2, ISL1; Cluster 3: SOX2; Cluster 6: GLI2; Cluster 11: PITX2, CTNNB1, DKK1; Cluster 12: BMP4; Cluster 16: RUNX2; Cluster 17: BMP2.1; Cluster 19: KLF4.1). (B) 3275 genes vary as a function of tissue according to their positioning along the axis of dental development; this is represented by a heatmap of differentially expressed genes. (C) Each compartment consists of a list of DEGs (with some expected overlap) that include tissue specific transcripts. A subset of markers that validate the function of the compartment are represented (taken from clusters assigned to one of the functional compartments (BHTB, TTJ, SL, ET and LT), including representative gene expression. (D) Gene ontology enrichment analysis was undertaken on each of the super-clusters (groups of clusters assigned to any given tissue/compartment), and we identified enrichment pathways for each tissue compartment. Color gradients reflect the adjusted p-values; lower p-values (dark red) reflect higher enrichment, whereas higher p-values (lighter color) indicate lower enrichment.

Given the arrangement of tissue compartments along a developmental ‘conveyor belt’, gene expression can be visualized as a pseudo-time series with tissues ordered according to their position along the axis of dental development (i.e., BHTB <=TTJ=>SL=>ET=>LT). To elucidate expression dynamics during dental regeneration, differentially expressed genes identified from our LRT analysis were clustered into 25 clusters using Kohonen Self-Organizing Maps (SOM;(*25*)) (Figure 3A). Tissue identities (SL, ET, LT, TTJ, and BHTB; Figure 3A) were then assigned to each cluster by noting which tissue exhibits the highest median expression for any given gene cluster.

(***i***) ***Successional Lamina (SL) SOM Clusters.*** Clusters 1, 2, 3, 7 and 8 were found to be upregulated in the SL (Figure 3A and C). Interestingly, the clusters that show higher levels of expression in the ET replicates also show a similar level of expression in the SL, for example clusters 6, 11, 12 and 16, 17 and 21 (Figure 3B, where SL clusters with ET within the heatmaps). This is most likely due to the physical connection between these juxtaposed compartments (SL and ET; Figure 2) without a clearly defined boundary; teeth initiate within the SL and then become the ET before developing and moving away from the SL as the cycle repeats. We superimposed a number of the key/novel DEGs onto the SOM clusters to show the individual levels of expression for each (Figure 3A), prioritizing the SL compartments, i.e., Cluster 1 (119 DEGs) shows levels of *mycn, foxL1* and *pitx1* peaking in the SL; Cluster 3 (76 DEGs) shows the specific expression level of *sox2*, upregulated in the SL, BHTB and TTJ compartments compared to ET and LT. Interestingly, we do see a small cluster of genes upregulated specifically in the SL and with some of these showing upregulation in the TTJ; given the fact that these two compartments are linked developmentally and share stem characteristics, it would be expected that they share genes associated with stem regulation and maintenance. Cluster 5 (235 DEGs) shows the relative levels of *foxA2* in taste bud-rich territories of the BHTB and TTJ, versus the rest of the compartments; and Cluster 11 (192 DEGs) where we see higher levels of *lef1*, *pitx2* and *ctnnb1* (*ß-catenin*) associated with the SL and ET compartment replicates; then Cluster 12 (125 DEGs) shows the relative expression levels of *bmp4* in SL, ET and LT compartments, compared to the BHTB and TTJ compartments. We selected a number of key transcripts (see Figure 3C) from each of the tissue specific SOM clusters (listed in Figure 3C), coupled with selected in situ hybridization results, again further validation that genes identified from tissue-specific dissections and assigned a tissue cluster showing localization to those particular tissues within the dental lamina or taste territories. Note that some genes can show more generic expression across the lamina compartments, e.g., *smad3* from the BHTB cluster, and *foxQ1* in the TTJ and BHTB compartments, even though the assigned cluster is TTJ (Figure 3C). The successional lamina gene localization of *mycn* and *wnt11* (Figure 3C) also show the proximity of the early tooth compartment with expression extending beyond the shared and adjacent territory.
(***ii***) ***Early Tooth (ET) SOM Clusters***. Six clusters (6, 11, 12, 16, 17 and 21) were upregulated in the ET compartment, marking the onset of the next generation of tooth development in the compartment adjacent to the SL. The proximity of SL and ET compartments in juxtaposed regions of the dental lamina will likely produce an overlap in gene expression (see previous section). Markers upregulated in these clusters include genes associated with tooth initiation, and odontoblast and ameloblast differentiation e.g., *dkk1*, *cd34*, *sparc*, *bmp4*, –5, *DSPP*, *runx1*, –2, –3, *lef1*, *pitx2* and –3, *gli2*, –3, and *fgf10*. Interestingly, *sp6* is also upregulated. When mutated, this gene is associated with the human congenital disorder *Amelogenesis imperfecta*, which leads to abnormal enamel formation (*26*). Similarly, *sp7* (osterix) is upregulated in the ET cluster (11) and is linked to normal enamel and dentine formation, and necessary for the maturation of odontoblasts. Clearly, the genes associated with each of these ET clusters show high propensity for functions associated with general tooth development, and differentiation of the enameloid and dentine-producing cells. Importantly, many of these markers have not been described in association with vertebrate dentitions beyond mammals, demonstrating that elements of tooth development are highly conserved and evolutionarily stable, from sharks to mammals. [*N.B.* there are distinct differences in the development of enamel and enameloid; enamel is a mineralized product of the ameloblasts in teeth of tetrapods and some fish (Sarcopterigians), and enameloid is formed from a collagen-rich matrix of combined origin from both the ameloblasts (inner dental epithelium) and odontoblasts (dental mesenchyme) in some fishes and elasmobranchs (*27*)].
(***iii***) ***Late-Stage Tooth (LT) SOM Clusters***. Clusters 18, 22, and 23 were upregulated in the LT compartment (Figure 3A and C). Therefore, we expected this gene list to include markers associated with characters of tooth maturation and matrix secretion, i.e., tooth mineralization, patterning and movement. These functions are representative of the terminal stage of the shark dental conveyor-belt (Figure 2). Specifically, from Cluster 18: *alx1*, *pdgfd*, *sema3C*; Cluster 22: *mmp20* (and mmp –3, –12, – 13, –14), spp24SPP24, *bmp3* and 7; Cluster 23: *mast4*, *traf4*, and *dsp* (desmoplakin) are all expressed and involved in aspects of tooth mineralization, and differentiation of either odontoblasts or ameloblasts (some genes are expressed across the ET/LT division, reflecting the complete process of differentiation and mineralization that likely spans the entirety of mineralization), or both (full list of genes for each cluster are present in Supplemental Figure S5). After mineralization is complete, teeth become functional and remain at their marginal endpoint for a short period (in some species, 10 days (*2*, *5*)) before exfoliation and loss, followed closely by the next generation row. The production and subsequent loss of teeth in sharks can occur very rapidly (*2*).
(***iv***) ***Tooth-Taste Junction (TTJ) SOM Clusters***. Clusters 19 and 20 are composed of genes upregulated in the region that separates the tooth field from the taste territories (Figure 3). This zone has been described as a primary progenitor niche as the dental lamina initiates via an invagination event (*16*). Cells from the TTJ migrate through the dental lamina and contribute to the developing teeth and the successional lamina (*16*). Within this compartment (Figure 3C), the cluster gene list includes both taste (*foxQ1*) and tooth transcript signatures (β*-catenin*), in addition to known markers present in both (*cebpb*) and those genes commonly expressed in association with proliferation (progranulin [GRN], *tfap2a*, *actb* – a factor that could also play a role in cell movement within the dental lamina) and stem cell factors (*klf4*, *ceam6*). The TTJ is clearly a dynamic region of the dental cascade where cells can be held in a progenitor state and moved out into the invaginated extent of the dental lamina to contribute to tooth development. The role of the TTJ in the cyclical activation of repeated tooth initiation is yet to be determined, as is whether the activity of this site continues after embryogenesis. This TTJ region is conserved among disparate vertebrate clades that possess multiple tooth generations, as a similar SOX2+ tooth/taste region has been identified in the bearded dragon (*28*).
(***v***) ***Basihyal Taste Bud region (BHTB) SOM Clusters***. Clusters 4, 5, 9, 10, 13-15, 24, and 25 contain transcripts found to be upregulated in the BHTB compartment (Figure 3). As validation, the BHTB compartment presents the expression of a number of key taste-related genes, including taste-receptor markers *trpm5*, *trcg1*, *tas1r1*, *tas1r3*; and a host of other genes associated with endoderm development more generally, like *foxa2*, and taste bud homeostasis and regeneration, i.e., *lgr4*; keratinocyte proliferation regulators *pou3f2*, *ins1b*, *keratin19*; and more general oral development, i.e., *foxe1* and *tbx1*.

### Functional enrichment analysis of the tissue specific SOM Maps

We then took our super-cluster data (groups of clusters, which are then assigned to tissues) and ran gene ontology enrichment (functional enrichment analysis; Figure 3D) to identify enriched pathways within each of the dental compartments. For each cluster, we took the tissue/compartment for which it was associated and then ran GO enrichment analysis (Figure 3D). GO terms represent tissue specific processes, e.g., in the taste territory (BHTB) the most enriched pathway, was unsurprisingly, “*sensory perception of taste*” and in the early tooth (ET) compartment “*odontogenesis*”. Within the SL and ET, where dental regeneration is initiated and next generation teeth begin their development, “*canonical Wnt signaling pathway*”, “*morphogenesis of a branching epithelium*” and “*regulation of epithelial cell proliferation*”, “*regeneration*” and “*stem cell development*” were all upregulated, validating the presence of the epithelial stem cell niche (Figure 3D). Enriched gene ontology pathways associated with the Late Tooth (LT) compartment, included “*regulation of cell death*”, “*connective tissue development*”, “*tissue remodeling*”, and “*extracellular matrix organization*” (Figure 3D). The Late Tooth (LT) represents the terminal phase of the dental conveyor-belt where marginal teeth complete mineralization; odontoblasts and ameloblasts secrete a collagen matrix which acts as a scaffold for mineralization. Then as mineralization completes, ameloblasts undergo apoptosis, facilitating eruption of the functional teeth, before exfoliation.

### Regulatory Network Enrichment Analysis (RNEA) as a predictive GRN for continuous tooth production

We know that odontogenesis is initiated in the SL, and that sox2 marks epithelial stem cells that govern this process of tooth renewal. However, we still do not understand how dental stem cells within the shark successional lamina are regulated. Focusing on this tissue region and identifying predicted regulatory interactions allows us to identify novel regulators. Classically, GRNs serve as a tool for describing genes, regulators, and their interactions (*29*, *30*). These interactions require functional testing before we assume that they are real. However, predictive GRN analysis can also serve as a tool to identify novel candidates regulating a process of interest. RNEA functions through projecting differentially expressed genes derived from transcriptome analyses onto a reference map of known (published) regulatory interactions. RNEA is able to add supplementary nodes, which are not initially included in the dataset, if it deems them statistically important (*31*). This process allows the identification of potentially novel regulators of tooth regeneration (Figure 5). Therefore, to identify novel candidates involved in the regulation of initiation and continuation of shark dental regeneration, gene regulatory network (GRN) analysis using RNEA was implemented using genes which were differentially expressed in the SL relative to other tissues (748 genes upregulated and 642 genes downregulated in the SL relative to other tissues; absolute log2FC > 0.75 and FDR < 0.001; Supplemental table S1 <SLvsALL_dea_res.tsv>) using RNEA (Figure 4; filtered from a Global GRN, Figure 4A).

**Figure 4.**
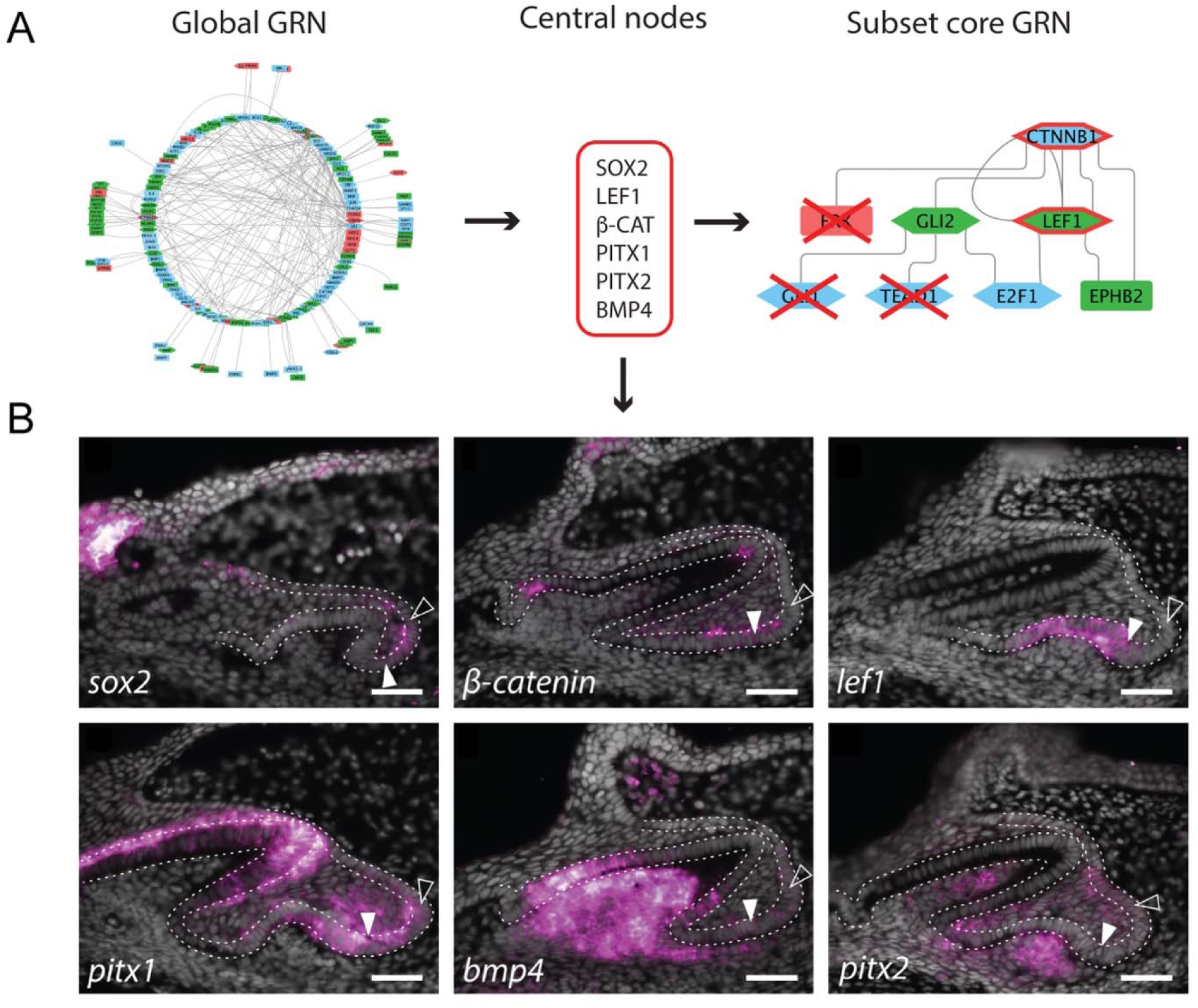
Filtering DEGs for gene regulatory network construction targeting potential candidates for stem cell maintenance and tooth initiation within the shark successional lamina. (A) Global GRN was constructed via CYTOSCAPE based on the initial set of 1282 DEGs. Four super-clusters associated with dental initiation (TTJ/SL, SL, SL/ET, TTJ/SL/ET) were filtered and run through a Regulatory Network Enrichment Analysis (RNEA) with central nodes (*sox2*, *lef1*, β*-cat*, *pitx1*, *pitx2*, and *bmp4*) included. A subset core GRN was then constructed, filtering genes based on minimal connectivity (see methods). (B) In situ hybridization assay of the central node genes, showing expression (false color, magenta; DAPI counterstain, greyscale) within the epithelial cells of the successional (and dental) lamina, at the location of new teeth (white arrowheads) and at sites related to the niche for continued tooth production (black arrowheads).

**Figure 5.**
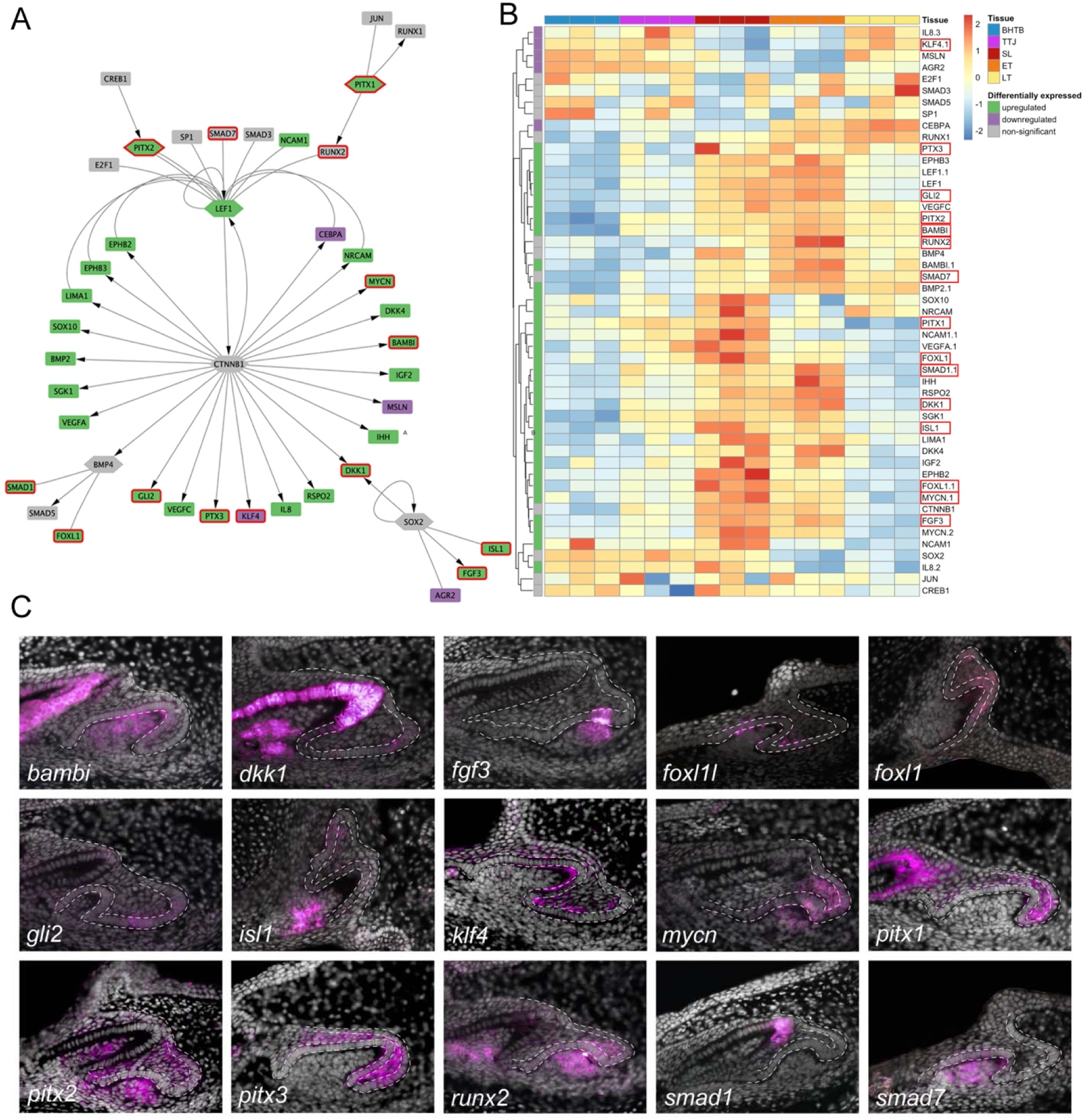
A predictive gene regulatory network (GRN) for the successional lamina and tooth regeneration in sharks. (A) Filtered gene regulatory network with super-cluster genes from the SL tissue compartment showing interactions based on RNEA metadata. Central nodes: β*-catenin* (CTNNB1), *lef1*, *sox2*, *pitx1*, *pitx2*, *bmp4*. Purple colored nodes are downregulated within the SL, while green-colored nodes are upregulated within the SL (see heatmap in B). Non-significant nodes are grey. Red outlined nodes represent the nodes for which in situ hybridization assays are presented (in C). (B) Expression-level heatmap of the genes represented within the GRN in A (red outline genes match those in A). (C) In situ hybridization of representative nodes within the network (red outlines) *bambi*, *dkk1*, *fgf3*, *foxI1L*, *foxI1*, *gli2*, *Isl1*, *klf4*, *mycn*, *pitx1*, *pitx2*, *pitx3*, *runx2*, *smad1*, *smad7*.

### Sub-setting GRNs based on prior knowledge of gene interactions

To construct our successional lamina GRN, we only used DEGs (abs(log2FC) > 0.75 & FDR < 0.001) – however we also include *sox2* and *bmp4* even though they are not significantly differentially expressed within the SL (Figure 4C). We are aware of their importance as central nodes, as they regulated stem cell fate in the SL (Figure 4A and B). *sox2* is expressed throughout the dental lamina, from the TTJ stem niche to the successional lamina, which is why we observed *sox2* within, but not limited to, the SL stem niche (Figure 4B). We characterized the underlying expression of several important known dental genes within our model to confirm their utility in focusing our analysis. Several markers were already well known from the developing shark dentition and specifically expressed in the dental lamina during tooth development and regeneration e.g., *sox2*, *lef1*, β*-catenin (CTNNB1), pitx1*, *pitx2*, and *bmp4* (*5, 16*).

Unsurprisingly, these markers are associated with vertebrate tooth development more generally. These genes became central nodes during the predictive gene network filtration (Figure 4A). Subsequently, we filtered our network by removing genes which are not within two degrees of separation of our central nodes (Figure 4A; subset core GRN). While constructing the GRN (Figure 5) we iteratively removed any markers which interact with only a single other gene, as these genes exhibit low network connectivity and are therefore less likely to play a significant role in our regulatory network.

After we obtained the predicted gene interactions using RNEA (Figure 4A; Global GRN and Full GRN Figure S4), we filtered the network to retain only the genes which interact with predefined ‘central nodes’ (Figure 4A), i.e., genes known to play a crucial role in stem cell regulation within the SL. By treating *lef1, sox2,* β*-catenin, pitx1, pitx2* and *bmp4* as central nodes in the global SL GRN (Figure 4) and retaining markers that interact with at least one of these six nodes, we reduced the number of GRN markers from 150 to 49 (Figure 5A). We identified transcription factor stem cell regulators specifically upregulated in the SL, compared to all other compartments (Figure 3A,B; Figure 4), from both the filtered networks and associated LRTs, these include *mycn* and *foxl1* (Figure 5B and 6A). After identifying putative novel stem cell regulators, their expression was then mapped back on the SOMs previously calculated (Figure 3A). Interestingly, this method revealed that *mycn* is not only identified as a novel regulator in our GRN (Figure 5 and 6), but it is also found within a SL upregulated cluster from the SOM analysis (see Figure 3). Our analysis also revealed markers significantly downregulated in the SL, compared to other compartments, including *cebpa* and *foxq1*, with opposing expression patterns (Figure 5 and 6). *cebpa* is upregulated during compartments undergoing tooth morphogenesis (ET and LT; Figure 5B, 6A and C), and *foxq1* is expressed almost exclusively in taste bud-linked territories i.e., BHTB and TTJ (however, expression of *foxQ1* is expressed again later in tooth development (LT; Figure 5B and 6A and C).

**Figure 6.**
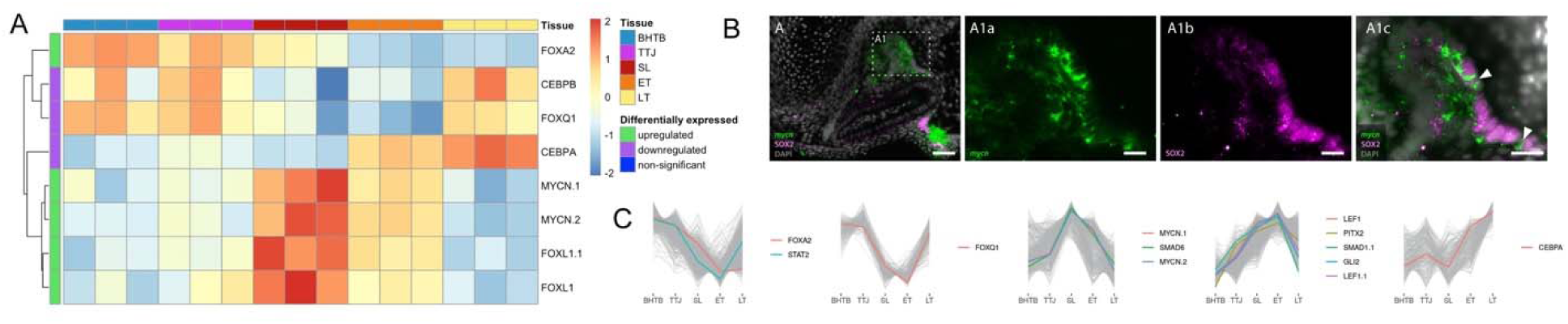
MYCN, a proto-oncogene identified as a key marker of the successional lamina in the shark. (A) Heatmap showing the filtered subset of DE transcription factors, identifying the proto-oncogene *mycn* as a key marker expressed within the successional lamina (red). (B) Expression validation of *mycn* (arrowheads; green) represented by in situ hybridization within the epithelial cells of the successional lamina of the catshark upper jaw (A-A1c); *mycn* in situ hybridization is combined with SOX2 immunohistochemistry – a core gene and known stem marker within the dental lamina. (C) Relative expression levels of transcripts across compartments showing representative markers observed from filtered DEGs of the SL clusters, including those represented in A.

*sox2* is a key marker of stem-potential within the DL, however, not necessarily confined solely to the SL (Figure 4B)(*16*). Simultaneously, canonical Wnt signaling (specifically β*-catenin* and *lef1;* Figure 4B) regulates dental initiation from *sox2+* cells (*15*, *16*). Markers including *bmp4, pitx1* and *pitx2* (Figure 4B) have also been shown to regulate the onset of odontogenesis in first generation teeth (*7*, *32*, *33*), whilst their importance in the regulation of regeneration is less well known. To further validate the methods and design of the GRN analysis we revisited the expression of our core central nodes for the RNEA analysis. We carried out *in situ* hybridization on sagittal thin paraffin sections of late stage 32 embryos (St32L) during which the second dental generation forms (*23*) (Figure 4), and in conjunction with Sox2 protein immunofluorescent staining (Figure 7).

**Figure 7.**
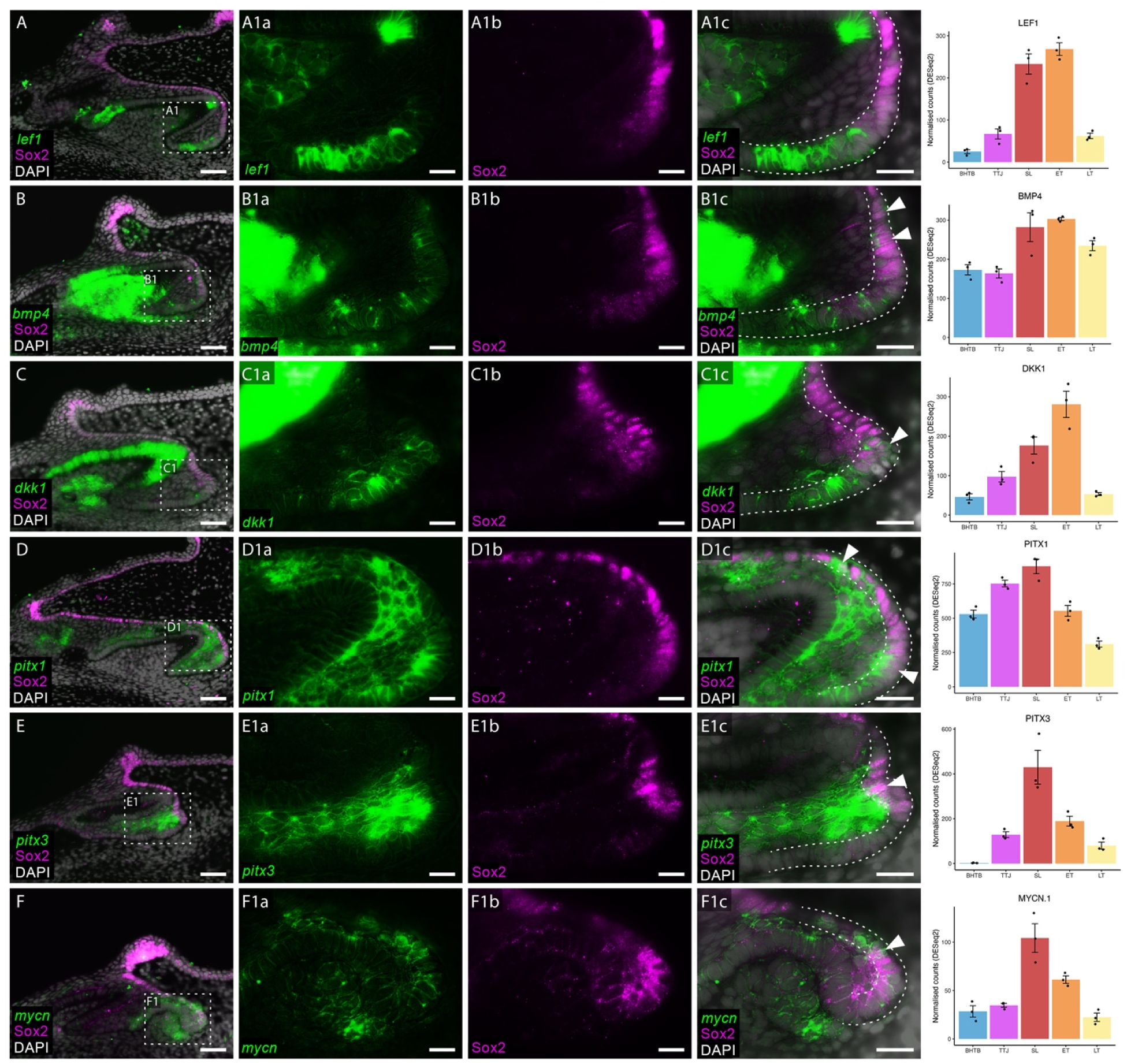
Co-expression of key SL markers in conjunction with the stem cell marker, Sox2. Double section *in situ* hybridization/immunohistochemistry on catshark lower jaws reveals co-expression of *in situ* hybridization markers with Sox2 within the SL. Genes investigated with *in situ* hybridization include *lef1* (A), *bmp4* (B), *dkk1* (C), *pitx1* (D), *pitx3* (E) and *mycn* (F). Sox2 is found within the taste bud on the oral surface and forms a stream of expression throughout the DL (A-F). Sox2 expression stops abruptly at the tip of the SL (A-F1b). *lef1* (A) is expressed within the enamel knot and the new tooth forming region (A1a) but is not co-expressed with Sox2 (A1c). *bmp4* (B) is expressed within the dental mesenchyme and dental epithelium, although it is absent from the enamel knot. It also shows weak expression within the SL (B1a) and is co-expressed within basal epithelial cells of the SL (B1c). *dkk1* is strongly expressed in the dental epithelium and dental mesenchyme and is also expressed within the new tooth forming region (C1a). The limit of its expression sees it co-expressed with Sox2 at their boundary (C1c). *pitx1* (D) and *pitx3* (E) are both expressed within the MDE. The expression of *pitx1* extends further orally and into the basal epithelium of the SL (D1a). Both *pitx1* and *pitx3* are co-expressed with Sox2 within the SL (D1c and E1c). *mycn* (F) is found within the dental mesenchyme and the MDE (F1a). It is also extensively co-expressed with Sox2 within the basal epithelium of the SL (F1c). Gene expression is shown in green, Sox2 protein expression is in magenta and DAPI nuclear stain in greyscale. Dotted white boxes in A-F depict a magnified region in A1-F1. White arrowheads in A1c-F1c highlight regions of co-expression between *in situ* markers and Sox2 in the SL. Scale bars are 50μm in A-F, and 25μm in A1(a-c)-F1(a-c). Transcript normalized counts for each gene are represented across the color-coded tissue compartments; BHTB, blue; TTJ, magenta; SL, red; ET, orange; LT, yellow. Each gene shows higher transcript counts in the SL and ET sub-compartments, relative to others. Bars show mean ± SEM of DESeq2 size-factor–normalized counts; points are individual replicates.

Sox2 is expressed within taste buds at the TTJ on the oral surface and throughout the DL and in the tip of the SL (Fig 7B: white arrowhead). Furthermore, a stream of *sox2*/Sox2 expression connects epithelial cells on the oral surface to the dental progenitor niche within the SL (Figure 4B; Figure 7A-F), with epithelial cell migration known to occur from the oral junction of the tooth and taste territories (TTJ) to the SL (*16*). β*-catenin* is expressed in cells positioned more labially where new dental placodes form (Figure 4B: white arrowhead). This confirms previous work in the catshark, which described the isolation of Sox2/β-catenin, except for during a short period in dental initiation when they are co-expressed (*16*). The Wnt readout factor, *lef1* is absent from the SL, but is highly expressed within the dental epithelium during placode formation (Figure 4B). *pitx1* and *pitx2* are both expressed within the ‘middle’ dental epithelium (a stellate reticulum-like group of stratified squamous epithelial cells (*16*)), and the columnar basal epithelium of the SL (Figure 4B; Figure 7D, E), seemingly within the same region of the SL which expresses *sox2* (Figure 4A, black arrowhead). However, *pitx1* and *pitx2* differ in their expression within the tooth-forming region. *pitx1* is expressed throughout the dental epithelium in both early and late teeth, whereas *pitx2* is primarily restricted to the dental mesenchyme, with low levels of expression within the epithelial compartments of the DL.

The importance of *bmp4* expression during dental morphogenesis is well known, with a role promoting Wnt signalling (*34–36*). The expression of *bmp4* during dental morphogenesis has been found within the lingual dental epithelium in the ferret (*9*), where *sox2* is also expressed (*17*). We find *bmp4* upregulated within the dental mesenchyme and dental epithelium of developing teeth and observed its absence within the cusp-forming enameloid knot. *bmp4* is also expressed within the mesenchyme underlying the dental initiation site (Figure 4B: white arrowhead, Figure 4B), with weak expression within the SL region, housing dental progenitors restricted to basal epithelial cells (Figure 4B: black arrowhead, Figure 7B1a). As described by Jussila et al.(*9*) we also note its co-expression with Sox2 within the SL, albeit within only a small number of cells (Figure 7B1c).

### Expression of predictive successional lamina GRN markers

We focused our investigation on the transcriptional signature of this important SL compartment to identify putative regulators of this stem/progenitor cell niche directly involved in the regulation of tooth regeneration (*16*). Representative genes associated with the SL tissue clusters and DEGs were validated via *in situ* hybridization assays (Figure 5); from this list, notable genes include *bambi, dkk1, fgf3, foxL1, gli2, isl1, klf4, mycn* (see later descriptions), *pitx1, pitx2, pitx3, smad1, smad7* and *runx2*.

From the wider analysis of the upregulated genes from the SL compartment (see Supplemental Table S1), WNT pathway genes were ever-present. Our analysis identified *wnt11* and *sfrp5* upregulated in the SL, which is key given the known role of canonical Wnt signaling in regulating dental initiation from the cycling stem cell niches. Furthermore “canonical WNT pathway signaling” was a key GO term covered by the SL compartment (Figure 3D). Additionally, we find *pitx3* upregulated in the SL compared to all other tissue compartments. Members of the Pituitary homeobox family (Pitx; see phylogenetic analyses: Figure S2) of transcription factors are well-known to be involved in multiple stages of dental development. *pitx1* and *pitx2* are both expressed in the oral epithelium prior to the invagination of the DL in the catshark (*5*, *37*). As a result of its early expression within the odontogenic band prior to dental initiation, *pitx2* has been described as an odontogenic-commissioning gene. Within our RNAseq dataset, we find *pitx1, pitx2* and *pitx3* differentially expressed within SL-related super-clusters (SL; TTJ/SL/ET; and TTJ/SL/ET respectively). This is the first report of *pitx3* expression throughout the entire odontogenic program (Figure 3; Figure 5B and Figure 7), although negative expression had previously been reported in the catshark dentition (*37*). *pitx1* is upregulated in the TTJ and SL, before being downregulated in the ET (Figure 4). In contrast, *pitx3* is dramatically upregulated (>2fold relative to any other tissue) in the SL before being downregulated in the ET and LT (Figure 3 and 5). Both *pitx1* and *pitx3* are expressed within the middle dental epithelium of the SL (Figure 7D, D1a, E and E1a). *pitx1* extends orally throughout the epithelium, towards the TTJ, reflecting its expression within the RNAseq analysis (Figure 7). We suggest that all three vertebrate Pitx family genes are important to the evolution, development and regeneration of the vertebrate dentition. Further work on this family of transcription factors, beyond Pitx2, will be necessary to appreciate the extent to their role in shark tooth regeneration and beyond, across the vertebrate phylogeny.

A readout of canonical Wnt signaling, *lef1* (Figure 3, 4, and 7) is expressed within the dental epithelium of bud stage teeth. *islet1* (*isl1*) shows weak expression within the teeth; its expression can be seen in both the dental epithelium and within the SL (Figure 5). Dickkopf (*dkk*) is a potent antagonist of the Wnt signaling pathway, with its ectopic expression leading to a complete loss of hair, tooth and mammary gland development in mice (*38*). During catshark tooth development (e.g., present in both the SL network and ET compartment SOM clusters; Figure 3), we observe *dkk1* strongly expressed throughout the dental epithelium and dental papilla, as well as weakly expressed within the SL (Figure 5C). Given that canonical Wnt signaling is critical in the regulation of epithelial appendage initiation, Dkk1 is a prime target deserving of further investigation. Intriguingly, *dkk1* is found in a similar pattern within the dental initiation site (Figure 7C1a). However, it is also expressed within the distal most Sox2+ cells of the SL (Figure 7C1c). Given its role as a canonical Wnt signaling antagonist, *dkk1* may be working in conjunction with other Wnt signals in mediating the activation of dental progenitors during repeated dental initiation in sharks. *smad1, smad7, bambi, fgf3, msx2* and *wnt11* (Figures 3 and 5) were upregulated within the developing teeth, and not as highly expressed within the SL, all but *wnt11* were found within either TTJ/SL/ET or SL/ET super-clusters (Figure 3A-C). These represent genes where expression is activated in the SL and continues to rise in the early developing teeth (ET; Figure 3).

Two *foxl1* transcripts were initially found differentially expressed within the SL, however phylogenetic analysis identifies these markers as *foxl1* and *foxl1-like* (Figure S3). *In situ* hybridization for both markers (Figure 5 and 6) reveal subtle differences in their expression patterns in the teeth. Both markers are weakly expressed within the dental epithelium, however *foxl1* is also found within the dental papilla. Furthermore, they are both expressed within the epithelium at the SL dental initiation site. *Foxl1* has been identified as a direct target of Hh signaling, with binding sites for the Hh family member Gli2, found within Foxl1 non-coding sequences throughout vertebrates (*39*). Concordant with *foxl1* expression, we find *gli2 (and gli3)* within the epithelium of the SL dental initiation site, whilst its expression also extends within the greater SL and during early tooth initiation (Figure 5C), and later in mesenchymal compartments of the developing tooth. Interactions between *gli2* and *foxl1* may be regulating the proliferation of the dental epithelium, as is the case in intestinal villi (*39*).

Runt-related transcription factor-2 (*runx2*) is expressed in the dental papilla during morphogenesis and dental mesenchyme in bell stage teeth (Fig 5C). Its expression is thought to regulate dental development prior to bell-stage, with its downregulation in later stages regulating odontoblastic differentiation (*40*). We also note *runx2* expression within the stratified epithelial cells of the middle dental epithelium, whilst it is absent from the distal most basal epithelial cells of the SL, corresponding with the Sox2+ progenitor cell niche (*16*); the exact role of *runx2* in the SL progenitor niche requires further investigation. *runx2* is implicated in the regulation of cell cycle progression, and control of osteogenic progenitor cell proliferation (*41*, *42*); therefore a potentially similar role for *runx2* in the SL could be investigated, given the conservation of the associated network genes are among those expressed in this specific dental compartment. Interestingly, we find all members of the Runx family of transcription factors expressed in the early tooth (ET) compartment (*runx1*, cluster 17; *runx2*, cluster 16; and *runx3*, cluster 21: Figure 3). The three Mammalian Runx genes have long been associated with tooth and craniofacial development (*43*) and we can confirm the association of all Runt-related transcription factors within the developing early tooth compartments of the shark dental lamina.

### Expression of SL markers in conjunction with the stem marker, Sox2

Given the expression of *pitx1, lef1, bmp4, dkk1* and *mycn* within the SL (Figure 7), we further investigated their roles during dental regeneration. These markers were chosen because of (*i*) previous research on their involvement in dental initiation; (*ii*) their expression patterns within the SL; and/or (*iii*) their known role in canonical Wnt signaling. Furthermore, *pitx3* was also included in this analysis, as it was more than 2-fold upregulated in the SL transcriptome than any other tissue (Figure 3 and 7), and given the importance of both *pitx1* and *pitx2* during the onset of odontogenesis (*7*, *32*, *33*) we find all these markers upregulated within the SL and/or ET relative to other dental tissues (Figure 3,5 and 7). To characterize their expression within the catshark SL, we carried out double sagittal section *in situ* hybridization and immunohistochemistry revealing their co-expression with Sox2 (Figure 7).

Sox2 is expressed within taste buds at the TTJ on the oral surface and throughout the DL. A stream of Sox2+ cells connects epithelial cells on the oral surface to the dental progenitor niche within the SL (Figure 7A-F), with epithelial cell migration known to occur from the TTJ to the SL (*16*). *lef1* can be seen expressed within the dental initiation site (Figure 7A1a) but is notably absent within Sox2+ cells of the SL (Figure 7A1c). There is an abrupt expression boundary between *lef1* and Sox2 (Figure 7A1c), suggesting a role for canonical Wnt signaling in dental initiation outside of Sox2+ dental progenitors.

We also identify co-expression of the novel dental marker *mycn* in Sox2+ SL cells (Figure 3, 5, 64AB(A1c) and 7F1c). Unlike other markers studied (Figure 5), we also note *mycn* upregulation within the lingual DL, which connects the SL to the oral surface, and within Sox2+ taste bud cells at the TTJ (Figure 6). *bmp4* is expressed throughout a range of dental tissues during dental development. We observe its expression strongly upregulated within the dental papilla, and throughout the dental epithelium –although absent from the enameloid knot (Figure 4B and 7B;(*44*)). Its mesenchymal expression extends below and surrounds the epithelial dental initiation site. Within the SL, *bmp4* is restricted to basal epithelial cells (Figure 7B1a). As described by Jussila et al. (*9*), we also note its co-expression with Sox2 within the SL, albeit within only a few cells (Figure 7B1c). *pitx1* is expressed throughout the basal epithelial cells of the SL, co-expressed with Sox2+ cells (Figure 7D1c). The extent of *pitx3*/Sox2 co-expression is more restricted, although there is still observable co-expression at the distal tip of the SL (Figure 7E1c). The expression of Pitx genes within the SL, and their co-expression with Sox2+ cells, highlights a potentially important role for this pathway in, not only dental initiation and development, but also repeated dental regeneration. These findings suggest that some of the core elements of the tooth initiation program might be cyclically redeployed and necessary for the continuation of next-generation tooth production, at least in sharks.

### mycn (n-myc): a potential regulator of SL progenitor maintenance, proliferation and dental lamina morphology

The bHLH transcription factor *mycn* (N-Myc Proto-oncogene) has been well described in the maintenance of neural progenitors (*45*). Mycn is a proto-oncogene which functions to maintain cells in an undifferentiated state and has been identified as critical for maintenance of neural crest stem cells (*46*). We asked whether it could be fulfilling a similar function in the undifferentiated SL progenitors. We observe both *mycn* and *myc* in the filtered DEGs specifically within the successional lamina clusters; this is interesting as both markers are known regulators of normal embryogenesis and important for the maintenance of stem cell properties (*47*). *mycn* is highly upregulated in the SL clusters (Figure 3A) and we find that it is expressed within the distal epithelial compartment of the DL and within the distal-most underlying dental mesenchymal cells of early bud-stage teeth (Figure 5, 6 and 7). It is also expressed throughout the SL (Figure 3C; Figure 5 and 6). *mycn* is co-expressed near or in Sox2+ SL cells (Figure 7F1c and 6B(A1c)). Unlike other genes studied (Figure 5), we also noted *mycn* upregulation within the lingual DL which connects the SL to the oral surface, and within Sox2+ taste bud cells at the TTJ (Figure 7). The *mycn* gene is commonly amplified in neuroblastoma, tumors which originate from undifferentiated neural crest cells (*48*). Given the pivotal role this gene has in this pathogenesis, it has been an attractive target for drug development. *10074-G5* (*G5*) and 10058-*F4* (*F4*) are small molecule inhibitors which disrupt MYCN/n-myc dimerization with MAX, essential for its role in transcriptional regulation (c-MYC / MAX dimerization is also blocked; (*49*)). We therefore treated whole shark embryos with these inhibitors, at the stage (100dpf) prior to tooth formation and during the early establishment of the DL.

Embryos were treated with 0.5μM and 2μM (G5) and 0.5μM and 2μM, 5μM and 10 μM concentrations of these pharmacological agents in seawater for 14 days, or with a DMSO vehicle control (Figure 8; Table 1). *mycn* and the other drug target *c-myc* have pleiotropic effects throughout development. 2μM *G5* proved lethal. However, samples were obtained from 0.5μM-*G5* and –*F4* treated sharks and 2μM-*F4* treated specimens. Tissue thin sections were processed through immunofluorescence with DAPI, PCNA and PH3 to assess lamina morphology and proliferation. PCNA, the DNA sliding clamp important in DNA synthesis, was expressed throughout the lamina and areas of adjacent mesenchyme associated with early tooth formation (Figure 8 and Figure S5). PH3 which strongly stains the condensed chromatin in cells in late G2/mitosis was quantified in sections. PH3/DAPI was quantified within the shark dental lamina; 2-way ANOVA revealed a significant effect of treatment (p= 0.0219), and also a significant difference between individual animals (p<0.0001). Testing for differences in treatment versus DMSO control with Dunnett’s post hoc test for multiple comparisons confirmed a significant reduction in proliferation in both the *F4* 2μM (p adj. = 0.0234) and a trend toward significance (p adj. = 0.1136) in the F4 0.5μM treated samples (Figure 8C). We investigated the effect upon the dental lamina when we increased the dosage of the drug to 10μM. At this concentration of *F4*, in particular (*G5* was lethal at this concentration), we observe modifications of the dental lamina, successional lamina and developing teeth, suggesting that our drug treatments target the myc-max interaction within the shark dental epithelial stem cells (Figure S5). Further work will be necessary to understand the disruption and role of the signaling in the neighboring mesenchymal populations that normally express myc/n-myc. Our discovery of *mycn* as a potential regulator of proliferation within the shark SL stem niche is significant. It reveals the potential for the discovery of novel and important markers, beyond *sox2*, that are likely involved in maintenance of perpetual tooth regeneration in sharks that also could be essential for the evolution of tooth regenerative mechanisms in vertebrates. Given the well-documented role for Mycn in stem self-renewal and proliferation, albeit during neurogenesis, we therefore suggest that Mycn (N-myc) and the MYC family of regulator genes could be part of the cascade of genes necessary for the maintenance of a progenitor cell state in the SL and regulation of tooth renewal in chondrichthyan fishes.

**Figure 8.**
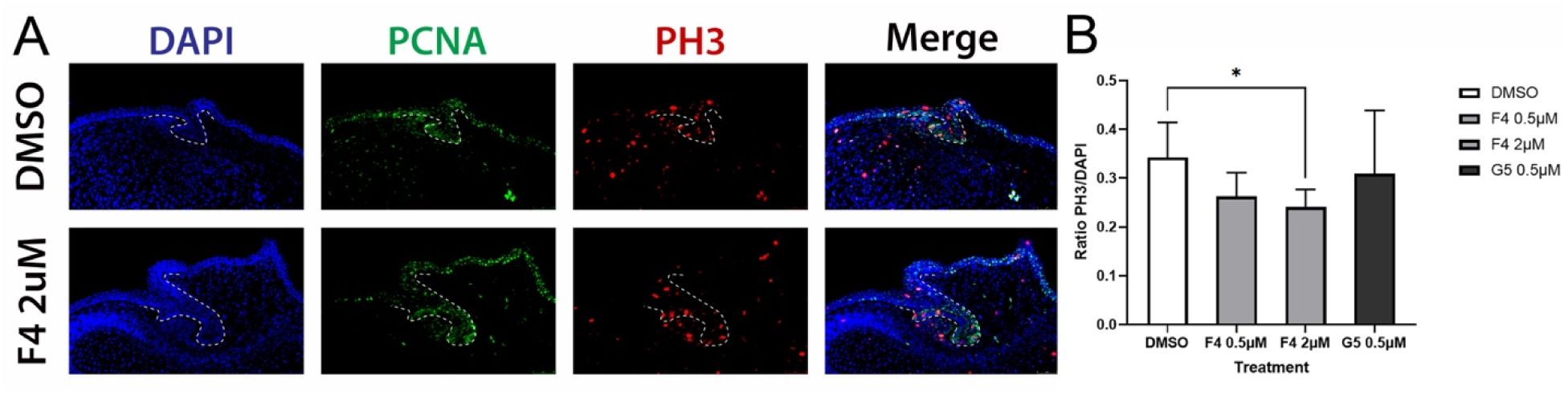
Pharmacological perturbation of MYC/MYCN in the developing dental lamina disturbs normal levels of cell proliferation. (A) following MYC/N inhibition the morphology of the dental lamina, successional lamina (arrow) and the TTJ (arrowhead) compartment shows altered morphological differences in development compared to the DMSO control. (B) Cell proliferation (PCNA; PH3) visualization shows marked changes in cell proliferation after MYCN inhibition, with F4 drug at 2μM concentration. (C) Reduced cell proliferation was apparent via cell counts particularly in the F4 treatment assay at 2μM (two-way ANOVA with Dunnett’s multiple comparison test).

**Table 1.** Replication numbers and treatment concentrations for the MYC inhibition assays. DMSO (0.1%) was used as a delivery control, and all treatment agents included DMSO.

| Treatment (pharmacological agent) | Concentration (μM) | Replicates |
| --- | --- | --- |
| 10074-G5 | 0.5 | 5 |
| 10058-F4 | 0.5 | 4 |
| 10058-F4 | 2 | 5 |
| 10058-F4 | 5 | 2 |
| 10058-F4 | 10 | 2 |
| DMSO (control) | 0.1% | 6 |

## Discussion

### Renewal and maintenance of pluripotency in the shark successional lamina

Sharks, and likely the vast majority of polyphyodont vertebrates, can maintain a typically embryonic epithelial tissue, the dental lamina (DL), throughout their lives. Our transcriptomic description of the DL expands our knowledge of this highly robust dental tissue, uncovering how it forms and how it is maintained. Importantly, this work paves the way for a clearer understanding of the mechanisms of embryonic tissue maintenance and why some vertebrates (i.e., mammals) have lost the ability to (*i*) preserve and maintain embryonic dental organogenesis, and (*ii*) recapitulate embryonic odontogenesis to produce unlimited or multiple generations of teeth.

While we know little about the maintenance of continuously regenerative and sequential organ systems, the shark dentition offers a unique model that can further uncover the genetic regulation of stem/progenitor mediated perpetual tooth regeneration. The conserved nature of the DL tissues and corresponding genetic networks that maintain them, suggests that studying shark tooth regeneration and progenitor cell maintenance has implications for other dental systems, including the human dentition (*50*). Several well-known stem cell markers are expressed in the shark DL that might be crucial for (i) the maintenance of embryonic epithelial stem cell niches, and (ii) growth and development of the DL, and (iii) continuous regeneration of the teeth. Loss of stem/progenitor maintenance factors with the SL may have prevented the continuation of polyphyodontism during the evolution of more restricted dentitions i.e., mono– and diphyodontism in mammals. Thus, it will be intriguing to understand the expression of these genes during developmental stage series in models with more restrictive dental generations (monophyodont and diphyodont animals).

Our *de novo* transcriptome assembly of an emerging polyphyodont model, which can serve as a reference point for further dental developmental and regenerative research. Our results show that predictive genetic regulatory networks (GRNs) serve as useful tools to identify key markers of interest from extensive RNAseq datasets. Through utilizing previous knowledge to subset a global reference GRN, we generate a sub-network, which contains multiple markers expressed within the SL. These markers include *runx2, gli2, isl1* and *foxl1*. Furthermore, this analysis led to the identification of the proto-oncogene, *mycn*, which we find expressed within Sox2+ dental progenitors for the first time. Previous polyphyodont research has been primarily based on the candidate approach for the study of successional dental regeneration. This has inevitably relied heavily on research in mammals, which do not possess lifelong successional whole tooth regeneration. Nevertheless, this approach has vastly improved our understanding of dental development and regeneration. Lineage tracing experiments revealed that Sox2+ cells in the adult originate from Sox2+ cells in the embryo (*51*). Although it had been previously found within taste buds (*52*), the first description of its expression in teeth came from the cervical loop stem cells of the continuously growing mouse incisor (*18*). Since then, Sox2 has been identified within dental stem cells of all studied polyphyodont vertebrates, including reptiles (*15*, *17*), teleosts (*14*) and sharks (*16*). Sox2 has now become a hallmark of dental regenerative potential and has revealed an ancient link between teeth and taste buds in the evolution of polyphyodonty (*16*).

The importance of the candidate approach cannot be understated. However, our results demonstrate the efficacy of taking a transcriptome-based approach in an emerging polyphyodont model in order to reveal novel markers expressed during successional and life-long regeneration. We describe the expression of *mycn* during dental regeneration for the first time. Its role in neural development has been extensively studied and is known to regulate proliferation and the cell cycle in neural progenitors (*45*, *46*). It is a target of canonical Wnt signaling, with its downregulation leading to differentiation of neuroblastoma cells (*53*, *54*). We identify its expression within Sox2+ taste bud cells at the TTJ (Figure 6 and 7) and notably within Sox2+ SL cells (Figure 7F1c and 6BA1c). The extensive co-expression of *mycn* with Sox2 in the SL together with its role in neuroblastoma formation, implicates *mycn* in the cell cycle regulation of dental progenitors and the onset of odontogenic differentiation.

Models aimed at recreating GRNs from gene expression data, have focused on inferring networks solely using gene expression stochasticity or changes in expression under different conditions (e.g. ARACNE (*55*); GENIE3 (*56*); NARROMI (*57*); LBN (*58*)). These models invariably lead to a number of false positives as they predict interactions based on co-correlated genes (*59*). As a result, networks derived from these models require experimental validation of predicted interactions. In this study, we identify *mycn* within our differentially expressed SL data set through predictive GRN reconstruction using RNEA and reveal its co-expression with Sox2 (Figure 7F). We use prior system knowledge to simplify our initial network to a key list of testable genes. *mycn*, for example, was not two-fold upregulated within the SL, nor was it labelled a key marker of interest following our GO enrichment analysis, revealing the importance of GRN reconstruction in this case. Although the goal is to both identify and understand interactions between genes during successional dental regeneration, we show that predictive GRNs can serve as a key tool in identifying novel candidates from transcriptome datasets. Our predictive model therefore presents a testable framework to uncover the functional relationships of these genes for the purpose of unlimited tooth production and provides insights regarding how this program is lost in mammalian vertebrates. Furthermore, our predictive successional lamina GRN is an important testable model to include in the conversation about dental gene network evolution.

We can now expand our understanding of odontode development, regeneration and the presumptive core network that may have existed at the dawn of tooth evolution (*60*). Importantly in this context, these two processes of development and regeneration are interlinked where tooth regeneration is a direct extension of DL development. Homeostasis regulates the continuation of tooth regeneration directly from an embryonic epithelial tissue – the successional lamina. Furthermore, the same genes that establish the first teeth are reused for the cyclical production of all tooth generations. However, we predict that separate but integrated networks must operate (i) to produce a unit odontode (tooth or denticle) and (ii) maintain the dental lamina and resident progenitor cells to initiate new tooth generations at cyclical intervals. It is possible that one core network originally regulated both processes and subsequently separated over time to allow the reduction of tooth generations, in some cases producing just a single generation e.g., the murine dentition. We hypothesize that although our shark model (*Scyliorhinus canicula*) is a representative of more derived chondrichthyan clades, the patterning of the dentition via the dental lamina might represent an evolutionary relic dental character leading to separate tooth units, interlinked and renewed in a cyclical and continuous manner (a tooth family/whorl), retained in modern shark lineages and toothed vertebrates more generally The presence or absence of the dental lamina in all tooth-forming gnathostomes has been a point of debate and it has been suggested (*21*, *61*) that this structure is not essential for tooth regeneration in some groups, e.g., in non-teleost actinopterygians.

However, we posit that tooth regeneration at all levels must be regulated by an active dental lamina, whether permanent or temporary (*3*, *62*). Even reports of tooth replacement from the expanded outer dental epithelium of teleosts and non-teleost fishes must be considered a modification or reduction (not loss) of the dental lamina from the initial competent field of dental epithelia. This information highlights the diversity of the dental arcade and the regenerative capacity in early diverging clades and gnathostomes more generally. Characterizing tooth regeneration from soft tissues in fossil specimens present challenges (*63*), however the conservation of the tooth regeneration mechanism from an active dental lamina or reduced lamina is plausible in all lineages. Our data suggests that the general tooth network and especially the developmental/regenerative gene network offers insights into a highly conserved, evolutionarily robust and well-preserved gene network. Thus, tinkering of this ‘stable’ network for over 400 million years, has therefore led to the vast diversity of dental form and regenerative capacity.

### Pitx homeobox family of genes as key markers of tooth regeneration

Aside from predictive GRN analysis, our differential expression analysis also identified the co-expression of a new marker within the SL. Two members of the Pitx family, Pitx1 and Pitx2, have both been implicated in the onset of odontogenesis (*7*, *32*, *33*), and related to shark odontogenesis (*5*, *37*).

Although *pitx3* has been described during molar morphogenesis (*64*), and aside from negative expression reported in the shark dentition (*37*), its expression during successional regeneration has not previously been described. We, however, find *pitx3* more than two-fold upregulated in the SL relative to any other tissue, and note its strong expression in situ, specifically within the SL (Figure 7EC). The expression of both *pitx1* and *pitx3* within Sox2+ dental progenitors emphasizes a potential key role for the Pitx gene family, not only in the regulation of the first dental generation (*7*, *32*), but also successional tooth regeneration. More generally, the Pitx family of genes have been implicated in several other regenerative systems, including murine satellite cells (*65*), and as one of the first families of genes expressed in the initiation of the dentition within vertebrates, it seems to correlate that all Pitx genes are implicated in the continued production of teeth in the SL of the shark. Further research will elucidate the complete role of this family of transcription factors in the maintenance of a polyphyodont dentition for over 400 million years of tooth evolution.

## Conservation of dental and stem markers across vertebrates

The tooth development program in extant vertebrates is now well-known and the genes expressed during this process of organogenesis appear highly conserved among vertebrate clades, from fish to mammals. However, this has mostly relied on a subset of well-studied signaling molecules and transcription factors, first described during the development of the mouse molar and incisor models (*17*, *33*, *66–69*). There is now a wealth of dental transcriptomic data from many vertebrate systems (*28*, *70–74*) with varying degrees of tooth regenerative capacity, some of which have characterized the dental transcriptome at the level of the single cell (*75*, *76*). This provides an enormous coverage of data available for comparison across vertebrates, including both bulk RNAseq of specific tissues or single cell sequencing. These data sets, together with functional readouts of the pathways and gene regulatory network analysis, provide a framework to uncover the network elements lost and gained over evolutionary time and how to manipulate and enhance this developmental program. The discovery of novel potential regulators of tooth regeneration and dental lamina maintenance in the shark i.e., MYCN, could pave the way for important avenues of future comparative research.

## Conclusion

Sharks are masters of tooth regeneration. We have taken advantage of this non-traditional model system to uncover the genetic signature of cyclical regeneration and unlimited tooth production. Understanding how vertebrates continuously regenerate their teeth via a highly conserved epithelial tissue, the dental lamina, is an important contribution to developmental biology with wider biomedical implications.

Humans make only two sets of teeth from a successional dental lamina; the termination of this productivity is due, in part, to a break-down of the dental lamina (*10*, *50*). The shark’s ability for lifelong dental regeneration is an intriguing prospect for further investigation, allowing us to learn the components necessary for natural whole tooth production and importantly maintenance of the dental lamina beyond two generations. Despite a lack of information on gene interactions during dental regeneration, we find that predictive gene regulatory network (GRN) reconstructions, used alongside more traditional transcriptome approaches, serve as a useful tool for filtering important candidates. This sets the groundwork for further in-depth study of gene interactions that regulate cyclical dental initiation and the diversity of vertebrate dental regeneration. Furthermore, the shark successional lamina provides an accessible system for the study of regenerative processes, and mechanisms of both controlled and aberrant tissue growth, active for the entirety of life. Comparable vertebrate tissues or structures are either rare or difficult to access across life stages, e.g., cancerous and non-malignant tumors (*50*) and intestinal epithelia. However, we show that the shark dental lamina and its lifelong stem cell niche has a genetic signature with overlapping elements known from a range of developmental, maintained, and tumorigenic tissues. This genetic conservation offers important avenues for translational research that will shed light on the conserved cellular dynamics and genetic mechanisms for the renewal and controlled growth of crucial tissues beyond embryogenesis.

## Materials and Methods

### Animal husbandry

Shark embryos were housed at The University of Sheffield and the University of Florida. The University of Sheffield is a licensed establishment under the Animals (Scientific Procedures) Act 1986. All animals were culled by approved methods cited under Schedule 1 to the Act. Catshark embryos (*Scyliorhinus canicula*) were obtained from North Wales Biologicals, Bangor, UK. Embryos were raised in filtered artificial seawater (Instant Ocean) at 12-16°C. At the required stage, embryos were anaesthetized using 300mg/L MS222 and fixed overnight in 4% paraformaldehyde at 4°C. Samples were then dehydrated through a graded series of DEPC-PBS/EtOH and kept at –20°C.

### Paraffin sectioning and histology

Following dehydration, samples were cleared with xylene or histoclear and embedded in paraffin. 14µm sagittal sections were obtained using a Leica RM2145 microtome. For histological study, sections were stained with 50% Haematoxylin Gill no.3 and Eosin Y. Slides were mounted with Fluoromount (Sigma) and imaged using a BX51 Olympus compound microscope.

### Micro-computed tomography (Micro-CT)

Catshark embryos were stained with 0.1% PTA (phosphotungstic acid) in 70% EtOH for 3 days to enhance contrast of soft tissues. Samples were imaged using an Xradia Micro-XCT scanner (Imaging and Analysis Centre, Natural History Museum, London). Volume rendering of the CT scans was carried out using the Avizo Lite software (Thermo Scientific) and in VgStudio Max. Manual segmentation of the DL in the lower jaw was carried out in order to visualize the DL three dimensionally.

### Sample collection, RNAseq, transcriptome assembly

Five dental sub-regions were dissected from stage 34 (hatchling) catshark samples. The dental compartments included: the basi-hyal taste buds (BHTB); the taste-tooth junction (TTJ); the successional lamina (SL); the early developing tooth (ET); and the late stage developing tooth (LT). Tissue samples were flash frozen in liquid N_2_ and added to TRIzol Reagent (Invitrogen) and homogenized in a tissue lyser. RNA was extracted using phenol/chloroform phase separation and further purified using the Qiagen RNeasy kit. RNA samples were sent for Illumina next-generation sequencing by the Sheffield Diagnostic Genetics Service at the Sheffield Children’s Hospital. Three libraries were sequenced for each dental sub-compartment, with each library containing RNA from three pooled samples. Given the lack of a reference genome, a de-novo transcriptome was assembled using TRINITY (*77*). FPKM (Fragments Per Kilobase of transcript per Million mapped reads) values were normalized to generate TMM (trimmed mean of M values) in order to compare transcript relative abundance between tissue samples (*78*). Predicted TRINITY ‘genes’ were annotated based on their closest UniProt BLAST match. Read quality was assessed using Fastqc v0.11.8 and multiqc v1.9. Raw reads were filtered and trimmed using prinseq-lite by quality score (>20), length (>50), sequence complexity and duplication levels using the following parameters [-min_qual_mean 20 –min_len 50 – min_qual_mean 20 –ns_max_p 5 –trim_qual_left 20 –trim_qual_right 20 – trim_ns_left 1 –trim_ns_right 1 –trim_qual_type min – trim_qual_rule lt – trim_tail_left 10 –trim_tail_right 10 –lc_method dust –lc_threshold 7 – out_format 3]. Trinity v2.8.4 (Grabherr et al. 2011) was used to assemble all fifteen libraries into two reference assemblies: *De novo* assembly – raw reads were assembled using default parameters. Genome-guided assembly – raw reads were aligned to the cloudy catshark genome (Hara et al. 2018) using bowtie2 v2.3.4.3 and then assembled using default parameters. Chimeric fragments and locally misassembled transcripts were filtered from both assemblies using Transrate v1.0.3 (Smith-Unna et al. 2016), where ‘good transcripts were retained, followed by detonate v1.11 (Li et al. 2014) with the bowtie2 option and transcript length parameters calculated using the cloudy catshark genome (Hara et al. 2018), where transcripts scoring < 0 were discarded. Transcripts of each assembly were clustered using cd-hit-est version 4.8.1 (Fu et al. 2012; Li and Godzik 2006) at an identity threshold of 95% (-c 0.95 –n 8 –g 1), and the representative sequence of each cluster was retained. The filtered transcripts of both assemblies were annotated using the Trinotate pipeline v3.1.1 with the following software versions: transdecoder v5.5.0, blast 2.8.1+, hmmer v3.2.1, tmhmm v2.0c, and signalp v4.1. Transcripts with bacterial identities were discarded from the assemblies producing two final transcript sets: Scan_de_novo_filtered.fasta and Scan_genome_guided.fasta. Read data were aligned to the transcriptomes using Trinity’s <u>align_and_estimate_abundance.pl</u> and <u>abundance_estimates_to_matrix.pl</u> scripts with the bowtie2 and RSEM options. Downstream analysis in R was carried out on the de-novo transcriptome counts.

The transcriptome data have been deposited in NCBI GEO with accession GSE198580. Data related to the downstream RNAseq analysis and figures produced in this manuscript can be located at the following GitHub page: https://github.com/alexthiery/s.canicula

### Differential expression analysis

Trinity gene counts were rounded (due to fractional counts), and genes with ≤ 5 reads in ≤ 2 samples were discarded. Counts were analyzed in DESeq2 v1.26.0: size-factor normalization (median-of-ratios) and rlog transformation were applied. A likelihood-ratio test compared the full model ‘∼ 0 + Tissue + Samplè with the reduced model ‘∼ 0 + Sampl’ to identify genes whose expression differed across tissues, yielding 3275 significant genes (FDR < 0.0001).

### Self-organizing maps (SOM)

The 3275 differentially expressed genes were z-score scaled (row-wise) and clustered on a 5 × 5 hexagonal grid using kohonen v3.0.10. Each SOM unit was assigned to the tissue showing the highest median expression, and neighboring units with the same dominant tissue were merged into tissue “super-clusters” (Fig. 3A).

### GO enrichment analysis

For each tissue super-cluster, we ran a one-tailed hypergeometric test in GOstats / GSEABase against the Biological Process ontology, using all GO-annotated transcripts in the RNA-seq dataset as background. Raw P-values were Benjamini–Hochberg–adjusted within each tissue; terms with FDR < 0.05 were considered significant. Enriched terms were manually curated for visualization in Fig. 3D.

### RNEA gene interaction analysis

To predict regulatory interactions in the successional lamina (SL), we carried out pairwise differential expression analysis comparing SL with all other tissues (Figure S7). Genes significantly regulated in SL (absolute log2FC > 0.75, FDR < 0.001) plus the known regulators sox2 and bmp4 which are not specifically differentially expressed in the SL, were submitted to Regulatory Network Enrichment Analysis (RNEA; Chouvardas et al., 2016). RNEA queries GO, KEGG, TarBase, TRED, TRRUST, TFactS and ORegAnno to infer transcription-factor/target relationships and returns a network that we visualized in Cytoscape 3.5.1. In order to determine a refined gene regulatory network (GRN), we used prior knowledge of gene expression within the catshark SL in order to filter the RNEA generated predictive GRN. We treated six markers (*ctnnb1/*β*-catenin, pitx1, pitx2, lef1, sox2 and bmp4*) as central nodes in the network. Nodes which interact directly with any of these six central nodes were retained (Figure 4C and D).

### RNAseq sequence of interest identity check

ORFs for RNAseq sequences were predicted using Geneious 10.0.5 and compared to *Callorhinchus milii*, *Latimeria chalumnae* and *Lepisosteus oculatus* genome annotations. In order to verify the identity of the RNAseq sequences of interest, protein coding sequences (CDS) were extracted for the gene of interest and closest sister genes, from ensembl.org. Sequences were extracted from a range of species. Species were chosen based on their phylogenetic position and included: *Anolis carolinensis, Ciona intestinalis, Danio rerio, Gallus gallus, Gasterosteus aculeatus, Latimeria chalumnae, Lepisosteus oculatus, Mus musculus, Oreochromis niloticus, Oryzias latipes, Pan troglodytes, Pelodiscus sinensis, Petromyzon marinus, Rattus norvegicus, Taeniopygia guttata, Takifugu rubripes* and *Xenopus tropicalis*. RNAseq and ENSEMBL sequences were aligned using MUSCLE (*79*). A maximum likelihood tree was generated from 100 bootstrap replications using PHYML (*80*) with an LG substitution model. Resulting gene trees were checked to see if the gene of interest falls within the expected node.

### Riboprobe synthesis

*Scyliorhinus canicula* total RNA was extracted using phenol/chloroform phase separation and cleaned through EtOH/LiCL precipitation. RT-cDNA was made using the RETROscript 1710 kit (Ambion).

Probes were made using forward and reverse primers designed through Primer3. Primer sequences are available in the supplementary information (Fig S7). Sequences of interest were amplified from the cDNA through PCR and ligated into the pGEM-T-Easy vector (Promega). Ligation products were cloned into JM109 cells. Plasmid DNA was then extracted from chosen colonies using a Qiaprep spin Mini-prep kit (Qiagen) and sequenced (Applied Biosystems’ 3730 DNA Analyser) through the Core Genomics Facility, University of Sheffield. Verified vectors were then amplified through PCR and used as a template for probe synthesis. Sense and anti-sense probes were made using a Riboprobe Systems kit (Promega) and SP6/T7 polymerases (Promega). Probes were labelled with Digoxigenin-11-UTP (Roche) for detection during *in situ* hybridization. A final EtOH precipitation step was carried out to purify the RNA probe.

### In situ hybridization

Sagittal paraffin sections were obtained as previously described. Slides were deparaffinized using Xylene and rehydrated through a graded series of EtOH/PBS. Slides were incubated in pre-heated pre-hybridization solution pH 6 [250ml deionized-formamide, 125ml 20x saline sodium citrate (SSC), 5ml 1M sodium citrate, 500ul tween and 119.7ml DEPC-treated ddH20] at 61°C for 2 hours. Slides were transferred to pre-heated pre-hybridization solution containing DIG labelled RNA probe (1:500) and incubated overnight at 61°C. The following day, slides underwent a series of 61°C SSC stringency washes to remove nonspecific probe binding [2×30m 50:50 pre-hybridization solution:2x SSC; 2×30m 2x SSC; 2×30m 0.2x SSC]. Following the stringency washes, samples were incubated in blocking solution (2% Roche Blocking Reagent (Roche)) for 2hr at room temperature and then incubated in blocking solution containing anti-Digoxigenin-AP antibody (1:2000; Roche) overnight at 4°C. Excess antibody was washed off through 6×1hr MAB-T (0.1% tween-20) washes. Slides were then color reacted with BM-purple (Roche) at room temperature and left until sufficient coloration had taken place. Following the color reaction, a DAPI nuclear counterstain (1µg/ml) was carried out before mounting the slides using Fluoromount (Sigma). Images were taken using a BX51 Olympus compound microscope. Images were contrast enhanced and merged in Adobe Photoshop.

### Double in situ hybridization/immunohistochemistry

For double *in situ* hybridization/immunohistochemistry, samples first underwent *in situ* hybridization as previously described. Immediately after color reaction, samples were fixed for 1 minute in 4% paraformaldehyde in PBS. Samples were then blocked with 5% goat serum and 1% bovine serum albumin in PBS-T (0.05% tween-20). Blocking solution was replaced with a blocking solution containing rabbit anti-Sox2 primary antibody (ab97959; Abcam) at a concentration of 1:500. Goat anti-rabbit Alexa-Fluor 647 (1:250) (A-20721245; Thermo) and goat anti-mouse Alexa-Fluor 488 (1:250) (A-11-001; Thermo) secondary antibodies were used for immunodetection. Samples were counterstained with DAPI (1µg/ml) and mounted using Fluoromount (Sigma). Images were taken using a BX51 Olympus compound microscope. Images were contrast enhanced and merged in Adobe Photoshop.

### MYC inhibition experiments

*Scyliorhinus* shark embryos (supplied by North Wales Biologicals, Wales, United Kingdom, and kept at the University of Florida) of ∼110 days post fertilization (dpf) (Stage 32) (Ballard et al., 1993), were moved from their egg cases and treated with the C-myc/N-myc inhibitors 10058-F4 and 10074-G5 (Sigma-Aldrich) (*81*), for a total of 7 days. Inhibitors were reconstituted in DMSO and dissolved in artificial saltwater (Instant Ocean) to a final concentration of either 0.5 or 2 µM for G4, and 0.5, 2, 5, 10 µM in F4. Embryos were treated in individual wells of a 6-well culture plate, with water changes taking place every 48 hours. Following the treatments, embryos were fixed overnight in 4% PFA in PBS at 4°C, before subsequent dehydration and paraffin embedding. For replicate numbers, see the table below.

## Gene nomenclature

The gene nomenclature conventions and the presentation of expressed genes (italic, lowercase) and protein (Capitalized first letter and lowercase) codes are reflective of the “fish” rules, set by the zebrafish community, ZFIN Nomenclature Conventions (*82*): (https://zfin.atlassian.net/wiki/spaces/general/pages/1818394635/ZFIN=Zebrafish=Nomenclature=Conventions)

## Acknowledgments

We would like to thank the Earth Sciences Department, Natural History Museum, for funding for the initial RNAseq project. We thank Farah Ahmed, Amin Garbout, and Brett Clark (Image and Analysis Centre, Natural History Museum, London) for assistance with embryo micro-CT scanning. We are grateful to past and present members of the Fraser Lab for stimulating discussion associated with this extended project. We also extend our gratitude to Edward Stanley, Gary Scheiffele and Jaimi Gray for additional assistance with micro-CT scanning at the Nanoscale Research Facility and Florida Museum of Natural History, University of Florida. We also thank Michael Hayle at the University of Bangor and North Wales Biologicals, and the Marine Biological Laboratory at Woods Hole, for the support and supply of embryonic specimens. Research presented in this manuscript was also funded by the National Science Foundation IntBIO Collaborative Research grant: 2128032 (GJF); Natural Environment Research Council, UK (NERC) Standard Grant NE/ K014595/1 (to G.J.F); NERC Ph.D studentship (to APT); and Leverhulme Trust, UK, Research Grant RPG-211 (to G.J.F.).

## Author contributions

*Conceptualization*: KJM, KJ, ZJ, GJF, APT; *Methodology*: KJM, APT, KJ, RLC, GJF, CH, WAD, ASIS, EFN, SRB; *Investigation*: KJM, APT, RLC, ASIS, KJ, ZJ, CH, WAD; *Visualization*: KJM, APT, RLC, ASIS, KJ, ZJ, CH, WAD, KEC; *Supervision*: GJF, KJ, ZJ; *Writing—original draft*: APT, KJM, GJF; *Writing—review & editing*: KJ, ZJ, GJF, RLC, KEC, ASIS, EFN, WAD

## Data and materials availability

All data are available in the main text (links in Methods) or the supplementary materials.

