## Supplemental Material for "Shark tooth regeneration: RNAseq reveals genes for unlimited dental renewal" (from another version of this preprint: version 3, 2025-09-01)

Thiery et al.

**This PDF file includes:**

Supplementary Text

Figs. S1 to S5

Tables S1 to S7

Movies S1

**Fig. S1.** Methods schematic showing tissue separation prior to dissection for RNAseq. (A) Tissue from the lower jaw (B) was removed, and the soft tissues that included the DL, teeth and taste buds were part of the same sheet of oral tissue that was flattened to separate the compartments for collection, digestion and RNAseq. Each of the compartments were separated and a sterile scalpel was used to cut each compartment: BHTB, TTJ, SL, ET and LT. (C) condensed lower jaw conveyor belt prior to the flattening and separation of the dental lamina. (D) condensed lower jaw unit prior to flattening and separation of the entire unit containing all compartments. First row teeth (at stage 34, hatching) have begun to erupt beyond the surface epithelia (yellow dotted line). Note the Late-stage teeth (LT) are juxtaposed with the TTJ prior to expansion and flattening of the dental lamina, which folds into the jaw. Hence, after flattening the order of compartments within the oral sheet is: BHTB, TTJ, SL, ET and LT (See Figure 1 and 2).

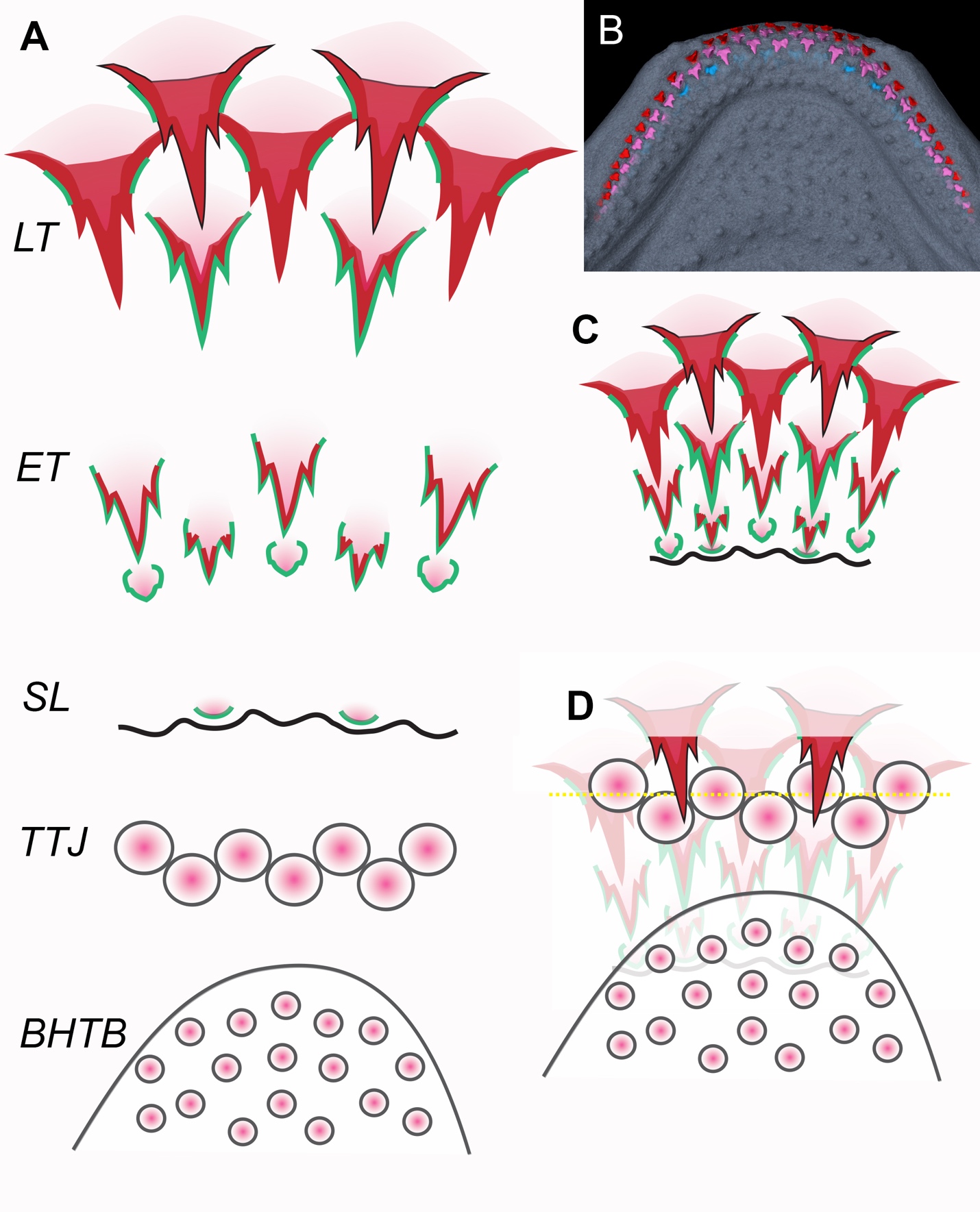

**Fig. S2.** Phylogenetic gene trees for bmp4, dkk1, pitx1, pitx3, lef1 and mycn reconstructed from protein coding sequences (cds) extracted from www.ensembl.org. Species included in the analysis were selected based on their phylogenetic position. Ensembl sequences were aligned to *S. canicula* sequences obtained from RNAseq. Sequences were aligned using MUSCLE (Edgar, 2004). A maximum likelihood tree was generated from 100 bootstrap replications using PHYML (Guindon and Gascuel, 2003) with a GTR substitution model. Node labels represent bootstrap confidence values. Trees were edited in figTree v1.4.3.

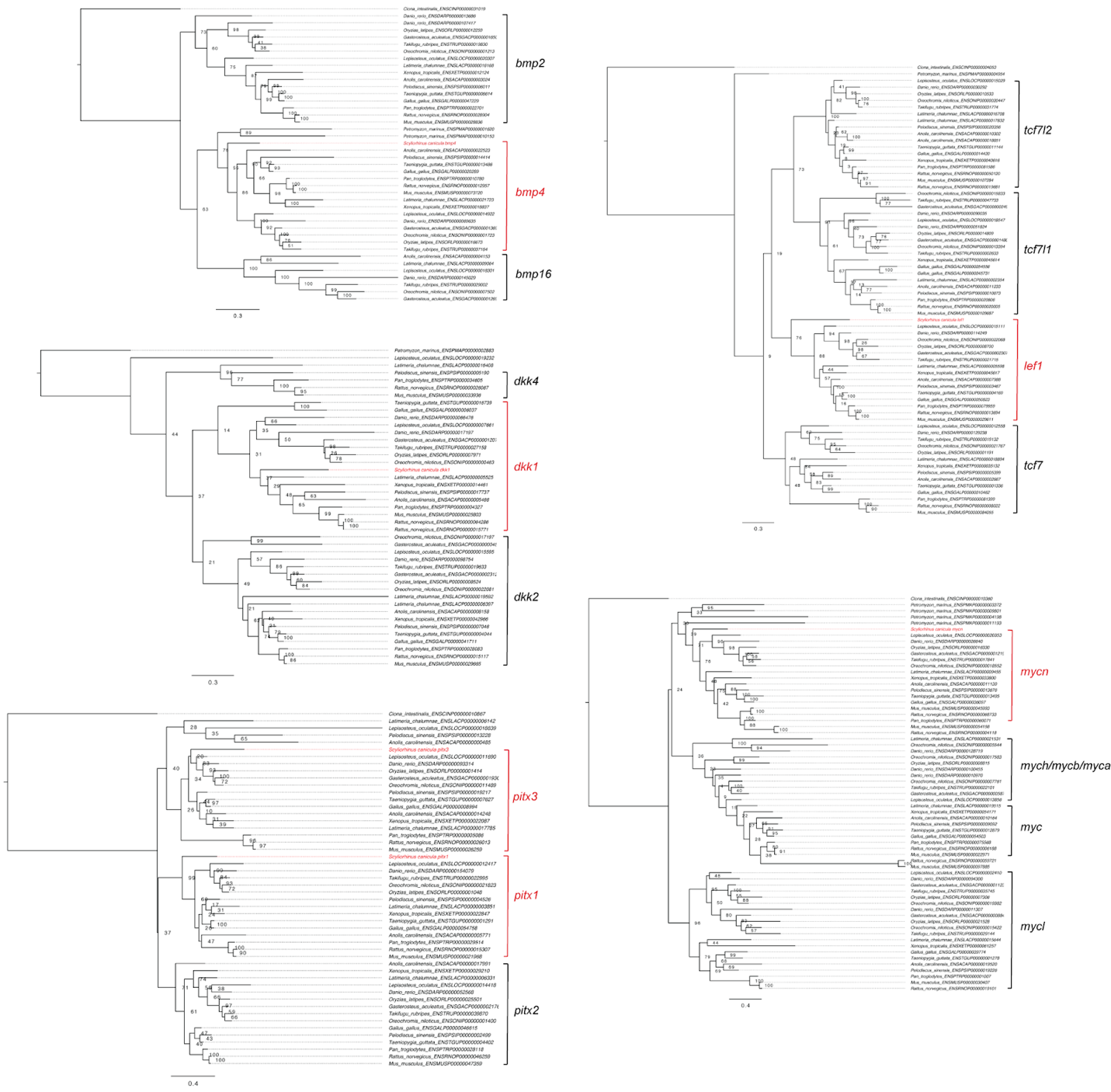

Fig. S3. Phylogenetic gene tree for foxl1 and foxl1l reconstructed from protein coding sequences (cds) extracted from [www.ensembl.org](http://www.ensembl.org) (foxl1) and protein sequences from genbank [www.ncbi.nlm.nih.gov/genbank](http://www.ncbi.nlm.nih.gov/genbank) (foxl1-like). Species included in the analysis were selected based on their phylogenetic position. Ensembl and genbank sequences were aligned to *S. canicula* sequences obtained from RNAseq. Sequences were aligned using MUSCLE (Edgar, 2004). A maximum likelihood tree was generated from 100 bootstrap replications using PHYML (Guindon and Gascuel, 2003) with an LG substitution model. Node labels represent bootstrap confidence values. Trees were edited in figTree v1.4.3.

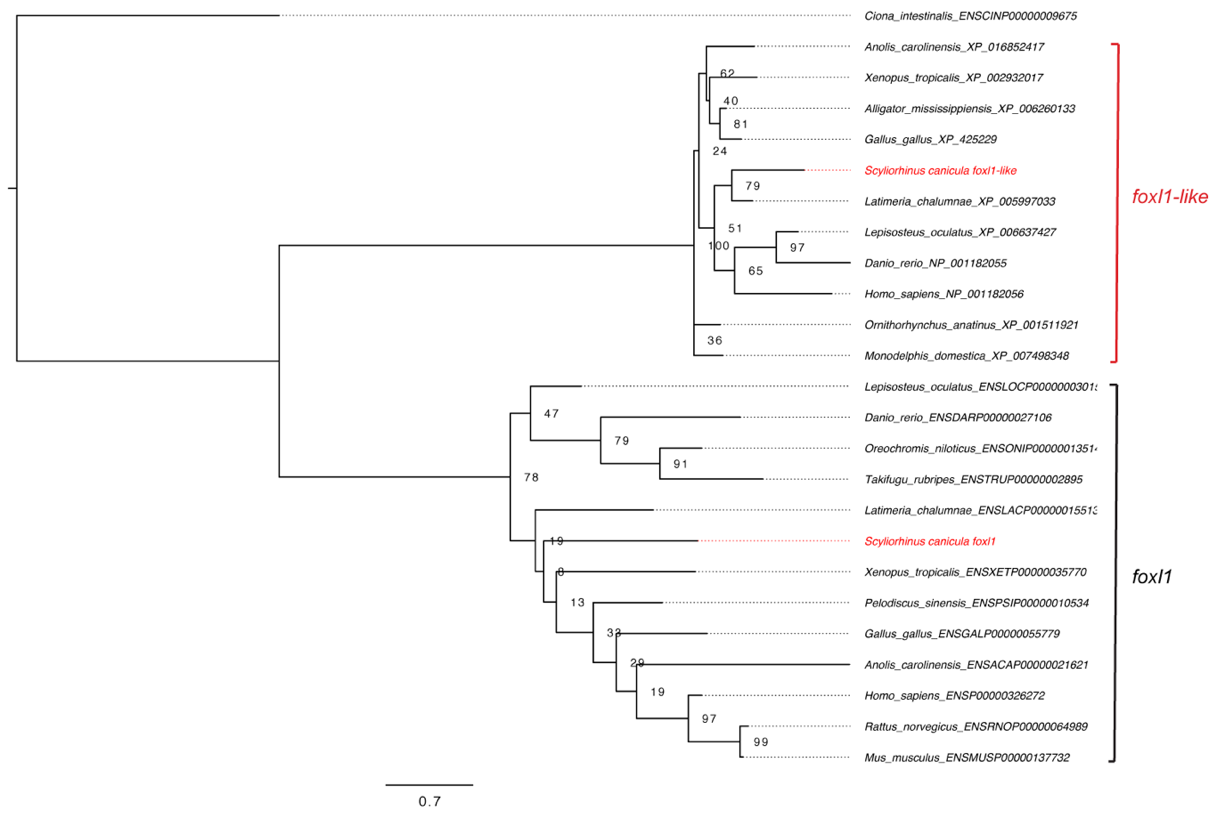

**Fig. S4.** Full RNEA/Cytoscape gene regulatory network representing the SL (Successional lamina) of the catshark.

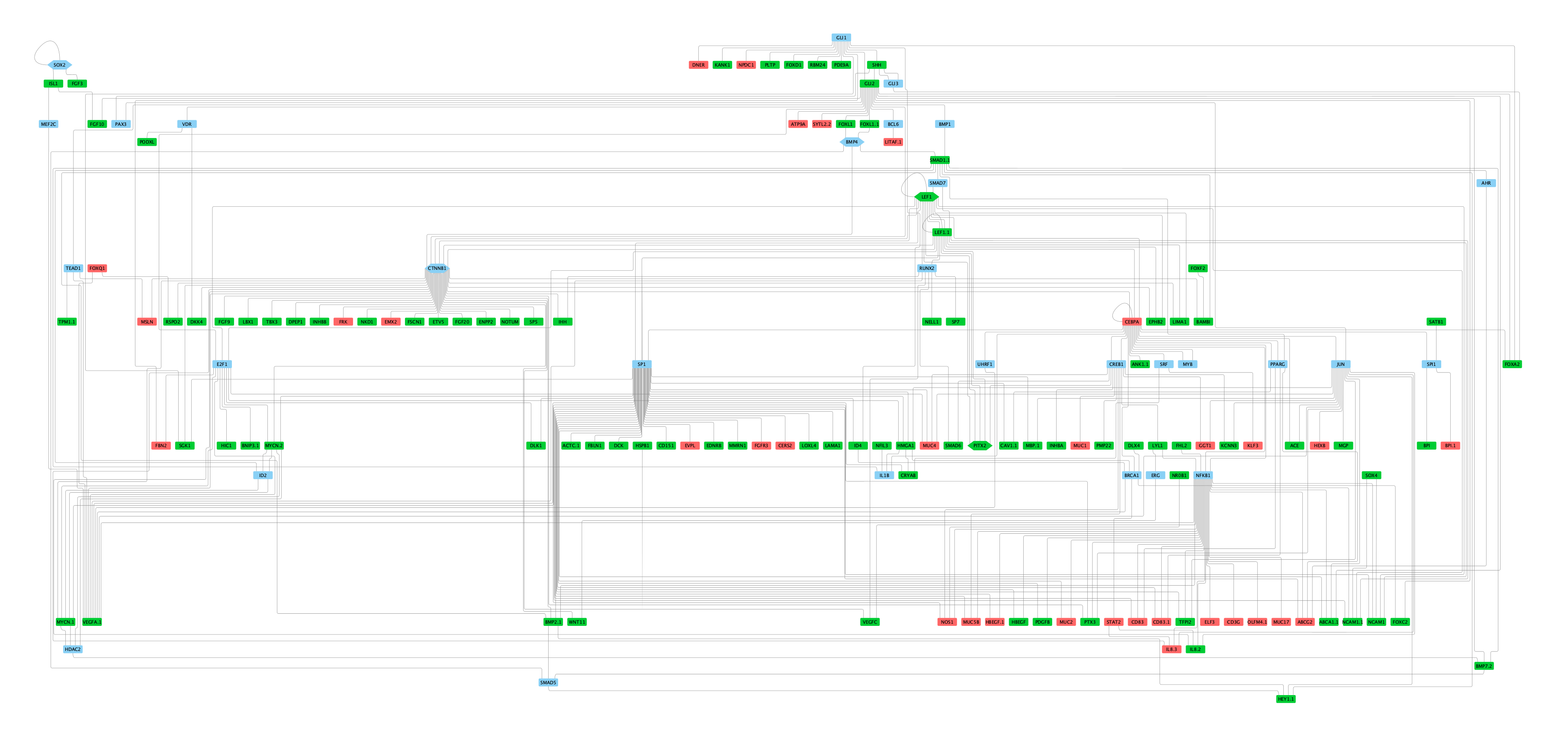

**Figure S5.** MYC/MYCN chemical perturbations results in shifts in dental lamina morphology and expression. A, DMSO control thin section showing equivalent stage of tooth development with advanced first tooth (normal) and second-generation tooth cap forming, with *mycn* expression (magenta) in the SL (arrowhead). B, dysmorphic dental lamina in the 10058-F4 (0.5μM) treated sharks, showing *mycn* expression lacking from the overgrown SL (arrowhead). Dotted line denotes the outline of the first-generation tooth that fails to form correctly in F4 treatments. C, DMSO treated control thin section through the shark upper jaw (stage 33) showing first generation tooth (dotted line) and second-generation tooth forming in the dental lamina, with Sox2 (magenta) and PCNA (green) immunofluorescence expressed throughout the lamina and within the SL (arrowhead). DAPI used as the nuclear counter stain in A-D). D, MYC/MYCN inhibitor 10058-F4 (10μM) treated shark lower jaw section showing PCNA immunofluorescence (green) and Sox2 expression (magenta) restricted to the taste/tooth junction (arrowhead). E, Haematoxylin and Eosin (H&E) stained thin section histology showing sections through three regions of the shark upper jaw treated with MYC/MYCN inhibitor 10058-F4 (10μM), E1, sagittal; E2, medio-lateral; and E3, lateral section along the jaw – E1, showing the first symphyseal tooth not forming correctly and second tooth fails to form; E2, enlarged medial valve (asterisk), epithelium connecting the dental lamina showing stunted growth of the first tooth and no sign of a second tooth; E3, lateral section without a first-generation tooth (black arrowhead) and abnormal rostral (aboral) outgrowth of the dental lamina (asterisk), neighboring a dental lamina (white arrowhead) with condensing mesenchyme.

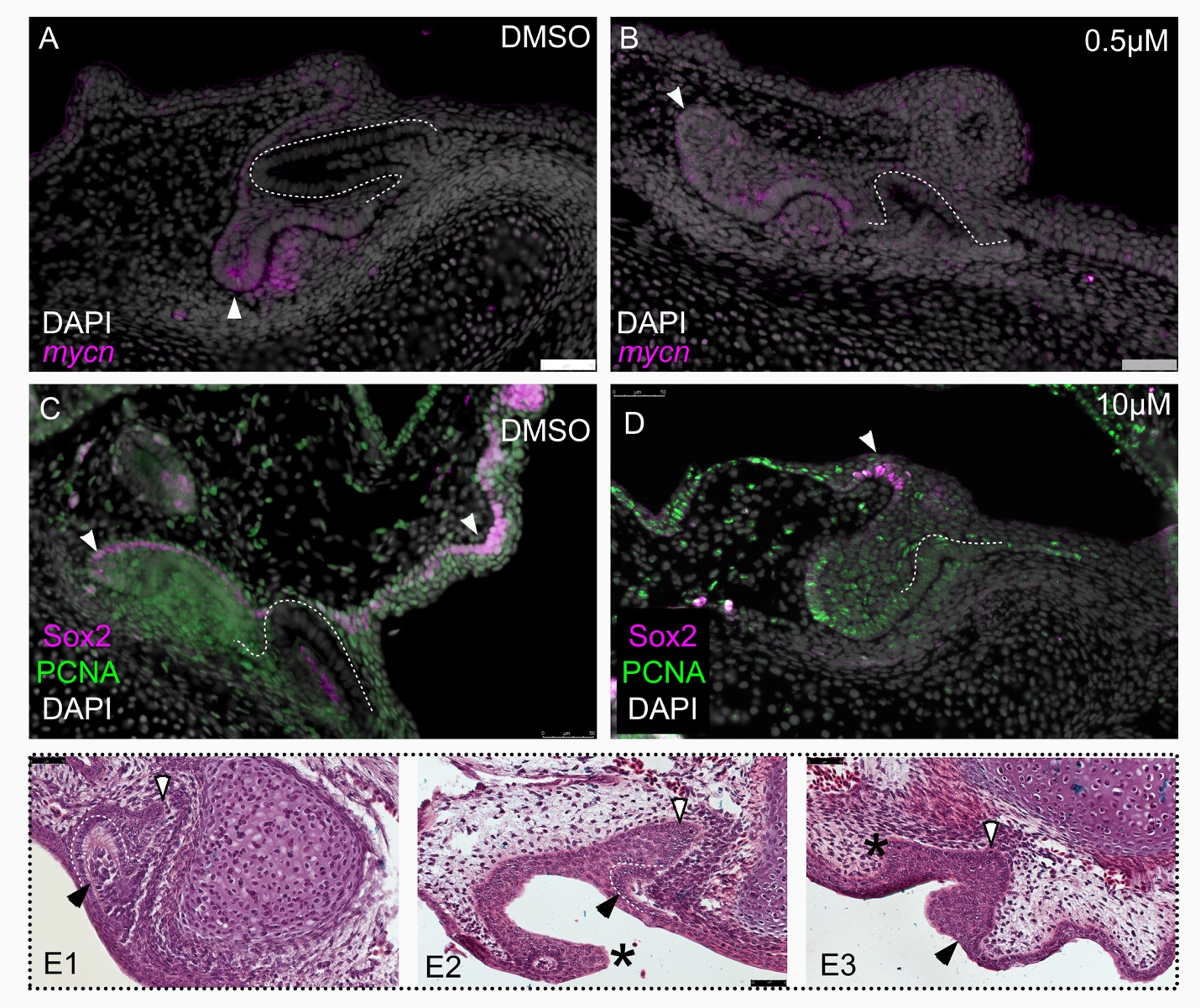

Table S1. List of forward and reverse primers used in the synthesis of probes for *in situ* hybridization.
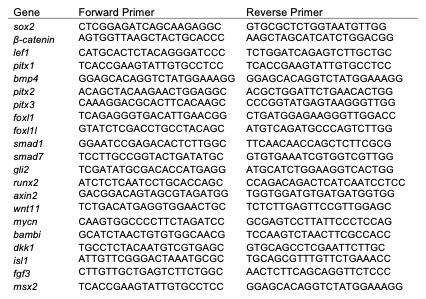

Table S2. Supplementary data to accompany Figure 3D. Top gene ontology (Functional Enrichment Analysis) for each compartment (BHTB).

GOBPID,Pvalue,OddsRatio,ExpCount,Count,Size,Term,FDR

GO:0002376,1.31658172533592e-15,1.95610463032334,131.548634590377,220,1863,immune system process,7.41235511364125e-13

GO:0006955,6.38712186906379e-14,2.06841674081681,89.1113133940182,160,1262,immune response,1.79797480614146e-11

GO:0043207,3.73076857977506e-12,2.10799102198438,69.1989596879064,128,980,response to external biotic stimulus,5.2510567760334e-10

GO:0051707,3.73076857977506e-12,2.10799102198438,69.1989596879064,128,980,response to other organism,5.2510567760334e-10

GO:0050896,1.26313536770484e-11,1.52625430938684,462.079583875163,568,6544,response to stimulus,1.42229042403565e-09

GO:0009607,1.75663372211628e-11,2.05156623973476,70.7524057217165,128,1002,response to biotic stimulus,1.64830797591911e-09

GO:0044419,5.51246658681998e-11,1.95834559905799,79.5788036410923,138,1127,biological process involved in interspecies interaction between organisms,4.43359812625664e-09

GO:0006952,9.49095489551014e-11,1.95423912193341,77.8841352405722,135,1103,defense response,6.67925950771526e-09

GO:0009605,1.89068893020509e-10,1.72507444842655,130.913133940182,200,1854,response to external stimulus,1.18273096411718e-08

GO:0098542,6.75831551416118e-10,2.10545709609539,52.8877763328999,99,749,defense response to other organism,3.80493163447275e-08

GO:0019882,1.32093176203548e-09,4.3380376344086,8.6851755526658,30,123,antigen processing and presentation,6.76076892750887e-08

GO:0002757,1.45963433876622e-08,2.49057399716892,25.9849154746424,57,368,immune response-activating signaling pathway,6.84811777271152e-07

GO:0002764,3.99036520374821e-08,2.38743072750855,27.3971391417425,58,388,immune response-regulating signaling pathway,1.72813508439249e-06

GO:0032101,6.37645317966989e-08,1.95118404522613,51.6167750325098,91,731,regulation of response to external stimulus,2.56424510011011e-06

GO:0045087,9.1627664459157e-08,2.02547676901554,43.778933680104,80,620,innate immune response,3.43909167270036e-06

GO:0002429,1.20331583053741e-07,2.76485167464115,17.0172951885566,41,241,immune response-activating cell surface receptor signaling pathway,4.23416757870351e-06

GO:0002253,1.43760248257447e-07,2.25065425995929,29.7979193758127,60,422,activation of immune response,4.76100116287897e-06

GO:0001775,2.35688015820535e-07,1.80790512409206,63.1970091027308,104,895,cell activation,7.37179738372008e-06

GO:0042221,2.7684249555408e-07,1.48569782110033,181.965019505852,244,2577,response to chemical,8.20328026299721e-06

GO:0007154,4.27686906781766e-07,1.38699125963975,335.120676202861,409,4746,cell communication,1.20393864259067e-05

GO:0009617,4.87677833864903e-07,2.22466190245708,28.0326397919376,56,397,response to bacterium,1.30744104983781e-05

GO:0002682,5.30820122465007e-07,1.7184613679193,73.8592977893368,116,1046,regulation of immune system process,1.34040291543087e-05

GO:0023052,5.47589112875845e-07,1.38496012336458,327.282834850455,400,4635,signaling,1.34040291543087e-05

GO:0046903,6.50036587533037e-07,1.7886877230078,60.5843953185956,99,858,secretion,1.52487749492125e-05

GO:0002768,8.71132844276119e-07,2.53332162767218,18.2882964889467,41,259,immune response-regulating cell surface receptor signaling pathway,1.96179116530982e-05

GO:0080134,1.22021961256391e-06,1.72586486551916,67.0100130039012,106,949,regulation of response to stress,2.55414782747647e-05

GO:0050776,1.22490215527291e-06,1.87203642802934,47.3801040312094,81,671,regulation of immune response,2.55414782747647e-05

GO:0032940,1.3884950297073e-06,1.79769076596857,55.2885565669701,91,783,secretion by cell,2.79186679187576e-05

GO:0050778,1.53698101672949e-06,1.96527584522241,38.5537061118335,69,546,positive regulation of immune response,2.9838631462714e-05

GO:0031347,1.59822833919773e-06,1.99521813179615,36.3647594278283,66,515,regulation of defense response,2.9993418498944e-05

GO:0006954,1.65283488187535e-06,1.9818331868785,37.1414824447334,67,526,inflammatory response,3.00176141450265e-05

GO:0008219,2.4984683301816e-06,1.56516880836459,102.880494148244,148,1457,cell death,4.17713522777603e-05

GO:1901700,2.51958497988659e-06,1.59393798588151,92.7830949284785,136,1314,response to oxygen-containing compound,4.17713522777603e-05

GO:0002758,2.54788871458904e-06,2.6259906766102,15.1107932379714,35,214,innate immune response-activating signaling pathway,4.17713522777603e-05

GO:0012501,2.59679809897267e-06,1.56609778472996,102.103771131339,147,1446,programmed cell death,4.17713522777603e-05

GO:0006950,3.02499422262878e-06,1.41657150759995,201.030039011704,259,2847,response to stress,4.60467376388528e-05

GO:0044281,3.19699636815756e-06,1.58789643960725,92.3594278283485,135,1308,small molecule metabolic process,4.60467376388528e-05

GO:0140352,3.21854580431599e-06,1.75004343610409,57.1950585175553,92,810,export from cell,4.60467376388528e-05

GO:0000165,3.23942758858068e-06,1.89830883208564,40.8838751625488,71,579,MAPK cascade,4.60467376388528e-05

GO:0007165,3.27152665284922e-06,1.36005655573043,303.133810143043,369,4293,signal transduction,4.60467376388528e-05

GO:0002274,4.82182518198003e-06,2.28802974090397,20.9009102730819,43,296,myeloid leukocyte activation,6.62118921330429e-05

GO:0006534,4.96214399664477e-06,18.5399443929564,0.847334200260078,7,12,cysteine metabolic process,6.65163588121668e-05

GO:0002684,5.25635014380505e-06,1.76535354889013,52.3228868660598,85,741,positive regulation of immune system process,6.74985202910342e-05

GO:0032103,5.2785187485674e-06,2.03474266624909,30.2215864759428,56,428,positive regulation of response to external stimulus,6.74985202910342e-05

GO:0006810,5.39508599129047e-06,1.366596716684,262.391157347204,324,3716,transport,6.74985202910342e-05

GO:0006629,5.55075212226452e-06,1.65126533742331,70.8936280884265,108,1004,lipid metabolic process,6.79363792355419e-05

GO:0042127,6.47870540446417e-06,1.51119803293108,114.107672301691,159,1616,regulation of cell population proliferation,7.76066200577304e-05

GO:0010033,6.76997006489478e-06,1.45272401739279,148.283485045514,198,2100,response to organic substance,7.78875250881823e-05

GO:0002444,6.77884321371391e-06,2.53419940853401,15.1107932379714,34,214,myeloid leukocyte mediated immunity,7.78875250881823e-05

GO:0006915,7.30040291236089e-06,1.54345679012346,98.9262678803641,141,1401,apoptotic process,8.22025367931836e-05

GO:0006887,7.73298051233754e-06,1.94205796269727,34.3170351105332,61,486,exocytosis,8.53660397734517e-05

GO:0002218,8.05841066802212e-06,2.47203164905604,15.8875162548765,35,225,activation of innate immune response,8.68647700915717e-05

GO:0140112,8.23365959631335e-06,11.7790146361575,1.20039011703511,8,17,extracellular vesicle biogenesis,8.68647700915717e-05

GO:0070371,8.33161205141185e-06,2.50599590523545,15.2520156046814,34,216,ERK1 and ERK2 cascade,8.68647700915717e-05

GO:0043408,8.90356182994322e-06,1.95523321305153,32.975422626788,59,467,regulation of MAPK cascade,9.06738700293145e-05

GO:0071702,9.06159286771309e-06,1.4323289917308,158.733940182055,209,2248,organic substance transport,9.06738700293145e-05

GO:0051716,9.18012538485066e-06,1.32099958093474,384.76033810143,451,5449,cellular response to stimulus,9.06738700293145e-05

GO:0031349,9.63586804650524e-06,2.15102831196581,23.5841352405722,46,334,positive regulation of defense response,9.35343743134905e-05

GO:0072593,1.03705190443535e-05,2.89720125786164,10.3092327698309,26,146,reactive oxygen species metabolic process,9.81315090123405e-05

GO:0002250,1.04580649036242e-05,2.1635446685879,22.9486345903771,45,325,adaptive immune response,9.81315090123405e-05

GO:0007186,1.06636106896924e-05,1.91878389109472,34.6700910273082,61,491,G protein-coupled receptor signaling pathway,9.84198822671613e-05

GO:0009636,1.1021737130121e-05,3.13425614836716,8.54395318595579,23,121,response to toxic substance,0.000100084483939647

GO:0043299,1.20032542514393e-05,2.53752961386883,14.1928478543563,32,201,leukocyte degranulation,0.000106325534437402

GO:0050865,1.20867392610901e-05,1.96945792880259,31.0689206762029,56,440,regulation of cell activation,0.000106325534437402

GO:0051241,1.34235611579345e-05,1.72083543755862,52.8171651495449,84,748,negative regulation of multicellular organismal process,0.000116268691260263

GO:0050909,1.39034708152416e-05,10.6003710575139,1.27100130039012,8,18,sensory perception of taste,0.000118600819226985

GO:0032496,1.42887907420198e-05,2.47123143771188,14.9695708712614,33,212,response to lipopolysaccharide,0.000120068495339659

GO:0045321,1.48313009638294e-05,1.69888539902887,55.3591677503251,87,784,leukocyte activation,0.000122757102165709

GO:0002443,1.50448313489058e-05,2.02231884057971,27.6089726918075,51,391,leukocyte mediated immunity,0.000122757102165709

GO:0002833,1.64264815084875e-05,2.25775615057801,19.1356306892068,39,271,positive regulation of response to biotic stimulus,0.000132115844132549

GO:0050866,1.70533264163169e-05,2.80295191722459,10.591677503251,26,150,negative regulation of cell activation,0.000135225672850513

GO:0002275,1.74315614206893e-05,2.44357872534668,15.1107932379714,33,214,myeloid cell activation involved in immune response,0.000136305126109001

GO:0051234,1.80418654374043e-05,1.33831198011859,270.65266579974,329,3833,establishment of localization,0.000138034982812988

GO:0002220,1.81431416130748e-05,3.63329928498468,5.93133940182055,18,84,innate immune response activating cell surface receptor signaling pathway,0.000138034982812988

GO:0002366,1.86032748347624e-05,2.05215157353886,25.6318595578674,48,363,leukocyte activation involved in immune response,0.00013964858309295

GO:0002263,2.00031392213624e-05,2.04551108509549,25.7024707412224,48,364,cell activation involved in immune response,0.000146278666982488

GO:0070372,2.00061409549762e-05,2.46373966716831,14.5459037711313,32,206,regulation of ERK1 and ERK2 cascade,0.000146278666982488

GO:0002252,2.04861624578743e-05,1.8418953241454,37.7063719115735,64,534,immune effector process,0.000147868070048503

GO:0006749,2.40770298689152e-05,6.63289962825279,2.1183355006502,10,30,glutathione metabolic process,0.000171586934382269

GO:2000707,2.47321230699434e-05,Inf,0.282444733420026,4,4,positive regulation of dense core granule biogenesis,0.000171903522078742

GO:0002159,2.47321230699434e-05,Inf,0.282444733420026,4,4,desmosome assembly,0.000171903522078742

GO:0002831,2.53847202617065e-05,2.04057251078891,25.2081924577373,47,357,regulation of response to biotic stimulus,0.000174287774479765

GO:0051179,2.68844136907073e-05,1.31922413519434,301.156697009103,360,4265,localization,0.000182360541058653

GO:0016192,2.91898381821516e-05,1.4892305790385,103.37477243173,143,1464,vesicle-mediated transport,0.000195641415435135

GO:0044703,2.99983043553378e-05,3.47459697117733,6.14317295188557,18,87,multi-organism reproductive process,0.000198694651200649

GO:0002237,3.10243431020719e-05,2.36416959946372,15.5344603381014,33,220,response to molecule of bacterial origin,0.000203101222865889

GO:0050727,3.21921481804908e-05,2.3246876697447,16.2405721716515,34,230,regulation of inflammatory response,0.000204920985003895

GO:0050912,3.23942587306334e-05,33.0527289546716,0.494278283485045,5,7,detection of chemical stimulus involved in sensory perception of taste,0.000204920985003895

GO:0070813,3.23942587306334e-05,33.0527289546716,0.494278283485045,5,7,hydrogen sulfide metabolic process,0.000204920985003895

GO:0009069,3.3305754400311e-05,6.31660470879802,2.1889466840052,10,31,serine family amino acid metabolic process,0.000208345996970834

GO:0048584,4.16028236322033e-05,1.45693710795642,113.68400520156,154,1610,positive regulation of response to stimulus,0.000257388897856378

GO:0030855,4.33213918841484e-05,1.86926848024608,32.481144343303,56,460,epithelial cell differentiation,0.000265108082943212

GO:0090066,4.61548484540266e-05,1.93982063578605,28.0326397919376,50,397,regulation of anatomical structure size,0.00027941053418943

GO:0050878,4.80226083308509e-05,2.1359803520262,20.0535760728218,39,284,regulation of body fluid levels,0.000287624771173075

GO:0070887,4.99496628210997e-05,1.41555451752868,136.138361508453,179,1928,cellular response to chemical stimulus,0.00029601747545557

GO:0048518,5.08124121543651e-05,1.29925831744014,326.153055916775,384,4619,positive regulation of biological process,0.000297993625446954

GO:0034109,5.20780011913851e-05,3.83591898297781,4.73094928478544,15,67,homotypic cell-cell adhesion,0.000302267161554122

GO:0007166,5.55948569983601e-05,1.39039340774372,154.285435630689,199,2185,cell surface receptor signaling pathway,0.000319386780510987

GO:0014070,6.20839246174663e-05,1.71235917564518,44.6262678803641,71,632,response to organic cyclic compound,0.000352144149470167

GO:0045088,6.25478063002073e-05,2.04327003165084,22.4543563068921,42,318,regulation of innate immune response,0.000352144149470167

GO:0045089,7.19135011370817e-05,2.16338244714926,18.2882964889467,36,259,positive regulation of innate immune response,0.000400864367724525

GO:0002934,7.79914566292614e-05,13.2296296296296,0.847334200260078,6,12,desmosome organization,0.00043048225570857

GO:0007567,8.13408987158482e-05,22.0336108541474,0.564889466840052,5,8,parturition,0.000444610931815753

GO:0051336,8.43274552360832e-05,1.65087425595238,50.6988296488947,78,718,regulation of hydrolase activity,0.000452691938594351

GO:0052547,8.44274485833159e-05,2.14384761904762,18.4295188556567,36,261,regulation of peptidase activity,0.000452691938594351

GO:0048002,9.0570478640468e-05,3.27849405512398,6.07256176853056,17,86,antigen processing and presentation of peptide antigen,0.000481048862967769

GO:0002504,9.32727895305631e-05,4.4252017380509,3.38933680104031,12,48,antigen processing and presentation of peptide or polysaccharide antigen via MHC class II,0.000490771780427169

GO:0044706,9.56791085702127e-05,3.01233989378319,7.27295188556567,19,103,multi-multicellular organism process,0.000498771649305831

GO:0042981,9.67481723002911e-05,1.53277218928123,73.2944083224967,105,1038,regulation of apoptotic process,0.000499717623899669

GO:0002683,0.000102784436475863,2.05075566043036,20.7596879063719,39,294,negative regulation of immune system process,0.000526069433962824

GO:0043409,0.000107332081177026,2.64454860998475,9.8149544863459,23,139,negative regulation of MAPK cascade,0.000544396051375368

GO:0000096,0.000113293783027812,7.06444032158318,1.62405721716515,8,23,sulfur amino acid metabolic process,0.000569503570041593

GO:1901607,0.000115675794170249,4.30529971312094,3.45994798439532,12,49,alpha-amino acid biosynthetic process,0.000576329647389878

GO:0001580,0.000116699076025659,52.8391866913124,0.353055916775033,4,5,detection of chemical stimulus involved in sensory perception of bitter taste,0.000576329647389878

GO:0002695,0.000120217776557497,2.69081534110506,9.25006501950585,22,131,negative regulation of leukocyte activation,0.000588544419146702

GO:0009084,0.000136725654834205,8.42370882129918,1.27100130039012,7,18,glutamine family amino acid biosynthetic process,0.000663590893721184

GO:0009611,0.00014200326036418,1.84177817739462,29.3036410923277,50,415,response to wounding,0.000683314834060112

GO:0008285,0.000144912347206844,1.68441876232404,42.6491547464239,67,604,negative regulation of cell population proliferation,0.000687483266913166

GO:0043067,0.000145311738477206,1.5107746554119,74.9184655396619,106,1061,regulation of programmed cell death,0.000687483266913166

GO:0010817,0.000149506101024215,1.97086313045566,22.5955786736021,41,320,regulation of hormone levels,0.000701432790638609

GO:0002712,0.000155609918965407,5.67966573816156,2.1183355006502,9,30,regulation of B cell mediated immunity,0.00071810151129118

GO:0002889,0.000155609918965407,5.67966573816156,2.1183355006502,9,30,regulation of immunoglobulin mediated immune response,0.00071810151129118

GO:0045216,0.000157091225930053,2.4529439485502,11.368400520156,25,161,cell-cell junction organization,0.000719043578850567

GO:0050907,0.000172350062393796,16.5240518038853,0.635500650195059,5,9,detection of chemical stimulus involved in sensory perception,0.000769533595745962

GO:0043129,0.000172350062393796,16.5240518038853,0.635500650195059,5,9,surfactant homeostasis,0.000769533595745962

GO:0097070,0.000172350062393796,16.5240518038853,0.635500650195059,5,9,ductus arteriosus closure,0.000769533595745962

GO:0042119,0.000173589283587455,2.34097347411795,12.7806241872562,27,181,neutrophil activation,0.000769533595745962

GO:0051046,0.000181372629513561,1.76653803684816,33.4697009102731,55,474,regulation of secretion,0.000797756175126052

GO:0043066,0.000190729466813108,1.67963194374083,41.4487646293888,65,587,negative regulation of apoptotic process,0.000832408448184338

GO:0006690,0.000192721509107451,3.75266420843632,4.16605981794538,13,59,icosanoid metabolic process,0.000834632381749962

GO:0016477,0.000199182709168693,1.47264079353895,83.2505851755527,115,1179,cell migration,0.000856029505816598

GO:0043069,0.000206279574138773,1.66732760519938,42.3667100130039,66,600,negative regulation of programmed cell death,0.000879813638182795

GO:1901615,0.000210608346756708,1.8315817130846,28.2444733420026,48,400,organic hydroxy compound metabolic process,0.000891522550556592

GO:0045649,0.000224113466014562,9.92083333333333,0.988556566970091,6,14,regulation of macrophage differentiation,0.000939183307412967

GO:2000377,0.000227075678174572,3.12831225948323,5.93133940182055,16,84,regulation of reactive oxygen species metabolic process,0.000939183307412967

GO:0010466,0.000227075678174572,3.12831225948323,5.93133940182055,16,84,negative regulation of peptidase activity,0.000939183307412967

GO:0036230,0.000228540165391788,2.29575431695567,12.9924577373212,27,184,granulocyte activation,0.000939183307412967

GO:0002694,0.000236780774308336,1.82097253518827,28.3856957087126,48,402,regulation of leukocyte activation,0.000965996927069515

GO:0034097,0.000277273864795559,1.60557701026718,48.5098829648895,73,687,response to cytokine,0.0011230588912223

GO:0050864,0.000280702545758139,2.93621906624713,6.63745123537061,17,94,regulation of B cell activation,0.00112882523758451

GO:0045055,0.000291402166946147,1.84060237576439,26.3379713914174,45,373,regulated exocytosis,0.00116354198574951

GO:1901606,0.000324727580158724,3.52216748768473,4.3778933680104,13,62,alpha-amino acid catabolic process,0.00127426730367898

GO:0050913,0.000330436256285178,26.4177449168207,0.423667100130039,4,6,sensory perception of bitter taste,0.00127426730367898

GO:0061110,0.000330436256285178,26.4177449168207,0.423667100130039,4,6,dense core granule biogenesis,0.00127426730367898

GO:2000705,0.000330436256285178,26.4177449168207,0.423667100130039,4,6,regulation of dense core granule biogenesis,0.00127426730367898

GO:1990858,0.000339568643872032,4.05266149870801,3.31872561768531,11,47,cellular response to lectin,0.00127426730367898

GO:1990840,0.000339568643872032,4.05266149870801,3.31872561768531,11,47,response to lectin,0.00127426730367898

GO:0002223,0.000339568643872032,4.05266149870801,3.31872561768531,11,47,stimulatory C-type lectin receptor signaling pathway,0.00127426730367898

GO:0070527,0.000339568643872032,4.05266149870801,3.31872561768531,11,47,platelet aggregation,0.00127426730367898

GO:0052548,0.000346192583160189,2.04255038514298,17.5821846553966,33,249,regulation of endopeptidase activity,0.00127426730367898

GO:0070445,0.00035115960890722,Inf,0.21183355006502,3,3,regulation of oligodendrocyte progenitor proliferation,0.00127426730367898

GO:0097734,0.00035127190124665,8.8179012345679,1.0591677503251,6,15,extracellular exosome biogenesis,0.00127426730367898

GO:1990182,0.00035127190124665,8.8179012345679,1.0591677503251,6,15,exosomal secretion,0.00127426730367898

GO:0030728,0.00035127190124665,8.8179012345679,1.0591677503251,6,15,ovulation,0.00127426730367898

GO:0031953,0.00035127190124665,8.8179012345679,1.0591677503251,6,15,negative regulation of protein autophosphorylation,0.00127426730367898

GO:0044283,0.00035323864015595,1.77327559542221,29.6566970091027,49,420,small molecule biosynthetic process,0.00127426730367898

GO:0140962,0.000355346299604972,4.41883519206939,2.82444733420026,10,40,multicellular organismal-level chemical homeostasis,0.00127426730367898

GO:0002221,0.000371036958857342,2.29922974422308,12.0039011703511,25,170,pattern recognition receptor signaling pathway,0.00131462634947433

GO:0008652,0.000371271029425254,3.70300116928674,3.88361508452536,12,55,amino acid biosynthetic process,0.00131462634947433

GO:0050790,0.000378614804870198,1.42319464982261,93.3479843953186,125,1322,regulation of catalytic activity,0.00133225084463701

GO:0048821,0.000398017574585967,5.57562738013866,1.90650195058518,8,27,erythrocyte development,0.00138323391661666

GO:0070266,0.000398017574585967,5.57562738013866,1.90650195058518,8,27,necroptotic process,0.00138323391661666

GO:0051248,0.000423317694174598,1.59465444775484,46.7446033810143,70,662,negative regulation of protein metabolic process,0.00146213412159692

GO:0002283,0.000431542671732549,2.31818553999836,11.4390117035111,24,162,neutrophil activation involved in immune response,0.00148145441576479

GO:0006536,0.000441502189169728,4.76958217270195,2.40078023407022,9,34,glutamate metabolic process,0.00150116772841596

GO:0007606,0.000442617838218559,3.61858811579482,3.95422626788036,12,56,sensory perception of chemical stimulus,0.00150116772841596

GO:0051607,0.000453704691826239,2.08658854166667,15.6756827048114,30,222,defense response to virus,0.00152045084225103

GO:0140546,0.000453704691826239,2.08658854166667,15.6756827048114,30,222,defense response to symbiont,0.00152045084225103

GO:0007565,0.000484057346202285,3.01926831338596,5.71950585175553,15,81,female pregnancy,0.00161256973912359

GO:0006879,0.000498798981936525,3.83882496940024,3.45994798439532,11,49,intracellular iron ion homeostasis,0.00164704054450614

GO:0030001,0.000501448355318532,1.58348966865719,47.0270481144343,70,666,metal ion transport,0.00164704054450614

GO:0071310,0.000503181125497435,1.38601216986295,107.258387516255,140,1519,cellular response to organic substance,0.00164704054450614

GO:1903530,0.000512391917644938,1.72973968077061,30.9276983094928,50,438,regulation of secretion by cell,0.00166749508459017

GO:0009065,0.000528602677096669,7.93555555555556,1.1297789336801,6,16,glutamine family amino acid catabolic process,0.00171036383451394

GO:0019221,0.000542319235272736,1.82076033180821,24.7845253576073,42,351,cytokine-mediated signaling pathway,0.00174471845404886

GO:0070493,0.000560619957505684,11.0144927536232,0.776723016905072,5,11,thrombin-activated receptor signaling pathway,0.00179106675504791

GO:0042060,0.00056308848249286,1.8466407331707,23.3016905071521,40,330,wound healing,0.00179106675504791

GO:0050900,0.000593234820854778,1.95039899476166,18.8531859557867,34,267,leukocyte migration,0.00187293207329221

GO:0045834,0.000595479291508535,2.6305099394987,7.69661898569571,18,109,positive regulation of lipid metabolic process,0.00187293207329221

GO:1901698,0.000600332785484991,1.50587944165926,59.8782834850455,85,848,response to nitrogen compound,0.00187770754571139

GO:0010243,0.000641497197132518,1.51725836835241,56.6301690507152,81,802,response to organonitrogen compound,0.00199537525958899

GO:1901184,0.000679521372265142,3.04719231710301,5.29583875162549,14,75,regulation of ERBB signaling pathway,0.00209433654938018

GO:0033993,0.000680752377507234,1.5908823791292,43.425877763329,65,615,response to lipid,0.00209433654938018

GO:0034113,0.000697354857564165,4.41566078613432,2.54200260078023,9,36,heterotypic cell-cell adhesion,0.002122220458425

GO:0097300,0.000697354857564165,4.41566078613432,2.54200260078023,9,36,programmed necrotic cell death,0.002122220458425

GO:1901605,0.000769416936655229,2.30650938369546,10.521066319896,22,149,alpha-amino acid metabolic process,0.00232893406095104

GO:0044248,0.000785967663996559,1.43977388211382,74.9890767230169,102,1062,cellular catabolic process,0.00234041515220703

GO:0051259,0.00078825283388318,1.67356405011387,33.116644993498,52,469,protein complex oligomerization,0.00234041515220703

GO:0050851,0.000789838150833632,2.25246250899231,11.227178153446,23,159,antigen receptor-mediated signaling pathway,0.00234041515220703

GO:0043270,0.000789838150833632,2.25246250899231,11.227178153446,23,159,positive regulation of monoatomic ion transport,0.00234041515220703

GO:0050852,0.000806164909552679,2.4776081004098,8.54395318595579,19,121,T cell receptor signaling pathway,0.0023460603054078

GO:0050817,0.000808310487419116,2.12643824516726,13.3455136540962,26,189,coagulation,0.0023460603054078

GO:0007596,0.000808310487419116,2.12643824516726,13.3455136540962,26,189,blood coagulation,0.0023460603054078

GO:0051924,0.00080841154395935,2.16348213520692,12.6394018205462,25,179,regulation of calcium ion transport,0.0023460603054078

GO:0030216,0.000820368165249511,2.84593837535014,6.00195058517555,15,85,keratinocyte differentiation,0.00236855013864346

GO:0016050,0.000830698943756104,1.95125332501538,17.7234070221066,32,251,vesicle organization,0.00238614033334024

GO:0007599,0.000875175768828672,2.11332259549011,13.4161248374512,26,190,hemostasis,0.00250113684188093

GO:0010959,0.000882082899440979,1.900623637781,19.2768530559168,34,273,regulation of metal ion transport,0.00250814481002662

GO:0043085,0.000892520623997781,1.50142261866907,56.4183355006502,80,799,positive regulation of catalytic activity,0.0025250709111093

GO:1901701,0.000910772535803531,1.46872240653585,64.1149544863459,89,908,cellular response to oxygen-containing compound,0.00256382468828694

GO:0001818,0.000981990011153692,2.30964963333721,10.0267880364109,21,142,negative regulation of cytokine production,0.00273848880718469

GO:0006811,0.000982548381973903,1.4708889114396,62.5615084525358,87,886,monoatomic ion transport,0.00273848880718469

GO:2000117,0.00100670262623399,3.4722480620155,3.74239271781534,11,53,negative regulation of cysteine-type endopeptidase activity,0.002778301855734

GO:0006826,0.00100670262623399,3.4722480620155,3.74239271781534,11,53,iron ion transport,0.002778301855734

GO:0002446,0.00103178344803373,2.16002443121087,12.1451235370611,24,172,neutrophil mediated immunity,0.00283362966459996

GO:0042742,0.00103896583417023,2.35451360640223,9.39128738621586,20,133,defense response to bacterium,0.00283950371183418

GO:0010611,0.00106283918331981,4.11055614254154,2.68322496749025,9,38,regulation of cardiac muscle hypertrophy,0.00287681952023584

GO:0042743,0.00106283918331981,4.11055614254154,2.68322496749025,9,38,hydrogen peroxide metabolic process,0.00287681952023584

GO:0032501,0.00109481998443899,1.21932981647968,412.016254876463,460,5835,multicellular organismal process,0.00294920407291461

GO:0016053,0.00111878841763709,1.98370327244499,15.8169050715215,29,224,organic acid biosynthetic process,0.00299941847204611

GO:0051249,0.00119456960253172,1.779956384404,23.4429128738622,39,332,regulation of lymphocyte activation,0.00318173370206488

GO:0002697,0.00120314198509476,1.92416620742863,17.3703511053316,31,246,regulation of immune effector process,0.00318173370206488

GO:0070374,0.00120374649829453,2.38344149528744,8.82639791937581,19,125,positive regulation of ERK1 and ERK2 cascade,0.00318173370206488

GO:0016064,0.00131530124573348,2.81535504296698,5.64889466840052,14,80,immunoglobulin mediated immune response,0.00339289040342056

GO:0010951,0.00131530124573348,2.81535504296698,5.64889466840052,14,80,negative regulation of endopeptidase activity,0.00339289040342056

GO:0019344,0.00133044206641905,39.5927977839335,0.282444733420026,3,4,cysteine biosynthetic process,0.00339289040342056

GO:0006526,0.00133044206641905,39.5927977839335,0.282444733420026,3,4,arginine biosynthetic process,0.00339289040342056

GO:0046351,0.00133044206641905,39.5927977839335,0.282444733420026,3,4,disaccharide biosynthetic process,0.00339289040342056

GO:0070444,0.00133044206641905,39.5927977839335,0.282444733420026,3,4,oligodendrocyte progenitor proliferation,0.00339289040342056

GO:0090330,0.00133428223696634,5.14529914529915,1.76527958387516,7,25,regulation of platelet aggregation,0.00339289040342056

GO:0006575,0.00133662020895071,2.69134680899387,6.28439531859558,15,89,cellular modified amino acid metabolic process,0.00339289040342056

GO:0009615,0.00134672238965178,1.86679704476315,18.9944083224967,33,269,response to virus,0.00339289040342056

GO:0048870,0.00134675940999135,1.37782134676403,91.1590377113134,119,1291,cell motility,0.00339289040342056

GO:0086103,0.00137431027399996,13.2070240295749,0.564889466840052,4,8,G protein-coupled receptor signaling pathway involved in heart process,0.00339289040342056

GO:0045651,0.00137431027399996,13.2070240295749,0.564889466840052,4,8,positive regulation of macrophage differentiation,0.00339289040342056

GO:0031663,0.00137516060532539,4.41249226963513,2.25955786736021,8,32,lipopolysaccharide-mediated signaling pathway,0.00339289040342056

GO:0061436,0.00138608311329792,8.25971322849214,0.917945383615084,5,13,establishment of skin barrier,0.00339289040342056

GO:0014741,0.00138608311329792,8.25971322849214,0.917945383615084,5,13,negative regulation of muscle hypertrophy,0.00339289040342056

GO:0010614,0.00138608311329792,8.25971322849214,0.917945383615084,5,13,negative regulation of cardiac muscle hypertrophy,0.00339289040342056

GO:2001026,0.00138608311329792,8.25971322849214,0.917945383615084,5,13,regulation of endothelial cell chemotaxis,0.00339289040342056

GO:0009063,0.00143822273927461,2.92313646200262,5.08400520156047,13,72,amino acid catabolic process,0.00350527879745284

GO:0043122,0.00145917952020631,1.99915014164306,14.6165149544863,27,207,regulation of canonical NF-kappaB signal transduction,0.00353500100122524

GO:0032879,0.00146297554757634,1.34495484385768,108.31755526658,138,1534,regulation of localization,0.00353500100122524

GO:0019724,0.00148899400085939,2.77313989752729,5.71950585175553,14,81,B cell mediated immunity,0.00355406914268334

GO:0048485,0.00149082055572608,6.10299145299145,1.34161248374512,6,19,sympathetic nervous system development,0.00355406914268334

GO:0010499,0.00149082055572608,6.10299145299145,1.34161248374512,6,19,proteasomal ubiquitin-independent protein catabolic process,0.00355406914268334

GO:0048583,0.00149611791619174,1.25843172587268,210.562548764629,249,2982,regulation of response to stimulus,0.00355406914268334

GO:0007034,0.00153536394147416,2.21721439749609,10.379843953186,21,147,vacuolar transport,0.0036319743657561

GO:0014743,0.00156961344358314,3.84481984005751,2.82444733420026,9,40,regulation of muscle hypertrophy,0.00369745760977953

GO:0007249,0.00157859562899994,1.9334247871334,16.1699609882965,29,229,canonical NF-kappaB signal transduction,0.00369835339399725

GO:0051345,0.00160153713898891,1.65634466339091,30.1509752925878,47,427,positive regulation of hydrolase activity,0.00369835339399725

GO:0071222,0.00160211743363709,2.3173520889401,9.03823146944083,19,128,cellular response to lipopolysaccharide,0.00369835339399725

GO:0042445,0.00160283877111071,2.45489486313906,7.69661898569571,17,109,hormone metabolic process,0.00369835339399725

GO:0007043,0.00160283877111071,2.45489486313906,7.69661898569571,17,109,cell-cell junction assembly,0.00369835339399725

GO:0043154,0.00163831506214206,3.48659753472902,3.38933680104031,10,48,negative regulation of cysteine-type endopeptidase activity involved in apoptotic process,0.00374947715441454

GO:0002474,0.00163831506214206,3.48659753472902,3.38933680104031,10,48,antigen processing and presentation of peptide antigen via MHC class I,0.00374947715441454

GO:0071705,0.00166229142123899,1.32271147319512,121.310013003901,152,1718,nitrogen compound transport,0.00378894765245971

GO:0071985,0.00170446401353717,4.23569573283859,2.33016905071521,8,33,multivesicular body sorting pathway,0.00384886203848979

GO:0006692,0.00171592250739065,4.87415248036681,1.83589076723017,7,26,prostanoid metabolic process,0.00384886203848979

GO:0006693,0.00171592250739065,4.87415248036681,1.83589076723017,7,26,prostaglandin metabolic process,0.00384886203848979

GO:0042744,0.00171592250739065,4.87415248036681,1.83589076723017,7,26,hydrogen peroxide catabolic process,0.00384886203848979

GO:0006812,0.00176674423124672,1.49046882111971,50.9812743823147,72,722,monoatomic cation transport,0.00394713096107897

GO:0051240,0.00182421026437141,1.39446160300687,77.8135240572172,103,1102,positive regulation of multicellular organismal process,0.0040489344262233

GO:0043312,0.00182669510525882,2.13664178694912,11.227178153446,22,159,neutrophil degranulation,0.0040489344262233

GO:0043086,0.00188070625552772,1.65076819962879,29.5860858257477,46,419,negative regulation of catalytic activity,0.00413731007043198

GO:0032482,0.0018812635488998,3.16942366026289,4.02483745123537,11,57,Rab protein signal transduction,0.00413731007043198

GO:0043407,0.00193049867245116,3.39695929844629,3.45994798439532,10,49,negative regulation of MAP kinase activity,0.00421687842377626

GO:0030162,0.00193242386027402,1.58936526396261,35.3055916775033,53,500,regulation of proteolysis,0.00421687842377626

GO:1903034,0.00196046003514943,2.40232470194853,7.83784135240572,17,111,regulation of response to wounding,0.00426154053972638

GO:0006520,0.00200005231897101,1.92348917428332,15.6756827048114,28,222,amino acid metabolic process,0.00433088252146416

GO:0061158,0.00203162180623311,7.34145338678179,0.988556566970091,5,14,3'-UTR-mediated mRNA destabilization,0.00434906112893248

GO:0009070,0.00203162180623311,7.34145338678179,0.988556566970091,5,14,serine family amino acid biosynthetic process,0.00434906112893248

GO:0060487,0.00203162180623311,7.34145338678179,0.988556566970091,5,14,lung epithelial cell differentiation,0.00434906112893248

GO:0038127,0.00210698547909776,2.25480318650422,9.25006501950585,19,131,ERBB signaling pathway,0.00449330615428802

GO:0040011,0.00212662279292508,1.43230519480519,62.4908972691808,85,885,locomotion,0.00451807031100687

GO:0050670,0.00215178994404206,2.14838569880823,10.662288686606,21,151,regulation of lymphocyte proliferation,0.00453729490073288

GO:0032944,0.00215178994404206,2.14838569880823,10.662288686606,21,151,regulation of mononuclear cell proliferation,0.00453729490073288

GO:0019752,0.00225563328472682,1.50530244182512,45.5442132639792,65,645,carboxylic acid metabolic process,0.0047369496844316

GO:0007032,0.00226330277995044,3.3118029739777,3.53055916775032,10,50,endosome organization,0.0047369496844316

GO:0002460,0.0022907865968471,1.95630529794265,14.3340702210663,26,203,adaptive immune response based on somatic recombination of immune receptors built from immunoglobulin superfamily domains,0.00475877699788238

GO:0032102,0.00231751651952884,1.7970695970696,19.6299089726918,33,278,negative regulation of response to external stimulus,0.00475877699788238

GO:0009064,0.00232148950338382,2.89263585576435,4.73094928478544,12,67,glutamine family amino acid metabolic process,0.00475877699788238

GO:0016054,0.00233506294842485,2.13183528652833,10.732899869961,21,152,organic acid catabolic process,0.00475877699788238

GO:0046395,0.00233506294842485,2.13183528652833,10.732899869961,21,152,carboxylic acid catabolic process,0.00475877699788238

GO:0060670,0.0023358784432849,10.5648798521257,0.635500650195059,4,9,branching involved in labyrinthine layer morphogenesis,0.00475877699788238

GO:0060644,0.0023358784432849,10.5648798521257,0.635500650195059,4,9,mammary gland epithelial cell differentiation,0.00475877699788238

GO:0000041,0.00234135209309666,2.52012197283977,6.63745123537061,15,94,transition metal ion transport,0.00475877699788238

GO:0140888,0.00238521832686135,2.73677125401263,5.36644993498049,13,76,interferon-mediated signaling pathway,0.00479838435606894

GO:0060840,0.00238640785026519,2.61617090603321,6.00195058517555,14,85,artery development,0.00479838435606894

GO:0061041,0.00238640785026519,2.61617090603321,6.00195058517555,14,85,regulation of wound healing,0.00479838435606894

GO:0043410,0.00245566403776132,1.78963704166956,19.7005201560468,33,279,positive regulation of MAPK cascade,0.00492006709345061

GO:0007041,0.00250387247233533,2.41393372982158,7.34356306892068,16,104,lysosomal transport,0.00498932566340145

GO:0061515,0.0025079558840899,3.03693798449612,4.16605981794538,11,59,myeloid cell development,0.00498932566340145

GO:0043436,0.00252484091166695,1.49237472766885,46.6033810143043,66,660,oxoacid metabolic process,0.00500523039883272

GO:0097529,0.00253400034122803,2.16156439237938,10.0973992197659,20,143,myeloid leukocyte migration,0.00500576207758379

GO:2000379,0.00254804515388431,3.92139077853364,2.47139141742523,8,35,positive regulation of reactive oxygen species metabolic process,0.00501590706866038

GO:0032528,0.00263968920543228,5.28851851851852,1.48283485045514,6,21,microvillus organization,0.00511107406325642

GO:0071875,0.00263968920543228,5.28851851851852,1.48283485045514,6,21,adrenergic receptor signaling pathway,0.00511107406325642

GO:0050672,0.00264067456397784,3.23080061655635,3.60117035110533,10,51,negative regulation of lymphocyte proliferation,0.00511107406325642

GO:0032945,0.00264067456397784,3.23080061655635,3.60117035110533,10,51,negative regulation of mononuclear cell proliferation,0.00511107406325642

GO:0030593,0.00264178073251797,2.84078212290503,4.80156046814044,12,68,neutrophil chemotaxis,0.00511107406325642

GO:0016266,0.00267526828319819,3.50483368834999,3.03628088426528,9,43,O-glycan processing,0.00515813713507048

GO:0006066,0.00275329379184333,1.7749517507582,19.8417425227568,33,281,alcohol metabolic process,0.00529045871948052

GO:0006082,0.00281419641077991,1.48455020199034,46.8152145643693,66,663,organic acid metabolic process,0.00538909040567717

GO:0034134,0.00287179675872634,6.60684551341351,1.0591677503251,5,15,toll-like receptor 2 signaling pathway,0.005436742268178

GO:0070254,0.00287179675872634,6.60684551341351,1.0591677503251,5,15,mucus secretion,0.005436742268178

GO:0060479,0.00287179675872634,6.60684551341351,1.0591677503251,5,15,lung cell differentiation,0.005436742268178

GO:0048844,0.00287918937289433,2.97475083056478,4.23667100130039,11,60,artery morphogenesis,0.005436742268178

GO:0032535,0.00288736401098618,1.72227342549923,22.242522756827,36,315,regulation of cellular component size,0.005436742268178

GO:0070663,0.00296581851566263,2.08366617616145,10.944733420026,21,155,regulation of leukocyte proliferation,0.00556585274772687

GO:0071219,0.00298301510220736,2.17643732023398,9.53250975292588,19,135,cellular response to molecule of bacterial origin,0.00556785588595279

GO:0006816,0.00298666514663898,1.69138676844784,23.8665799739922,38,338,calcium ion transport,0.00556785588595279

GO:0070664,0.00306674634501669,3.15365551425031,3.67178153446034,10,52,negative regulation of leukocyte proliferation,0.00564982432732849

GO:0034340,0.00306674634501669,3.15365551425031,3.67178153446034,10,52,response to type I interferon,0.00564982432732849

GO:0010543,0.00307681380997578,3.78107606679035,2.54200260078023,8,36,regulation of platelet activation,0.00564982432732849

GO:1902532,0.00312152934757558,1.60221354166667,30.3628088426528,46,430,negative regulation of intracellular signal transduction,0.00564982432732849

GO:0050917,0.00315105655200914,19.7950138504155,0.353055916775033,3,5,sensory perception of umami taste,0.00564982432732849

GO:0061817,0.00315105655200914,19.7950138504155,0.353055916775033,3,5,endoplasmic reticulum-plasma membrane tethering,0.00564982432732849

GO:0086042,0.00315105655200914,19.7950138504155,0.353055916775033,3,5,cardiac muscle cell-cardiac muscle cell adhesion,0.00564982432732849

GO:0040034,0.00315105655200914,19.7950138504155,0.353055916775033,3,5,regulation of development, heterochronic,0.00564982432732849

GO:0048505,0.00315105655200914,19.7950138504155,0.353055916775033,3,5,regulation of timing of cell differentiation,0.00564982432732849

GO:0099509,0.00315105655200914,19.7950138504155,0.353055916775033,3,5,regulation of presynaptic cytosolic calcium ion concentration,0.00564982432732849

GO:0005984,0.00315105655200914,19.7950138504155,0.353055916775033,3,5,disaccharide metabolic process,0.00564982432732849

GO:0006537,0.00315105655200914,19.7950138504155,0.353055916775033,3,5,glutamate biosynthetic process,0.00564982432732849

GO:0002822,0.00321922012209756,2.21379525593009,8.89700910273082,18,126,regulation of adaptive immune response based on somatic recombination of immune receptors built from immunoglobulin superfamily domains,0.00575371723409818

GO:0030041,0.0032376802958669,2.10965186574943,10.3092327698309,20,146,actin filament polymerization,0.00576662156348481

GO:0042326,0.00324692546292129,1.78885537842884,18.5001300390117,31,262,negative regulation of phosphorylation,0.00576662156348481

GO:0098771,0.00331092868547796,1.64490136069083,26.4085825747724,41,374,inorganic ion homeostasis,0.00585254505412141

GO:0045471,0.00331892365138802,2.50958047599839,6.21378413524057,14,88,response to ethanol,0.00585254505412141

GO:0043068,0.00332649097214716,1.58759393744957,31.2807542262679,47,443,positive regulation of programmed cell death,0.00585254505412141

GO:0051250,0.00335594644051294,2.33386053199137,7.5553966189857,16,107,negative regulation of lymphocyte activation,0.00587231034750583

GO:1901575,0.00335858602468362,1.30560644448775,113.472171651495,141,1607,organic substance catabolic process,0.00587231034750583

GO:0007200,0.00338099789322484,4.20861066644199,2.04772431729519,7,29,phospholipase C-activating G protein-coupled receptor signaling pathway,0.00589173639081271

GO:0032801,0.0034156022358848,4.95763888888889,1.55344603381014,6,22,receptor catabolic process,0.00589173639081271

GO:0002714,0.0034156022358848,4.95763888888889,1.55344603381014,6,22,positive regulation of B cell mediated immunity,0.00589173639081271

GO:0002891,0.0034156022358848,4.95763888888889,1.55344603381014,6,22,positive regulation of immunoglobulin mediated immune response,0.00589173639081271

GO:0051260,0.00342913391256611,1.73093779932081,20.9009102730819,34,296,protein homooligomerization,0.00589173639081271

GO:0035556,0.00343248585468307,1.26059386903449,161.34655396619,193,2285,intracellular signal transduction,0.00589173639081271

GO:0043065,0.00356342304845203,1.58945040747366,30.5746423927178,46,433,positive regulation of apoptotic process,0.00609789415282217

GO:0051047,0.00362779559248533,1.79257541785569,17.8646293888166,30,253,positive regulation of secretion,0.00618923914717952

GO:0061626,0.00367670020871125,8.80345040049292,0.706111833550065,4,10,pharyngeal arch artery morphogenesis,0.00620567044588253

GO:0034351,0.00367670020871125,8.80345040049292,0.706111833550065,4,10,negative regulation of glial cell apoptotic process,0.00620567044588253

GO:0070257,0.00367670020871125,8.80345040049292,0.706111833550065,4,10,positive regulation of mucus secretion,0.00620567044588253

GO:0065008,0.00368151674764612,1.26605128205128,150.472431729519,181,2131,regulation of biological quality,0.00620567044588253

GO:0030838,0.00370341983904934,3.30965645311049,3.17750325097529,9,45,positive regulation of actin filament polymerization,0.00622395632652173

GO:0046394,0.00372901999902137,1.85304167518472,15.6050715214564,27,221,carboxylic acid biosynthetic process,0.00624832815312211

GO:0071345,0.00374653840094514,1.48215728052585,43.990767230169,62,623,cellular response to cytokine stimulus,0.00625310425554711

GO:0002224,0.0037540839047512,2.85769265845873,4.3778933680104,11,62,toll-like receptor signaling pathway,0.00625310425554711

GO:0045595,0.0037852290565364,1.35040816326531,82.3326397919376,106,1166,regulation of cell differentiation,0.00628638335938051

GO:0044093,0.00381871159831789,1.35757925626594,78.8020806241873,102,1116,positive regulation of molecular function,0.00631268005178797

GO:0032680,0.00382371913723813,2.69576744626456,5.01339401820546,12,71,regulation of tumor necrosis factor production,0.00631268005178797

GO:0022407,0.00383470084851063,1.66306469778501,24.2196358907672,38,343,regulation of cell-cell adhesion,0.00631268005178797

GO:0006979,0.00385155748005461,1.74821911842474,19.4886866059818,32,276,response to oxidative stress,0.00632194420195552

GO:0023056,0.00389908784395813,1.34347473645675,85.0864759427828,109,1205,positive regulation of signaling,0.00637847243934865

GO:0051641,0.00390865540244278,1.23494113656977,199.264759427828,233,2822,cellular localization,0.00637847243934865

GO:0030033,0.00393688497764893,6.00580270793037,1.1297789336801,5,16,microvillus assembly,0.00638751078506152

GO:0006525,0.00393688497764893,6.00580270793037,1.1297789336801,5,16,arginine metabolic process,0.00638751078506152

GO:0002696,0.00407661292202577,1.7579066249874,18.7825747724317,31,266,positive regulation of leukocyte activation,0.00658562812480048

GO:0002532,0.00408238759423688,3.00988509631632,3.81300390117035,10,54,production of molecular mediator involved in inflammatory response,0.00658562812480048

GO:0061082,0.00414580694900828,4.02534552927429,2.1183355006502,7,30,myeloid leukocyte cytokine production,0.00666775452608929

GO:0030833,0.00415698372763293,2.15350744002429,9.10884265279584,18,129,regulation of actin filament polymerization,0.00666775452608929

GO:0008284,0.00417094068820043,1.3916983260049,64.0443433029909,85,907,positive regulation of cell population proliferation,0.00667113524845693

GO:0097746,0.00423228210185843,2.5346472232882,5.71950585175553,13,81,blood vessel diameter maintenance,0.00671204175590506

GO:0035150,0.00423228210185843,2.5346472232882,5.71950585175553,13,81,regulation of tube size,0.00671204175590506

GO:0035296,0.00423228210185843,2.5346472232882,5.71950585175553,13,81,regulation of tube diameter,0.00671204175590506

GO:0019884,0.00426480176274477,2.80254025044723,4.44850455136541,11,63,antigen processing and presentation of exogenous antigen,0.00673674943246794

GO:0050867,0.00429693190885205,1.73375912127243,19.6299089726918,32,278,positive regulation of cell activation,0.00673674943246794

GO:0032640,0.00430081841683083,2.65065176908752,5.08400520156047,12,72,tumor necrosis factor production,0.00673674943246794

GO:0019751,0.00430150649868372,2.34124237930466,7.06111833550065,15,100,polyol metabolic process,0.00673674943246794

GO:0051239,0.00430769057848749,1.26006128702758,151.037321196359,181,2139,regulation of multicellular organismal process,0.00673674943246794

GO:0002752,0.00438690340716444,3.52850958565244,2.68322496749025,8,38,cell surface pattern recognition receptor signaling pathway,0.00680393007777845

GO:0006809,0.00438690340716444,3.52850958565244,2.68322496749025,8,38,nitric oxide biosynthetic process,0.00680393007777845

GO:0046209,0.00438690340716444,3.52850958565244,2.68322496749025,8,38,nitric oxide metabolic process,0.00680393007777845

GO:0008064,0.00442326007109184,2.0441622167701,10.591677503251,20,150,regulation of actin polymerization or depolymerization,0.00682404220251087

GO:0048522,0.00450709986727866,1.20050566962918,295.225357607282,333,4181,positive regulation of cellular process,0.00682404220251087

GO:2000145,0.00451396042136491,1.43805309734513,50.3457737321196,69,713,regulation of cell motility,0.00682404220251087

GO:0051289,0.00452985409809213,2.41129579375848,6.42561768530559,14,91,protein homotetramerization,0.00682404220251087

GO:0001816,0.00454029819267603,1.57430179398585,30.1509752925878,45,427,cytokine production,0.00682404220251087

GO:0001933,0.00455285499878734,1.77980895413068,17.3703511053316,29,246,negative regulation of protein phosphorylation,0.00682404220251087

GO:0033559,0.00468105642367669,2.9427922346138,3.88361508452536,10,55,unsaturated fatty acid metabolic process,0.00682404220251087

GO:0030168,0.00473088278654694,2.31385577486809,7.13172951885566,15,101,platelet activation,0.00682404220251087

GO:0003014,0.00473088278654694,2.31385577486809,7.13172951885566,15,101,renal system process,0.00682404220251087

GO:0065009,0.00473754148775653,1.27321772860159,131.054356306892,159,1856,regulation of molecular function,0.00682404220251087

GO:0009967,0.00488869773398552,1.35201331018279,75.907022106632,98,1075,positive regulation of signal transduction,0.00682404220251087

GO:0010647,0.00498160505907088,1.33242112582856,84.8746423927178,108,1202,positive regulation of cell communication,0.00682404220251087

GO:0016068,0.00498167201639781,Inf,0.141222366710013,2,2,type I hypersensitivity,0.00682404220251087

GO:0042636,0.00498167201639781,Inf,0.141222366710013,2,2,negative regulation of hair cycle,0.00682404220251087

GO:0032911,0.00498167201639781,Inf,0.141222366710013,2,2,negative regulation of transforming growth factor beta1 production,0.00682404220251087

GO:1902958,0.00498167201639781,Inf,0.141222366710013,2,2,positive regulation of mitochondrial electron transport, NADH to ubiquinone,0.00682404220251087

GO:0071877,0.00498167201639781,Inf,0.141222366710013,2,2,regulation of adenylate cyclase-inhibiting adrenergic receptor signaling pathway,0.00682404220251087

GO:0008292,0.00498167201639781,Inf,0.141222366710013,2,2,acetylcholine biosynthetic process,0.00682404220251087

GO:1903860,0.00498167201639781,Inf,0.141222366710013,2,2,negative regulation of dendrite extension,0.00682404220251087

GO:0061872,0.00498167201639781,Inf,0.141222366710013,2,2,hepatic stellate cell contraction,0.00682404220251087

GO:0061873,0.00498167201639781,Inf,0.141222366710013,2,2,regulation of hepatic stellate cell contraction,0.00682404220251087

GO:0061586,0.00498167201639781,Inf,0.141222366710013,2,2,positive regulation of transcription by transcription factor localization,0.00682404220251087

GO:0086096,0.00498167201639781,Inf,0.141222366710013,2,2,adenylate cyclase-inhibiting adrenergic receptor signaling pathway involved in heart process,0.00682404220251087

GO:1904387,0.00498167201639781,Inf,0.141222366710013,2,2,cellular response to L-phenylalanine derivative,0.00682404220251087

GO:0099156,0.00498167201639781,Inf,0.141222366710013,2,2,cell-cell signaling via exosome,0.00682404220251087

GO:0014043,0.00498167201639781,Inf,0.141222366710013,2,2,negative regulation of neuron maturation,0.00682404220251087

GO:0014031,0.00498167201639781,Inf,0.141222366710013,2,2,mesenchymal cell development,0.00682404220251087

GO:1904799,0.00498167201639781,Inf,0.141222366710013,2,2,regulation of neuron remodeling,0.00682404220251087

GO:1904800,0.00498167201639781,Inf,0.141222366710013,2,2,negative regulation of neuron remodeling,0.00682404220251087

GO:1900620,0.00498167201639781,Inf,0.141222366710013,2,2,acetate ester biosynthetic process,0.00682404220251087

GO:0001810,0.00498167201639781,Inf,0.141222366710013,2,2,regulation of type I hypersensitivity,0.00682404220251087

GO:0001812,0.00498167201639781,Inf,0.141222366710013,2,2,positive regulation of type I hypersensitivity,0.00682404220251087

GO:0001802,0.00498167201639781,Inf,0.141222366710013,2,2,type III hypersensitivity,0.00682404220251087

GO:0001803,0.00498167201639781,Inf,0.141222366710013,2,2,regulation of type III hypersensitivity,0.00682404220251087

GO:0001805,0.00498167201639781,Inf,0.141222366710013,2,2,positive regulation of type III hypersensitivity,0.00682404220251087

GO:0010813,0.00498167201639781,Inf,0.141222366710013,2,2,neuropeptide catabolic process,0.00682404220251087

GO:0005988,0.00498167201639781,Inf,0.141222366710013,2,2,lactose metabolic process,0.00682404220251087

GO:0005989,0.00498167201639781,Inf,0.141222366710013,2,2,lactose biosynthetic process,0.00682404220251087

GO:0006059,0.00498167201639781,Inf,0.141222366710013,2,2,hexitol metabolic process,0.00682404220251087

GO:0140193,0.00498167201639781,Inf,0.141222366710013,2,2,regulation of adenylate cyclase-inhibiting adrenergic receptor signaling pathway involved in heart process,0.00682404220251087

GO:0140194,0.00498167201639781,Inf,0.141222366710013,2,2,negative regulation of adenylate cyclase-inhibiting adrenergic receptor signaling pathway involved in heart process,0.00682404220251087

GO:0019343,0.00498167201639781,Inf,0.141222366710013,2,2,cysteine biosynthetic process via cystathionine,0.00682404220251087

GO:0021551,0.00498167201639781,Inf,0.141222366710013,2,2,central nervous system morphogenesis,0.00682404220251087

GO:0015886,0.00498167201639781,Inf,0.141222366710013,2,2,heme transport,0.00682404220251087

GO:0015840,0.00498167201639781,Inf,0.141222366710013,2,2,urea transport,0.00682404220251087

GO:0097037,0.00498167201639781,Inf,0.141222366710013,2,2,heme export,0.00682404220251087

GO:0060158,0.00498167201639781,Inf,0.141222366710013,2,2,phospholipase C-activating dopamine receptor signaling pathway,0.00682404220251087

GO:1902178,0.00498167201639781,Inf,0.141222366710013,2,2,fibroblast growth factor receptor apoptotic signaling pathway,0.00682404220251087

GO:0045861,0.00500932993819011,1.97924283288383,11.4390117035111,21,162,negative regulation of proteolysis,0.00682869916513567

GO:0002440,0.00500932993819011,1.97924283288383,11.4390117035111,21,162,production of molecular mediator of immune response,0.00682869916513567

GO:0050829,0.00503461721245172,3.85735248687056,2.1889466840052,7,31,defense response to Gram-negative bacterium,0.00683009515809715

GO:0043552,0.00503461721245172,3.85735248687056,2.1889466840052,7,31,positive regulation of phosphatidylinositol 3-kinase activity,0.00683009515809715

GO:0060627,0.0051226542161889,1.58879554556999,27.8914174252276,42,395,regulation of vesicle-mediated transport,0.00693282289354411

GO:0030832,0.00513898674513874,2.0129057933936,10.732899869961,20,152,regulation of actin filament length,0.00693824829139834

GO:2001057,0.0051844518125777,3.41444730384823,2.75383615084525,8,39,reactive nitrogen species metabolic process,0.00696298716244214

GO:0002381,0.0051844518125777,3.41444730384823,2.75383615084525,8,39,immunoglobulin production involved in immunoglobulin-mediated immune response,0.00696298716244214

GO:0003018,0.00519441315848259,2.287098747545,7.20234070221066,15,102,vascular process in circulatory system,0.00696298716244214

GO:1990266,0.00523638658986674,2.4618825722274,5.86072821846554,13,83,neutrophil migration,0.0069805150350947

GO:0002115,0.00525708414721164,5.50493370336109,1.20039011703511,5,17,store-operated calcium entry,0.0069805150350947

GO:0006622,0.00525708414721164,5.50493370336109,1.20039011703511,5,17,protein targeting to lysosome,0.0069805150350947

GO:0070213,0.00525708414721164,5.50493370336109,1.20039011703511,5,17,protein auto-ADP-ribosylation,0.0069805150350947

GO:0044255,0.00534555844531824,1.42976270393112,49.8514954486346,68,706,cellular lipid metabolic process,0.00706604427447942

GO:0055117,0.00534659833202172,2.87861645385486,3.95422626788036,10,56,regulation of cardiac muscle contraction,0.00706604427447942

GO:1903555,0.00539771057610285,2.56478644800865,5.22522756827048,12,74,regulation of tumor necrosis factor superfamily cytokine production,0.00708036934591023

GO:0006869,0.00539870932163248,1.75516470284682,17.5821846553966,29,249,lipid transport,0.00708036934591023

GO:0051049,0.0054242350625687,1.31988783664122,89.6055916775033,113,1269,regulation of transport,0.00708036934591023

GO:0090280,0.00545731749511444,7.5452865064695,0.776723016905072,4,11,positive regulation of calcium ion import,0.00708036934591023

GO:0006750,0.00545731749511444,7.5452865064695,0.776723016905072,4,11,glutathione biosynthetic process,0.00708036934591023

GO:0001913,0.00545804670714927,4.40617283950617,1.69466840052016,6,24,T cell mediated cytotoxicity,0.00708036934591023

GO:0045191,0.00545804670714927,4.40617283950617,1.69466840052016,6,24,regulation of isotype switching,0.00708036934591023

GO:0060428,0.00545804670714927,4.40617283950617,1.69466840052016,6,24,lung epithelium development,0.00708036934591023

GO:0050870,0.00557572703383977,1.92374851602691,12.2863459037711,22,174,positive regulation of T cell activation,0.00721410742290357

GO:0060326,0.00558676880352745,1.83723788149406,14.5459037711313,25,206,cell chemotaxis,0.00721410742290357

GO:0023061,0.00561670447637015,1.61963149037875,24.7845253576073,38,351,signal release,0.00723616617893911

GO:1902533,0.00567502554432105,1.44573775760254,45.6854356306892,63,647,positive regulation of intracellular signal transduction,0.00729461045993779

GO:0060759,0.00574693614821383,2.07801660967269,9.39128738621586,18,133,regulation of response to cytokine stimulus,0.00732400613443852

GO:0030334,0.00577599020204741,1.43190557776509,48.2980494148244,66,684,regulation of cell migration,0.00732400613443852

GO:0010634,0.00579191646229043,2.12856310782428,8.6851755526658,17,123,positive regulation of epithelial cell migration,0.00732400613443852

GO:0009123,0.00579779427593896,3.05442468395115,3.38933680104031,9,48,nucleoside monophosphate metabolic process,0.00732400613443852

GO:0032966,0.0059716768477331,13.1957525392428,0.423667100130039,3,6,negative regulation of collagen biosynthetic process,0.00732400613443852

GO:0000098,0.0059716768477331,13.1957525392428,0.423667100130039,3,6,sulfur amino acid catabolic process,0.00732400613443852

GO:0051643,0.0059716768477331,13.1957525392428,0.423667100130039,3,6,endoplasmic reticulum localization,0.00732400613443852

GO:1900407,0.0059716768477331,13.1957525392428,0.423667100130039,3,6,regulation of cellular response to oxidative stress,0.00732400613443852

GO:0034154,0.0059716768477331,13.1957525392428,0.423667100130039,3,6,toll-like receptor 7 signaling pathway,0.00732400613443852

GO:0009133,0.0059716768477331,13.1957525392428,0.423667100130039,3,6,nucleoside diphosphate biosynthetic process,0.00732400613443852

GO:0010713,0.0059716768477331,13.1957525392428,0.423667100130039,3,6,negative regulation of collagen metabolic process,0.00732400613443852

GO:0006004,0.0059716768477331,13.1957525392428,0.423667100130039,3,6,fucose metabolic process,0.00732400613443852

GO:0002713,0.0059716768477331,13.1957525392428,0.423667100130039,3,6,negative regulation of B cell mediated immunity,0.00732400613443852

GO:0002890,0.0059716768477331,13.1957525392428,0.423667100130039,3,6,negative regulation of immunoglobulin mediated immune response,0.00732400613443852

GO:0031666,0.0059716768477331,13.1957525392428,0.423667100130039,3,6,positive regulation of lipopolysaccharide-mediated signaling pathway,0.00732400613443852

GO:0021561,0.0059716768477331,13.1957525392428,0.423667100130039,3,6,facial nerve development,0.00732400613443852

GO:0021610,0.0059716768477331,13.1957525392428,0.423667100130039,3,6,facial nerve morphogenesis,0.00732400613443852

GO:0021612,0.0059716768477331,13.1957525392428,0.423667100130039,3,6,facial nerve structural organization,0.00732400613443852

GO:0021637,0.0059716768477331,13.1957525392428,0.423667100130039,3,6,trigeminal nerve structural organization,0.00732400613443852

GO:0032460,0.0059716768477331,13.1957525392428,0.423667100130039,3,6,negative regulation of protein oligomerization,0.00732400613443852

GO:0050793,0.0059898252160373,1.25939750511644,136.491417425228,164,1933,regulation of developmental process,0.00732400613443852

GO:0051346,0.00600039851812761,1.80155267295597,15.3932379713914,26,218,negative regulation of hydrolase activity,0.00732400613443852

GO:0071706,0.0060238237824601,2.52389819987585,5.29583875162549,12,75,tumor necrosis factor superfamily cytokine production,0.00732400613443852

GO:0006790,0.00603332910588082,1.77864415336718,16.1699609882965,27,229,sulfur compound metabolic process,0.00732400613443852

GO:0048247,0.0060592650624471,3.70279888785913,2.25955786736021,7,32,lymphocyte chemotaxis,0.00732400613443852

GO:0034110,0.0060592650624471,3.70279888785913,2.25955786736021,7,32,regulation of homotypic cell-cell adhesion,0.00732400613443852

GO:0097530,0.00607565741516741,2.32038246268657,6.63745123537061,14,94,granulocyte migration,0.00732400613443852

GO:0007589,0.00608390360502639,2.81717155738353,4.02483745123537,10,57,body fluid secretion,0.00732400613443852

GO:0061462,0.00608816140482634,3.30751391465677,2.82444733420026,8,40,protein localization to lysosome,0.00732400613443852

GO:0002448,0.00608816140482634,3.30751391465677,2.82444733420026,8,40,mast cell mediated immunity,0.00732400613443852

GO:0045926,0.00611800001877539,1.87577119969527,13.1336801040312,23,186,negative regulation of growth,0.00734420897776236

GO:0070373,0.00613486953999546,2.64911627906977,4.66033810143043,11,66,negative regulation of ERK1 and ERK2 cascade,0.00734879053407967

GO:0048523,0.00622829751596934,1.19788886397801,265.99232769831,301,3767,negative regulation of cellular process,0.00744486518363214

GO:0050777,0.00629482002969195,2.16608811748999,8.04967490247074,16,114,negative regulation of immune response,0.00750843999304357

GO:0071216,0.00638922582283381,1.96775762629421,10.944733420026,20,155,cellular response to biotic stimulus,0.00759197871272673

GO:0072659,0.00639182577234897,1.74954959752212,17.0172951885566,28,241,protein localization to plasma membrane,0.00759197871272673

GO:0030595,0.00656708771685505,2.00229095074456,10.2386215864759,19,145,leukocyte chemotaxis,0.00778372712545136

GO:0002886,0.00666793578672371,2.97785515320334,3.45994798439532,9,49,regulation of myeloid leukocyte mediated immunity,0.00788665514270052

GO:0071621,0.00670606230538487,2.48428770949721,5.36644993498049,12,76,granulocyte chemotaxis,0.0079151217566702

GO:0001959,0.00680848168081092,2.08885077781242,8.82639791937581,17,125,regulation of cytokine-mediated signaling pathway,0.00796560407188707

GO:0048878,0.00683469401102669,1.40825599511501,51.2637191157347,69,726,chemical homeostasis,0.00796560407188707

GO:0043031,0.00686202126974286,5.08112146872554,1.27100130039012,5,18,negative regulation of macrophage activation,0.00796560407188707

GO:1903715,0.00686202126974286,5.08112146872554,1.27100130039012,5,18,regulation of aerobic respiration,0.00796560407188707

GO:0034114,0.00686202126974286,5.08112146872554,1.27100130039012,5,18,regulation of heterotypic cell-cell adhesion,0.00796560407188707

GO:0035767,0.00686202126974286,5.08112146872554,1.27100130039012,5,18,endothelial cell chemotaxis,0.00796560407188707

GO:0002823,0.00686202126974286,5.08112146872554,1.27100130039012,5,18,negative regulation of adaptive immune response based on somatic recombination of immune receptors built from immunoglobulin superfamily domains,0.00796560407188707

GO:0042219,0.00686202126974286,5.08112146872554,1.27100130039012,5,18,cellular modified amino acid catabolic process,0.00796560407188707

GO:0055080,0.00687778684987859,1.53303828736105,30.8570871261378,45,437,monoatomic cation homeostasis,0.00796747735901573

GO:0040012,0.00691898189272153,1.40045494059693,53.0289986996099,71,751,regulation of locomotion,0.00799874087392653

GO:0050801,0.00693857952276692,1.524589616339,31.7044213263979,46,449,monoatomic ion homeostasis,0.00800495957237249

GO:0010876,0.0069658169238811,1.68495599187542,19.4886866059818,31,276,lipid localization,0.008019948728313

GO:0062197,0.00705398604033996,1.82327683615819,14.0516254876463,24,199,cellular response to chemical stress,0.00810488600145183

GO:0070661,0.00711913332862695,1.79700435591105,14.8283485045514,25,210,leukocyte proliferation,0.00816307956011604

GO:0046649,0.0071359726237436,1.47656905453372,37.6357607282185,53,533,lymphocyte activation,0.00816575729099116

GO:0090218,0.00723171149869207,3.56013402723319,2.33016905071521,7,33,positive regulation of lipid kinase activity,0.00822515873487603

GO:1990874,0.00723171149869207,3.56013402723319,2.33016905071521,7,33,vascular associated smooth muscle cell proliferation,0.00822515873487603

GO:1904705,0.00723171149869207,3.56013402723319,2.33016905071521,7,33,regulation of vascular associated smooth muscle cell proliferation,0.00822515873487603

GO:0006793,0.00727134466136445,1.24623584042778,143.693758127438,171,2035,phosphorus metabolic process,0.00825356258941166

GO:0006959,0.00732238364792889,2.26346923917,6.77867360208062,14,96,humoral immune response,0.00829477262330778

GO:0050830,0.0076348064467471,2.90502072151641,3.53055916775032,9,50,defense response to Gram-positive bacterium,0.00853695549891331

GO:0043112,0.0076348064467471,2.90502072151641,3.53055916775032,9,50,receptor metabolic process,0.00853695549891331

GO:0031400,0.00770644146876377,1.5834088740458,25.2788036410923,38,358,negative regulation of protein modification process,0.00853695549891331

GO:1903541,0.00773329894217724,6.60166358595194,0.847334200260078,4,12,regulation of exosomal secretion,0.00853695549891331

GO:1903543,0.00773329894217724,6.60166358595194,0.847334200260078,4,12,positive regulation of exosomal secretion,0.00853695549891331

GO:0061549,0.00773329894217724,6.60166358595194,0.847334200260078,4,12,sympathetic ganglion development,0.00853695549891331

GO:0034350,0.00773329894217724,6.60166358595194,0.847334200260078,4,12,regulation of glial cell apoptotic process,0.00853695549891331

GO:0048925,0.00773329894217724,6.60166358595194,0.847334200260078,4,12,lateral line system development,0.00853695549891331

GO:0002468,0.00773329894217724,6.60166358595194,0.847334200260078,4,12,dendritic cell antigen processing and presentation,0.00853695549891331

GO:0031664,0.00773329894217724,6.60166358595194,0.847334200260078,4,12,regulation of lipopolysaccharide-mediated signaling pathway,0.00853695549891331

GO:0006590,0.00773329894217724,6.60166358595194,0.847334200260078,4,12,thyroid hormone generation,0.00853695549891331

GO:0002604,0.00773329894217724,6.60166358595194,0.847334200260078,4,12,regulation of dendritic cell antigen processing and presentation,0.00853695549891331

GO:0070255,0.00773329894217724,6.60166358595194,0.847334200260078,4,12,regulation of mucus secretion,0.00853695549891331

GO:0043124,0.00779389294912027,2.70180562931492,4.16605981794538,10,59,negative regulation of canonical NF-kappaB signal transduction,0.00858700925705423

GO:0051928,0.00781341000426989,2.3281529432508,6.14317295188557,13,87,positive regulation of calcium ion transport,0.00859169889141396

GO:0007015,0.00784188571617024,1.54391881194389,28.5975292587776,42,405,actin filament organization,0.00860620206277552

GO:0032943,0.00796167943295561,1.8024194835456,14.1928478543563,24,201,mononuclear cell proliferation,0.00870874020508088

GO:0042110,0.00796625436166368,1.56005054062818,26.9734720416125,40,382,T cell activation,0.00870874020508088

GO:0051251,0.00800027960569269,1.73506589481463,16.5230169050715,27,234,positive regulation of lymphocyte activation,0.00872898724419571

GO:0006081,0.00824838301203696,3.11251773436647,2.96566970091027,8,42,cellular aldehyde metabolic process,0.0089347862012396

GO:0002495,0.00824838301203696,3.11251773436647,2.96566970091027,8,42,antigen processing and presentation of peptide antigen via MHC class II,0.0089347862012396

GO:0019886,0.00824838301203696,3.11251773436647,2.96566970091027,8,42,antigen processing and presentation of exogenous peptide antigen via MHC class II,0.0089347862012396

GO:0043648,0.00825237801890692,2.4086676824107,5.50767230169051,12,78,dicarboxylic acid metabolic process,0.0089347862012396

GO:0030003,0.00830985827296564,1.55539079876693,27.0440832249675,40,383,intracellular monoatomic cation homeostasis,0.00897975087846383

GO:0002819,0.00839799141406281,1.99073033707865,9.7443433029909,18,138,regulation of adaptive immune response,0.00905760376650836

GO:1903532,0.00845093544153445,1.72660165613423,16.5936280884265,27,235,positive regulation of secretion by cell,0.00909727849633632

GO:0009893,0.00852659133428536,1.21780006176328,180.835240572172,210,2561,positive regulation of metabolic process,0.00916120404809668

GO:0032941,0.00856395249603342,3.42803693406103,2.40078023407022,7,34,secretion by tissue,0.00918381953384155

GO:0060429,0.00864425761072532,1.35101000030212,64.1149544863459,83,908,epithelium development,0.00923612111415867

GO:0031348,0.00864553432888387,1.8719160601191,12.0039011703511,21,170,negative regulation of defense response,0.00923612111415867

GO:0030888,0.00870497756448338,2.83565459610028,3.60117035110533,9,51,regulation of B cell proliferation,0.00928201206212906

GO:0060713,0.00878030723903456,4.71785383903793,1.34161248374512,5,19,labyrinthine layer morphogenesis,0.0093270056142952

GO:0050869,0.00878030723903456,4.71785383903793,1.34161248374512,5,19,negative regulation of B cell activation,0.0093270056142952

GO:0002700,0.00880398847325881,2.13864040239752,7.6260078023407,15,108,regulation of production of molecular mediator of immune response,0.00933454898388834

GO:0019216,0.00885197372340765,1.70089166256657,17.4409622886866,28,247,regulation of lipid metabolic process,0.00935461983378049

GO:0030036,0.00885614986039965,1.42362407200033,44.0613784135241,60,624,actin cytoskeleton organization,0.00935461983378049

GO:0043588,0.00890422454065384,1.80840759934761,13.5573472041612,23,192,skin development,0.00937166220581955

GO:0051258,0.00892222192241435,1.71821841495317,16.6642392717815,27,236,protein polymerization,0.00937166220581955

GO:0046777,0.00892222192241435,1.71821841495317,16.6642392717815,27,236,protein autophosphorylation,0.00937166220581955

GO:0044092,0.00906766303962513,1.4017153464839,48.4392717815345,65,686,negative regulation of molecular function,0.00950669327990494

GO:0009056,0.00911050123119879,1.24174257646391,138.044863459038,164,1955,catabolic process,0.00952956834225955

GO:0050729,0.00912333452305132,2.37255065454849,5.57828348504551,12,79,positive regulation of inflammatory response,0.00952956834225955

GO:0048519,0.00916116382856759,1.18245159610129,283.292067620286,317,4012,negative regulation of biological process,0.00955136154719176

GO:0043269,0.00922559882310477,1.52665288609101,28.8799739921977,42,409,regulation of monoatomic ion transport,0.00960076180666911

GO:1903039,0.00936509907957644,1.8265272556391,12.8512353706112,22,182,positive regulation of leukocyte cell-cell adhesion,0.00970058795226108

GO:0008154,0.00936509907957644,1.8265272556391,12.8512353706112,22,182,actin polymerization or depolymerization,0.00970058795226108

GO:0009913,0.00937321464658975,1.92518797799352,10.591677503251,19,150,epidermal cell differentiation,0.00970058795226108

GO:0006873,0.00941562125107664,1.54157318272748,27.2559167750325,40,386,intracellular monoatomic ion homeostasis,0.0097248567641268

GO:0006906,0.00943121100037875,2.26656692990631,6.28439531859558,13,89,vesicle fusion,0.0097248567641268

GO:0007162,0.00947489717368663,1.72850943396226,15.9581274382315,26,226,negative regulation of cell adhesion,0.00972961866688507

GO:0044282,0.00947489717368663,1.72850943396226,15.9581274382315,26,226,small molecule catabolic process,0.00972961866688507

GO:0006796,0.00950509461430572,1.23713213117314,141.928478543563,168,2010,phosphate-containing compound metabolic process,0.00972961866688507

GO:0071357,0.00952223780720013,3.02337662337662,3.03628088426528,8,43,cellular response to type I interferon,0.00972961866688507

GO:0060337,0.00952223780720013,3.02337662337662,3.03628088426528,8,43,type I interferon-mediated signaling pathway,0.00972961866688507

GO:0050764,0.00956541284542134,2.4688214426488,4.94278283485045,11,70,regulation of phagocytosis,0.0097560279564714

GO:0070727,0.00978706789690553,1.2345478432436,143.905591677503,170,2038,cellular macromolecule localization,0.009922189075426

GO:0046173,0.00985201470782669,2.59548800933013,4.3072821846554,10,61,polyol biosynthetic process,0.009922189075426

GO:0048671,0.0099045652937645,9.89612188365651,0.494278283485045,3,7,negative regulation of collateral sprouting,0.009922189075426

GO:0009265,0.0099045652937645,9.89612188365651,0.494278283485045,3,7,2'-deoxyribonucleotide biosynthetic process,0.009922189075426

GO:1901725,0.0099045652937645,9.89612188365651,0.494278283485045,3,7,regulation of histone deacetylase activity,0.009922189075426

GO:0021877,0.0099045652937645,9.89612188365651,0.494278283485045,3,7,forebrain neuron fate commitment,0.009922189075426

GO:0021902,0.0099045652937645,9.89612188365651,0.494278283485045,3,7,commitment of neuronal cell to specific neuron type in forebrain,0.009922189075426

GO:0046385,0.0099045652937645,9.89612188365651,0.494278283485045,3,7,deoxyribose phosphate biosynthetic process,0.009922189075426

GO:0036446,0.0099045652937645,9.89612188365651,0.494278283485045,3,7,myofibroblast differentiation,0.009922189075426

GO:0106070,0.0099045652937645,9.89612188365651,0.494278283485045,3,7,regulation of adenylate cyclase-activating G protein-coupled receptor signaling pathway,0.009922189075426

GO:0046883,0.00995858882100519,1.81505393919582,12.9218465539662,22,183,regulation of hormone secretion,0.00995858882100519

Table S3. Supplementary data to accompany Figure 3D. Top gene ontology (Functional Enrichment Analysis) for each compartment (ET).

GOBPID,Pvalue,OddsRatio,ExpCount,Count,Size,Term,FDR

GO:0009653,6.11330342322517e-33,2.69152046783626,129.395838751625,263,2372,anatomical structure morphogenesis,5.63035245279039e-30

GO:0007275,8.04517857189509e-32,2.36272408020514,229.715799739922,384,4211,multicellular organism development,3.70480473235769e-29

GO:0001501,1.81732953616855e-31,5.30368747718145,23.0206762028609,92,422,skeletal system development,5.57920167603744e-29

GO:0048731,1.73517695787725e-30,2.41540435386331,178.764824447334,320,3277,system development,3.99524494551237e-28

GO:0048856,5.42317315473655e-30,2.26344012874122,272.429518855657,427,4994,anatomical structure development,9.98948495102472e-28

GO:0032502,1.96285577021828e-29,2.23700637987178,289.504096228869,444,5307,developmental process,3.01298360728506e-27

GO:0048513,3.75017005922132e-28,2.4843303242123,134.360013003901,258,2463,animal organ development,4.93415232077548e-26

GO:0032501,1.7254529166209e-26,2.13426197558519,318.30721716515,466,5835,multicellular organismal process,1.98642767025981e-24

GO:0001503,3.8215193959364e-26,5.37634342971949,18.0565019505852,74,331,ossification,3.91068818184158e-24

GO:0009887,4.92009835438214e-25,3.23483779971791,49.6417425227568,130,910,animal organ morphogenesis,4.53141058438595e-23

GO:0030198,2.27248910674061e-24,5.41598615030922,16.4199609882965,68,301,extracellular matrix organization,1.90269315209828e-22

GO:0043062,2.78558733058687e-24,5.39246399946789,16.4745123537061,68,302,extracellular structure organization,2.13793827622542e-22

GO:0035295,6.52912599402998e-24,3.17264937723195,49.1507802340702,127,901,tube development,4.6256346465397e-22

GO:0048646,1.61912894373283e-23,3.03492473331844,54.0604031209363,134,991,anatomical structure formation involved in morphogenesis,1.06515554084138e-21

GO:0009888,3.52011161048479e-23,2.57587193746791,86.4093628088427,181,1584,tissue development,2.16134852883766e-21

GO:0035239,7.40774689579735e-23,3.30500796117977,41.8408972691808,113,767,tube morphogenesis,4.26408430689335e-21

GO:0051216,4.53533160857077e-22,6.86507142380666,9.98289986996099,50,183,cartilage development,2.45708259499628e-20

GO:0048705,5.40192670701963e-22,6.34106670658085,11.2375812743823,53,206,skeletal system morphogenesis,2.76398583175838e-20

GO:0045229,6.90547871785184e-22,4.788132542301,18.0565019505852,68,331,external encapsulating structure organization,3.3473399469166e-20

GO:0050793,1.32452534805716e-21,2.36361017195623,105.447789336801,203,1933,regulation of developmental process,6.09943922780324e-20

GO:0060348,1.67292166527691e-21,6.6140408883011,10.2556566970091,50,188,bone development,7.3369564462859e-20

GO:0030154,2.50013174749459e-21,2.08016749129735,184.547269180754,301,3383,cell differentiation,1.0011397128011e-19

GO:0048869,2.50013174749459e-21,2.08016749129735,184.547269180754,301,3383,cellular developmental process,1.0011397128011e-19

GO:0061448,1.13151095465136e-20,5.59704506802721,12.8195708712614,55,235,connective tissue development,4.3421732884746e-19

GO:0001568,1.25292119816905e-20,3.42923464343097,33.6581924577373,95,617,blood vessel development,4.61576169405477e-19

GO:0001944,2.92721475210721e-20,3.33177205308353,35.2401820546164,97,646,vasculature development,1.03690953334259e-18

GO:0007155,1.26134995588351e-19,2.48185373989852,76.9719765929779,159,1411,cell adhesion,4.30260484951376e-18

GO:0001649,5.79118074188698e-19,6.71758577369607,8.67366710013004,43,159,osteoblast differentiation,1.90488480831354e-17

GO:0016477,7.91246941683873e-19,2.57780082417582,64.3160598179454,139,1179,cell migration,2.51289114927878e-17

GO:0072359,3.20910840576163e-18,2.66293808493297,55.7514954486346,125,1022,circulatory system development,9.85196280568821e-17

GO:0022603,4.21199578255813e-18,2.91594205802212,42.822821846554,105,785,regulation of anatomical structure morphogenesis,1.25137035991485e-16

GO:0060485,5.73069330398466e-18,5.23250732048424,12.2195058517555,50,224,mesenchyme development,1.64936516655308e-16

GO:0048762,8.11676518613311e-18,5.6800061764933,10.5284135240572,46,193,mesenchymal cell differentiation,2.26531537467533e-16

GO:0007399,1.05999375033954e-17,2.14740897798086,113.084980494148,202,2073,nervous system development,2.87133601194917e-16

GO:0045595,1.31013020878314e-17,2.51279924609823,63.6068920676203,135,1166,regulation of cell differentiation,3.44751406368364e-16

GO:0002062,1.40602014951152e-17,8.71422055174427,5.50968790637191,33,101,chondrocyte differentiation,3.59706821583363e-16

GO:0048514,3.44201898984655e-17,3.27902160480675,29.8395968790637,82,547,blood vessel morphogenesis,8.56783645850993e-16

GO:2000026,5.03221638260985e-17,2.51693737501618,60.4429128738622,129,1108,regulation of multicellular organismal development,1.21965033904834e-15

GO:0048699,1.06334209887661e-16,2.37006111107692,71.6259427828349,144,1313,generation of neurons,2.51112326427015e-15

GO:0048870,1.41809489760825e-16,2.37454813804781,70.4258127438231,142,1291,cell motility,3.26516350174301e-15

GO:0060349,2.76711419402565e-16,8.41443519351935,5.29148244473342,31,97,bone morphogenesis,6.21588334804299e-15

GO:0007178,2.94835150365728e-16,4.4097389243264,14.8379713914174,53,272,transmembrane receptor protein serine/threonine kinase signaling pathway,6.46531365444846e-15

GO:0045597,6.31815374692876e-16,2.95840443788547,35.0219765929779,88,642,positive regulation of cell differentiation,1.3532603723073e-14

GO:0022008,8.67600120123917e-16,2.23426533722906,81.4997399219766,155,1494,neurogenesis,1.81604479689574e-14

GO:0048729,2.49755768436614e-15,3.00455302551678,32.0762028608583,82,588,tissue morphogenesis,5.11166806066937e-14

GO:0051094,2.97397538364489e-15,2.48168180586427,54.4968140442133,116,999,positive regulation of developmental process,5.95441593116726e-14

GO:0060350,3.47972594235203e-15,10.6024043524044,3.65494148244473,25,67,endochondral bone morphogenesis,6.81878211256642e-14

GO:0030182,3.5612671261229e-15,2.31354141141877,68.0801040312094,135,1248,neuron differentiation,6.83318129824832e-14

GO:0042127,3.79623652210988e-15,2.15378348121314,88.1550065019506,162,1616,regulation of cell population proliferation,7.13537517727184e-14

GO:0001525,4.55954880117733e-15,3.32144896987829,24.6026657997399,69,451,angiogenesis,8.39868889176864e-14

GO:0060429,6.02983730440829e-15,2.53766757865937,49.5326397919376,108,908,epithelium development,1.08891767791373e-13

GO:0051093,8.6875010982519e-15,2.84608446783277,35.294733420026,86,647,negative regulation of developmental process,1.53869009836346e-13

GO:0051241,1.50633510893141e-14,2.67918231635983,40.8044213263979,94,748,negative regulation of multicellular organismal process,2.61761251948269e-13

GO:0008283,1.66983009372681e-14,1.90673508986928,141.560793237971,227,2595,cell population proliferation,2.84798799318961e-13

GO:0051239,2.36491055784485e-14,1.97639570688415,116.685370611183,196,2139,regulation of multicellular organismal process,3.89061452147698e-13

GO:0002009,2.36562880784702e-14,3.22569098824641,24.8208712613784,68,455,morphogenesis of an epithelium,3.89061452147698e-13

GO:0048598,6.69870812434626e-14,3.00483642034801,28.3667100130039,73,520,embryonic morphogenesis,1.08237020746016e-12

GO:0000902,2.69577371663343e-13,2.37081323982468,52.4238621586476,108,961,cell morphogenesis,4.28070274658515e-12

GO:0045669,3.22386245902628e-13,8.70935269813447,3.98224967490247,24,73,positive regulation of osteoblast differentiation,5.03250394027661e-12

GO:0009790,4.20857098912624e-13,2.38136061215312,50.6782184655397,105,929,embryo development,6.46015646830877e-12

GO:0031214,4.95979803385108e-13,5.96039603960396,6.76436931079324,31,124,biomineral tissue development,7.48848194947024e-12

GO:0045667,6.54584547447838e-13,6.39140374942844,6.00065019505852,29,110,regulation of osteoblast differentiation,9.72374787418481e-12

GO:0007166,7.20365671934266e-13,1.89589587251157,119.194733420026,194,2185,cell surface receptor signaling pathway,1.05310600611343e-11

GO:0031589,8.34544253611801e-13,3.45329685117072,18.4383615084525,54,338,cell-substrate adhesion,1.19150678788795e-11

GO:0001837,8.40911413818854e-13,6.31302318578741,6.05520156046814,29,111,epithelial to mesenchymal transition,1.19150678788795e-11

GO:0030111,1.96911185234704e-12,3.74203488630801,14.9470741222367,47,274,regulation of Wnt signaling pathway,2.7478060848661e-11

GO:0043009,2.76533650611918e-12,2.86681370175785,27.3847854356307,68,502,chordate embryonic development,3.80130585393398e-11

GO:0070848,2.86319183463065e-12,2.72772659264509,31.2033810143043,74,572,response to growth factor,3.87794070543357e-11

GO:0010717,4.53484871108736e-12,8.48517907508914,3.70949284785436,22,68,regulation of epithelial to mesenchymal transition,6.0530371926253e-11

GO:0060562,4.99256063888933e-12,3.62831844081844,15.328933680104,47,281,epithelial tube morphogenesis,6.56878335488153e-11

GO:0045165,6.00574217784133e-12,4.54601582095312,9.71014304291287,36,178,cell fate commitment,7.68583503068888e-11

GO:0009792,6.0084703822975e-12,2.78682377225424,28.4758127438231,69,522,embryo development ending in birth or egg hatching,7.68583503068888e-11

GO:0090287,1.09173202992876e-11,4.12109558823529,11.4012353706112,39,209,regulation of cellular response to growth factor stimulus,1.37737698570465e-10

GO:0016055,1.12042578193005e-11,3.081555991962,21.4386866059818,57,393,Wnt signaling pathway,1.3847051495098e-10

GO:0042476,1.12761005660407e-11,6.33823653305026,5.40058517555267,26,99,odontogenesis,1.3847051495098e-10

GO:0001763,1.17225049782517e-11,4.54345122215012,9.43738621586476,35,173,morphogenesis of a branching structure,1.42058251118024e-10

GO:0007389,1.35320798917103e-11,3.13769493646515,20.3476592977893,55,373,pattern specification process,1.61857734808639e-10

GO:0008284,1.6759506529169e-11,2.27382760235483,49.478088426528,99,907,positive regulation of cell population proliferation,1.97891096325188e-10

GO:0007423,1.83833033518931e-11,2.7304285856815,28.5303641092328,68,523,sensory organ development,2.11187478865839e-10

GO:0040007,1.84359055806143e-11,2.34358923296492,44.6230169050715,92,818,growth,2.11187478865839e-10

GO:0198738,1.87719865172661e-11,3.03530311780456,21.7114434330299,57,398,cell-cell signaling by wnt,2.11187478865839e-10

GO:0071363,1.88027939923983e-11,2.70657990078591,29.1849804941482,69,535,cellular response to growth factor stimulus,2.11187478865839e-10

GO:0010718,2.00633012033245e-11,12.0083698296837,2.29115734720416,17,42,positive regulation of epithelial to mesenchymal transition,2.22630125400746e-10

GO:0048568,2.83333837292221e-11,3.03286631974228,21.3295838751625,56,391,embryonic organ development,3.10655314459685e-10

GO:0048589,3.73702366257955e-11,2.57991421111526,32.2398569570871,73,591,developmental growth,4.04917505086561e-10

GO:0030278,4.77192041110409e-11,5.85442259485886,5.7278933680104,26,105,regulation of ossification,5.11039383561264e-10

GO:0007167,4.96494789062347e-11,2.30498888668823,44.7321196358908,91,820,enzyme-linked receptor protein signaling pathway,5.25599656007381e-10

GO:0085029,6.16992535607053e-11,12.2715410217127,2.12750325097529,16,39,extracellular matrix assembly,6.45738778743291e-10

GO:0030334,6.36728785134928e-11,2.43209521067281,37.3131339401821,80,684,regulation of cell migration,6.58906978774459e-10

GO:0061035,7.74828366961729e-11,9.63879968995682,2.78211963589077,18,51,regulation of cartilage development,7.92907695524169e-10

GO:0040011,1.13256668071651e-10,2.222582346536,48.2779583875163,95,885,locomotion,1.13602072939388e-09

GO:0030282,1.13478726497543e-10,5.84548364153627,5.50968790637191,25,101,bone mineralization,1.13602072939388e-09

GO:0048754,1.35811074978993e-10,5.08825083670953,6.87347204161248,28,126,branching morphogenesis of an epithelial tube,1.3449677425339e-09

GO:0060173,1.44461236468875e-10,4.30468566046621,9.27373211963589,33,170,limb development,1.41541275306206e-09

GO:0070167,1.9390502214802e-10,6.27730015082956,4.80052015604681,23,88,regulation of biomineral tissue development,1.87985816208765e-09

GO:0040012,1.9907297173907e-10,2.31423890703655,40.9680754226268,84,751,regulation of locomotion,1.9098563226217e-09

GO:2000145,2.02472707368563e-10,2.35176889549447,38.8951235370611,81,713,regulation of cell motility,1.92244704625203e-09

GO:0003170,2.45232542134197e-10,9.68016639196295,2.6184655396619,17,48,heart valve development,2.30468542148567e-09

GO:0048736,3.20407135817306e-10,4.15167056932164,9.54648894668401,33,175,appendage development,2.98075729381554e-09

GO:0061138,3.34869574269853e-10,4.3892916077322,8.56456436931079,31,157,morphogenesis of a branching epithelium,3.08414877902535e-09

GO:0048562,3.5976162538352e-10,3.4561050513608,13.8560468140442,41,254,embryonic organ morphogenesis,3.28059858394279e-09

GO:0048468,3.76091554260933e-10,1.74587594486438,123.558842652796,189,2265,cell development,3.39588550465019e-09

GO:0030509,4.14939660176681e-10,5.19302900894177,6.27340702210663,26,115,BMP signaling pathway,3.71028569924974e-09

GO:0007411,6.05590630489867e-10,3.44974487339944,13.5287386215865,40,248,axon guidance,5.31189495886826e-09

GO:0097485,6.05590630489867e-10,3.44974487339944,13.5287386215865,40,248,neuron projection guidance,5.31189495886826e-09

GO:0008285,6.51842543679772e-10,2.42966307066917,32.9490247074122,71,604,negative regulation of cell population proliferation,5.65767131779267e-09

GO:0032963,6.57297319222383e-10,6.5234847509973,4.25500650195058,21,78,collagen metabolic process,5.65767131779267e-09

GO:0010975,6.85373769754061e-10,2.7236636997348,23.7843953185956,57,436,regulation of neuron projection development,5.84471520318046e-09

GO:0060070,7.11213496784685e-10,3.56426195004935,12.4922626788036,38,229,canonical Wnt signaling pathway,6.00942780310729e-09

GO:0045596,7.8467798112853e-10,2.66041905855339,25.1481794538361,59,461,negative regulation of cell differentiation,6.56989473290342e-09

GO:0010171,8.69775408531014e-10,7.94817295980512,3.16397919375813,18,58,body morphogenesis,7.21678514646003e-09

GO:0120035,9.95699908186523e-10,2.43563122923588,31.9125487646294,69,585,regulation of plasma membrane bounded cell projection organization,8.18785370928382e-09

GO:0071772,1.10965924702422e-09,4.91510298081704,6.54616384915475,26,120,response to BMP,8.94525125423162e-09

GO:0071773,1.10965924702422e-09,4.91510298081704,6.54616384915475,26,120,cellular response to BMP stimulus,8.94525125423162e-09

GO:0031344,1.11694233901915e-09,2.41110518410722,32.6762678803641,70,599,regulation of cell projection organization,8.94525125423162e-09

GO:0070169,1.18851707366819e-09,7.75378033926502,3.21853055916775,18,59,positive regulation of biomineral tissue development,9.43641573145175e-09

GO:0009719,1.52413067792628e-09,1.92771480516881,70.3712613784135,121,1290,response to endogenous stimulus,1.19976440544454e-08

GO:0035108,1.72143882307142e-09,4.61584643804471,7.1462288686606,27,131,limb morphogenesis,1.34359758987184e-08

GO:0006029,1.91742397898607e-09,5.7312621499028,4.90962288686606,22,90,proteoglycan metabolic process,1.48398948289594e-08

GO:0051960,2.2472386338382e-09,2.79578171045932,20.7295188556567,51,380,regulation of nervous system development,1.71693041504397e-08

GO:0071559,2.26149631648936e-09,3.70260566580821,10.8011703511053,34,198,response to transforming growth factor beta,1.71693041504397e-08

GO:0001823,2.27432693415162e-09,5.9953269185267,4.5277633289987,21,83,mesonephros development,1.71693041504397e-08

GO:0072073,2.33880722135201e-09,4.71318121344479,6.76436931079324,26,124,kidney epithelium development,1.75125321208553e-08

GO:0001657,2.59919170192629e-09,6.3165009593581,4.14590377113134,20,76,ureteric bud development,1.93052867538235e-08

GO:0048666,2.68910003891537e-09,2.01116399715047,57.2243823146944,103,1049,neuron development,1.98132890867284e-08

GO:0030500,3.33365823558683e-09,6.20525673157252,4.20045513654096,20,77,regulation of bone mineralization,2.43674542458371e-08

GO:0003179,3.58903282740256e-09,9.43546289875173,2.34570871261378,15,43,heart valve morphogenesis,2.582421276592e-08

GO:0060351,3.58903282740256e-09,9.43546289875173,2.34570871261378,15,43,cartilage development involved in endochondral bone morphogenesis,2.582421276592e-08

GO:0045778,3.8635331575431e-09,7.0626065773447,3.43673602080624,18,63,positive regulation of ossification,2.75838297526914e-08

GO:0098868,5.17290769581954e-09,9.10947438901908,2.40026007802341,15,44,bone growth,3.66480614449984e-08

GO:0072163,5.41132224547252e-09,5.99408124831854,4.30955786736021,20,79,mesonephric epithelium development,3.77562711218196e-08

GO:0072164,5.41132224547252e-09,5.99408124831854,4.30955786736021,20,79,mesonephric tubule development,3.77562711218196e-08

GO:0035107,5.69773481381165e-09,4.32266009852217,7.52808842652796,27,138,appendage morphogenesis,3.93838786028606e-08

GO:0048522,5.73011914525876e-09,1.54579378144168,228.079258777633,302,4181,positive regulation of cellular process,3.93838786028606e-08

GO:0007409,7.61099822542104e-09,2.54805035209795,24.7117685305592,56,453,axonogenesis,5.19239212267613e-08

GO:0090092,7.77405513122591e-09,3.87031762240017,9.16462938881664,30,168,regulation of transmembrane receptor protein serine/threonine kinase signaling pathway,5.26463586460225e-08

GO:0031175,8.5363554760259e-09,2.01632304005563,52.4238621586476,95,961,neuron projection development,5.73867400979551e-08

GO:0007179,8.90932840662368e-09,4.08049160114006,8.18270481144343,28,150,transforming growth factor beta receptor signaling pathway,5.94600830615972e-08

GO:0030855,1.31764400873219e-08,2.50266176042892,25.0936280884265,56,460,epithelial cell differentiation,8.73057648951331e-08

GO:0060560,1.48478668647121e-08,3.32350878585794,12.1649544863459,35,223,developmental growth involved in morphogenesis,9.76777527314272e-08

GO:0061564,1.60405055075026e-08,2.44291431999499,26.5665149544863,58,487,axon development,1.04431375166506e-07

GO:0060828,1.61012543687772e-08,3.54168109784725,10.5284135240572,32,193,regulation of canonical Wnt signaling pathway,1.04431375166506e-07

GO:0007507,1.70561438734656e-08,2.34287876542687,30.0032509752926,63,550,heart development,1.09851108443789e-07

GO:0048812,1.78063775653287e-08,2.24555927943025,34.2037061118335,69,627,neuron projection morphogenesis,1.13838242009722e-07

GO:0120039,1.79224159515849e-08,2.23112723290671,34.9128738621586,70,640,plasma membrane bounded cell projection morphogenesis,1.13838242009722e-07

GO:0048858,2.03432619381784e-08,2.22300781143434,35.0219765929779,70,642,cell projection morphogenesis,1.28329755103166e-07

GO:0030199,2.0747363418267e-08,6.66212489862125,3.38218465539662,17,62,collagen fibril organization,1.29988583049142e-07

GO:0030501,2.13497069557998e-08,7.22908683054491,3.00032509752926,16,55,positive regulation of bone mineralization,1.32858649366835e-07

GO:2000027,2.30388162981799e-08,4.86753365973072,5.56423927178153,22,102,regulation of animal organ morphogenesis,1.42407716849824e-07

GO:0003002,2.34258126370616e-08,2.76562661743901,18.3838101430429,45,337,regionalization,1.43834489591558e-07

GO:0071560,2.3788521479734e-08,3.56089108910891,10.1465539661899,31,186,cellular response to transforming growth factor beta stimulus,1.45094227038642e-07

GO:0003417,2.47573210186313e-08,9.93441414885777,1.96384915474642,13,36,growth plate cartilage development,1.5000982011947e-07

GO:0050767,2.76404618724537e-08,2.82329228560384,17.2382314694408,43,316,regulation of neurogenesis,1.66384741075359e-07

GO:0023052,2.81558185753934e-08,1.50093213682776,252.845578673602,325,4635,signaling,1.68386421480113e-07

GO:0001704,3.03781275135607e-08,5.01894865525672,5.18237971391417,21,95,formation of primary germ layer,1.80504873806384e-07

GO:0060572,3.08909471326768e-08,34.9867629362214,0.654616384915475,8,12,morphogenesis of an epithelial bud,1.82375399417919e-07

GO:0016525,3.20197214748154e-08,5.27545423067811,4.74596879063719,20,87,negative regulation of angiogenesis,1.87835436167548e-07

GO:0003416,3.30337465933555e-08,8.49187042842215,2.34570871261378,14,43,endochondral bone growth,1.92557472230889e-07

GO:0007154,3.40894240699704e-08,1.49441595848084,258.90078023407,331,4746,cell communication,1.97461380933602e-07

GO:0042475,3.49020096883193e-08,6.37775016824559,3.49128738621586,17,64,odontogenesis of dentin-containing tooth,2.00306660007118e-07

GO:0072001,3.51909466573131e-08,2.87840188707062,16.1472041612484,41,296,renal system development,2.00306660007118e-07

GO:0030326,3.52330932911543e-08,4.52577614379085,6.16430429128739,23,113,embryonic limb morphogenesis,2.00306660007118e-07

GO:0051240,3.59207284613555e-08,1.89704106280193,60.1156046814044,103,1102,positive regulation of multicellular organismal process,2.02963134435021e-07

GO:0072132,3.62725396809037e-08,9.51982445520581,2.01840052015605,13,37,mesenchyme morphogenesis,2.03701274671416e-07

GO:0032990,4.02852834764642e-08,2.17934677144138,35.6220416124837,70,653,cell part morphogenesis,2.24865127768627e-07

GO:0071495,4.1252597164156e-08,1.89094230018224,60.2792587776333,103,1105,cellular response to endogenous stimulus,2.28877361374624e-07

GO:0007165,4.21095389807309e-08,1.50310925137586,234.189011703511,304,4293,signal transduction,2.32232846714091e-07

GO:0009966,4.3012169862302e-08,1.62495461070166,127.704746423927,185,2341,regulation of signal transduction,2.35798859780834e-07

GO:0048706,4.47356553109847e-08,4.88619547033844,5.29148244473342,21,97,embryonic skeletal system development,2.43796086043887e-07

GO:2000181,4.81859826791183e-08,5.12183468705208,4.85507152145644,20,89,negative regulation of blood vessel morphogenesis,2.59528011973497e-07

GO:1901343,4.81859826791183e-08,5.12183468705208,4.85507152145644,20,89,negative regulation of vasculature development,2.59528011973497e-07

GO:2000147,5.08220758225111e-08,2.48240809709504,23.4025357607282,52,429,positive regulation of cell motility,2.72134487398446e-07

GO:0030335,5.53970721475508e-08,2.49959888795142,22.8024707412224,51,418,positive regulation of cell migration,2.94917360970487e-07

GO:0030203,5.75849222110934e-08,4.5796241630067,5.83699609882965,22,107,glycosaminoglycan metabolic process,3.04802950324236e-07

GO:0001822,5.92052582029525e-08,2.904890625,15.2198309492848,39,279,kidney development,3.10185739128608e-07

GO:0001667,5.92754506912433e-08,2.60396791994014,20.2385565669701,47,371,ameboidal-type cell migration,3.10185739128608e-07

GO:0050654,6.34967963765293e-08,7.94291300097752,2.45481144343303,14,45,chondroitin sulfate proteoglycan metabolic process,3.303985845355e-07

GO:0048332,6.40987639171747e-08,6.55480516544689,3.21853055916775,16,59,mesoderm morphogenesis,3.31657087459089e-07

GO:0003156,6.70742372732715e-08,19.698343373494,0.927373211963589,9,17,regulation of animal organ formation,3.43196514048239e-07

GO:0060571,6.70742372732715e-08,19.698343373494,0.927373211963589,9,17,morphogenesis of an epithelial fold,3.43196514048239e-07

GO:0035113,6.95684802545775e-08,4.33199050896954,6.38250975292588,23,117,embryonic appendage morphogenesis,3.53992101184894e-07

GO:0030155,7.50181736969782e-08,2.1788820569111,34.0400520156047,67,624,regulation of cell adhesion,3.79624933928115e-07

GO:0060322,8.36191940947159e-08,2.14542335880243,35.5674902470741,69,652,head development,4.20837583394718e-07

GO:0045765,8.64896529111278e-08,3.12350643529487,12.437711313394,34,228,regulation of angiogenesis,4.32918317017112e-07

GO:0010646,1.00154926191114e-07,1.57365249263853,144.833875162549,203,2655,regulation of cell communication,4.98609119037923e-07

GO:0040013,1.03989393195579e-07,3.15986952693508,11.9467490247074,33,219,negative regulation of locomotion,5.14915221145852e-07

GO:0030166,1.06912273520452e-07,6.26261644390441,3.327633289987,16,61,proteoglycan biosynthetic process,5.26557240172921e-07

GO:0023051,1.12465425893649e-07,1.57056969967017,145.052080624187,203,2659,regulation of signaling,5.50699088101422e-07

GO:0048880,1.13009910587588e-07,2.56968097017124,20.0203511053316,46,367,sensory system development,5.50699088101422e-07

GO:1901342,1.1945808575339e-07,3.07529715925214,12.6013654096229,34,231,regulation of vasculature development,5.7905735252038e-07

GO:0022604,1.2167298177086e-07,2.95931730664386,13.8014954486346,36,253,regulation of cell morphogenesis,5.86705844036452e-07

GO:0040017,1.24689893219468e-07,2.40379169211774,24.0571521456437,52,441,positive regulation of locomotion,5.98121831537134e-07

GO:0045664,1.28432132692958e-07,3.52598578062282,9.21918075422627,28,169,regulation of neuron differentiation,6.12880799016655e-07

GO:0001654,1.29676926077387e-07,2.61835809356652,18.8202210663199,44,345,eye development,6.15631179985948e-07

GO:0060323,1.57456963046069e-07,7.24057040998217,2.6184655396619,14,48,head morphogenesis,7.43681348540665e-07

GO:0007369,1.66547554316195e-07,3.57500367620028,8.78276983094928,27,161,gastrulation,7.82603558802119e-07

GO:0030030,1.78441004162038e-07,1.73038901189218,79.699544863459,125,1461,cell projection organization,8.34234339255011e-07

GO:0048667,1.89632505877781e-07,2.17171391752577,32.0216514954486,63,587,cell morphogenesis involved in neuron differentiation,8.82078474310286e-07

GO:0001707,1.97644574942206e-07,6.4379587970637,3.05487646293888,15,56,mesoderm formation,9.14726902119455e-07

GO:0002063,2.09468611876816e-07,7.03321212121212,2.67301690507152,14,49,chondrocyte development,9.6460295769274e-07

GO:0048585,2.19680160543973e-07,1.78751848276368,67.0981794538362,109,1230,negative regulation of response to stimulus,1.00659416846268e-06

GO:0016049,2.44433397925355e-07,2.43899678751392,21.8750975292588,48,401,cell growth,1.11447108658045e-06

GO:0003197,2.50352445381898e-07,10.1539537248919,1.63654096228869,11,30,endocardial cushion development,1.13583547880162e-06

GO:0007417,2.58058236264454e-07,1.96294933137038,44.7866710013004,80,821,central nervous system development,1.16505703725276e-06

GO:0150063,2.93737586873044e-07,2.53238690367854,19.3657347204161,44,355,visual system development,1.31966984151255e-06

GO:0048518,3.20843126203958e-07,1.45318166868304,251.972756827048,318,4619,positive regulation of biological process,1.43444912249439e-06

GO:0006024,3.46924229450037e-07,5.25521833781534,4.03680104031209,17,74,glycosaminoglycan biosynthetic process,1.5435614266835e-06

GO:0098609,3.5510864510971e-07,1.97139885017899,42.8773732119636,77,786,cell-cell adhesion,1.57238010647136e-06

GO:0050650,3.69001876493025e-07,9.64559178743961,1.69109232769831,11,31,chondroitin sulfate proteoglycan biosynthetic process,1.62608003947405e-06

GO:0016331,3.83608099348629e-07,3.87521008403361,6.98257477243173,23,128,morphogenesis of embryonic epithelium,1.6744220829388e-06

GO:0030177,3.83608099348629e-07,3.87521008403361,6.98257477243173,23,128,positive regulation of Wnt signaling pathway,1.6744220829388e-06

GO:0120036,4.03151887788016e-07,1.71358291721757,77.6265929778934,121,1423,plasma membrane bounded cell projection organization,1.7514287200602e-06

GO:0032989,4.08248253487182e-07,1.97247143925886,42.2773081924577,76,775,cellular component morphogenesis,1.76524244817697e-06

GO:0048583,4.14075774852071e-07,1.51571231541846,162.672171651495,220,2982,regulation of response to stimulus,1.78207377868578e-06

GO:0060411,4.16067925283672e-07,5.9977658870256,3.21853055916775,15,59,cardiac septum morphogenesis,1.7823188799361e-06

GO:0060541,4.67599597293077e-07,3.464454494169,8.67366710013004,26,159,respiratory system development,1.99379272734687e-06

GO:0048645,4.94844107796584e-07,7.13593371670702,2.45481144343303,13,45,animal organ formation,2.10023697364357e-06

GO:0014033,5.08881532908822e-07,4.52987804878049,5.07327698309493,19,93,neural crest cell differentiation,2.14990776059186e-06

GO:0007160,5.52032933019717e-07,3.02693531715809,11.6194408322497,31,213,cell-matrix adhesion,2.32156315667196e-06

GO:0040008,5.7147173927214e-07,2.19915563809556,28.0394018205462,56,514,regulation of growth,2.39057541484535e-06

GO:0051128,5.73634274354857e-07,1.61924969718531,101.465539661899,149,1860,regulation of cellular component organization,2.39057541484535e-06

GO:0060993,5.78231437589275e-07,4.73634264729943,4.63686605981795,18,85,kidney morphogenesis,2.39887907216091e-06

GO:0031346,5.95709287559768e-07,2.55173648406731,17.8928478543563,41,328,positive regulation of cell projection organization,2.46030607104281e-06

GO:0003007,6.40784886416452e-07,2.78237717661692,14.1288036410923,35,259,heart morphogenesis,2.62958792888799e-06

GO:0006022,6.42407474484036e-07,3.88864137086903,6.65526657997399,22,122,aminoglycan metabolic process,2.62958792888799e-06

GO:0007267,6.48792682060286e-07,1.76901281711396,63.7705461638492,103,1169,cell-cell signaling,2.64397371759966e-06

GO:0003151,6.55616031477049e-07,6.9192163768435,2.50936280884265,13,46,outflow tract morphogenesis,2.66001041845974e-06

GO:0043010,6.67005021166369e-07,2.64722807428215,16.0381014304291,38,294,camera-type eye development,2.69434923023783e-06

GO:0007162,6.80312540198453e-07,2.93248508539966,12.3286085825748,32,226,negative regulation of cell adhesion,2.73610414638766e-06

GO:0001558,6.99782137490063e-07,2.53372184787228,18.0019505851756,41,330,regulation of cell growth,2.80217108099282e-06

GO:0032330,7.62719318008576e-07,8.76751207729469,1.80019505851756,11,33,regulation of chondrocyte differentiation,3.04097182634588e-06

GO:0010810,8.09369949156739e-07,3.10036442064554,10.6375162548765,29,195,regulation of cell-substrate adhesion,3.2130591516093e-06

GO:0045666,8.23673518909592e-07,5.21560685837721,3.8185955786736,16,70,positive regulation of neuron differentiation,3.24291522444496e-06

GO:0051130,8.23932858328035e-07,1.87582425175709,48.3870611183355,83,887,positive regulation of cellular component organization,3.24291522444496e-06

GO:0060324,9.9445446526592e-07,6.00147819660015,3.00032509752926,14,55,face development,3.89741515961665e-06

GO:0048704,1.01012650363824e-06,5.12042416878383,3.87314694408323,16,71,embryonic skeletal system morphogenesis,3.94206148241873e-06

GO:0072028,1.03799398572076e-06,5.49643507281553,3.43673602080624,15,63,nephron morphogenesis,4.03372346349714e-06

GO:0060325,1.16833912337177e-06,7.26114330984447,2.23660598179454,12,41,face morphogenesis,4.52117786817395e-06

GO:0060536,1.18110742567666e-06,9.73260956976277,1.52743823146944,10,28,cartilage morphogenesis,4.5324997460342e-06

GO:0003413,1.18110742567666e-06,9.73260956976277,1.52743823146944,10,28,chondrocyte differentiation involved in endochondral bone morphogenesis,4.5324997460342e-06

GO:1903510,1.35479843336295e-06,4.67816681873479,4.41866059817945,17,81,mucopolysaccharide metabolic process,5.17746621214637e-06

GO:0034330,1.49328779897434e-06,2.05262955577445,32.5126137841352,61,596,cell junction organization,5.68313249113789e-06

GO:0061383,1.55509009770608e-06,7.0186215235792,2.29115734720416,12,42,trabecula morphogenesis,5.89398345673787e-06

GO:0035265,1.63427876184722e-06,3.30205544757784,8.67366710013004,25,159,organ growth,6.16873253959546e-06

GO:0003279,1.6620528656433e-06,4.34525219828809,4.96417425227568,18,91,cardiac septum development,6.24796199696932e-06

GO:0061430,1.80167169330321e-06,20.3816105769231,0.709167750325097,7,13,bone trabecula morphogenesis,6.71797420863262e-06

GO:0070831,1.80167169330321e-06,20.3816105769231,0.709167750325097,7,13,basement membrane assembly,6.71797420863262e-06

GO:0010721,1.85823771914368e-06,3.02664581516345,10.4738621586476,28,192,negative regulation of cell development,6.90095540052957e-06

GO:0060513,1.88737157086884e-06,34.9051620648259,0.490962288686606,6,9,prostatic bud formation,6.95307686708079e-06

GO:0042481,1.88737157086884e-06,34.9051620648259,0.490962288686606,6,9,regulation of odontogenesis,6.95307686708079e-06

GO:0014032,1.939427614043e-06,4.53577748285777,4.5277633289987,17,83,neural crest cell development,7.08814616084763e-06

GO:0006023,1.939427614043e-06,4.53577748285777,4.5277633289987,17,83,aminoglycan biosynthetic process,7.08814616084763e-06

GO:0003205,1.95639773836025e-06,3.47486173956762,7.6371911573472,23,140,cardiac chamber development,7.09825913409255e-06

GO:0021915,1.95760892514605e-06,3.59859921120631,7.09167750325097,22,130,neural tube development,7.09825913409255e-06

GO:1905314,2.0345296202086e-06,13.9889290012034,0.981924577373212,8,18,semi-lunar valve development,7.32634776259714e-06

GO:0060412,2.03642239655252e-06,7.71381642512077,1.96384915474642,11,36,ventricular septum morphogenesis,7.32634776259714e-06

GO:0050770,2.17903323027812e-06,3.15312810662353,9.38283485045514,26,172,regulation of axonogenesis,7.80890896920681e-06

GO:1904888,2.47768284743726e-06,5.46650505050505,3.21853055916775,14,59,cranial skeletal system development,8.84475156003764e-06

GO:0048863,2.50562761823267e-06,3.04640582347589,10.0374512353706,27,184,stem cell differentiation,8.90997311348375e-06

GO:0001958,2.55023278333814e-06,10.5007228915663,1.30923276983095,9,24,endochondral ossification,8.99909729292883e-06

GO:0036075,2.55023278333814e-06,10.5007228915663,1.30923276983095,9,24,replacement ossification,8.99909729292883e-06

GO:0048519,2.59300063719928e-06,1.42599436893481,218.860078023407,277,4012,negative regulation of biological process,9.08743816992856e-06

GO:0090101,2.5950013449416e-06,3.98784843205575,5.61879063719116,19,103,negative regulation of transmembrane receptor protein serine/threonine kinase signaling pathway,9.08743816992856e-06

GO:0060284,2.61797139847761e-06,1.97711945996791,35.294733420026,64,647,regulation of cell development,9.13315021968895e-06

GO:0050896,2.83740838616126e-06,1.38468336892742,356.984135240572,421,6544,response to stimulus,9.86133254209255e-06

GO:0006935,2.86527777219155e-06,2.43852631578947,17.6746423927178,39,324,chemotaxis,9.88359860744727e-06

GO:0042330,2.86527777219155e-06,2.43852631578947,17.6746423927178,39,324,taxis,9.88359860744727e-06

GO:0030510,2.92024740501955e-06,4.97618611467302,3.70949284785436,15,68,regulation of BMP signaling pathway,1.00356263433694e-05

GO:0060675,3.01706085352099e-06,5.85230024213075,2.83667100130039,13,52,ureteric bud morphogenesis,1.02915298003438e-05

GO:0072171,3.01706085352099e-06,5.85230024213075,2.83667100130039,13,52,mesonephric tubule morphogenesis,1.02915298003438e-05

GO:0009952,3.2147264230205e-06,3.35905849943283,7.8553966189857,23,144,anterior/posterior pattern specification,1.09253248546195e-05

GO:0048864,3.24723852648923e-06,4.33767058076801,4.69141742522757,17,86,stem cell development,1.0995245157708e-05

GO:0030308,3.62766413439749e-06,3.33129419800707,7.90994798439532,23,145,negative regulation of cell growth,1.22383833984618e-05

GO:0072175,4.67260181399625e-06,3.80552937915743,5.83699609882965,19,107,epithelial tube formation,1.56489682570565e-05

GO:0048565,4.67260181399625e-06,3.80552937915743,5.83699609882965,19,107,digestive tract development,1.56489682570565e-05

GO:0061387,4.84337919158352e-06,3.63631786312199,6.38250975292588,20,117,regulation of extent of cell growth,1.61037986839293e-05

GO:0051961,4.84337919158352e-06,3.63631786312199,6.38250975292588,20,117,negative regulation of nervous system development,1.61037986839293e-05

GO:0048675,5.53352508862e-06,3.46313095537326,6.98257477243173,21,128,axon extension,1.8332289951867e-05

GO:0045785,5.6237212566334e-06,2.30538491338757,19.529388816645,41,358,positive regulation of cell adhesion,1.85643271589941e-05

GO:0051962,5.68339438936445e-06,2.6174658375162,13.5832899869961,32,249,positive regulation of nervous system development,1.86943079735881e-05

GO:0007420,5.83615364225762e-06,1.96672694733094,33.1672301690507,60,608,brain development,1.91284608701753e-05

GO:0048638,5.8693268027341e-06,2.52744613945269,14.892522756827,34,273,regulation of developmental growth,1.91689715791422e-05

GO:0003281,5.92226381522993e-06,5.43315461777931,3.00032509752926,13,55,ventricular septum development,1.92735158085752e-05

GO:0030323,6.09336408726769e-06,3.31971251922292,7.58263979193758,22,139,respiratory tube development,1.97605222689209e-05

GO:0050919,6.41080394779196e-06,6.648009328669,2.18205461638492,11,40,negative chemotaxis,2.06445819437636e-05

GO:0030204,6.41080394779196e-06,6.648009328669,2.18205461638492,11,40,chondroitin sulfate metabolic process,2.06445819437636e-05

GO:0048588,6.57276605647842e-06,2.74817455601769,11.7830949284785,29,216,developmental cell growth,2.10923956028454e-05

GO:0003231,6.78918412149604e-06,3.86592792846321,5.45513654096229,18,100,cardiac ventricle development,2.1636119639785e-05

GO:0003206,6.78918412149604e-06,3.86592792846321,5.45513654096229,18,100,cardiac chamber morphogenesis,2.1636119639785e-05

GO:0003203,7.88184995419305e-06,8.74879518072289,1.47288686605982,9,27,endocardial cushion morphogenesis,2.49456488240955e-05

GO:1901861,7.88184995419305e-06,8.74879518072289,1.47288686605982,9,27,regulation of muscle tissue development,2.49456488240955e-05

GO:0110110,8.17000202337516e-06,10.7584930112006,1.14557867360208,8,21,positive regulation of animal organ morphogenesis,2.57690817244127e-05

GO:1902284,8.75089080651804e-06,7.29644149577805,1.85474642392718,10,34,neuron projection extension involved in neuron projection guidance,2.741350487348e-05

GO:0048846,8.75089080651804e-06,7.29644149577805,1.85474642392718,10,34,axon extension involved in axon guidance,2.741350487348e-05

GO:0055123,9.32204445807738e-06,3.59968528717545,6.10975292587776,19,112,digestive system development,2.91037387996246e-05

GO:1990138,9.3801231824874e-06,3.02913234005259,8.94642392717815,24,164,neuron projection extension,2.91861265238881e-05

GO:0060346,9.44186832980628e-06,20.9402160864346,0.600065019505852,6,11,bone trabecula formation,2.92793290631366e-05

GO:0061333,1.01020084619926e-05,4.72834498834499,3.60039011703511,14,66,renal tubule morphogenesis,3.12213080318631e-05

GO:0035988,1.03924510580768e-05,13.5849358974359,0.872821846553966,7,16,chondrocyte proliferation,3.20115298477885e-05

GO:0050768,1.06437593285291e-05,3.56114426569798,6.16430429128739,19,113,negative regulation of neurogenesis,3.26449968856176e-05

GO:0050673,1.06689946390563e-05,2.378589582745,16.638166449935,36,305,epithelial cell proliferation,3.26449968856176e-05

GO:0072078,1.10629083319355e-05,5.06989507667474,3.16397919375813,13,58,nephron tubule morphogenesis,3.37382071977237e-05

GO:0030324,1.15289640943119e-05,3.27788476101867,7.30988296488947,21,134,lung development,3.50434849203342e-05

GO:1903053,1.16564354259851e-05,7.00410132689988,1.9092977893368,10,35,regulation of extracellular matrix organization,3.53143981162246e-05

GO:0090596,1.19491485837638e-05,2.47108709991519,14.7288686605982,33,270,sensory organ morphogenesis,3.60825109693327e-05

GO:0034329,1.31675457456622e-05,2.23911965428484,19.529388816645,40,358,cell junction assembly,3.95162848065256e-05

GO:0051716,1.31720949355085e-05,1.36024759318206,297.250390117035,355,5449,cellular response to stimulus,3.95162848065256e-05

GO:0043583,1.33881186867812e-05,2.63396011396011,12.2195058517555,29,224,ear development,4.0033952306901e-05

GO:0060395,1.34891677079893e-05,4.95933782503421,3.21853055916775,13,59,SMAD protein signal transduction,4.02055775374049e-05

GO:0034446,1.37211908422491e-05,3.64248813473896,5.7278933680104,18,105,substrate adhesion-dependent cell spreading,4.07652153732625e-05

GO:0061351,1.38032182010914e-05,3.48647103658537,6.27340702210663,19,115,neural precursor cell proliferation,4.08770545440681e-05

GO:0001655,1.41793876080799e-05,5.39559110780393,2.78211963589077,12,51,urogenital system development,4.18564614969282e-05

GO:0045446,1.45274403831969e-05,4.55259259259259,3.70949284785436,14,68,endothelial cell differentiation,4.27468772936881e-05

GO:0072080,1.46712162180325e-05,4.25119401816474,4.20045513654096,15,77,nephron tubule development,4.30324526649935e-05

GO:0048841,1.52592919451865e-05,7.8728313253012,1.58198959687906,9,29,regulation of axon extension involved in axon guidance,4.46152631159263e-05

GO:0072088,1.6370693342667e-05,4.85348513729329,3.27308192457737,13,60,nephron epithelium morphogenesis,4.77133182550517e-05

GO:0048771,1.72068310103081e-05,3.29419282690311,6.92802340702211,20,127,tissue remodeling,4.99920863107057e-05

GO:0008361,1.73521813339571e-05,2.98546532583622,8.67366710013004,23,159,regulation of cell size,5.02558459389134e-05

GO:0072009,1.75337511448457e-05,3.73839720194647,5.29148244473342,17,97,nephron epithelium development,5.06225228978147e-05

GO:0003433,1.78541093705616e-05,9.32274368231047,1.25468140442133,8,23,chondrocyte development involved in endochondral bone morphogenesis,5.12262764183404e-05

GO:0003418,1.78541093705616e-05,9.32274368231047,1.25468140442133,8,23,growth plate cartilage chondrocyte differentiation,5.12262764183404e-05

GO:0002719,1.80110218774474e-05,17.4489795918367,0.654616384915475,6,12,negative regulation of cytokine production involved in immune response,5.15159973575437e-05

GO:0048546,2.0023243768841e-05,6.48438547111647,2.01840052015605,10,37,digestive tract morphogenesis,5.70941409012463e-05

GO:0008360,2.04642261601341e-05,3.13790352658406,7.58263979193758,21,139,regulation of cell shape,5.81714576959366e-05

GO:0050789,2.05831213184538e-05,1.35420254217427,484.907087126138,542,8889,regulation of biological process,5.8329399182449e-05

GO:0002011,2.15460671447373e-05,5.13168372312502,2.89122236671001,12,53,morphogenesis of an epithelial sheet,6.08709442954081e-05

GO:0001658,2.19120820090589e-05,5.66840721795965,2.45481144343303,11,45,branching involved in ureteric bud morphogenesis,6.17156805209275e-05

GO:1903844,2.21616486955894e-05,3.85306013748564,4.85507152145644,16,89,regulation of cellular response to transforming growth factor beta stimulus,6.2228287953164e-05

GO:1904948,2.32876696027054e-05,29.052757793765,0.436410923276983,5,8,midbrain dopaminergic neuron differentiation,6.51913182495189e-05

GO:0003181,2.55060476641734e-05,8.73947051744886,1.30923276983095,8,24,atrioventricular valve morphogenesis,7.1185060299102e-05

GO:0035148,2.56766573124266e-05,3.31272639459068,6.54616384915475,19,120,tube formation,7.1444717174456e-05

GO:0030850,2.58568891527171e-05,6.25236946407031,2.07295188556567,10,38,prostate gland development,7.1729502739917e-05

GO:0022407,2.60584641663537e-05,2.21431407461984,18.7111183355007,38,343,regulation of cell-cell adhesion,7.2071608099735e-05

GO:0001935,2.64160411526432e-05,3.44333527511518,6.00065019505852,18,110,endothelial cell proliferation,7.28418380286958e-05

GO:0045926,2.74679223288441e-05,2.74314404749187,10.1465539661899,25,186,negative regulation of growth,7.55162879548222e-05

GO:0090497,2.85002823128979e-05,4.56133171912833,3.43673602080624,13,63,mesenchymal cell migration,7.7889495579166e-05

GO:0001755,2.85002823128979e-05,4.56133171912833,3.43673602080624,13,63,neural crest cell migration,7.7889495579166e-05

GO:0061053,2.87529257916668e-05,4.23745036572623,3.92769830949285,14,72,somite development,7.83474693909026e-05

GO:0065007,2.99868631544884e-05,1.34978833424469,501.599804941482,557,9195,biological regulation,8.14687344108667e-05

GO:0042063,3.04205936974723e-05,2.55326884208118,12.1104031209363,28,222,gliogenesis,8.24040199863881e-05

GO:0003176,3.1904750606714e-05,14.9552392385526,0.709167750325097,6,13,aortic valve development,8.56684411334798e-05

GO:0060601,3.1904750606714e-05,14.9552392385526,0.709167750325097,6,13,lateral sprouting from an epithelium,8.56684411334798e-05

GO:1901201,3.1904750606714e-05,14.9552392385526,0.709167750325097,6,13,regulation of extracellular matrix assembly,8.56684411334798e-05

GO:0003158,3.21302816110342e-05,3.93258223445877,4.47321196358908,15,82,endothelium development,8.60232248946585e-05

GO:0007498,3.39645624171774e-05,3.70019824774573,5.01872561768531,16,92,mesoderm development,9.06706144528128e-05

GO:0003171,3.57241869205402e-05,8.2248177249239,1.36378413524057,8,25,atrioventricular valve development,9.45459084879815e-05

GO:0048843,3.57241869205402e-05,8.2248177249239,1.36378413524057,8,25,negative regulation of axon extension involved in axon guidance,9.45459084879815e-05

GO:0048566,3.57241869205402e-05,8.2248177249239,1.36378413524057,8,25,embryonic digestive tract development,9.45459084879815e-05

GO:0043408,3.69255512217094e-05,1.99521103896104,25.4754876462939,47,467,regulation of MAPK cascade,9.74453658315025e-05

GO:0048523,3.73421932147526e-05,1.3712257582613,205.494993498049,255,3767,negative regulation of cellular process,9.82633141451061e-05

GO:0048333,3.96192156451171e-05,10.1865985576923,1.03647594278283,7,19,mesodermal cell differentiation,0.000103727130855732

GO:0030879,3.96438111413874e-05,4.09563636363636,4.03680104031209,14,74,mammary gland development,0.000103727130855732

GO:0009968,3.99215991139989e-05,1.65559303992515,55.2059817945384,85,1012,negative regulation of signal transduction,0.000104158053212445

GO:0060685,4.20689881833634e-05,69.6526946107784,0.272756827048114,4,5,regulation of prostatic bud formation,0.000108530919094896

GO:0060686,4.20689881833634e-05,69.6526946107784,0.272756827048114,4,5,negative regulation of prostatic bud formation,0.000108530919094896

GO:0061309,4.20689881833634e-05,69.6526946107784,0.272756827048114,4,5,cardiac neural crest cell development involved in outflow tract morphogenesis,0.000108530919094896

GO:1904953,4.20689881833634e-05,69.6526946107784,0.272756827048114,4,5,Wnt signaling pathway involved in midbrain dopaminergic neuron differentiation,0.000108530919094896

GO:0061326,4.31230800271859e-05,3.81806669480794,4.58231469440832,15,84,renal tubule development,0.000110939543868822

GO:0022408,4.80403525089383e-05,2.93704156479218,8.01905071521456,21,147,negative regulation of cell-cell adhesion,0.000123245584013182

GO:0030206,4.91479951769416e-05,7.76734857601284,1.4183355006502,8,26,chondroitin sulfate biosynthetic process,0.000125736954327676

GO:0030336,4.93401356661771e-05,2.69794849751866,9.87379713914174,24,181,negative regulation of cell migration,0.000125878850272989

GO:0060840,4.9766519880714e-05,3.76326282940361,4.63686605981795,15,85,artery development,0.000126615924889883

GO:0060710,5.00380421425012e-05,21.7880695443645,0.490962288686606,5,9,chorio-allantoic fusion,0.000126956024278908

GO:0010631,5.62110414430922e-05,2.37015495320501,13.8560468140442,30,254,epithelial cell migration,0.000142226288926066

GO:0048839,5.71372253505892e-05,2.55149651496515,11.2375812743823,26,206,inner ear development,0.000144173656295596

GO:0060603,5.80928795907555e-05,9.40236686390532,1.09102730819246,7,20,mammary gland duct morphogenesis,0.000145576758693489

GO:0040036,5.80928795907555e-05,9.40236686390532,1.09102730819246,7,20,regulation of fibroblast growth factor receptor signaling pathway,0.000145576758693489

GO:0001838,5.81674779578761e-05,3.5142162818955,5.2369310793238,16,96,embryonic epithelial tube formation,0.000145576758693489

GO:0010632,6.08987483578746e-05,2.59629281688105,10.6375162548765,25,195,regulation of epithelial cell migration,0.000151999315007053

GO:0050769,6.20412059944919e-05,2.53722316228687,11.2921326397919,26,207,positive regulation of neurogenesis,0.000154432299245749

GO:2000242,6.24719603611463e-05,6.29609638554217,1.85474642392718,9,34,negative regulation of reproductive process,0.000155085378686296

GO:0035567,6.27249488289673e-05,3.89979797979798,4.20045513654096,14,77,non-canonical Wnt signaling pathway,0.000155294832987846

GO:0090132,6.50282208605153e-05,2.3488519640767,13.9651495448635,30,256,epithelium migration,0.000160565660623417

GO:0017015,6.57953119637224e-05,3.65822208737864,4.74596879063719,15,87,regulation of transforming growth factor beta receptor signaling pathway,0.000162025353793017

GO:0071711,6.65232142180991e-05,7.35803407435556,1.47288686605982,8,27,basement membrane organization,0.000163381014119651

GO:0014706,6.76225535215647e-05,2.47196202047015,12.001300390117,27,220,striated muscle tissue development,0.000165639286684471

GO:0072006,7.1352668825116e-05,3.03978935698448,7.03712613784135,19,129,nephron development,0.000174312488031649

GO:0023057,7.32125763006381e-05,1.60514152514153,59.460988296489,89,1090,negative regulation of signaling,0.000177912355601287

GO:0010648,7.32125763006381e-05,1.60514152514153,59.460988296489,89,1090,negative regulation of cell communication,0.000177912355601287

GO:0065008,7.46058417474071e-05,1.44095742476975,116.248959687906,155,2131,regulation of biological quality,0.000180821000656216

GO:0060711,8.00619497617788e-05,6.05352177942539,1.9092977893368,9,35,labyrinthine layer development,0.000193028941703137

GO:0022617,8.00619497617788e-05,6.05352177942539,1.9092977893368,9,35,extracellular matrix disassembly,0.000193028941703137

GO:0090130,8.06042076308588e-05,2.31759517653473,14.1288036410923,30,259,tissue migration,0.000193828917044441

GO:0050922,8.18271635996351e-05,5.30321307160873,2.34570871261378,10,43,negative regulation of chemotaxis,0.000195747578377309

GO:0048701,8.18271635996351e-05,5.30321307160873,2.34570871261378,10,43,embryonic cranial skeleton morphogenesis,0.000195747578377309

GO:0060343,8.30543183744457e-05,8.73016826923077,1.14557867360208,7,21,trabecula formation,0.000198168464307939

GO:0010692,8.467401677211e-05,11.6302521008403,0.818270481144343,6,15,regulation of alkaline phosphatase activity,0.000200991673832766

GO:0010935,8.467401677211e-05,11.6302521008403,0.818270481144343,6,15,regulation of macrophage cytokine production,0.000200991673832766

GO:0030178,8.82034419628054e-05,2.98509364111498,7.1462288686606,19,131,negative regulation of Wnt signaling pathway,0.000208308451055785

GO:0010001,8.82087903493554e-05,2.65061514801999,9.60104031209363,23,176,glial cell differentiation,0.000208308451055785

GO:0006027,8.8710145549837e-05,6.98965102286402,1.52743823146944,8,28,glycosaminoglycan catabolic process,0.000208956634402557

GO:0043542,9.2334206937547e-05,2.71123246398644,9.00097529258778,22,165,endothelial cell migration,0.000216938277013982

GO:0009100,9.44681049192989e-05,2.06861821496136,19.8566970091027,38,364,glycoprotein metabolic process,0.000221387085574235

GO:0072498,9.55761527468742e-05,17.4292565947242,0.545513654096229,5,10,embryonic skeletal joint development,0.000223415321522516

GO:0090263,9.82845383194563e-05,3.51116504854369,4.90962288686606,15,90,positive regulation of canonical Wnt signaling pathway,0.000229164708334732

GO:2000146,9.88186638273388e-05,2.56571109871723,10.3102080624187,24,189,negative regulation of cell motility,0.000229828256022674

GO:0010594,0.000107217335232772,3.04349761079359,6.65526657997399,18,122,regulation of endothelial cell migration,0.000248733414985853

GO:0035904,0.00011033772779167,4.5861801242236,2.89122236671001,11,53,aorta development,0.000255329264563135

GO:0003272,0.000116107300319555,8.14759615384615,1.2001300390117,7,22,endocardial cushion formation,0.000264711331742724

GO:1903055,0.000116107300319555,8.14759615384615,1.2001300390117,7,22,positive regulation of extracellular matrix organization,0.000264711331742724

GO:0061036,0.000116107300319555,8.14759615384615,1.2001300390117,7,22,positive regulation of cartilage development,0.000264711331742724

GO:0009954,0.000116107300319555,8.14759615384615,1.2001300390117,7,22,proximal/distal pattern formation,0.000264711331742724

GO:0003422,0.000116107300319555,8.14759615384615,1.2001300390117,7,22,growth plate cartilage morphogenesis,0.000264711331742724

GO:0001569,0.000116691423113514,6.65635207151453,1.58198959687906,8,29,branching involved in blood vessel morphogenesis,0.000264711331742724

GO:0060037,0.000116691423113514,6.65635207151453,1.58198959687906,8,29,pharyngeal system development,0.000264711331742724

GO:0060317,0.000116691423113514,6.65635207151453,1.58198959687906,8,29,cardiac epithelial to mesenchymal transition,0.000264711331742724

GO:1903010,0.00012073406982967,34.8239520958084,0.327308192457737,4,6,regulation of bone development,0.00026711447496571

GO:0061307,0.00012073406982967,34.8239520958084,0.327308192457737,4,6,cardiac neural crest cell differentiation involved in heart development,0.00026711447496571

GO:0061308,0.00012073406982967,34.8239520958084,0.327308192457737,4,6,cardiac neural crest cell development involved in heart development,0.00026711447496571

GO:0071895,0.00012073406982967,34.8239520958084,0.327308192457737,4,6,odontoblast differentiation,0.00026711447496571

GO:0044851,0.00012073406982967,34.8239520958084,0.327308192457737,4,6,hair cycle phase,0.00026711447496571

GO:0044848,0.00012073406982967,34.8239520958084,0.327308192457737,4,6,biological phase,0.00026711447496571

GO:0048819,0.00012073406982967,34.8239520958084,0.327308192457737,4,6,regulation of hair follicle maturation,0.00026711447496571

GO:0010936,0.00012073406982967,34.8239520958084,0.327308192457737,4,6,negative regulation of macrophage cytokine production,0.00026711447496571

GO:1905331,0.00012073406982967,34.8239520958084,0.327308192457737,4,6,negative regulation of morphogenesis of an epithelium,0.00026711447496571

GO:0060272,0.00012073406982967,34.8239520958084,0.327308192457737,4,6,embryonic skeletal joint morphogenesis,0.00026711447496571

GO:0030900,0.000120941081499133,2.16406120760959,16.5290637191157,33,303,forebrain development,0.00026711447496571

GO:0010463,0.000123306895208252,4.99948302602102,2.45481144343303,10,45,mesenchymal cell proliferation,0.00027168815905933

GO:0014070,0.000126090981454238,1.77040364728122,34.4764629388817,57,632,response to organic cyclic compound,0.000277029365850953

GO:0050920,0.000126332609834311,2.71919495181936,8.56456436931079,21,157,regulation of chemotaxis,0.000277029365850953

GO:0010720,0.00012737304858106,2.05682502208429,19.4202860858257,37,356,positive regulation of cell development,0.000278647453071629

GO:0060563,0.00012924295620053,10.4665066026411,0.872821846553966,6,16,neuroepithelial cell differentiation,0.000281892425362477

GO:0010811,0.000129468508065502,3.11188412408759,6.16430429128739,17,113,positive regulation of cell-substrate adhesion,0.000281892425362477

GO:0009798,0.000130414503219396,4.12262607582332,3.43673602080624,12,63,axis specification,0.000283282446851565

GO:0030512,0.000131824802834675,4.47921581844737,2.94577373211964,11,54,negative regulation of transforming growth factor beta receptor signaling pathway,0.000285672102142908

GO:0007265,0.000134867793244117,1.95326059109094,23.1297789336801,42,424,Ras protein signal transduction,0.000291580369901014

GO:0048732,0.000137072727841748,2.14784794190629,16.638166449935,33,305,gland development,0.000295653354431498

GO:0007264,0.000138422872217186,1.82504892667115,29.948699609883,51,549,small GTPase mediated signal transduction,0.000297867909607544

GO:0030902,0.000147206092267685,2.95755119696745,6.81892067620286,18,125,hindbrain development,0.000316029862420835

GO:0034405,0.000151579160236096,6.35335302483317,1.63654096228869,8,30,response to fluid shear stress,0.00032466141064522

GO:0008652,0.000156770615586875,4.37711352657005,3.00032509752926,11,55,amino acid biosynthetic process,0.000334226242952574

GO:0001892,0.000156770615586875,4.37711352657005,3.00032509752926,11,55,embryonic placenta development,0.000334226242952574

GO:0007435,0.000159517895689289,5.42617366015787,2.07295188556567,9,38,salivary gland morphogenesis,0.000338516087395935

GO:0061180,0.000159517895689289,5.42617366015787,2.07295188556567,9,38,mammary gland epithelium development,0.000338516087395935

GO:0044087,0.000160205612155286,1.68836234215787,41.1317295188557,65,754,regulation of cellular component biogenesis,0.000339193951252916

GO:0046946,0.000161788313719201,Inf,0.163654096228869,3,3,hydroxylysine metabolic process,0.000340198714464348

GO:0046947,0.000161788313719201,Inf,0.163654096228869,3,3,hydroxylysine biosynthetic process,0.000340198714464348

GO:0042482,0.000161788313719201,Inf,0.163654096228869,3,3,positive regulation of odontogenesis,0.000340198714464348

GO:0010977,0.000163173482900284,2.9299634591961,6.87347204161248,18,126,negative regulation of neuron projection development,0.000342329789865972

GO:0051963,0.000166296481238782,3.50812121212121,4.58231469440832,14,84,regulation of synapse assembly,0.000348088770956633

GO:0001710,0.000167354723094431,14.523381294964,0.600065019505852,5,11,mesodermal cell fate commitment,0.000348718778212604

GO:0003344,0.000167354723094431,14.523381294964,0.600065019505852,5,11,pericardium morphogenesis,0.000348718778212604

GO:0001841,0.000189274220098119,3.45847204438754,4.63686605981795,14,85,neural tube formation,0.000393501807568466

GO:0061311,0.000190555707030983,9.51435119502346,0.927373211963589,6,17,cell surface receptor signaling pathway involved in heart development,0.000393501807568466

GO:0010934,0.000190555707030983,9.51435119502346,0.927373211963589,6,17,macrophage cytokine production,0.000393501807568466

GO:0060039,0.000190555707030983,9.51435119502346,0.927373211963589,6,17,pericardium development,0.000393501807568466

GO:0030516,0.00019460285142882,3.1215876873228,5.78244473342003,16,106,regulation of axon extension,0.000400960237507704

GO:0051129,0.000208701526330945,1.81304771178189,28.9122236671001,49,530,negative regulation of cellular component organization,0.00042904934319375

GO:0061037,0.000214099492881444,7.18806561085973,1.30923276983095,7,24,negative regulation of cartilage development,0.000439166220364833

GO:0014020,0.000215866472214641,3.61685691225643,4.14590377113134,13,76,primary neural tube formation,0.000441806713132632

GO:0050678,0.00021756938477606,2.20817901234568,14.2379063719116,29,261,regulation of epithelial cell proliferation,0.000444077268919602

GO:0048013,0.000217940201467601,4.60383467716335,2.6184655396619,10,48,ephrin receptor signaling pathway,0.000444077268919602

GO:0002040,0.000221573102972597,2.95680662956807,6.4370611183355,17,118,sprouting angiogenesis,0.000450483063659519

GO:0050794,0.000229359111436868,1.29087425543969,459.595253576073,509,8425,regulation of cellular process,0.000465285774522809

GO:0022612,0.000232811381437048,3.20987745678428,5.29148244473342,15,97,gland morphogenesis,0.000469188801539433

GO:0048640,0.000232811381437048,3.20987745678428,5.29148244473342,15,97,negative regulation of developmental growth,0.000469188801539433

GO:0001936,0.000232811381437048,3.20987745678428,5.29148244473342,15,97,regulation of endothelial cell proliferation,0.000469188801539433

GO:0050771,0.000239874197335898,3.82174343190063,3.65494148244473,12,67,negative regulation of axonogenesis,0.000482367108616511

GO:0060688,0.00024258234377848,5.07539836766421,2.18205461638492,9,40,regulation of morphogenesis of a branching structure,0.000486750193071853

GO:0090288,0.000246973344152357,3.56009760895884,4.20045513654096,13,77,negative regulation of cellular response to growth factor stimulus,0.000492809872632306

GO:0009953,0.000246973344152357,3.56009760895884,4.20045513654096,13,77,dorsal/ventral pattern formation,0.000492809872632306

GO:0032964,0.000247207558258551,5.82310469314079,1.74564369310793,8,32,collagen biosynthetic process,0.000492809872632306

GO:0007416,0.000248021534366818,2.64544144243392,8.3463589076723,20,153,synapse assembly,0.000493364650435939

GO:0061458,0.000251173723832693,2.34477102828167,11.6194408322497,25,213,reproductive system development,0.000498558188900668

GO:0001708,0.000256932532306427,4.09687532120465,3.16397919375813,11,58,cell fate specification,0.000508892176890794

GO:0007501,0.000269519242466023,23.214371257485,0.38185955786736,4,7,mesodermal cell fate specification,0.000530400047673519

GO:0051964,0.000269519242466023,23.214371257485,0.38185955786736,4,7,negative regulation of synapse assembly,0.000530400047673519

GO:1905809,0.000269519242466023,23.214371257485,0.38185955786736,4,7,negative regulation of synapse organization,0.000530400047673519

GO:0048641,0.000272705800234515,8.72088835534214,0.981924577373212,6,18,regulation of skeletal muscle tissue development,0.000530868312324966

GO:0002701,0.000272705800234515,8.72088835534214,0.981924577373212,6,18,negative regulation of production of molecular mediator of immune response,0.000530868312324966

GO:0070723,0.000272705800234515,8.72088835534214,0.981924577373212,6,18,response to cholesterol,0.000530868312324966

GO:0003180,0.000274028795392276,12.4477560808496,0.654616384915475,5,12,aortic valve morphogenesis,0.000530868312324966

GO:0003177,0.000274028795392276,12.4477560808496,0.654616384915475,5,12,pulmonary valve development,0.000530868312324966

GO:0001759,0.000274028795392276,12.4477560808496,0.654616384915475,5,12,organ induction,0.000530868312324966

GO:0060438,0.000274028795392276,12.4477560808496,0.654616384915475,5,12,trachea development,0.000530868312324966

GO:0000165,0.000274368422005086,1.75703377223333,31.5852405721717,52,579,MAPK cascade,0.000530868312324966

GO:0048592,0.000277477975578462,2.54882387657027,9.0555266579974,21,166,eye morphogenesis,0.000535759361651495

GO:0009948,0.000283467667421795,6.78826121794872,1.36378413524057,7,25,anterior/posterior axis specification,0.000545039084959234

GO:0060443,0.000283467667421795,6.78826121794872,1.36378413524057,7,25,mammary gland morphogenesis,0.000545039084959234

GO:0030517,0.000310072194254951,4.37303980699638,2.72756827048114,10,50,negative regulation of axon extension,0.000594554637260085

GO:0060425,0.000310873440183289,5.58979542719615,1.80019505851756,8,33,lung morphogenesis,0.000594554637260085

GO:0097435,0.000311156715699632,1.69311659272702,36.4948634590377,58,669,supramolecular fiber organization,0.000594554637260085

GO:0050772,0.000349790449206909,3.22982456140351,4.90962288686606,14,90,positive regulation of axonogenesis,0.000666199007794536

GO:0048844,0.000350098067071179,3.92911367445529,3.27308192457737,11,60,artery morphogenesis,0.000666199007794536

GO:0007431,0.000358665269399726,4.7671412924425,2.29115734720416,9,42,salivary gland development,0.00067991294020179

GO:0007266,0.000358781421213974,2.28323000343726,11.8921976592978,25,218,Rho protein signal transduction,0.00067991294020179

GO:0060795,0.00038028422389116,8.04949672176563,1.03647594278283,6,19,cell fate commitment involved in formation of primary germ layer,0.000713323360903786

GO:0060713,0.00038028422389116,8.04949672176563,1.03647594278283,6,19,labyrinthine layer morphogenesis,0.000713323360903786

GO:0010092,0.00038028422389116,8.04949672176563,1.03647594278283,6,19,specification of animal organ identity,0.000713323360903786

GO:0035115,0.00038028422389116,8.04949672176563,1.03647594278283,6,19,embryonic forelimb morphogenesis,0.000713323360903786

GO:0036314,0.00038028422389116,8.04949672176563,1.03647594278283,6,19,response to sterol,0.000713323360903786

GO:0071526,0.000387304664743949,5.37443302786263,1.85474642392718,8,34,semaphorin-plexin signaling pathway,0.00072062140652359

GO:1903670,0.000387304664743949,5.37443302786263,1.85474642392718,8,34,regulation of sprouting angiogenesis,0.00072062140652359

GO:0030574,0.000387304664743949,5.37443302786263,1.85474642392718,8,34,collagen catabolic process,0.00072062140652359

GO:0006026,0.000387304664743949,5.37443302786263,1.85474642392718,8,34,aminoglycan catabolic process,0.00072062140652359

GO:0001756,0.000406412616058347,3.85026570048309,3.327633289987,11,61,somitogenesis,0.000754649232640601

GO:0035282,0.000412669475008208,3.34975430850306,4.41866059817945,13,81,segmentation,0.000764725526121851

GO:0060537,0.000420951583997356,1.95964402670193,19.1475292587776,35,351,muscle tissue development,0.000778506845103544

GO:0090090,0.00045327714300747,2.98978100176523,5.61879063719116,15,103,negative regulation of canonical Wnt signaling pathway,0.000836609716853467

GO:1902742,0.000475941431449082,6.10859375,1.47288686605982,7,27,apoptotic process involved in development,0.000876684116729208

GO:0031345,0.00048137516703684,2.56856472795497,8.12815344603381,19,149,negative regulation of cell projection organization,0.00088492321125934

GO:0045216,0.0004877676032844,2.49395994076845,8.78276983094928,20,161,cell-cell junction organization,0.000894888371762814

GO:0001890,0.000493403427318062,3.10652857690832,5.07327698309493,14,93,placenta development,0.000903428541868658

GO:0003306,0.000515750815329341,17.4095808383234,0.436410923276983,4,8,Wnt signaling pathway involved in heart development,0.000936108424573762

GO:0021536,0.000517104260741717,4.49411359724613,2.40026007802341,9,44,diencephalon development,0.000936108424573762

GO:0090171,0.000518366228591334,7.47401817870005,1.09102730819246,6,20,chondrocyte morphogenesis,0.000936108424573762

GO:0048730,0.000518366228591334,7.47401817870005,1.09102730819246,6,20,epidermis morphogenesis,0.000936108424573762

GO:0032331,0.000518366228591334,7.47401817870005,1.09102730819246,6,20,negative regulation of chondrocyte differentiation,0.000936108424573762

GO:0003429,0.000518366228591334,7.47401817870005,1.09102730819246,6,20,growth plate cartilage chondrocyte morphogenesis,0.000936108424573762

GO:0003414,0.000518366228591334,7.47401817870005,1.09102730819246,6,20,chondrocyte morphogenesis involved in endochondral bone morphogenesis,0.000936108424573762

GO:0050773,0.000558486504065527,2.92293689320388,5.7278933680104,15,105,regulation of dendrite development,0.00100658722161321

GO:0048608,0.000563126673495948,2.24821824827046,11.5648894668401,24,212,reproductive structure development,0.00101099350153951

GO:0021537,0.000563126673495948,2.24821824827046,11.5648894668401,24,212,telencephalon development,0.00101099350153951

GO:0000003,0.000566497561897075,1.54070001026408,53.0239271781534,77,972,reproduction,0.00101506664301013

GO:0016043,0.00057010978040186,1.27403064595635,280.830429128739,325,5148,cellular component organization,0.00101955554902935

GO:0042060,0.000578112502467049,1.96360628618693,18.0019505851756,33,330,wound healing,0.00103186359451968

GO:0009880,0.000593006378516466,3.97439412216252,2.94577373211964,10,54,embryonic pattern specification,0.00105436076180244

GO:0002065,0.000593006378516466,3.97439412216252,2.94577373211964,10,54,columnar/cuboidal epithelial cell differentiation,0.00105436076180244

GO:0048663,0.000615553602125084,4.36897590361446,2.45481144343303,9,45,neuron fate commitment,0.00107064604965825

GO:0010812,0.000615553602125084,4.36897590361446,2.45481144343303,9,45,negative regulation of cell-substrate adhesion,0.00107064604965825

GO:0042640,0.000620765462016836,52.177033492823,0.218205461638492,3,4,anagen,0.00107064604965825

GO:0003253,0.000620765462016836,52.177033492823,0.218205461638492,3,4,cardiac neural crest cell migration involved in outflow tract morphogenesis,0.00107064604965825

GO:0043378,0.000620765462016836,52.177033492823,0.218205461638492,3,4,positive regulation of CD8-positive, alpha-beta T cell differentiation,0.00107064604965825

GO:0051884,0.000620765462016836,52.177033492823,0.218205461638492,3,4,regulation of timing of anagen,0.00107064604965825

GO:0044340,0.000620765462016836,52.177033492823,0.218205461638492,3,4,canonical Wnt signaling pathway involved in regulation of cell proliferation,0.00107064604965825

GO:0048341,0.000620765462016836,52.177033492823,0.218205461638492,3,4,paraxial mesoderm formation,0.00107064604965825

GO:0048818,0.000620765462016836,52.177033492823,0.218205461638492,3,4,positive regulation of hair follicle maturation,0.00107064604965825

GO:1905069,0.000620765462016836,52.177033492823,0.218205461638492,3,4,allantois development,0.00107064604965825

GO:0045113,0.000620765462016836,52.177033492823,0.218205461638492,3,4,regulation of integrin biosynthetic process,0.00107064604965825

GO:2001187,0.000620765462016836,52.177033492823,0.218205461638492,3,4,positive regulation of CD8-positive, alpha-beta T cell activation,0.00107064604965825

GO:0021873,0.000620765462016836,52.177033492823,0.218205461638492,3,4,forebrain neuroblast division,0.00107064604965825

GO:0021502,0.000620765462016836,52.177033492823,0.218205461638492,3,4,neural fold elevation formation,0.00107064604965825

GO:0060439,0.000620765462016836,52.177033492823,0.218205461638492,3,4,trachea morphogenesis,0.00107064604965825

GO:0003419,0.000620765462016836,52.177033492823,0.218205461638492,3,4,growth plate cartilage chondrocyte proliferation,0.00107064604965825

GO:2000178,0.000632077497441715,9.68025579536371,0.76371911573472,5,14,negative regulation of neural precursor cell proliferation,0.00108608838646235

GO:0034616,0.000632077497441715,9.68025579536371,0.76371911573472,5,14,response to laminar fluid shear stress,0.00108608838646235

GO:0009611,0.000658645950415367,1.83655252659574,22.6388166449935,39,415,response to wounding,0.00112963299875708

GO:0051493,0.000684076516010218,1.7893120837757,24.9845253576073,42,458,regulation of cytoskeleton organization,0.00117106779041898

GO:0060740,0.000692490992678089,6.97527010804322,1.14557867360208,6,21,prostate gland epithelium morphogenesis,0.00117889871396769

GO:0060512,0.000692490992678089,6.97527010804322,1.14557867360208,6,21,prostate gland morphogenesis,0.00117889871396769

GO:0071542,0.000692490992678089,6.97527010804322,1.14557867360208,6,21,dopaminergic neuron differentiation,0.00117889871396769

GO:0030901,0.000712345710678477,4.81746130544836,2.01840052015605,8,37,midbrain development,0.00120999609716299

GO:0048534,0.000713385321128669,3.56407675791734,3.54583875162549,11,65,hematopoietic or lymphoid organ development,0.00120999609716299

GO:1905330,0.000728808647295481,4.25060240963855,2.50936280884265,9,46,regulation of morphogenesis of an epithelium,0.00123388375764548

GO:0015012,0.000760395332450974,5.55250218531468,1.58198959687906,7,29,heparan sulfate proteoglycan biosynthetic process,0.00128499835080247

GO:0009725,0.000766275424462084,1.67131805032501,33.0035760728218,52,605,response to hormone,0.00129256349071352

GO:0042698,0.000858485996661815,4.13845909955612,2.56391417425228,9,47,ovulation cycle,0.00144282044329477

GO:0035272,0.000858485996661815,4.13845909955612,2.56391417425228,9,47,exocrine system development,0.00144282044329477

GO:0016202,0.000888321033878811,13.9267065868263,0.490962288686606,4,9,regulation of striated muscle tissue development,0.00147946414503144

GO:0010172,0.000888321033878811,13.9267065868263,0.490962288686606,4,9,embryonic body morphogenesis,0.00147946414503144

GO:0010694,0.000888321033878811,13.9267065868263,0.490962288686606,4,9,positive regulation of alkaline phosphatase activity,0.00147946414503144

GO:1901862,0.000888321033878811,13.9267065868263,0.490962288686606,4,9,negative regulation of muscle tissue development,0.00147946414503144

GO:0060033,0.000888321033878811,13.9267065868263,0.490962288686606,4,9,anatomical structure regression,0.00147946414503144

GO:0050808,0.0008983044600247,1.86886564158062,19.965799739922,35,366,synapse organization,0.00148641691297486

GO:0007223,0.000905779237776513,8.71163069544365,0.818270481144343,5,15,Wnt signaling pathway, calcium modulating pathway,0.00148641691297486

GO:0055057,0.000905779237776513,8.71163069544365,0.818270481144343,5,15,neuroblast division,0.00148641691297486

GO:0033688,0.000905779237776513,8.71163069544365,0.818270481144343,5,15,regulation of osteoblast proliferation,0.00148641691297486

GO:0044089,0.000907766215484417,1.83298057377564,21.4932379713914,37,394,positive regulation of cellular component biogenesis,0.00148641691297486

GO:0003401,0.000908634877312536,6.53886554621849,1.2001300390117,6,22,axis elongation,0.00148641691297486

GO:0060669,0.000908634877312536,6.53886554621849,1.2001300390117,6,22,embryonic placenta morphogenesis,0.00148641691297486

GO:0030857,0.000908634877312536,6.53886554621849,1.2001300390117,6,22,negative regulation of epithelial cell differentiation,0.00148641691297486

GO:0031128,0.000908634877312536,6.53886554621849,1.2001300390117,6,22,developmental induction,0.00148641691297486

GO:0021983,0.000908634877312536,6.53886554621849,1.2001300390117,6,22,pituitary gland development,0.00148641691297486

GO:0048738,0.000940804060032701,2.19929534313725,11.2921326397919,23,207,cardiac muscle tissue development,0.00153631301292574

GO:0061082,0.000945687210951157,5.31072324414716,1.63654096228869,7,30,myeloid leukocyte cytokine production,0.00153883024962194

GO:0090102,0.000945687210951157,5.31072324414716,1.63654096228869,7,30,cochlea development,0.00153883024962194

GO:0033002,0.000991236049850998,2.54962879780398,7.30988296488947,17,134,muscle cell proliferation,0.00161010300160982

GO:0007405,0.00100630425709125,4.03206672845227,2.6184655396619,9,48,neuroblast proliferation,0.00163170109292437

GO:0022414,0.00101488106362952,1.51295456962878,52.4238621586476,75,961,reproductive process,0.00164271609772019

GO:0035987,0.00103068764226564,4.50603625635651,2.12750325097529,8,39,endodermal cell differentiation,0.00166537424302922

GO:0045185,0.001033346080025,2.8522903453136,5.45513654096229,14,100,maintenance of protein location,0.00166674560368306

GO:0010634,0.00105389754501464,2.62254573534255,6.70981794538362,16,123,positive regulation of epithelial cell migration,0.00169692244573162

GO:0045639,0.00106194687916276,3.64218938480096,3.16397919375813,10,58,positive regulation of myeloid cell differentiation,0.00170689891048674

GO:0042221,0.0011231011489018,1.32166370983639,140.578868660598,174,2577,response to chemical,0.00180204905599052

GO:0048545,0.00114395370373213,2.16356034392367,11.4557867360208,23,210,response to steroid hormone,0.00183231541067356

GO:0001953,0.00117317904411941,6.15380269754961,1.25468140442133,6,23,negative regulation of cell-matrix adhesion,0.00187081483072963

GO:0031290,0.00117317904411941,6.15380269754961,1.25468140442133,6,23,retinal ganglion cell axon guidance,0.00187081483072963

GO:1901607,0.00117408357455128,3.93099397590361,2.67301690507152,9,49,alpha-amino acid biosynthetic process,0.00187081483072963

GO:0030514,0.00122771396688743,4.36492178098676,2.18205461638492,8,40,negative regulation of BMP signaling pathway,0.0019528921649453

GO:0033687,0.00125875027485651,7.9191192500545,0.872821846553966,5,16,osteoblast proliferation,0.00199880862610835

GO:0010976,0.00127058836668117,2.48537104622871,7.47353706111834,17,137,positive regulation of neuron projection development,0.0020141340545841

GO:0032956,0.00128011176012552,1.95123456790123,15.8744473342003,29,291,regulation of actin cytoskeleton organization,0.00202574386782749

GO:2000241,0.00132742783176032,2.65556045895852,6.21885565669701,15,114,regulation of reproductive process,0.00209701720935035

GO:0007422,0.00136374460925047,3.83485160152806,2.72756827048114,9,50,peripheral nervous system development,0.00214702356430715

GO:0042246,0.00136374460925047,3.83485160152806,2.72756827048114,9,50,tissue regeneration,0.00214702356430715

GO:0021700,0.00138580028262863,2.09036020166434,12.3286085825748,24,226,developmental maturation,0.00217802399368766

GO:0090100,0.00139079636680993,3.49601930036188,3.27308192457737,10,60,positive regulation of transmembrane receptor protein serine/threonine kinase signaling pathway,0.00218215239153653

GO:0060831,0.0014168176213001,11.6047904191617,0.545513654096229,4,10,smoothened signaling pathway involved in dorsal/ventral neural tube patterning,0.00221167632070745

GO:0048069,0.0014168176213001,11.6047904191617,0.545513654096229,4,10,eye pigmentation,0.00221167632070745

GO:0048820,0.0014168176213001,11.6047904191617,0.545513654096229,4,10,hair follicle maturation,0.00221167632070745

GO:0010562,0.00144075939894239,1.64290876025939,31.530689206762,49,578,positive regulation of phosphorus metabolic process,0.00224145170004382

GO:0045937,0.00144075939894239,1.64290876025939,31.530689206762,49,578,positive regulation of phosphate metabolic process,0.00224145170004382

GO:0048048,0.00145359638511963,4.23235969806367,2.23660598179454,8,41,embryonic eye morphogenesis,0.00225399233092465

GO:0008544,0.00147192556235238,2.07991779745535,12.3831599479844,24,227,epidermis development,0.00225399233092465

GO:0048593,0.00148533911454853,2.5273390036452,6.92802340702211,16,127,camera-type eye morphogenesis,0.00225399233092465

GO:0060768,0.00148881017276705,26.0867224880383,0.272756827048114,3,5,regulation of epithelial cell proliferation involved in prostate gland development,0.00225399233092465

GO:0043569,0.00148881017276705,26.0867224880383,0.272756827048114,3,5,negative regulation of insulin-like growth factor receptor signaling pathway,0.00225399233092465

GO:0043932,0.00148881017276705,26.0867224880383,0.272756827048114,3,5,ossification involved in bone remodeling,0.00225399233092465

GO:0018057,0.00148881017276705,26.0867224880383,0.272756827048114,3,5,peptidyl-lysine oxidation,0.00225399233092465

GO:0030854,0.00148881017276705,26.0867224880383,0.272756827048114,3,5,positive regulation of granulocyte differentiation,0.00225399233092465

GO:0002051,0.00148881017276705,26.0867224880383,0.272756827048114,3,5,osteoblast fate commitment,0.00225399233092465

GO:0045112,0.00148881017276705,26.0867224880383,0.272756827048114,3,5,integrin biosynthetic process,0.00225399233092465

GO:0035905,0.00148881017276705,26.0867224880383,0.272756827048114,3,5,ascending aorta development,0.00225399233092465

GO:0070487,0.00148881017276705,26.0867224880383,0.272756827048114,3,5,monocyte aggregation,0.00225399233092465

GO:0042487,0.00148881017276705,26.0867224880383,0.272756827048114,3,5,regulation of odontogenesis of dentin-containing tooth,0.00225399233092465

GO:1902455,0.00148881017276705,26.0867224880383,0.272756827048114,3,5,negative regulation of stem cell population maintenance,0.00225399233092465

GO:0046884,0.00148881017276705,26.0867224880383,0.272756827048114,3,5,follicle-stimulating hormone secretion,0.00225399233092465

GO:0021953,0.00149169720638629,2.44428822145108,7.58263979193758,17,139,central nervous system neuron differentiation,0.00225399233092465

GO:0030513,0.0014928722278654,5.81152460984394,1.30923276983095,6,24,positive regulation of BMP signaling pathway,0.00225399233092465

GO:2000677,0.0014928722278654,5.81152460984394,1.30923276983095,6,24,regulation of transcription regulatory region DNA binding,0.00225399233092465

GO:0071840,0.00150822637451225,1.24599163237982,290.431469440832,331,5324,cellular component organization or biogenesis,0.00227344761198982

GO:0001656,0.00151629631386397,3.20634057971014,3.87314694408323,11,71,metanephros development,0.00228187729586392

GO:0009101,0.00152711673186173,1.90256200478431,16.8018205461638,30,308,glycoprotein biosynthetic process,0.00229441192503206

GO:0050803,0.00155882696817883,2.1483884128927,11.0193758127438,22,202,regulation of synapse structure or activity,0.00233824045226824

GO:0032355,0.00156408986400147,2.8449303874092,5.07327698309493,13,93,response to estradiol,0.00234231994267537

GO:0003006,0.00157221637381851,1.61067610449916,34.0946033810143,52,625,developmental process involved in reproduction,0.00235066766280333

GO:0022409,0.00166558124485427,2.0954231433506,11.7830949284785,23,216,positive regulation of cell-cell adhesion,0.00248622419207582

GO:1901135,0.00169446460651483,1.5292383632821,44.1320546163849,64,809,carbohydrate derivative metabolic process,0.00251882611323577

GO:0045665,0.00170342378156858,3.1535598321058,3.92769830949285,11,72,negative regulation of neuron differentiation,0.00251882611323577

GO:0042693,0.00170383134478381,7.25869304556355,0.927373211963589,5,17,muscle cell fate commitment,0.00251882611323577

GO:0051220,0.00170383134478381,7.25869304556355,0.927373211963589,5,17,cytoplasmic sequestering of protein,0.00251882611323577

GO:0048670,0.00170383134478381,7.25869304556355,0.927373211963589,5,17,regulation of collateral sprouting,0.00251882611323577

GO:1901889,0.00170383134478381,7.25869304556355,0.927373211963589,5,17,negative regulation of cell junction assembly,0.00251882611323577

GO:0021955,0.00171122723587725,4.10759538472429,2.29115734720416,8,42,central nervous system neuron axonogenesis,0.00252210096453005

GO:0002064,0.00171426189337222,2.33958587088916,8.3463589076723,18,153,epithelial cell development,0.00252210096453005

GO:1901888,0.00171426189337222,2.33958587088916,8.3463589076723,18,153,regulation of cell junction assembly,0.00252210096453005

GO:0021952,0.00172071719418862,4.69697670118343,1.80019505851756,7,33,central nervous system projection neuron axonogenesis,0.00252756066323401

GO:0045598,0.00172664350256584,2.80961348757959,5.12782834850455,13,94,regulation of fat cell differentiation,0.0025322271749413

GO:0022038,0.00187478945747917,5.50527579452834,1.36378413524057,6,25,corpus callosum development,0.00274076363545764

GO:0060445,0.00187478945747917,5.50527579452834,1.36378413524057,6,25,branching involved in salivary gland morphogenesis,0.00274076363545764

GO:0060021,0.00190897897591154,3.1024816892629,3.98224967490247,11,73,roof of mouth development,0.00278632272078372

GO:0045444,0.00191795362546296,2.25335365853659,9.11007802340702,19,167,fat cell differentiation,0.00279499254596738

GO:0016358,0.00195805418159782,1.940243289434,14.8379713914174,27,272,dendrite development,0.00284892243483664

GO:0001952,0.00200128540511981,2.63632453567937,5.83699609882965,14,107,regulation of cell-matrix adhesion,0.00290723006011884

GO:0046578,0.00201013306727023,1.96383869418093,14.1288036410923,26,259,regulation of Ras protein signal transduction,0.0029154843385132

GO:0050679,0.00203223650807093,2.36603522187464,7.80084525357607,17,143,positive regulation of epithelial cell proliferation,0.00294222976567198

GO:0008589,0.00203496238950386,3.29744861961445,3.43673602080624,10,63,regulation of smoothened signaling pathway,0.00294222976567198

GO:0010596,0.00206641673885947,4.52270299145299,1.85474642392718,7,34,negative regulation of endothelial cell migration,0.00298177452146027

GO:0032970,0.00206878818589914,1.84003400340034,17.8928478543563,31,328,regulation of actin filament-based process,0.00298177452146027

GO:0003184,0.00213079642027477,9.94627887082977,0.600065019505852,4,11,pulmonary valve morphogenesis,0.00304730357620041

GO:0051280,0.00213079642027477,9.94627887082977,0.600065019505852,4,11,negative regulation of release of sequestered calcium ion into cytosol,0.00304730357620041

GO:0072074,0.00213079642027477,9.94627887082977,0.600065019505852,4,11,kidney mesenchyme development,0.00304730357620041

GO:0035116,0.00213079642027477,9.94627887082977,0.600065019505852,4,11,embryonic hindlimb morphogenesis,0.00304730357620041

GO:0097094,0.00213079642027477,9.94627887082977,0.600065019505852,4,11,craniofacial suture morphogenesis,0.00304730357620041

GO:0060606,0.00213427796983495,3.05302507476421,4.03680104031209,11,74,tube closure,0.00304755040343874

GO:0032535,0.0021569694262784,1.85492160562097,17.1836801040312,30,315,regulation of cellular component size,0.0030751839653288

GO:0007224,0.00218690069412571,2.60809800128949,5.89154746423927,14,108,smoothened signaling pathway,0.00311303792780491

GO:0018126,0.00225426424653354,6.69987087253274,0.981924577373212,5,18,protein hydroxylation,0.00320397742447128

GO:0032504,0.00227044927430663,1.61050453970808,31.4215864759428,48,576,multicellular organism reproduction,0.00322200890853067

GO:0046639,0.00232628750440616,5.2296518607443,1.4183355006502,6,26,negative regulation of alpha-beta T cell differentiation,0.0032810272458776

GO:0048668,0.00232628750440616,5.2296518607443,1.4183355006502,6,26,collateral sprouting,0.0032810272458776

GO:0035136,0.00232628750440616,5.2296518607443,1.4183355006502,6,26,forelimb morphogenesis,0.0032810272458776

GO:0021532,0.00232628750440616,5.2296518607443,1.4183355006502,6,26,neural tube patterning,0.0032810272458776

GO:0001706,0.00233397441652427,3.87886081026875,2.40026007802341,8,44,endoderm formation,0.0032868355315273

GO:0007169,0.00242461364436321,1.61296834625323,30.7124187256177,47,563,transmembrane receptor protein tyrosine kinase signaling pathway,0.0034092659029901

GO:0051056,0.00249375827544771,1.81507187263577,18.1110533159948,31,332,regulation of small GTPase mediated signal transduction,0.00350114538366973

GO:0048814,0.00258430932089315,3.17710275249479,3.54583875162549,10,65,regulation of dendrite morphogenesis,0.00362275324892328

GO:0051965,0.00271428878639236,3.4168412781561,3.00032509752926,9,55,positive regulation of synapse assembly,0.00379917928916013

GO:1901654,0.00272810054034085,2.2925977915029,8.01905071521456,17,147,response to ketone,0.00381271714363266

GO:0060767,0.00285688627570442,17.38995215311,0.327308192457737,3,6,epithelial cell proliferation involved in prostate gland development,0.00387223551140303

GO:0016198,0.00285688627570442,17.38995215311,0.327308192457737,3,6,axon choice point recognition,0.00387223551140303

GO:0016199,0.00285688627570442,17.38995215311,0.327308192457737,3,6,axon midline choice point recognition,0.00387223551140303

GO:0060923,0.00285688627570442,17.38995215311,0.327308192457737,3,6,cardiac muscle cell fate commitment,0.00387223551140303

GO:0072038,0.00285688627570442,17.38995215311,0.327308192457737,3,6,mesenchymal stem cell maintenance involved in nephron morphogenesis,0.00387223551140303

GO:0001842,0.00285688627570442,17.38995215311,0.327308192457737,3,6,neural fold formation,0.00387223551140303

GO:2000643,0.00285688627570442,17.38995215311,0.327308192457737,3,6,positive regulation of early endosome to late endosome transport,0.00387223551140303

GO:1901621,0.00285688627570442,17.38995215311,0.327308192457737,3,6,negative regulation of smoothened signaling pathway involved in dorsal/ventral neural tube patterning,0.00387223551140303

GO:0021957,0.00285688627570442,17.38995215311,0.327308192457737,3,6,corticospinal tract morphogenesis,0.00387223551140303

GO:0060437,0.00285688627570442,17.38995215311,0.327308192457737,3,6,lung growth,0.00387223551140303

GO:0060174,0.00285688627570442,17.38995215311,0.327308192457737,3,6,limb bud formation,0.00387223551140303

GO:0044344,0.00288527739626071,2.76172595939668,4.80052015604681,12,88,cellular response to fibroblast growth factor stimulus,0.00387223551140303

GO:0021675,0.00290031842798457,3.1201533689471,3.60039011703511,10,66,nerve development,0.00387223551140303

GO:0051098,0.0029047326638253,1.90562730627306,14.5106631989597,26,266,regulation of binding,0.00387223551140303

GO:0034111,0.00292354282604867,6.22088043850634,1.03647594278283,5,19,negative regulation of homotypic cell-cell adhesion,0.00387223551140303

GO:0060751,0.00297249783557214,Inf,0.109102730819246,2,2,branch elongation involved in mammary gland duct branching,0.00387223551140303

GO:0042695,0.00297249783557214,Inf,0.109102730819246,2,2,thelarche,0.00387223551140303

GO:0060744,0.00297249783557214,Inf,0.109102730819246,2,2,mammary gland branching involved in thelarche,0.00387223551140303

GO:1902746,0.00297249783557214,Inf,0.109102730819246,2,2,regulation of lens fiber cell differentiation,0.00387223551140303

GO:1902871,0.00297249783557214,Inf,0.109102730819246,2,2,positive regulation of amacrine cell differentiation,0.00387223551140303

GO:0060535,0.00297249783557214,Inf,0.109102730819246,2,2,trachea cartilage morphogenesis,0.00387223551140303

GO:0032696,0.00297249783557214,Inf,0.109102730819246,2,2,negative regulation of interleukin-13 production,0.00387223551140303

GO:0016340,0.00297249783557214,Inf,0.109102730819246,2,2,calcium-dependent cell-matrix adhesion,0.00387223551140303

GO:1903392,0.00297249783557214,Inf,0.109102730819246,2,2,negative regulation of adherens junction organization,0.00387223551140303

GO:0061325,0.00297249783557214,Inf,0.109102730819246,2,2,cell proliferation involved in outflow tract morphogenesis,0.00387223551140303

GO:0061316,0.00297249783557214,Inf,0.109102730819246,2,2,canonical Wnt signaling pathway involved in heart development,0.00387223551140303

GO:1903463,0.00297249783557214,Inf,0.109102730819246,2,2,regulation of mitotic cell cycle DNA replication,0.00387223551140303

GO:0061209,0.00297249783557214,Inf,0.109102730819246,2,2,cell proliferation involved in mesonephros development,0.00387223551140303

GO:0051885,0.00297249783557214,Inf,0.109102730819246,2,2,positive regulation of timing of anagen,0.00387223551140303

GO:0072050,0.00297249783557214,Inf,0.109102730819246,2,2,S-shaped body morphogenesis,0.00387223551140303

GO:0072138,0.00297249783557214,Inf,0.109102730819246,2,2,mesenchymal cell proliferation involved in ureteric bud development,0.00387223551140303

GO:0099566,0.00297249783557214,Inf,0.109102730819246,2,2,regulation of postsynaptic cytosolic calcium ion concentration,0.00387223551140303

GO:0018400,0.00297249783557214,Inf,0.109102730819246,2,2,peptidyl-proline hydroxylation to 3-hydroxy-L-proline,0.00387223551140303

GO:1904954,0.00297249783557214,Inf,0.109102730819246,2,2,canonical Wnt signaling pathway involved in midbrain dopaminergic neuron differentiation,0.00387223551140303

GO:0045720,0.00297249783557214,Inf,0.109102730819246,2,2,negative regulation of integrin biosynthetic process,0.00387223551140303

GO:1901202,0.00297249783557214,Inf,0.109102730819246,2,2,negative regulation of extracellular matrix assembly,0.00387223551140303

GO:0021775,0.00297249783557214,Inf,0.109102730819246,2,2,smoothened signaling pathway involved in ventral spinal cord interneuron specification,0.00387223551140303

GO:0021776,0.00297249783557214,Inf,0.109102730819246,2,2,smoothened signaling pathway involved in spinal cord motor neuron cell fate specification,0.00387223551140303

GO:0021869,0.00297249783557214,Inf,0.109102730819246,2,2,forebrain ventricular zone progenitor cell division,0.00387223551140303

GO:0021521,0.00297249783557214,Inf,0.109102730819246,2,2,ventral spinal cord interneuron specification,0.00387223551140303

GO:0060066,0.00297249783557214,Inf,0.109102730819246,2,2,oviduct development,0.00387223551140303

GO:0036022,0.00297249783557214,Inf,0.109102730819246,2,2,limb joint morphogenesis,0.00387223551140303

GO:0060364,0.00297249783557214,Inf,0.109102730819246,2,2,frontal suture morphogenesis,0.00387223551140303

GO:0097338,0.00297249783557214,Inf,0.109102730819246,2,2,response to clozapine,0.00387223551140303

GO:0042262,0.00297249783557214,Inf,0.109102730819246,2,2,DNA protection,0.00387223551140303

GO:0097168,0.00297249783557214,Inf,0.109102730819246,2,2,mesenchymal stem cell proliferation,0.00387223551140303

GO:0060278,0.00297249783557214,Inf,0.109102730819246,2,2,regulation of ovulation,0.00387223551140303

GO:0060279,0.00297249783557214,Inf,0.109102730819246,2,2,positive regulation of ovulation,0.00387223551140303

GO:0035023,0.00301573260537149,2.20747340357572,8.78276983094928,18,161,regulation of Rho protein signal transduction,0.00392300809258071

GO:0030029,0.00305212097780492,1.52843304843305,38.5132639791938,56,706,actin filament-based process,0.00392957460855154

GO:0090091,0.00305918023271602,8.70239520958084,0.654616384915475,4,12,positive regulation of extracellular matrix disassembly,0.00392957460855154

GO:0071372,0.00305918023271602,8.70239520958084,0.654616384915475,4,12,cellular response to follicle-stimulating hormone stimulus,0.00392957460855154

GO:0071107,0.00305918023271602,8.70239520958084,0.654616384915475,4,12,response to parathyroid hormone,0.00392957460855154

GO:1903651,0.00305918023271602,8.70239520958084,0.654616384915475,4,12,positive regulation of cytoplasmic transport,0.00392957460855154

GO:0048557,0.00305918023271602,8.70239520958084,0.654616384915475,4,12,embryonic digestive tract morphogenesis,0.00392957460855154

GO:0048535,0.00305918023271602,8.70239520958084,0.654616384915475,4,12,lymph node development,0.00392957460855154

GO:0048634,0.00305918023271602,8.70239520958084,0.654616384915475,4,12,regulation of muscle organ development,0.00392957460855154

GO:1905048,0.00305918023271602,8.70239520958084,0.654616384915475,4,12,regulation of metallopeptidase activity,0.00392957460855154

GO:0034109,0.00324641598631379,3.06520220938353,3.65494148244473,10,67,homotypic cell-cell adhesion,0.00415001162586138

GO:0035050,0.00324641598631379,3.06520220938353,3.65494148244473,10,67,embryonic heart tube development,0.00415001162586138

GO:0009799,0.00324881474728127,2.31686131166966,7.47353706111834,16,137,specification of symmetry,0.00415001162586138

GO:0009855,0.00324881474728127,2.31686131166966,7.47353706111834,16,137,determination of bilateral symmetry,0.00415001162586138

GO:0001701,0.0033807185473179,1.88170622446965,14.6743172951886,26,269,in utero embryonic development,0.00431252324387782

GO:0032924,0.00343094736624871,4.06959134615385,2.01840052015605,7,37,activin receptor signaling pathway,0.00436450624905395

GO:0010770,0.00343094736624871,4.06959134615385,2.01840052015605,7,37,positive regulation of cell morphogenesis,0.00436450624905395

GO:0003209,0.00346857761575164,4.7535741569355,1.52743823146944,6,28,cardiac atrium morphogenesis,0.00440022036378411

GO:0033363,0.00346857761575164,4.7535741569355,1.52743823146944,6,28,secretory granule organization,0.00440022036378411

GO:1900024,0.00348551832265856,3.27402108433735,3.1094278283485,9,57,regulation of substrate adhesion-dependent cell spreading,0.00441562912677927

GO:0006790,0.00352180403081433,1.9614089567866,12.4922626788036,23,229,sulfur compound metabolic process,0.00445546911041209

GO:0001570,0.00362457423514713,3.01214591739112,3.70949284785436,10,68,vasculogenesis,0.00457919460983608

GO:0043537,0.00372526835853445,5.80575539568345,1.09102730819246,5,20,negative regulation of blood vessel endothelial cell migration,0.00469996186056195

GO:0071407,0.00380273354650443,1.72567759607101,20.1840052015605,33,370,cellular response to organic cyclic compound,0.0047911321427231

GO:0050807,0.00392431216958474,2.05213889424416,10.419310793238,20,191,regulation of synapse organization,0.00493755670517424

GO:1903035,0.00393066409799453,3.20698303417753,3.16397919375813,9,58,negative regulation of response to wounding,0.0049388016838376

GO:0032370,0.00401153660284049,3.93804280397022,2.07295188556567,7,38,positive regulation of lipid transport,0.00503354933408187

GO:0048609,0.00403184453795744,1.58171875547721,29.8941482444733,45,548,multicellular organismal reproductive process,0.00505151086316382

GO:1902106,0.00403682084178998,2.96088814377134,3.76404421326398,10,69,negative regulation of leukocyte differentiation,0.00505151086316382

GO:0043588,0.00416320685925409,2.04006587727518,10.4738621586476,20,192,skin development,0.00520259636007194

GO:0035909,0.00417505995052573,4.54658385093168,1.58198959687906,6,29,aorta morphogenesis,0.00521033904394877

GO:0060973,0.00422976863686391,7.73493013972056,0.709167750325097,4,13,cell migration involved in heart development,0.00522200658787086

GO:0071397,0.00422976863686391,7.73493013972056,0.709167750325097,4,13,cellular response to cholesterol,0.00522200658787086

GO:0061323,0.00422976863686391,7.73493013972056,0.709167750325097,4,13,cell proliferation involved in heart morphogenesis,0.00522200658787086

GO:0051284,0.00422976863686391,7.73493013972056,0.709167750325097,4,13,positive regulation of sequestering of calcium ion,0.00522200658787086

GO:0048339,0.00422976863686391,7.73493013972056,0.709167750325097,4,13,paraxial mesoderm development,0.00522200658787086

GO:0010715,0.00422976863686391,7.73493013972056,0.709167750325097,4,13,regulation of extracellular matrix disassembly,0.00522200658787086

GO:0009950,0.00422976863686391,7.73493013972056,0.709167750325097,4,13,dorsal/ventral axis specification,0.00522200658787086

GO:0042474,0.00422976863686391,7.73493013972056,0.709167750325097,4,13,middle ear morphogenesis,0.00522200658787086

GO:0046847,0.00441906123859991,3.1426265060241,3.21853055916775,9,59,filopodium assembly,0.00544840080421756

GO:0008543,0.00448523561946696,2.91133896260555,3.8185955786736,10,70,fibroblast growth factor receptor signaling pathway,0.00550786934070542

GO:0001843,0.00448523561946696,2.91133896260555,3.8185955786736,10,70,neural tube closure,0.00550786934070542

GO:0021761,0.00448523561946696,2.91133896260555,3.8185955786736,10,70,limbic system development,0.00550786934070542

GO:0071774,0.00456887519983414,2.59035335214295,5.07327698309493,12,93,response to fibroblast growth factor,0.00560310793481656

GO:0050819,0.00466359731296622,3.81471604567308,2.12750325097529,7,39,negative regulation of coagulation,0.00568539417941976

GO:0060914,0.00467301128536456,5.44252098321343,1.14557867360208,5,21,heart formation,0.00568539417941976

GO:0062009,0.00467301128536456,5.44252098321343,1.14557867360208,5,21,secondary palate development,0.00568539417941976

GO:0001946,0.00467301128536456,5.44252098321343,1.14557867360208,5,21,lymphangiogenesis,0.00568539417941976

GO:1905207,0.00467301128536456,5.44252098321343,1.14557867360208,5,21,regulation of cardiocyte differentiation,0.00568539417941976

GO:0060441,0.00467301128536456,5.44252098321343,1.14557867360208,5,21,epithelial tube branching involved in lung morphogenesis,0.00568539417941976

GO:0008219,0.00479353143550812,1.3468103961908,79.4813394018205,102,1457,cell death,0.00573076010134008

GO:0060638,0.00479741155063323,13.0415669856459,0.38185955786736,3,7,mesenchymal-epithelial cell signaling,0.00573076010134008

GO:1902961,0.00479741155063323,13.0415669856459,0.38185955786736,3,7,positive regulation of aspartic-type endopeptidase activity involved in amyloid precursor protein catabolic process,0.00573076010134008

GO:0043376,0.00479741155063323,13.0415669856459,0.38185955786736,3,7,regulation of CD8-positive, alpha-beta T cell differentiation,0.00573076010134008

GO:0051798,0.00479741155063323,13.0415669856459,0.38185955786736,3,7,positive regulation of hair follicle development,0.00573076010134008

GO:0017185,0.00479741155063323,13.0415669856459,0.38185955786736,3,7,peptidyl-lysine hydroxylation,0.00573076010134008

GO:0048340,0.00479741155063323,13.0415669856459,0.38185955786736,3,7,paraxial mesoderm morphogenesis,0.00573076010134008

GO:2000015,0.00479741155063323,13.0415669856459,0.38185955786736,3,7,regulation of determination of dorsal identity,0.00573076010134008

GO:0072178,0.00479741155063323,13.0415669856459,0.38185955786736,3,7,nephric duct morphogenesis,0.00573076010134008

GO:0048570,0.00479741155063323,13.0415669856459,0.38185955786736,3,7,notochord morphogenesis,0.00573076010134008

GO:0018158,0.00479741155063323,13.0415669856459,0.38185955786736,3,7,protein oxidation,0.00573076010134008

GO:0045602,0.00479741155063323,13.0415669856459,0.38185955786736,3,7,negative regulation of endothelial cell differentiation,0.00573076010134008

GO:0021546,0.00479741155063323,13.0415669856459,0.38185955786736,3,7,rhombomere development,0.00573076010134008

GO:0060391,0.00479741155063323,13.0415669856459,0.38185955786736,3,7,positive regulation of SMAD protein signal transduction,0.00573076010134008

GO:0044283,0.00491148095942719,1.65282534246575,22.9115734720416,36,420,small molecule biosynthetic process,0.00585942223268451

GO:0003143,0.00495346092185698,3.08079376328845,3.27308192457737,9,60,embryonic heart tube morphogenesis,0.00589819188653116

GO:1903707,0.00497194698104122,2.86341434475667,3.87314694408323,10,71,negative regulation of hemopoiesis,0.00589819188653116

GO:0003230,0.00498240313541937,4.35684273709484,1.63654096228869,6,30,cardiac atrium development,0.00589819188653116

GO:0022602,0.00498240313541937,4.35684273709484,1.63654096228869,6,30,ovulation cycle process,0.00589819188653116

GO:0071825,0.00498240313541937,4.35684273709484,1.63654096228869,6,30,protein-lipid complex organization,0.00589819188653116

GO:2001239,0.00498240313541937,4.35684273709484,1.63654096228869,6,30,regulation of extrinsic apoptotic signaling pathway in absence of ligand,0.00589819188653116

GO:0043069,0.00499773248465141,1.53784422580113,32.7308192457737,48,600,negative regulation of programmed cell death,0.00590874405438247

GO:0050680,0.00501638822067938,2.44505194095134,5.78244473342003,13,106,negative regulation of epithelial cell proliferation,0.00591645830717454

GO:0042981,0.00501710525288091,1.40598850690594,56.6243172951886,76,1038,regulation of apoptotic process,0.00591645830717454

GO:0001934,0.00507479866287572,1.60439185345318,26.184655396619,40,480,positive regulation of protein phosphorylation,0.00597684088044571

GO:0032102,0.00522118049771405,1.81336027645991,15.1652795838752,26,278,negative regulation of response to external stimulus,0.00614138855478243

GO:0006898,0.00524269124259224,1.99314574314574,10.6920676202861,20,196,receptor-mediated endocytosis,0.00615882478881053

GO:0072089,0.00530346869832603,3.3233625580196,2.72756827048114,8,50,stem cell proliferation,0.00622228620529717

GO:0043066,0.00535478412050511,1.53864337822671,32.0216514954486,47,587,negative regulation of apoptotic process,0.00627449895036286

GO:0001942,0.00549912814224647,2.81703568232227,3.92769830949285,10,72,hair follicle development,0.00643544729226048

GO:0016053,0.00554663709521812,1.91147278742562,12.2195058517555,22,224,organic acid biosynthetic process,0.00648280807702523

GO:0003183,0.0056688450926328,6.96095808383234,0.76371911573472,4,14,mitral valve morphogenesis,0.00650998295550475

GO:0003174,0.0056688450926328,6.96095808383234,0.76371911573472,4,14,mitral valve development,0.00650998295550475

GO:0042994,0.0056688450926328,6.96095808383234,0.76371911573472,4,14,cytoplasmic sequestering of transcription factor,0.00650998295550475

GO:0090103,0.0056688450926328,6.96095808383234,0.76371911573472,4,14,cochlea morphogenesis,0.00650998295550475

GO:0051797,0.0056688450926328,6.96095808383234,0.76371911573472,4,14,regulation of hair follicle development,0.00650998295550475

GO:0048070,0.0056688450926328,6.96095808383234,0.76371911573472,4,14,regulation of developmental pigmentation,0.00650998295550475

GO:0072176,0.0056688450926328,6.96095808383234,0.76371911573472,4,14,nephric duct development,0.00650998295550475

GO:0009067,0.0056688450926328,6.96095808383234,0.76371911573472,4,14,aspartate family amino acid biosynthetic process,0.00650998295550475

GO:0014029,0.0056688450926328,6.96095808383234,0.76371911573472,4,14,neural crest formation,0.00650998295550475

GO:0010453,0.0056688450926328,6.96095808383234,0.76371911573472,4,14,regulation of cell fate commitment,0.00650998295550475

GO:0120305,0.0056688450926328,6.96095808383234,0.76371911573472,4,14,regulation of pigmentation,0.00650998295550475

GO:0021904,0.0056688450926328,6.96095808383234,0.76371911573472,4,14,dorsal/ventral neural tube patterning,0.00650998295550475

GO:0060444,0.0056688450926328,6.96095808383234,0.76371911573472,4,14,branching involved in mammary gland duct morphogenesis,0.00650998295550475

GO:0036315,0.0056688450926328,6.96095808383234,0.76371911573472,4,14,cellular response to sterol,0.00650998295550475

GO:1902895,0.00578018081714457,5.122020031034,1.2001300390117,5,22,positive regulation of miRNA transcription,0.00661310128272069

GO:0060561,0.00578018081714457,5.122020031034,1.2001300390117,5,22,apoptotic process involved in morphogenesis,0.00661310128272069

GO:0044060,0.00578018081714457,5.122020031034,1.2001300390117,5,22,regulation of endocrine process,0.00661310128272069

GO:0006915,0.00580412427439767,1.34364898609355,76.4264629388817,98,1401,apoptotic process,0.00663225614977699

GO:0009066,0.00589864034832628,4.18228091236495,1.69109232769831,6,31,aspartate family amino acid metabolic process,0.0067163883718621

GO:0001974,0.00589864034832628,4.18228091236495,1.69109232769831,6,31,blood vessel remodeling,0.0067163883718621

GO:0043067,0.00589962887387236,1.39220840535177,57.8789986996099,77,1061,regulation of programmed cell death,0.0067163883718621

GO:0001947,0.00599670809294679,3.24585117398483,2.78211963589077,8,51,heart looping,0.00681007170604685

GO:1905954,0.00599670809294679,3.24585117398483,2.78211963589077,8,51,positive regulation of lipid localization,0.00681007170604685

GO:0022404,0.00606899308928376,2.77212935837785,3.98224967490247,10,73,molting cycle process,0.00687520619339525

GO:0022405,0.00606899308928376,2.77212935837785,3.98224967490247,10,73,hair cycle process,0.00687520619339525

GO:2000177,0.00617154053194115,2.96412821095704,3.38218465539662,9,62,regulation of neural precursor cell proliferation,0.00698278726034128

GO:0010799,0.00620335554349847,3.58982607466063,2.23660598179454,7,41,regulation of peptidyl-threonine phosphorylation,0.00700158144063982

GO:0003382,0.00620335554349847,3.58982607466063,2.23660598179454,7,41,epithelial cell morphogenesis,0.00700158144063982

GO:0090066,0.0062307167153871,1.64964837534008,21.6568920676203,34,397,regulation of anatomical structure size,0.00702385568527726

GO:0098727,0.0063503784260965,2.36815223970944,5.9460988296489,13,109,maintenance of cell number,0.00714126804692903

GO:0019827,0.0063503784260965,2.36815223970944,5.9460988296489,13,109,stem cell population maintenance,0.00714126804692903

GO:0030036,0.00650757338906782,1.50651403412218,34.0400520156047,49,624,actin cytoskeleton organization,0.00730911596503836

GO:0072659,0.00654411404319375,1.851889953229,13.1468790637191,23,241,protein localization to plasma membrane,0.00734120466964853

GO:0008037,0.0066465961659721,2.26781144781145,6.65526657997399,14,122,cell recognition,0.00744709862391764

GO:0034110,0.00693178979143523,4.02114692030658,1.74564369310793,6,32,regulation of homotypic cell-cell adhesion,0.00775720339964987

GO:1902903,0.00702139967788356,1.72699071970926,17.074577373212,28,313,regulation of supramolecular fiber organization,0.00784794794093539

GO:0043371,0.00705990353187377,4.83713029576339,1.25468140442133,5,23,negative regulation of CD4-positive, alpha-beta T cell differentiation,0.00787187790902633

GO:2000050,0.00705990353187377,4.83713029576339,1.25468140442133,5,23,regulation of non-canonical Wnt signaling pathway,0.00787187790902633

GO:0042471,0.00713568154305367,2.2468501529052,6.70981794538362,14,123,ear morphogenesis,0.00793715302071549

GO:0007368,0.00713568154305367,2.2468501529052,6.70981794538362,14,123,determination of left/right symmetry,0.00793715302071549

GO:0002761,0.00734580842238963,2.68646190962234,4.09135240572172,10,75,regulation of myeloid leukocyte differentiation,0.00803797215295384

GO:0031099,0.00734581301939327,2.10606711188664,8.12815344603381,16,149,regeneration,0.00803797215295384

GO:0012501,0.00735893421221488,1.32653861388528,78.8812743823147,100,1446,programmed cell death,0.00803797215295384

GO:0032274,0.00736640799181555,10.4325358851675,0.436410923276983,3,8,gonadotropin secretion,0.00803797215295384

GO:0060716,0.00736640799181555,10.4325358851675,0.436410923276983,3,8,labyrinthine layer blood vessel development,0.00803797215295384

GO:1902959,0.00736640799181555,10.4325358851675,0.436410923276983,3,8,regulation of aspartic-type endopeptidase activity involved in amyloid precursor protein catabolic process,0.00803797215295384

GO:1903054,0.00736640799181555,10.4325358851675,0.436410923276983,3,8,negative regulation of extracellular matrix organization,0.00803797215295384

GO:0071694,0.00736640799181555,10.4325358851675,0.436410923276983,3,8,maintenance of protein location in extracellular region,0.00803797215295384

GO:0062042,0.00736640799181555,10.4325358851675,0.436410923276983,3,8,regulation of cardiac epithelial to mesenchymal transition,0.00803797215295384

GO:0062043,0.00736640799181555,10.4325358851675,0.436410923276983,3,8,positive regulation of cardiac epithelial to mesenchymal transition,0.00803797215295384

GO:0099545,0.00736640799181555,10.4325358851675,0.436410923276983,3,8,trans-synaptic signaling by trans-synaptic complex,0.00803797215295384

GO:1901203,0.00736640799181555,10.4325358851675,0.436410923276983,3,8,positive regulation of extracellular matrix assembly,0.00803797215295384

GO:0035583,0.00736640799181555,10.4325358851675,0.436410923276983,3,8,sequestering of TGFbeta in extracellular matrix,0.00803797215295384

GO:1901620,0.00736640799181555,10.4325358851675,0.436410923276983,3,8,regulation of smoothened signaling pathway involved in dorsal/ventral neural tube patterning,0.00803797215295384

GO:0006563,0.00736640799181555,10.4325358851675,0.436410923276983,3,8,L-serine metabolic process,0.00803797215295384

GO:0060019,0.00736640799181555,10.4325358851675,0.436410923276983,3,8,radial glial cell differentiation,0.00803797215295384

GO:0071371,0.00740086903985327,6.32770821992379,0.818270481144343,4,15,cellular response to gonadotropin stimulus,0.00803797215295384

GO:0006103,0.00740086903985327,6.32770821992379,0.818270481144343,4,15,2-oxoglutarate metabolic process,0.00803797215295384

GO:0045601,0.00740086903985327,6.32770821992379,0.818270481144343,4,15,regulation of endothelial cell differentiation,0.00803797215295384

GO:0060413,0.00740086903985327,6.32770821992379,0.818270481144343,4,15,atrial septum morphogenesis,0.00803797215295384

GO:0035306,0.00758724985405795,3.10116325711994,2.89122236671001,8,53,positive regulation of dephosphorylation,0.00823069153779431

GO:0007492,0.00760788764537662,2.85594742606791,3.49128738621586,9,64,endoderm development,0.00823862694202177

GO:0010647,0.00761245551320361,1.35480362566461,65.5707412223667,85,1202,positive regulation of cell communication,0.00823862694202177

GO:0035051,0.00782038280517438,2.09020511053481,8.18270481144343,16,150,cardiocyte differentiation,0.00845372366615682

GO:0046849,0.00805735136744899,2.64557517271631,4.14590377113134,10,76,bone remodeling,0.00867560390470432

GO:0021510,0.00805735136744899,2.64557517271631,4.14590377113134,10,76,spinal cord development,0.00867560390470432

GO:0021954,0.00805735136744899,2.64557517271631,4.14590377113134,10,76,central nervous system neuron development,0.00867560390470432

GO:0060986,0.0080898059241567,3.8719487795118,1.80019505851756,6,33,endocrine hormone secretion,0.00867560390470432

GO:0043392,0.0080898059241567,3.8719487795118,1.80019505851756,6,33,negative regulation of DNA binding,0.00867560390470432

GO:0002763,0.0080898059241567,3.8719487795118,1.80019505851756,6,33,positive regulation of myeloid leukocyte differentiation,0.00867560390470432

GO:0071695,0.00810017961178473,1.98313147794912,9.65559167750325,18,177,anatomical structure maturation,0.00867560390470432

GO:0023056,0.0081009982172049,1.35087272641152,65.7343953185956,85,1205,positive regulation of signaling,0.00867560390470432

GO:0060972,0.00819964530710258,2.20606060606061,6.81892067620286,14,125,left/right pattern formation,0.00875272104197151

GO:1901136,0.00819964530710258,2.20606060606061,6.81892067620286,14,125,carbohydrate derivative catabolic process,0.00875272104197151

GO:0048660,0.00821102169409705,2.49551414768806,4.80052015604681,11,88,regulation of smooth muscle cell proliferation,0.00875272104197151

GO:0021782,0.00821102169409705,2.49551414768806,4.80052015604681,11,88,glial cell development,0.00875272104197151

GO:0030307,0.00831275763872449,2.1338503433578,7.52808842652796,15,138,positive regulation of cell growth,0.00877494173593687

GO:0050886,0.00849233105205521,3.03353738293308,2.94577373211964,8,54,endocrine process,0.00877494173593687

GO:0090398,0.00849233105205521,3.03353738293308,2.94577373211964,8,54,cellular senescence,0.00877494173593687

GO:0060762,0.00859391687927802,34.7431302270012,0.163654096228869,2,3,regulation of branching involved in mammary gland duct morphogenesis,0.00877494173593687

GO:0060770,0.00859391687927802,34.7431302270012,0.163654096228869,2,3,negative regulation of epithelial cell proliferation involved in prostate gland development,0.00877494173593687

GO:0046543,0.00859391687927802,34.7431302270012,0.163654096228869,2,3,development of secondary female sexual characteristics,0.00877494173593687

GO:1902730,0.00859391687927802,34.7431302270012,0.163654096228869,2,3,positive regulation of proteoglycan biosynthetic process,0.00877494173593687

GO:1902732,0.00859391687927802,34.7431302270012,0.163654096228869,2,3,positive regulation of chondrocyte proliferation,0.00877494173593687

GO:1902869,0.00859391687927802,34.7431302270012,0.163654096228869,2,3,regulation of amacrine cell differentiation,0.00877494173593687

GO:0060573,0.00859391687927802,34.7431302270012,0.163654096228869,2,3,cell fate specification involved in pattern specification,0.00877494173593687

GO:0060648,0.00859391687927802,34.7431302270012,0.163654096228869,2,3,mammary gland bud morphogenesis,0.00877494173593687

GO:0090009,0.00859391687927802,34.7431302270012,0.163654096228869,2,3,primitive streak formation,0.00877494173593687

GO:0061073,0.00859391687927802,34.7431302270012,0.163654096228869,2,3,ciliary body morphogenesis,0.00877494173593687

GO:0048073,0.00859391687927802,34.7431302270012,0.163654096228869,2,3,regulation of eye pigmentation,0.00877494173593687

GO:0038031,0.00859391687927802,34.7431302270012,0.163654096228869,2,3,non-canonical Wnt signaling pathway via JNK cascade,0.00877494173593687

GO:0072125,0.00859391687927802,34.7431302270012,0.163654096228869,2,3,negative regulation of glomerular mesangial cell proliferation,0.00877494173593687

GO:1900126,0.00859391687927802,34.7431302270012,0.163654096228869,2,3,negative regulation of hyaluronan biosynthetic process,0.00877494173593687

GO:1904796,0.00859391687927802,34.7431302270012,0.163654096228869,2,3,regulation of core promoter binding,0.00877494173593687

GO:1904798,0.00859391687927802,34.7431302270012,0.163654096228869,2,3,positive regulation of core promoter binding,0.00877494173593687

GO:0048696,0.00859391687927802,34.7431302270012,0.163654096228869,2,3,regulation of collateral sprouting in absence of injury,0.00877494173593687

GO:0048669,0.00859391687927802,34.7431302270012,0.163654096228869,2,3,collateral sprouting in absence of injury,0.00877494173593687

GO:2000833,0.00859391687927802,34.7431302270012,0.163654096228869,2,3,positive regulation of steroid hormone secretion,0.00877494173593687

GO:1905071,0.00859391687927802,34.7431302270012,0.163654096228869,2,3,tight junction disassembly,0.00877494173593687

GO:1905073,0.00859391687927802,34.7431302270012,0.163654096228869,2,3,regulation of tight junction disassembly,0.00877494173593687

GO:1905075,0.00859391687927802,34.7431302270012,0.163654096228869,2,3,positive regulation of tight junction disassembly,0.00877494173593687

GO:0010908,0.00859391687927802,34.7431302270012,0.163654096228869,2,3,regulation of heparan sulfate proteoglycan biosynthetic process,0.00877494173593687

GO:0010909,0.00859391687927802,34.7431302270012,0.163654096228869,2,3,positive regulation of heparan sulfate proteoglycan biosynthetic process,0.00877494173593687

GO:1905310,0.00859391687927802,34.7431302270012,0.163654096228869,2,3,regulation of cardiac neural crest cell migration involved in outflow tract morphogenesis,0.00877494173593687

GO:1905312,0.00859391687927802,34.7431302270012,0.163654096228869,2,3,positive regulation of cardiac neural crest cell migration involved in outflow tract morphogenesis,0.00877494173593687

GO:0045843,0.00859391687927802,34.7431302270012,0.163654096228869,2,3,negative regulation of striated muscle tissue development,0.00877494173593687

GO:0019287,0.00859391687927802,34.7431302270012,0.163654096228869,2,3,isopentenyl diphosphate biosynthetic process, mevalonate pathway,0.00877494173593687

GO:0021910,0.00859391687927802,34.7431302270012,0.163654096228869,2,3,smoothened signaling pathway involved in ventral spinal cord patterning,0.00877494173593687

GO:0021943,0.00859391687927802,34.7431302270012,0.163654096228869,2,3,formation of radial glial scaffolds,0.00877494173593687

GO:0070483,0.00859391687927802,34.7431302270012,0.163654096228869,2,3,detection of hypoxia,0.00877494173593687

GO:0060363,0.00859391687927802,34.7431302270012,0.163654096228869,2,3,cranial suture morphogenesis,0.00877494173593687

GO:0097187,0.00859391687927802,34.7431302270012,0.163654096228869,2,3,dentinogenesis,0.00877494173593687

GO:0042701,0.00859391687927802,34.7431302270012,0.163654096228869,2,3,progesterone secretion,0.00877494173593687

GO:0003420,0.00859391687927802,34.7431302270012,0.163654096228869,2,3,regulation of growth plate cartilage chondrocyte proliferation,0.00877494173593687

GO:0060326,0.00897644586878029,1.88467446531963,11.2375812743823,20,206,cell chemotaxis,0.00915537834456993

GO:0050714,0.00918607994385625,2.25013784671445,6.21885565669701,13,114,positive regulation of protein secretion,0.00935882702244646

GO:0032870,0.00934268307644447,1.57681083994721,23.8934980494148,36,438,cellular response to hormone stimulus,0.00949981533467126

GO:0045746,0.00938053217814899,3.7334076487738,1.85474642392718,6,34,negative regulation of Notch signaling pathway,0.00949981533467126

GO:0044272,0.00938544543747417,2.16671493698042,6.92802340702211,14,127,sulfur compound biosynthetic process,0.00949981533467126

GO:0007010,0.00944299143741337,1.33257975890104,69.716644993498,89,1278,cytoskeleton organization,0.00949981533467126

GO:0060674,0.00944824196151886,5.8,0.872821846553966,4,16,placenta blood vessel development,0.00949981533467126

GO:0071498,0.00944824196151886,5.8,0.872821846553966,4,16,cellular response to fluid shear stress,0.00949981533467126

GO:0043567,0.00944824196151886,5.8,0.872821846553966,4,16,regulation of insulin-like growth factor receptor signaling pathway,0.00949981533467126

GO:0030033,0.00944824196151886,5.8,0.872821846553966,4,16,microvillus assembly,0.00949981533467126

GO:0072111,0.00944824196151886,5.8,0.872821846553966,4,16,cell proliferation involved in kidney development,0.00949981533467126

GO:2000725,0.00944824196151886,5.8,0.872821846553966,4,16,regulation of cardiac muscle cell differentiation,0.00949981533467126

GO:0030539,0.00944824196151886,5.8,0.872821846553966,4,16,male genitalia development,0.00949981533467126

GO:1901522,0.00944824196151886,5.8,0.872821846553966,4,16,positive regulation of transcription from RNA polymerase II promoter involved in cellular response to chemical stimulus,0.00949981533467126

GO:0042327,0.00947524478045756,1.52810143752616,28.0394018205462,41,514,positive regulation of phosphorylation,0.00951657627350208

GO:0046394,0.00953581274812648,1.84083740831296,12.0558517555267,21,221,carboxylic acid biosynthetic process,0.00956697553488506

GO:0046887,0.00963836589629839,2.56740935216065,4.25500650195058,10,78,positive regulation of hormone secretion,0.00965934166538718

GO:0048659,0.00967840712643494,2.43200024460344,4.90962288686606,11,90,smooth muscle cell proliferation,0.00968892713418107

GO:1902904,0.00996957234046918,2.02905594590311,8.40091027308192,16,154,negative regulation of supramolecular fiber organization,0.00996957234046918

Table S4. Supplementary data to accompany Figure 3D. Top gene ontology (Functional Enrichment Analysis) for each compartment (LT).

GOBPID,Pvalue,OddsRatio,ExpCount,Count,Size,Term,FDR

GO:0030574,1.46779885225061e-06,15.2216457702839,0.583615084525358,7,34,collagen catabolic process,0.00031117335667713

GO:0032963,7.3234824257708e-06,7.69667519181586,1.33888166449935,9,78,collagen metabolic process,0.000776289137131705

GO:0032940,1.26263097329453e-05,2.54134325632362,13.4403120936281,31,783,secretion by cell,0.000874382695537579

GO:2000147,1.68551299362343e-05,3.11534737351731,7.36384915474642,21,429,positive regulation of cell motility,0.000874382695537579

GO:0140352,2.43936121800892e-05,2.44864936338543,13.9037711313394,31,810,export from cell,0.000874382695537579

GO:0040017,2.53746654850016e-05,3.0238683127572,7.56983094928479,21,441,positive regulation of locomotion,0.000874382695537579

GO:0001934,2.90685061419445e-05,2.90948789202064,8.23927178153446,22,480,positive regulation of protein phosphorylation,0.000874382695537579

GO:0055088,3.36465090176848e-05,6.24124567474048,1.61352405721717,9,94,lipid homeostasis,0.000874382695537579

GO:0030335,3.71200200935765e-05,3.03113930307274,7.17503250975293,20,418,positive regulation of cell migration,0.000874382695537579

GO:0046849,4.66199369142282e-05,6.91544117647059,1.30455136540962,8,76,bone remodeling,0.000898570959608968

GO:0016477,5.5971730028985e-05,2.12500584795322,20.237711313394,39,1179,cell migration,0.000898570959608968

GO:0010562,6.10454300878458e-05,2.62851985559567,9.92145643693108,24,578,positive regulation of phosphorus metabolic process,0.000898570959608968

GO:0045937,6.10454300878458e-05,2.62851985559567,9.92145643693108,24,578,positive regulation of phosphate metabolic process,0.000898570959608968

GO:0032501,6.39683721479544e-05,1.62525704162229,100.158647594278,131,5835,multicellular organismal process,0.000898570959608968

GO:0030198,6.64523217243972e-05,3.35732880588568,5.1667100130039,16,301,extracellular matrix organization,0.000898570959608968

GO:0043062,6.91066197625368e-05,3.34536431310625,5.18387516254876,16,302,extracellular structure organization,0.000898570959608968

GO:0046903,7.20552184592097e-05,2.29880482222833,14.7276983094928,31,858,secretion,0.000898570959608968

GO:0042327,8.03655697100347e-05,2.70214338507021,8.82288686605982,22,514,positive regulation of phosphorylation,0.000946527821029297

GO:0002685,8.73480011592143e-05,4.87896141081538,2.24863459037711,10,131,regulation of leukocyte migration,0.000974619802408075

GO:0002283,0.000112509623791962,4.30895479412612,2.78075422626788,11,162,neutrophil activation involved in immune response,0.00119260201219479

GO:0002687,0.000133141253976019,5.8734375,1.51053315994798,8,88,positive regulation of leukocyte migration,0.00134409265918647

GO:0048870,0.000182501920272907,1.97913098092954,22.1602080624187,40,1291,cell motility,0.00162600637219674

GO:0023052,0.000186771374125183,1.59364520267381,79.5604681404421,107,4635,signaling,0.00162600637219674

GO:0002446,0.000190969814235258,4.03861733729409,2.95240572171652,11,172,neutrophil mediated immunity,0.00162600637219674

GO:0007154,0.000191746034457163,1.58919629627053,81.465799739922,109,4746,cell communication,0.00162600637219674

GO:0045229,0.000199559059148439,3.03143881208397,5.6816644993498,16,331,external encapsulating structure organization,0.00162717386690266

GO:0050900,0.000218115984929467,3.28983399209486,4.58309492847854,14,267,leukocyte migration,0.00166249464424825

GO:0050896,0.000219574764334675,1.55990522735151,112.328738621586,141,6544,response to stimulus,0.00166249464424825

GO:0065009,0.000228673083191717,1.80998201062628,31.8585175552666,52,1856,regulation of molecular function,0.00167167909091876

GO:2000145,0.000245760775523322,2.29443567820141,12.2387516254876,26,713,regulation of cell motility,0.00173670948036481

GO:0072124,0.000262717908697786,34.7379310344828,0.137321196358908,3,8,regulation of glomerular mesangial cell proliferation,0.00179665150464292

GO:0090170,0.00029354537553151,Inf,0.0343302990897269,2,2,regulation of Golgi inheritance,0.00190938802377936

GO:0042119,0.000297216060305277,3.82250639386189,3.10689206762029,11,181,neutrophil activation,0.00190938802377936

GO:0030334,0.000313308980869232,2.29474733493756,11.7409622886866,25,684,regulation of cell migration,0.00195357364541992

GO:0036230,0.000342196061502796,3.75546619753707,3.15838751625488,11,184,granulocyte activation,0.0020118316037224

GO:0048771,0.000343234202204144,4.48594217347956,2.17997399219766,9,127,tissue remodeling,0.0020118316037224

GO:0001932,0.000376022676586444,2.19139850541406,13.2858257477243,27,774,regulation of protein phosphorylation,0.0020118316037224

GO:0072110,0.000389080640342539,28.9463601532567,0.154486345903771,3,9,glomerular mesangial cell proliferation,0.0020118316037224

GO:1901722,0.000389080640342539,28.9463601532567,0.154486345903771,3,9,regulation of cell proliferation involved in kidney development,0.0020118316037224

GO:2001268,0.000389080640342539,28.9463601532567,0.154486345903771,3,9,negative regulation of cysteine-type endopeptidase activity involved in apoptotic signaling pathway,0.0020118316037224

GO:0070486,0.000389080640342539,28.9463601532567,0.154486345903771,3,9,leukocyte aggregation,0.0020118316037224

GO:0043312,0.000424804603620521,3.954711198013,2.72925877763329,10,159,neutrophil degranulation,0.00212736852092566

GO:0038127,0.000431494558489639,4.33770491803279,2.24863459037711,9,131,ERBB signaling pathway,0.00212736852092566

GO:0071677,0.000512539985128749,8.56347944583239,0.669440832249675,5,39,positive regulation of mononuclear cell migration,0.00241463281882877

GO:2000401,0.000512539985128749,8.56347944583239,0.669440832249675,5,39,regulation of lymphocyte migration,0.00241463281882877

GO:0040012,0.000541706619241988,2.16844972471747,12.8910273081925,26,751,regulation of locomotion,0.00247538564163181

GO:0032310,0.000548788326210826,24.8095238095238,0.171651495448635,3,10,prostaglandin secretion,0.00247538564163181

GO:0007188,0.000567275867864182,4.16553960166744,2.33446033810143,9,136,adenylate cyclase-modulating G protein-coupled receptor signaling pathway,0.00250546841640014

GO:0009653,0.000607318288594714,1.66499691985002,40.7157347204161,61,2372,anatomical structure morphogenesis,0.0025598572304073

GO:0071902,0.000614095325509979,4.59987745098039,1.88816644993498,8,110,positive regulation of protein serine/threonine kinase activity,0.0025598572304073

GO:0002274,0.000615814711088549,2.94575886524823,5.08088426527958,14,296,myeloid leukocyte activation,0.0025598572304073

GO:0040011,0.000629508385313805,2.05577649632134,15.1911573472042,29,885,locomotion,0.00256645726320244

GO:0071675,0.00066429720201823,6.25415282392027,1.06423927178153,6,62,regulation of mononuclear cell migration,0.00265718880807292

GO:0045055,0.000742021719741301,2.66720881901148,6.40260078023407,16,373,regulated exocytosis,0.00289874920180636

GO:0006887,0.000752033991034669,2.43264432111174,8.34226267880364,19,486,exocytosis,0.00289874920180636

GO:0032368,0.000787392787673393,6.03769045709703,1.09856957087126,6,64,regulation of lipid transport,0.00297675806318713

GO:2000403,0.000807944699435631,11.0586080586081,0.429128738621586,4,25,positive regulation of lymphocyte migration,0.00297675806318713

GO:0097529,0.000814396073890818,3.94609306409131,2.45461638491547,9,143,myeloid leukocyte migration,0.00297675806318713

GO:0006575,0.000843261275728379,4.99373635759704,1.52769830949285,7,89,cellular modified amino acid metabolic process,0.00303002356702401

GO:0071593,0.000870633670047612,115.381679389313,0.0514954486345904,2,3,lymphocyte aggregation,0.00307623896750156

GO:0007189,0.000962298188366461,4.87418936446174,1.56202860858257,7,91,adenylate cyclase-activating G protein-coupled receptor signaling pathway,0.00324894683774906

GO:0048856,0.000977716332047143,1.4964314964315,85.7227568270481,110,4994,anatomical structure development,0.00324894683774906

GO:0072109,0.00098081413969783,19.29374201788,0.205981794538362,3,12,glomerular mesangium development,0.00324894683774906

GO:0015732,0.00098081413969783,19.29374201788,0.205981794538362,3,12,prostaglandin transport,0.00324894683774906

GO:0051649,0.00101794343132776,1.70425334881112,31.4465539661899,49,1832,establishment of localization in cell,0.00328972807632225

GO:0045860,0.00102770967204778,2.9137389834601,4.75474642392718,13,277,positive regulation of protein kinase activity,0.00328972807632225

GO:0061572,0.00103967821279996,3.80287769784173,2.54044213263979,9,148,actin filament bundle organization,0.00328972807632225

GO:0050790,0.0011260003290806,1.81131971438811,22.6923276983095,38,1322,regulation of catalytic activity,0.00351047161419246

GO:0016192,0.00117137351164911,1.76918277245383,25.1297789336801,41,1464,vesicle-mediated transport,0.00353488589444247

GO:0031401,0.00119529398604644,2.14463544754001,11.4491547464239,23,667,positive regulation of protein modification process,0.00353488589444247

GO:0002444,0.00120052728490499,3.19404583422574,3.67334200260078,11,214,myeloid leukocyte mediated immunity,0.00353488589444247

GO:0002275,0.00120052728490499,3.19404583422574,3.67334200260078,11,214,myeloid cell activation involved in immune response,0.00353488589444247

GO:0045453,0.00122144548358839,6.92866335723479,0.806762028608583,5,47,bone resorption,0.00354721154137998

GO:0042325,0.0012457336029813,2.00947416067053,14.3843953185956,27,838,regulation of phosphorylation,0.00356885843016265

GO:0007264,0.00130153243281225,2.2602187858316,9.42366710013004,20,549,small GTPase mediated signal transduction,0.00367899834341597

GO:0042058,0.00136138131730982,5.38497316636851,1.21872561768531,6,71,regulation of epidermal growth factor receptor signaling pathway,0.0037975373588116

GO:0007165,0.00158469806936302,1.48664011709044,73.6899869960988,96,4293,signal transduction,0.00434478122858858

GO:0006468,0.00160693782424967,1.84556695606419,19.2078023407022,33,1119,protein phosphorylation,0.00434478122858858

GO:0051716,0.0016319634172433,1.46047736423902,93.532899869961,117,5449,cellular response to stimulus,0.00434478122858858

GO:1903225,0.00172151709057283,57.6870229007634,0.0686605981794538,2,4,negative regulation of endodermal cell differentiation,0.00434478122858858

GO:0051913,0.00172151709057283,57.6870229007634,0.0686605981794538,2,4,regulation of synaptic plasticity by chemical substance,0.00434478122858858

GO:0051914,0.00172151709057283,57.6870229007634,0.0686605981794538,2,4,positive regulation of synaptic plasticity by chemical substance,0.00434478122858858

GO:0051915,0.00172151709057283,57.6870229007634,0.0686605981794538,2,4,induction of synaptic plasticity by chemical substance,0.00434478122858858

GO:2000562,0.00172151709057283,57.6870229007634,0.0686605981794538,2,4,negative regulation of CD4-positive, alpha-beta T cell proliferation,0.00434478122858858

GO:1901184,0.00180822297171677,5.07145264577014,1.28738621586476,6,75,regulation of ERBB signaling pathway,0.00450992082357593

GO:0065008,0.00187577159963638,1.61429259233785,36.578933680104,54,2131,regulation of biological quality,0.00462399510608038

GO:2001267,0.00195263432901857,14.4674329501916,0.257477243172952,3,15,regulation of cysteine-type endopeptidase activity involved in apoptotic signaling pathway,0.00475814342243606

GO:0007010,0.00216571209699184,1.76379354182558,21.9370611183355,36,1278,cytoskeleton organization,0.00521739732457125

GO:0009611,0.00223295467386348,2.37965882448056,7.12353706111834,16,415,response to wounding,0.00524026636054881

GO:0048731,0.00233541124118552,1.50474918077462,56.2501950585176,76,3277,system development,0.00524026636054881

GO:0090199,0.0023471228789118,8.00371352785146,0.566449934980494,4,33,regulation of release of cytochrome c from mitochondria,0.00524026636054881

GO:0047497,0.00237295080477682,13.3536693191866,0.274642392717815,3,16,mitochondrion transport along microtubule,0.00524026636054881

GO:0051654,0.00237295080477682,13.3536693191866,0.274642392717815,3,16,establishment of mitochondrion localization,0.00524026636054881

GO:0072111,0.00237295080477682,13.3536693191866,0.274642392717815,3,16,cell proliferation involved in kidney development,0.00524026636054881

GO:0034643,0.00237295080477682,13.3536693191866,0.274642392717815,3,16,establishment of mitochondrion localization, microtubule-mediated,0.00524026636054881

GO:0002693,0.00237295080477682,13.3536693191866,0.274642392717815,3,16,positive regulation of cellular extravasation,0.00524026636054881

GO:0071674,0.00246026579907806,4.08996108949416,1.83667100130039,7,107,mononuclear cell migration,0.00524855426242755

GO:0030029,0.00248125867478753,2.01672509006519,12.1185955786736,23,706,actin filament-based process,0.00524855426242755

GO:0043299,0.0025195187811627,3.07643154553325,3.45019505851756,10,201,leukocyte degranulation,0.00524855426242755

GO:0030036,0.00259650005784727,2.07995004401859,10.7110533159948,21,624,actin cytoskeleton organization,0.00524855426242755

GO:0097435,0.00273876719894907,2.03301953070114,11.4834850455137,22,669,supramolecular fiber organization,0.00524855426242755

GO:0086023,0.00283669676606869,38.4554707379135,0.0858257477243173,2,5,adenylate cyclase-activating adrenergic receptor signaling pathway involved in heart process,0.00524855426242755

GO:0048313,0.00283669676606869,38.4554707379135,0.0858257477243173,2,5,Golgi inheritance,0.00524855426242755

GO:0048308,0.00283669676606869,38.4554707379135,0.0858257477243173,2,5,organelle inheritance,0.00524855426242755

GO:0072126,0.00283669676606869,38.4554707379135,0.0858257477243173,2,5,positive regulation of glomerular mesangial cell proliferation,0.00524855426242755

GO:2000561,0.00283669676606869,38.4554707379135,0.0858257477243173,2,5,regulation of CD4-positive, alpha-beta T cell proliferation,0.00524855426242755

GO:1901724,0.00283669676606869,38.4554707379135,0.0858257477243173,2,5,positive regulation of cell proliferation involved in kidney development,0.00524855426242755

GO:0035739,0.00283669676606869,38.4554707379135,0.0858257477243173,2,5,CD4-positive, alpha-beta T cell proliferation,0.00524855426242755

GO:0006599,0.00283669676606869,38.4554707379135,0.0858257477243173,2,5,phosphagen metabolic process,0.00524855426242755

GO:0006603,0.00283669676606869,38.4554707379135,0.0858257477243173,2,5,phosphocreatine metabolic process,0.00524855426242755

GO:0042396,0.00283669676606869,38.4554707379135,0.0858257477243173,2,5,phosphagen biosynthetic process,0.00524855426242755

GO:0046314,0.00283669676606869,38.4554707379135,0.0858257477243173,2,5,phosphocreatine biosynthetic process,0.00524855426242755

GO:0072012,0.00284515222058198,12.3990147783251,0.291807542262679,3,17,glomerulus vasculature development,0.00524855426242755

GO:0034377,0.00284515222058198,12.3990147783251,0.291807542262679,3,17,plasma lipoprotein particle assembly,0.00524855426242755

GO:1905952,0.00284709311405268,4.60220318237454,1.4075422626788,6,82,regulation of lipid localization,0.00524855426242755

GO:0032502,0.00290624849198036,1.42954810313479,91.0954486345904,113,5307,developmental process,0.00531141965775721

GO:0002526,0.00302575194719054,4.54213228631833,1.42470741222367,6,83,acute inflammatory response,0.00548255908379824

GO:0045859,0.00311486421949586,2.23288309268747,8.05045513654096,17,469,regulation of protein kinase activity,0.00559238640164506

GO:0001775,0.00315290660637802,1.87003441637986,15.3628088426528,27,895,cell activation,0.00559238640164506

GO:0032535,0.00319188091792006,2.54059259650141,5.40702210663199,13,315,regulation of cellular component size,0.00559238640164506

GO:0033674,0.00319188091792006,2.54059259650141,5.40702210663199,13,315,positive regulation of kinase activity,0.00559238640164506

GO:0007173,0.00334204175844784,3.85691212098965,1.93966189856957,7,113,epidermal growth factor receptor signaling pathway,0.00567221485182243

GO:0061440,0.00337122203457371,11.5716475095785,0.308972691807542,3,18,kidney vasculature development,0.00567221485182243

GO:0061437,0.00337122203457371,11.5716475095785,0.308972691807542,3,18,renal system vasculature development,0.00567221485182243

GO:0010743,0.00337122203457371,11.5716475095785,0.308972691807542,3,18,regulation of macrophage derived foam cell differentiation,0.00567221485182243

GO:0070168,0.00337122203457371,11.5716475095785,0.308972691807542,3,18,negative regulation of biomineral tissue development,0.00567221485182243

GO:0019220,0.00345490177707066,1.8156588667227,17.0106631989597,29,991,regulation of phosphate metabolic process,0.00575962495813638

GO:0010256,0.00349962042629546,2.33608116677235,6.78023407022107,15,395,endomembrane system organization,0.00575962495813638

GO:0051174,0.00350467745094148,1.81364530169462,17.0278283485046,29,992,regulation of phosphorus metabolic process,0.00575962495813638

GO:0001836,0.00359067217690178,7.03170163170163,0.635110533159948,4,37,release of cytochrome c from mitochondria,0.00581085879010058

GO:0031638,0.00359067217690178,7.03170163170163,0.635110533159948,4,37,zymogen activation,0.00581085879010058

GO:0090066,0.00366747435590159,2.32353497760676,6.81456436931079,15,397,regulation of anatomical structure size,0.00589018608675104

GO:0072676,0.00390095023447629,5.19167126309983,1.04707412223667,5,61,lymphocyte migration,0.00606711857728206

GO:0015908,0.00390095023447629,5.19167126309983,1.04707412223667,5,61,fatty acid transport,0.00606711857728206

GO:0006869,0.00390993558317061,2.7179393496529,4.274122236671,11,249,lipid transport,0.00606711857728206

GO:0065005,0.00395299542304691,10.8477011494253,0.326137841352406,3,19,protein-lipid complex assembly,0.00606711857728206

GO:0006809,0.00395979685612813,6.82443438914027,0.652275682704811,4,38,nitric oxide biosynthetic process,0.00606711857728206

GO:0046209,0.00395979685612813,6.82443438914027,0.652275682704811,4,38,nitric oxide metabolic process,0.00606711857728206

GO:0065007,0.00407652906971684,1.42478730622684,157.83355006502,179,9195,biological regulation,0.00606711857728206

GO:0051347,0.00412289498666544,2.3694899713467,6.23094928478544,14,363,positive regulation of transferase activity,0.00606711857728206

GO:0046504,0.00420691712670029,28.8396946564886,0.102990897269181,2,6,glycerol ether biosynthetic process,0.00606711857728206

GO:0061307,0.00420691712670029,28.8396946564886,0.102990897269181,2,6,cardiac neural crest cell differentiation involved in heart development,0.00606711857728206

GO:0061308,0.00420691712670029,28.8396946564886,0.102990897269181,2,6,cardiac neural crest cell development involved in heart development,0.00606711857728206

GO:1903224,0.00420691712670029,28.8396946564886,0.102990897269181,2,6,regulation of endodermal cell differentiation,0.00606711857728206

GO:0008611,0.00420691712670029,28.8396946564886,0.102990897269181,2,6,ether lipid biosynthetic process,0.00606711857728206

GO:0044036,0.00420691712670029,28.8396946564886,0.102990897269181,2,6,cell wall macromolecule metabolic process,0.00606711857728206

GO:1901503,0.00420691712670029,28.8396946564886,0.102990897269181,2,6,ether biosynthetic process,0.00606711857728206

GO:2001057,0.00435376906995663,6.62901098901099,0.669440832249675,4,39,reactive nitrogen species metabolic process,0.00623648001912706

GO:0032970,0.00448562791736035,2.43360526149371,5.63016905071521,13,328,regulation of actin filament-based process,0.00638223569449929

GO:0050772,0.00452026921725468,4.16168327796235,1.54486345903771,6,90,positive regulation of axonogenesis,0.00638864716038662

GO:0009719,0.00456114324739612,1.68804258946746,22.1430429128739,35,1290,response to endogenous stimulus,0.00640372429435746

GO:0015718,0.00477142011341148,4.1124487004104,1.56202860858257,6,91,monocarboxylic acid transport,0.00661403810362527

GO:1900026,0.00477333882006918,6.44444444444444,0.686605981794538,4,40,positive regulation of substrate adhesion-dependent cell spreading,0.00661403810362527

GO:0043406,0.00512487746819076,4.84427284427284,1.11573472041612,5,65,positive regulation of MAP kinase activity,0.00705502612504183

GO:0035556,0.00516136301623695,1.52065140429549,39.222366710013,55,2285,intracellular signal transduction,0.0070594126415628

GO:0007166,0.0052195109619013,1.52972977779359,37.5058517555267,53,2185,cell surface receptor signaling pathway,0.00709318156360945

GO:0045780,0.00529028543826468,9.64112388250319,0.360468140442133,3,21,positive regulation of bone resorption,0.00714249087872688

GO:1903169,0.00532317716433418,3.5220716490004,2.11131339401821,7,123,regulation of calcium ion transmembrane transport,0.00714249087872688

GO:0044093,0.00540760687014568,1.72053954629062,19.1563068920676,31,1116,positive regulation of molecular function,0.00721014249352757

GO:0016310,0.00570347817601783,1.68561751542277,20.8384915474642,33,1214,phosphorylation,0.00731210868304489

GO:0071900,0.00576369868896687,2.71580635753864,3.87932379713914,10,226,regulation of protein serine/threonine kinase activity,0.00731210868304489

GO:0033700,0.00582316040156942,23.0702290076336,0.120156046814044,2,7,phospholipid efflux,0.00731210868304489

GO:0038007,0.00582316040156942,23.0702290076336,0.120156046814044,2,7,netrin-activated signaling pathway,0.00731210868304489

GO:0014012,0.00582316040156942,23.0702290076336,0.120156046814044,2,7,peripheral nervous system axon regeneration,0.00731210868304489

GO:0072537,0.00582316040156942,23.0702290076336,0.120156046814044,2,7,fibroblast activation,0.00731210868304489

GO:0099550,0.00582316040156942,23.0702290076336,0.120156046814044,2,7,trans-synaptic signaling, modulating synaptic transmission,0.00731210868304489

GO:0097384,0.00582316040156942,23.0702290076336,0.120156046814044,2,7,cellular lipid biosynthetic process,0.00731210868304489

GO:0032308,0.00582316040156942,23.0702290076336,0.120156046814044,2,7,positive regulation of prostaglandin secretion,0.00731210868304489

GO:0034109,0.00582899229921975,4.68738323577033,1.15006501950585,5,67,homotypic cell-cell adhesion,0.00731210868304489

GO:0014902,0.00587955170553299,3.92657434021427,1.63068920676203,6,95,myotube differentiation,0.00733214683278231

GO:0032288,0.00604878026655701,9.13309134906231,0.377633289986996,3,22,myelin assembly,0.00745547335180282

GO:2000406,0.00604878026655701,9.13309134906231,0.377633289986996,3,22,positive regulation of T cell migration,0.00745547335180282

GO:0008361,0.00620864733510466,3.09706125827815,2.72925877763329,8,159,regulation of cell size,0.00759092634789705

GO:0051049,0.00625141724656526,1.66151733849674,21.7825747724317,34,1269,regulation of transport,0.00759092634789705

GO:0051641,0.00626609486265086,1.46422172523845,48.4400520156047,65,2822,cellular localization,0.00759092634789705

GO:0042981,0.00655529028991108,1.72533580751956,17.8174252275683,29,1038,regulation of apoptotic process,0.00789614512193834

GO:0007265,0.00665481008127156,2.16617079565205,7.27802340702211,15,424,Ras protein signal transduction,0.00797073297869814

GO:0043496,0.00686894539645687,8.67586206896552,0.39479843953186,3,23,regulation of protein homodimerization activity,0.00809009124471587

GO:0045742,0.00686894539645687,8.67586206896552,0.39479843953186,3,23,positive regulation of epidermal growth factor receptor signaling pathway,0.00809009124471587

GO:0042398,0.00686894539645687,8.67586206896552,0.39479843953186,3,23,cellular modified amino acid biosynthetic process,0.00809009124471587

GO:0048878,0.00710989563471371,1.86105371900826,12.4618985695709,22,726,chemical homeostasis,0.008327612566626

GO:0007584,0.00716089335354536,3.32007845370283,2.23146944083225,7,130,response to nutrient,0.00834126038984405

GO:0061448,0.00751552235397097,2.60559930008749,4.03381014304291,10,235,connective tissue development,0.0085544970600059

GO:0010035,0.00762785728527385,2.19531914893617,6.69440832249675,14,390,response to inorganic substance,0.0085544970600059

GO:0046642,0.00767664123252662,19.2239185750636,0.137321196358908,2,8,negative regulation of alpha-beta T cell proliferation,0.0085544970600059

GO:0032689,0.00767664123252662,19.2239185750636,0.137321196358908,2,8,negative regulation of type II interferon production,0.0085544970600059

GO:0086103,0.00767664123252662,19.2239185750636,0.137321196358908,2,8,G protein-coupled receptor signaling pathway involved in heart process,0.0085544970600059

GO:0032305,0.00767664123252662,19.2239185750636,0.137321196358908,2,8,positive regulation of icosanoid secretion,0.0085544970600059

GO:0032306,0.00767664123252662,19.2239185750636,0.137321196358908,2,8,regulation of prostaglandin secretion,0.0085544970600059

GO:0090077,0.00775195483351294,8.26217843459223,0.411963589076723,3,24,foam cell differentiation,0.0085544970600059

GO:0010742,0.00775195483351294,8.26217843459223,0.411963589076723,3,24,macrophage derived foam cell differentiation,0.0085544970600059

GO:1901186,0.00775195483351294,8.26217843459223,0.411963589076723,3,24,positive regulation of ERBB signaling pathway,0.0085544970600059

GO:0023061,0.00778782043670348,2.26448053938094,6.02496749024707,13,351,signal release,0.0085544970600059

GO:0010876,0.00830210202760043,2.43658736669401,4.73758127438231,11,276,lipid localization,0.00907240015387263

GO:0007169,0.00858945108734285,1.95627657194003,9.66397919375813,18,563,transmembrane receptor protein tyrosine kinase signaling pathway,0.009267132889168

GO:0090200,0.00869886530634166,7.88610240334378,0.429128738621586,3,25,positive regulation of release of cytochrome c from mitochondria,0.009267132889168

GO:0010800,0.00869886530634166,7.88610240334378,0.429128738621586,3,25,positive regulation of peptidyl-threonine phosphorylation,0.009267132889168

GO:0140353,0.00869886530634166,7.88610240334378,0.429128738621586,3,25,lipid export from cell,0.009267132889168

GO:0032309,0.00869886530634166,7.88610240334378,0.429128738621586,3,25,icosanoid secretion,0.009267132889168

GO:0043067,0.00876774694027424,1.68413326735939,18.2122236671001,29,1061,regulation of programmed cell death,0.00928306431830251

GO:0022603,0.00880139588669247,1.79774779189946,13.4746423927178,23,785,regulation of anatomical structure morphogenesis,0.00928306431830251

GO:0034329,0.00910461694570406,2.21748368843467,6.14512353706112,13,358,cell junction assembly,0.0095407533716587

GO:0034103,0.00913572138889961,5.26993006993007,0.823927178153446,4,48,regulation of tissue remodeling,0.0095407533716587

GO:0007275,0.00925943910229048,1.38002158833226,72.28244473342,90,4211,multicellular organism development,0.00962255436120383

GO:0090504,0.00971062090760996,7.54272863568216,0.44629388816645,3,26,epiboly,0.00985174236745276

GO:2000114,0.00971062090760996,7.54272863568216,0.44629388816645,3,26,regulation of establishment of cell polarity,0.00985174236745276

GO:0048881,0.00975880140172208,16.4765539803708,0.154486345903771,2,9,mechanosensory lateral line system development,0.00985174236745276

GO:0048915,0.00975880140172208,16.4765539803708,0.154486345903771,2,9,posterior lateral line system development,0.00985174236745276

GO:0030913,0.00975880140172208,16.4765539803708,0.154486345903771,2,9,paranodal junction assembly,0.00985174236745276

GO:0035313,0.00975880140172208,16.4765539803708,0.154486345903771,2,9,wound healing, spreading of epidermal cells,0.00985174236745276

GO:0048513,0.00986932598603582,1.45463635596998,42.2777633289987,57,2463,animal organ development,0.00991610004284168

GO:0033993,0.00991933472281277,1.88933022873579,10.556566970091,19,615,response to lipid,0.00991933472281277

Table S5. Supplementary data to accompany Figure 3D. Top gene ontology (Functional Enrichment Analysis) for each compartment (SL).

GOBPID,Pvalue,OddsRatio,ExpCount,Count,Size,Term,FDR

GO:0009653,4.67719085470563e-19,2.73801153537818,65.5461638491548,138,2372,anatomical structure morphogenesis,2.30117790051517e-16

GO:0032502,1.99048222187005e-16,2.25046099046237,146.649869960988,228,5307,developmental process,4.89658626580033e-14

GO:0048856,7.25808650178254e-16,2.22281646028244,138.000650195059,217,4994,anatomical structure development,1.19032618629234e-13

GO:0009888,2.80385605570766e-15,2.75460122699387,43.7711313394018,99,1584,tissue development,3.44874294852042e-13

GO:0048646,8.32904354431508e-15,3.17111371063577,27.3845903771131,73,991,anatomical structure formation involved in morphogenesis,8.19577884760604e-13

GO:0048513,5.11430137090308e-14,2.35662735357465,68.0607932379714,129,2463,animal organ development,4.19372712414052e-12

GO:0030154,1.85570070798746e-13,2.1749811097222,93.4834200260078,159,3383,cell differentiation,1.14125593541229e-11

GO:0048869,1.85570070798746e-13,2.1749811097222,93.4834200260078,159,3383,cellular developmental process,1.14125593541229e-11

GO:0048731,1.2812683267212e-12,2.1302616837388,90.5542912873862,153,3277,system development,7.00426685274256e-11

GO:0007275,5.7803675496819e-12,2.01096794532807,116.363784135241,181,4211,multicellular organism development,2.84394083444349e-10

GO:0032501,1.9682407645633e-11,1.92955325060498,161.240247074122,228,5835,multicellular organismal process,8.80340414695586e-10

GO:0061061,1.48533033714846e-10,3.40187344926712,14.424577373212,43,522,muscle structure development,6.08985438230867e-09

GO:0009887,7.32263049101983e-10,2.67167523903091,25.1462938881665,59,910,animal organ morphogenesis,2.77133400121674e-08

GO:0035295,1.36096267061865e-09,2.64559232790637,24.8975942782835,58,901,tube development,4.19351267809935e-08

GO:0072359,1.3612491186026e-09,2.53990402064766,28.24122236671,63,1022,circulatory system development,4.19351267809935e-08

GO:0031032,1.36374396035751e-09,5.05479159244745,5.49902470741222,24,199,actomyosin structure organization,4.19351267809935e-08

GO:0048729,1.86983105902158e-09,3.05930600586691,16.2483745123537,44,588,tissue morphogenesis,5.41151106493305e-08

GO:0007167,2.43275679668544e-09,2.69614266712646,22.6592977893368,54,820,enzyme-linked receptor protein signaling pathway,6.64953524427353e-08

GO:0035239,5.13714151566413e-09,2.71184611477908,21.194733420026,51,767,tube morphogenesis,1.33024927668777e-07

GO:0030239,1.20820585120981e-08,7.00951483194391,2.73569570871261,16,99,myofibril assembly,2.97218639397614e-07

GO:0055002,1.87895853788192e-08,6.76363222835049,2.8185955786736,16,102,striated muscle cell development,4.40213143160906e-07

GO:0006936,3.34785242712424e-08,3.92873577386088,7.46098829648895,26,270,muscle contraction,7.48701542793239e-07

GO:0010927,4.74740347201049e-08,5.46744471744472,3.81339401820546,18,138,cellular component assembly involved in morphogenesis,1.01553152531703e-06

GO:0051146,6.35770416719806e-08,3.91489361702128,7.18465539661899,25,260,striated muscle cell differentiation,1.3033293542756e-06

GO:0003008,6.64250113926351e-08,2.16948884317139,36.3654096228869,70,1316,system process,1.30724422420706e-06

GO:0003012,7.85474294511523e-08,3.44827698203904,9.39531859557867,29,340,muscle system process,1.48635904961411e-06

GO:0007517,1.0693900759259e-07,3.68295739348371,7.90312093628088,26,286,muscle organ development,1.94866636057609e-06

GO:0001525,1.19866967819304e-07,3.03159211761026,12.4626137841352,34,451,angiogenesis,2.10623386311063e-06

GO:0045214,1.43276675947123e-07,7.458622283865,2.1001300390117,13,76,sarcomere organization,2.43076291606843e-06

GO:0055001,1.70714315768139e-07,4.48140574111944,5.05689206762029,20,183,muscle cell development,2.79845322398918e-06

GO:0048514,1.76325304763546e-07,2.78677855449455,15.1154096228869,38,547,blood vessel morphogenesis,2.79845322398918e-06

GO:0001568,1.86953425560177e-07,2.66537814670139,17.0497399219766,41,617,blood vessel development,2.87440891798772e-06

GO:0001944,2.37745902531483e-07,2.60552798575208,17.8511053315995,42,646,vasculature development,3.54457527410575e-06

GO:0007169,3.64980696536642e-07,2.69885566629753,15.5575422626788,38,563,transmembrane receptor protein tyrosine kinase signaling pathway,5.28148537341259e-06

GO:0042692,3.76871154025253e-07,3.33664125999123,8.98081924577373,27,325,muscle cell differentiation,5.29773165086928e-06

GO:0060537,5.22855728326482e-07,3.19498405221826,9.69928478543563,28,351,muscle tissue development,7.14569495379526e-06

GO:0071363,8.07749765596008e-07,2.68102271380808,14.7838101430429,36,535,cellular response to growth factor stimulus,1.07408887749523e-05

GO:0030334,1.08062438734779e-06,2.44481589029062,18.9011703511053,42,684,regulation of cell migration,1.39912420677662e-05

GO:0050793,1.17634785535489e-06,1.85549202673036,53.4151495448635,88,1933,regulation of developmental process,1.48400806367848e-05

GO:0070848,1.45425108374644e-06,2.57028615473552,15.8062418725618,37,572,response to growth factor,1.78872883300812e-05

GO:0007166,1.64788149774473e-06,1.79713392338796,60.3787386215865,96,2185,cell surface receptor signaling pathway,1.97745779729368e-05

GO:2000145,3.12224920834276e-06,2.33441377780718,19.7025357607282,42,713,regulation of cell motility,3.65749192977295e-05

GO:0071495,3.42173978319784e-06,2.0554135205775,30.5347854356307,57,1105,cellular response to endogenous stimulus,3.91510691472869e-05

GO:0007507,4.06918468870843e-06,2.51630570077172,15.1983094928479,35,550,heart development,4.55008833373761e-05

GO:0023057,4.75411130534848e-06,2.04320317864687,30.1202860858257,56,1090,negative regulation of signaling,5.0848320918075e-05

GO:0010648,4.75411130534848e-06,2.04320317864687,30.1202860858257,56,1090,negative regulation of cell communication,5.0848320918075e-05

GO:0048585,5.25922254164285e-06,1.9762970135083,33.9889466840052,61,1230,negative regulation of response to stimulus,5.50539891593251e-05

GO:0090287,5.63160159101474e-06,3.63669950738916,5.77535760728219,19,209,regulation of cellular response to growth factor stimulus,5.77239163079011e-05

GO:0033275,6.4418774432698e-06,12.5053827751196,0.746098829648895,7,27,actin-myosin filament sliding,6.46817082058927e-05

GO:0048598,8.1881890378427e-06,2.50096383522608,14.369310793238,33,520,embryonic morphogenesis,8.05717801323722e-05

GO:0048522,8.68115133509168e-06,1.57480431319125,115.534785435631,156,4181,positive regulation of cellular process,8.37475775855903e-05

GO:0070371,9.0724821429395e-06,3.50581380810682,5.96879063719116,19,216,ERK1 and ERK2 cascade,8.54595482386602e-05

GO:0048518,9.20600824522153e-06,1.5584137991573,127.638166449935,169,4619,positive regulation of biological process,8.54595482386602e-05

GO:0009968,1.07235934218656e-05,2.03233914209115,27.9648894668401,52,1012,negative regulation of signal transduction,9.77038511769978e-05

GO:0040012,1.12144951765614e-05,2.20341966584054,20.7526007802341,42,751,regulation of locomotion,0.000100318756852149

GO:0016477,1.17573891869978e-05,1.95031197816274,32.5796488946684,58,1179,cell migration,0.000103297062142909

GO:0051239,1.248102920837e-05,1.71707902554266,59.1076072821847,91,2139,regulation of multicellular organismal process,0.000107730988956457

GO:0001667,1.40361338796736e-05,2.75950746431296,10.2519505851756,26,371,ameboidal-type cell migration,0.000119065135668955

GO:0070372,1.70544866737181e-05,3.47386167598934,5.6924577373212,18,206,regulation of ERK1 and ERK2 cascade,0.000142217075313039

GO:0050789,1.87517341723131e-05,1.53596821096543,245.632314694408,287,8889,regulation of biological process,0.000153764220212967

GO:0060070,2.07855400418738e-05,3.28589021815623,6.32802340702211,19,229,canonical Wnt signaling pathway,0.000164431583196652

GO:0072338,2.09561194396757e-05,Inf,0.0828998699609883,3,3,lactam metabolic process,0.000164431583196652

GO:0008543,2.10552637020104e-05,5.98192771084337,1.93433029908973,10,70,fibroblast growth factor receptor signaling pathway,0.000164431583196652

GO:0048468,2.3418866354541e-05,1.67227153872594,62.5894018205462,94,2265,cell development,0.000179285409964057

GO:0032989,2.36860805846824e-05,2.12768443287181,21.415799739922,42,775,cellular component morphogenesis,0.000179285409964057

GO:0030036,2.54448115644908e-05,2.26121399716699,17.2431729518856,36,624,actin cytoskeleton organization,0.00018967950438984

GO:0044344,2.97913836190565e-05,5.13388543823326,2.43172951885566,11,88,cellular response to fibroblast growth factor stimulus,0.000216266965302109

GO:0065007,3.00342677098553e-05,1.52643969156909,254.088101430429,294,9195,biological regulation,0.000216266965302109

GO:0061138,3.03301231826128e-05,3.8164720027482,4.33842652795839,15,157,morphogenesis of a branching epithelium,0.000216266965302109

GO:0030029,3.08050121063226e-05,2.16432716284345,19.5091027308192,39,706,actin filament-based process,0.000216515227947296

GO:2000026,3.14302435218521e-05,1.9196642747178,30.6176853055917,54,1108,regulation of multicellular organismal development,0.000217798307221848

GO:0051094,3.20956422898339e-05,1.96782578152441,27.6056566970091,50,999,positive regulation of developmental process,0.000219320222313865

GO:0048880,3.2947361063383e-05,2.67050438596491,10.1414174252276,25,367,sensory system development,0.000222056186892937

GO:0007267,3.57883335501833e-05,1.88740604387176,32.3033159947984,56,1169,cell-cell signaling,0.000237944055495813

GO:0050678,3.98613398767219e-05,3.01500947697352,7.21228868660598,20,261,regulation of epithelial cell proliferation,0.000261490389591296

GO:0009719,4.32362263394036e-05,1.83427998663548,35.646944083225,60,1290,response to endogenous stimulus,0.000279662932144169

GO:0022618,4.376838572175e-05,3.6854156296665,4.47659297789337,15,162,protein-RNA complex assembly,0.000279662932144169

GO:0030049,4.87083940128761e-05,11.256877276724,0.690832249674902,6,25,muscle filament sliding,0.00030420837694833

GO:0002455,4.88464670303213e-05,8.61920475169114,0.99479843953186,7,36,humoral immune response mediated by circulating immunoglobulin,0.00030420837694833

GO:0071774,5.01488851334096e-05,4.81922351832214,2.56989596879064,11,93,response to fibroblast growth factor,0.000308415643570469

GO:0048519,5.1067658041371e-05,1.51724105838227,110.864759427828,147,4012,negative regulation of biological process,0.000310188737732772

GO:0060485,5.20820560163015e-05,3.16645595286372,6.18985695708713,18,224,mesenchyme development,0.000312492336097809

GO:0150063,5.33061366499104e-05,2.64426546925737,9.80981794538362,24,355,visual system development,0.000315983364238023

GO:0040011,5.64450082534287e-05,1.98989661654135,24.4554616384915,45,885,locomotion,0.00033060647691294

GO:0050794,5.7949736513541e-05,1.48318955531025,232.810468140442,272,8425,regulation of cellular process,0.000335426710172496

GO:0072006,5.870344622772e-05,4.03638684298627,3.5646944083225,13,129,nephron development,0.000335838320279514

GO:0007492,6.18372216108386e-05,5.86101398601399,1.76853055916775,9,64,endoderm development,0.000347662753319342

GO:0071826,6.21835818945164e-05,3.56298138639281,4.61475942782835,15,167,protein-RNA complex organization,0.000347662753319342

GO:0030178,6.88919146297046e-05,3.96743870330755,3.61996098829649,13,131,negative regulation of Wnt signaling pathway,0.000380840696604659

GO:0030111,7.86372638609946e-05,2.85817050646447,7.57152145643693,20,274,regulation of Wnt signaling pathway,0.000425835468265836

GO:0072001,7.87622512442909e-05,2.77479747974797,8.17945383615085,21,296,renal system development,0.000425835468265836

GO:0003013,8.1325456174925e-05,2.51239669421488,10.7217165149545,25,388,circulatory system process,0.000431806237646358

GO:0018108,8.20580517571832e-05,3.04643045764541,6.4109232769831,18,232,peptidyl-tyrosine phosphorylation,0.000431806237646358

GO:0048870,8.24995657291822e-05,1.79559869775034,35.674577373212,59,1291,cell motility,0.000431806237646358

GO:0030324,8.7042825212456e-05,3.86829013881088,3.70286085825748,13,134,lung development,0.000448450510797255

GO:0009790,8.75025386921473e-05,1.9342550731047,25.6713263979194,46,929,embryo development,0.000448450510797255

GO:0018212,9.15989611773074e-05,3.01781326781327,6.46618985695709,18,234,peptidyl-tyrosine modification,0.00046232758946652

GO:0001763,9.29704336250223e-05,3.42628897807965,4.78055916775032,15,173,morphogenesis of a branching structure,0.00046232758946652

GO:0060828,9.33789522718857e-05,3.26617214371555,5.33322496749025,16,193,regulation of canonical Wnt signaling pathway,0.00046232758946652

GO:0001654,9.39690222492927e-05,2.6000355366027,9.53348504551365,23,345,eye development,0.00046232758946652

GO:0016055,9.96613731070459e-05,2.47741168478261,10.8598829648895,25,393,Wnt signaling pathway,0.000485479164046204

GO:0001822,0.000100793270680007,2.80204013537347,7.70968790637191,20,279,kidney development,0.000486039793499739

GO:0048523,0.000101752233192019,1.5006754531585,104.094603381014,138,3767,negative regulation of cellular process,0.000486039793499739

GO:0031327,0.000103971045282854,1.71941645228563,42.417100130039,67,1535,negative regulation of cellular biosynthetic process,0.000490288637829897

GO:0030335,0.000104634770268576,2.42083525139379,11.5507152145644,26,418,positive regulation of cell migration,0.000490288637829897

GO:0060538,0.000109194570615981,3.77393908549953,3.78576072821847,13,137,skeletal muscle organ development,0.000506827629651535

GO:0090130,0.000110611957188459,2.86930418719212,7.15702210663199,19,259,tissue migration,0.000508608251745063

GO:0050673,0.000120233924508455,2.68521649700181,8.42815344603381,21,305,epithelial cell proliferation,0.000543919375338373

GO:0009890,0.000121250650788082,1.70908275423536,42.638166449935,67,1543,negative regulation of biosynthetic process,0.000543919375338373

GO:0198738,0.000121607990421181,2.44336461126005,10.9980494148244,25,398,cell-cell signaling by wnt,0.000543919375338373

GO:0030323,0.000126530808995426,3.71353444290337,3.84102730819246,13,139,respiratory tube development,0.00056083926149324

GO:0090090,0.000128191296921079,4.292506826297,2.8462288686606,11,103,negative regulation of canonical Wnt signaling pathway,0.000563126054331885

GO:0014706,0.000135110202388499,3.02791461412151,6.07932379713914,17,220,striated muscle tissue development,0.000588267429868507

GO:0008015,0.000146651712813379,2.46145200379333,10.4730169050715,24,379,blood circulation,0.000628224263086879

GO:0022008,0.00014684103710364,1.70902340408988,41.2841352405722,65,1494,neurogenesis,0.000628224263086879

GO:0007369,0.000153425826101328,3.43135210288495,4.44895968790637,14,161,gastrulation,0.000650653307280906

GO:0051253,0.000155246704289491,1.86867758893482,27.0806241872562,47,980,negative regulation of RNA metabolic process,0.000650653307280906

GO:0042127,0.000156559983060441,1.67985880609807,44.6553966189857,69,1616,regulation of cell population proliferation,0.000650653307280906

GO:0010558,0.000158494819711224,1.70374893386059,41.3946684005202,65,1498,negative regulation of macromolecule biosynthetic process,0.000650653307280906

GO:2000147,0.000158695928605099,2.35297922224917,11.8546814044213,26,429,positive regulation of cell motility,0.000650653307280906

GO:0050680,0.000165935273566406,4.15611492499364,2.92912873862159,11,106,negative regulation of epithelial cell proliferation,0.000674712021443567

GO:0002009,0.000168061096055532,2.30257009345794,12.5731469440832,27,455,morphogenesis of an epithelium,0.0006777545840928

GO:0040036,0.000173096293720027,11.8571428571429,0.552665799739922,5,20,regulation of fibroblast growth factor receptor signaling pathway,0.000692385174880109

GO:0070374,0.000178652692198884,3.81632346954081,3.45416124837451,12,125,positive regulation of ERK1 and ERK2 cascade,0.000708847778724603

GO:0006816,0.000187052007867797,2.5289600150768,9.34005201560468,22,338,calcium ion transport,0.000736236702967649

GO:0007411,0.000190416046523727,2.83142826621087,6.85305591677503,18,248,axon guidance,0.000737674762910817

GO:0097485,0.000190416046523727,2.83142826621087,6.85305591677503,18,248,neuron projection guidance,0.000737674762910817

GO:0048754,0.000192629080697188,3.78259207340385,3.48179453836151,12,126,branching morphogenesis of an epithelial tube,0.000740418028929815

GO:0044057,0.000197649429777884,2.31750525457899,12.0204811443433,26,435,regulation of system process,0.000751759218978938

GO:0043542,0.00019863556599037,3.3395530204154,4.55949284785436,14,165,endothelial cell migration,0.000751759218978938

GO:0010511,0.00020102900924391,53.1504739336493,0.13816644993498,3,5,regulation of phosphatidylinositol biosynthetic process,0.000755009714106899

GO:0048738,0.000210608282619618,3.02389943547664,5.72009102730819,16,207,cardiac muscle tissue development,0.00078499450794585

GO:0045595,0.000224148732935075,1.77188940092166,32.2204161248374,53,1166,regulation of cell differentiation,0.00082918177897787

GO:0040017,0.000244881314591297,2.28305704019084,12.1862808842653,26,441,positive regulation of locomotion,0.000894983535142472

GO:0010605,0.00024557475049641,1.60617618805447,52.8901170351105,78,1914,negative regulation of macromolecule metabolic process,0.000894983535142472

GO:0010631,0.000255608921587415,2.75831841086078,7.01885565669701,18,254,epithelial cell migration,0.000924702863389764

GO:0090132,0.000281291374491708,2.73476761712056,7.074122236671,18,256,epithelium migration,0.00101018508211621

GO:0007519,0.000297632331052418,3.59200968523002,3.64759427828348,12,132,skeletal muscle tissue development,0.0010575913239965

GO:0001501,0.000298791044787629,2.2918765743073,11.6612483745124,25,422,skeletal system development,0.0010575913239965

GO:0048762,0.000310767092039278,3.03720197314333,5.33322496749025,15,193,mesenchymal cell differentiation,0.00109212435202375

GO:0006959,0.000316465598610149,4.16615298402914,2.65279583875163,10,96,humoral immune response,0.00110426293983116

GO:0006956,0.000323894537524458,6.09160928929863,1.32639791937581,7,48,complement activation,0.00112222614409883

GO:0001732,0.000337133342195895,15.778305621536,0.359232769830949,4,13,formation of cytoplasmic translation initiation complex,0.00115381964401857

GO:0010562,0.000340048472322546,2.07243942501322,15.9720416124837,31,578,positive regulation of phosphorus metabolic process,0.00115381964401857

GO:0045937,0.000340048472322546,2.07243942501322,15.9720416124837,31,578,positive regulation of phosphate metabolic process,0.00115381964401857

GO:0090329,0.000353374596310559,7.37025759196774,0.967165149544863,6,35,regulation of DNA-templated DNA replication,0.00119082398208764

GO:0140694,0.000356354042956507,2.40445155311071,9.78218465539662,22,354,non-membrane-bounded organelle assembly,0.00119269516418096

GO:0045892,0.000364325251192248,1.84035361572381,25.0081274382315,43,905,negative regulation of DNA-templated transcription,0.00120437205936216

GO:0048736,0.000364738692774311,3.13001163651751,4.83582574772432,14,175,appendage development,0.00120437205936216

GO:0007015,0.000391294512912408,2.28939462367703,11.1914824447334,24,405,actin filament organization,0.00121095029184866

GO:0034285,0.000393804972958912,35.4312796208531,0.165799739921977,3,6,response to disaccharide,0.00121095029184866

GO:2000589,0.000393804972958912,35.4312796208531,0.165799739921977,3,6,regulation of metanephric mesenchymal cell migration,0.00121095029184866

GO:0009744,0.000393804972958912,35.4312796208531,0.165799739921977,3,6,response to sucrose,0.00121095029184866

GO:0035787,0.000393804972958912,35.4312796208531,0.165799739921977,3,6,cell migration involved in kidney development,0.00121095029184866

GO:0035788,0.000393804972958912,35.4312796208531,0.165799739921977,3,6,cell migration involved in metanephros development,0.00121095029184866

GO:0035789,0.000393804972958912,35.4312796208531,0.165799739921977,3,6,metanephric mesenchymal cell migration,0.00121095029184866

GO:0019375,0.000393804972958912,35.4312796208531,0.165799739921977,3,6,galactolipid biosynthetic process,0.00121095029184866

GO:0006682,0.000393804972958912,35.4312796208531,0.165799739921977,3,6,galactosylceramide biosynthetic process,0.00121095029184866

GO:0021631,0.000393804972958912,35.4312796208531,0.165799739921977,3,6,optic nerve morphogenesis,0.00121095029184866

GO:0046476,0.000393804972958912,35.4312796208531,0.165799739921977,3,6,glycosylceramide biosynthetic process,0.00121095029184866

GO:0045934,0.000400701704978978,1.76183897954968,29.8163198959688,49,1079,negative regulation of nucleobase-containing compound metabolic process,0.00122450458912831

GO:1902679,0.000408787647254594,1.82909110672295,25.1462938881665,43,910,negative regulation of RNA biosynthetic process,0.00124150322499543

GO:2000351,0.000414280222166608,7.12410501193317,0.99479843953186,6,36,regulation of endothelial cell apoptotic process,0.00125046545586485

GO:0002040,0.000425082757553253,3.68702875976342,3.26072821846554,11,118,sprouting angiogenesis,0.00127524827265976

GO:0051241,0.000447267534457767,1.91043542563835,20.6697009102731,37,748,negative regulation of multicellular organismal process,0.00133367046638316

GO:0022607,0.000464883447848614,1.50649636233526,70.492522756827,97,2551,cellular component assembly,0.00137784732735854

GO:0045597,0.000469037886200349,1.98308535236755,17.7405721716515,33,642,positive regulation of cell differentiation,0.00138183616772797

GO:0060541,0.000476242984442187,3.2005087112648,4.39369310793238,13,159,respiratory system development,0.00139471159729498

GO:0045445,0.000498341991819853,4.8436369548429,1.85143042912874,8,67,myoblast differentiation,0.00145079443772407

GO:2000027,0.000516129441057992,3.89287585123101,2.8185955786736,10,102,regulation of animal organ morphogenesis,0.00149373932353254

GO:0051093,0.000536958670691717,1.9662509140464,17.8787386215865,33,647,negative regulation of developmental process,0.00154493371918319

GO:0043010,0.000552053849860144,2.49816390506046,8.12418725617685,19,294,camera-type eye development,0.0015791307798325

GO:0010594,0.000565062096632173,3.55320537929234,3.37126137841352,11,122,regulation of endothelial cell migration,0.00160191090173932

GO:0022603,0.000566529465249273,1.86760478606934,21.6921326397919,38,785,regulation of anatomical structure morphogenesis,0.00160191090173932

GO:0051216,0.000572315760388367,2.98023294317511,5.05689206762029,14,183,cartilage development,0.00160774210253399

GO:0031324,0.00057512725619102,1.55777252502781,53.5256827048114,77,1937,negative regulation of cellular metabolic process,0.00160774210253399

GO:0045765,0.000617579913493301,2.72048715228122,6.30039011703511,16,228,regulation of angiogenesis,0.00171097215816988

GO:0048705,0.000619010252346013,2.82799131656238,5.6924577373212,15,206,skeletal system morphogenesis,0.00171097215816988

GO:0010464,0.000640636316786877,8.46598639455782,0.718465539661899,5,26,regulation of mesenchymal cell proliferation,0.00174711804833262

GO:0051240,0.000641812886237526,1.72050978965873,30.4518855656697,49,1102,positive regulation of multicellular organismal process,0.00174711804833262

GO:0065003,0.000642740582821554,1.6352915220923,39.4050715214564,60,1426,protein-containing complex assembly,0.00174711804833262

GO:0034446,0.000649665352060364,3.76918199112238,2.90149544863459,10,105,substrate adhesion-dependent cell spreading,0.00175623820447087

GO:1901342,0.000711255753334928,2.68198100869961,6.3832899869961,16,231,regulation of vasculature development,0.00190197811035282

GO:0008285,0.000711308886798616,1.97483190262312,16.6905071521456,31,604,negative regulation of cell population proliferation,0.00190197811035282

GO:0048699,0.000716860681514424,1.65380212274676,36.2825097529259,56,1313,generation of neurons,0.00190646192056809

GO:0001656,0.000738562197164179,4.534886376613,1.96196358907672,8,71,metanephros development,0.00194212933443232

GO:0048565,0.000753621575684145,3.69097006583033,2.95676202860858,10,107,digestive tract development,0.00194212933443232

GO:0007495,0.00076185154785658,Inf,0.0552665799739922,2,2,visceral mesoderm-endoderm interaction involved in midgut development,0.00194212933443232

GO:0061837,0.00076185154785658,Inf,0.0552665799739922,2,2,neuropeptide processing,0.00194212933443232

GO:0031443,0.00076185154785658,Inf,0.0552665799739922,2,2,fast-twitch skeletal muscle fiber contraction,0.00194212933443232

GO:0031446,0.00076185154785658,Inf,0.0552665799739922,2,2,regulation of fast-twitch skeletal muscle fiber contraction,0.00194212933443232

GO:0031448,0.00076185154785658,Inf,0.0552665799739922,2,2,positive regulation of fast-twitch skeletal muscle fiber contraction,0.00194212933443232

GO:0046449,0.00076185154785658,Inf,0.0552665799739922,2,2,creatinine metabolic process,0.00194212933443232

GO:0010646,0.000790284934023544,1.47313790719779,73.3663849154746,99,2655,regulation of cell communication,0.00198863062037457

GO:1904018,0.000792218702425643,3.39890471430951,3.5094278283485,11,127,positive regulation of vasculature development,0.00198863062037457

GO:0045766,0.000792218702425643,3.39890471430951,3.5094278283485,11,127,positive regulation of angiogenesis,0.00198863062037457

GO:0090092,0.000801330548158859,3.01284058878797,4.64239271781535,13,168,regulation of transmembrane receptor protein serine/threonine kinase signaling pathway,0.00200129253651857

GO:0008283,0.000835437502148685,1.47475175262161,71.7083875162549,97,2595,cell population proliferation,0.00207593561139976

GO:0001932,0.000841876152155933,1.83967463525857,21.388166449935,37,774,regulation of protein phosphorylation,0.00208142244653628

GO:0072577,0.00085130297232756,6.10433003750426,1.13296488946684,6,41,endothelial cell apoptotic process,0.00209239854175782

GO:0061448,0.000854821355474233,2.63228053722745,6.49382314694408,16,235,connective tissue development,0.00209239854175782

GO:0048732,0.000859541761364962,2.40028075372903,8.42815344603381,19,305,gland development,0.00209353735936417

GO:0045834,0.000870903283942963,3.61591821832786,3.01202860858257,10,109,positive regulation of lipid metabolic process,0.0021107606684726

GO:0051172,0.000893596750706397,1.57745463500113,44.9593628088427,66,1627,negative regulation of nitrogen compound metabolic process,0.0021469354359395

GO:0060173,0.000894556431641459,2.97405849978356,4.69765929778934,13,170,limb development,0.0021469354359395

GO:0006958,0.000914664604751026,7.7287784679089,0.773732119635891,5,28,complement activation, classical pathway,0.00217398543737925

GO:0003413,0.000914664604751026,7.7287784679089,0.773732119635891,5,28,chondrocyte differentiation involved in endochondral bone morphogenesis,0.00217398543737925

GO:0016043,0.000932005527909375,1.37715561936667,142.256176853056,173,5148,cellular component organization,0.00220455153717025

GO:0032835,0.00093985390544645,4.9921052631579,1.57509752925878,7,57,glomerulus development,0.00220890076019714

GO:0035051,0.000942823495206095,3.11969681019055,4.14499349804941,12,150,cardiocyte differentiation,0.00220890076019714

GO:0097435,0.000950351065277247,1.89532393145938,18.4866710013004,33,669,supramolecular fiber organization,0.00221598447448533

GO:0050878,0.000958217081892792,2.44224196855776,7.84785435630689,18,284,regulation of body fluid levels,0.00222378681269459

GO:0042325,0.001012314102442,1.79007438054044,23.1566970091027,39,838,regulation of phosphorylation,0.00233830299718998

GO:0006688,0.00102793700694255,10.9205189110177,0.469765929778934,4,17,glycosphingolipid biosynthetic process,0.00236026539529468

GO:0001708,0.00104390007274152,4.89389248522375,1.60273081924577,7,58,cell fate specification,0.00236026539529468

GO:0090269,0.00105794122967812,21.2559241706161,0.221066319895969,3,8,fibroblast growth factor production,0.00236026539529468

GO:2000020,0.00105794122967812,21.2559241706161,0.221066319895969,3,8,positive regulation of male gonad development,0.00236026539529468

GO:2000018,0.00105794122967812,21.2559241706161,0.221066319895969,3,8,regulation of male gonad development,0.00236026539529468

GO:0048541,0.00105794122967812,21.2559241706161,0.221066319895969,3,8,Peyer's patch development,0.00236026539529468

GO:0048537,0.00105794122967812,21.2559241706161,0.221066319895969,3,8,mucosa-associated lymphoid tissue development,0.00236026539529468

GO:0010632,0.00106698109941231,2.78039010095307,5.38849154746424,14,195,regulation of epithelial cell migration,0.00236026539529468

GO:0014033,0.00106865013068536,3.83009958791209,2.56989596879064,9,93,neural crest cell differentiation,0.00236026539529468

GO:0007399,0.0010699712254955,1.50811899619683,57.2838101430429,80,2073,nervous system development,0.00236026539529468

GO:0055123,0.00107459237509351,3.50885896527286,3.09492847854356,10,112,digestive system development,0.00236026539529468

GO:0060351,0.00110185209497938,5.77359220795975,1.18823146944083,6,43,cartilage development involved in endochondral bone morphogenesis,0.00240938324768825

GO:0010811,0.00115056455036829,3.4745584278863,3.12256176853056,10,113,positive regulation of cell-substrate adhesion,0.00250030758520534

GO:0006413,0.00115359719886507,3.7847850678733,2.59752925877763,9,94,translational initiation,0.00250030758520534

GO:1901566,0.00117301834024152,1.60179840570036,38.7142392717815,58,1401,organonitrogen compound biosynthetic process,0.00253125010262645

GO:0031589,0.00119299722516293,2.27300256231074,9.34005201560468,20,338,cell-substrate adhesion,0.00256312067589591

GO:0033002,0.00123314479378749,3.2039589961117,3.70286085825748,11,134,muscle cell proliferation,0.00263673952683651

GO:0009892,0.00123798136320983,1.4957938876312,58.4444083224967,81,2115,negative regulation of metabolic process,0.00263673952683651

GO:0070252,0.00124391948263911,3.74052437388193,2.62516254876463,9,95,actin-mediated cell contraction,0.00263796717870017

GO:0023051,0.00124959523929923,1.45037476998544,73.4769180754226,98,2659,regulation of signaling,0.00263863029070911

GO:0032373,0.00129302549821863,10.1398031896844,0.49739921976593,4,18,positive regulation of sterol transport,0.00270710019201518

GO:0032376,0.00129302549821863,10.1398031896844,0.49739921976593,4,18,positive regulation of cholesterol transport,0.00270710019201518

GO:0050731,0.00131005223455873,3.17790634252766,3.73049414824447,11,135,positive regulation of peptidyl-tyrosine phosphorylation,0.0027303693450368

GO:0008284,0.00131523889181651,1.73706035892995,25.0633940182055,41,907,positive regulation of cell population proliferation,0.0027303693450368

GO:0048568,0.00132800403631373,2.15788093364805,10.8046163849155,22,391,embryonic organ development,0.00274528565490064

GO:0007155,0.00138550254494432,1.58879148365425,38.9905721716515,58,1411,cell adhesion,0.00285216423478077

GO:0007423,0.0014241896228124,1.97759462635759,14.452210663199,27,523,sensory organ development,0.00290699377557511

GO:0051924,0.00143435626155912,2.81110363785238,4.9463589076723,13,179,regulation of calcium ion transport,0.00290699377557511

GO:0022612,0.00144167983991937,3.65502076048951,2.68042912873862,9,97,gland morphogenesis,0.00290699377557511

GO:0072009,0.00144167983991937,3.65502076048951,2.68042912873862,9,97,nephron epithelium development,0.00290699377557511

GO:0001936,0.00144167983991937,3.65502076048951,2.68042912873862,9,97,regulation of endothelial cell proliferation,0.00290699377557511

GO:0043552,0.00147736850852527,6.83562271062271,0.856631989596879,5,31,positive regulation of phosphatidylinositol 3-kinase activity,0.00296679716814055

GO:0010038,0.00150330294304698,2.4825900008431,6.85305591677503,16,248,response to metal ion,0.00300660588609396

GO:0007178,0.0015276977625918,2.40196078431373,7.51625487646294,17,272,transmembrane receptor protein serine/threonine kinase signaling pathway,0.00303492060837915

GO:0043933,0.00154034918644118,1.51733259556953,50.2096879063719,71,1817,protein-containing complex organization,0.00303492060837915

GO:0030174,0.00155447153112103,17.7120853080569,0.248699609882965,3,9,regulation of DNA-templated DNA replication initiation,0.00303492060837915

GO:0030949,0.00155447153112103,17.7120853080569,0.248699609882965,3,9,positive regulation of vascular endothelial growth factor receptor signaling pathway,0.00303492060837915

GO:1905941,0.00155447153112103,17.7120853080569,0.248699609882965,3,9,positive regulation of gonad development,0.00303492060837915

GO:1905939,0.00155447153112103,17.7120853080569,0.248699609882965,3,9,regulation of gonad development,0.00303492060837915

GO:0051049,0.00159573937868268,1.60973533885116,35.066644993498,53,1269,regulation of transport,0.00307963920775988

GO:0048025,0.00160241389672059,9.46318289786223,0.525032509752926,4,19,negative regulation of mRNA splicing, via spliceosome,0.00307963920775988

GO:2000352,0.00160241389672059,9.46318289786223,0.525032509752926,4,19,negative regulation of endothelial cell apoptotic process,0.00307963920775988

GO:1900273,0.00160241389672059,9.46318289786223,0.525032509752926,4,19,positive regulation of long-term synaptic potentiation,0.00307963920775988

GO:0071688,0.00171177212203198,6.58201058201058,0.884265279583875,5,32,striated muscle myosin thick filament assembly,0.00326430962806098

GO:0030241,0.00171177212203198,6.58201058201058,0.884265279583875,5,32,skeletal muscle myosin thick filament assembly,0.00326430962806098

GO:0071709,0.00176876443230288,5.20891786483497,1.29876462938882,6,47,membrane assembly,0.00335996950074524

GO:0006941,0.00185860699361227,3.03000743218135,3.89629388816645,11,141,striated muscle contraction,0.00350498058184232

GO:0030182,0.00185934945500172,1.60379997668726,34.4863459037711,52,1248,neuron differentiation,0.00350498058184232

GO:0006096,0.00187217149929728,4.37698312767565,1.76853055916775,7,64,glycolytic process,0.00351150394685606

GO:0001934,0.00187708442687631,1.99175824175824,13.2639791937581,25,480,positive regulation of protein phosphorylation,0.00351150394685606

GO:0002062,0.00191293349604567,3.49516617892977,2.79096228868661,9,101,chondrocyte differentiation,0.00355155954737535

GO:0003014,0.00191293349604567,3.49516617892977,2.79096228868661,9,101,renal system process,0.00355155954737535

GO:0032331,0.00195974329747073,8.87114014251781,0.552665799739922,4,20,negative regulation of chondrocyte differentiation,0.00361258666653325

GO:0090218,0.0019723311258987,6.34651360544218,0.911898569570871,5,33,positive regulation of lipid kinase activity,0.00361258666653325

GO:0031034,0.0019723311258987,6.34651360544218,0.911898569570871,5,33,myosin filament assembly,0.00361258666653325

GO:1904035,0.00197517441727123,5.08455506307535,1.32639791937581,6,48,regulation of epithelial cell apoptotic process,0.00361258666653325

GO:0043410,0.00200091243786423,2.33667302798982,7.70968790637191,17,279,positive regulation of MAPK cascade,0.00364610710899705

GO:0007530,0.00217534091675846,15.1807718348003,0.276332899869961,3,10,sex determination,0.00387749255655618

GO:0019374,0.00217534091675846,15.1807718348003,0.276332899869961,3,10,galactolipid metabolic process,0.00387749255655618

GO:0006681,0.00217534091675846,15.1807718348003,0.276332899869961,3,10,galactosylceramide metabolic process,0.00387749255655618

GO:0030048,0.00218000244407084,3.16494295767139,3.39889466840052,10,123,actin filament-based movement,0.00387749255655618

GO:0019220,0.00220425539505536,1.66319007223805,27.3845903771131,43,991,regulation of phosphate metabolic process,0.00387749255655618

GO:0001503,0.00221443225878829,2.19635752178856,9.14661898569571,19,331,ossification,0.00387749255655618

GO:0061333,0.00223909806108781,4.22804314329738,1.82379713914174,7,66,renal tubule morphogenesis,0.00387749255655618

GO:0046689,0.00224364240469039,70.7044917257683,0.0828998699609883,2,3,response to mercury ion,0.00387749255655618

GO:0097646,0.00224364240469039,70.7044917257683,0.0828998699609883,2,3,calcitonin family receptor signaling pathway,0.00387749255655618

GO:1990408,0.00224364240469039,70.7044917257683,0.0828998699609883,2,3,calcitonin gene-related peptide receptor signaling pathway,0.00387749255655618

GO:2000532,0.00224364240469039,70.7044917257683,0.0828998699609883,2,3,regulation of renal albumin absorption,0.00387749255655618

GO:0010512,0.00224364240469039,70.7044917257683,0.0828998699609883,2,3,negative regulation of phosphatidylinositol biosynthetic process,0.00387749255655618

GO:0097017,0.00224364240469039,70.7044917257683,0.0828998699609883,2,3,renal protein absorption,0.00387749255655618

GO:0097018,0.00224364240469039,70.7044917257683,0.0828998699609883,2,3,renal albumin absorption,0.00387749255655618

GO:0051174,0.00224610849312706,1.66131888623461,27.4122236671001,43,992,regulation of phosphorus metabolic process,0.00387749255655618

GO:0070296,0.00226071506316985,6.12725779967159,0.939531859557867,5,34,sarcoplasmic reticulum calcium ion transport,0.00388906227650198

GO:0042327,0.00229682535289239,1.93178643329636,14.203511053316,26,514,positive regulation of phosphorylation,0.0039340175224789

GO:0044085,0.00232275379343234,1.41186616074508,76.7100130039012,100,2776,cellular component biogenesis,0.0039340175224789

GO:0050817,0.00232682743707594,2.64958903353928,5.22269180754226,13,189,coagulation,0.0039340175224789

GO:0007596,0.00232682743707594,2.64958903353928,5.22269180754226,13,189,blood coagulation,0.0039340175224789

GO:2000146,0.00232682743707594,2.64958903353928,5.22269180754226,13,189,negative regulation of cell motility,0.0039340175224789

GO:0003015,0.00234517157080752,2.53874511538745,5.85825747724317,14,212,heart process,0.0039514534686209

GO:0060429,0.00239611025168057,1.68615716081154,25.0910273081925,40,908,epithelium development,0.00402350253865816

GO:0007599,0.00243683076751222,2.63444133618562,5.25032509752926,13,190,hemostasis,0.00405930960956482

GO:0007422,0.00244204955064828,4.85278802343241,1.3816644993498,6,50,peripheral nervous system development,0.00405930960956482

GO:0060350,0.00244218626916908,4.1572966507177,1.85143042912874,7,67,endochondral bone morphogenesis,0.00405930960956482

GO:0043534,0.00257413026809298,3.65910348644162,2.37646293888166,8,86,blood vessel endothelial cell migration,0.00424305595104357

GO:0051155,0.00257860514097973,5.92261904761905,0.967165149544863,5,35,positive regulation of striated muscle cell differentiation,0.00424305595104357

GO:0031033,0.00257860514097973,5.92261904761905,0.967165149544863,5,35,myosin filament organization,0.00424305595104357

GO:1902074,0.00259147478298009,2.27478314745973,7.90312093628088,17,286,response to salt,0.00425001864408734

GO:0045664,0.00260329051253959,2.73863759041347,4.67002600780234,12,169,regulation of neuron differentiation,0.00425521239923414

GO:0001570,0.00265916137076798,4.08886971527179,1.87906371911573,7,68,vasculogenesis,0.00431836884645574

GO:0065009,0.00265948325300018,1.47979869914067,51.2873862158648,71,1856,regulation of molecular function,0.00431836884645574

GO:0055003,0.00270441135686226,4.74463007159905,1.4092977893368,6,51,cardiac myofibril assembly,0.00436252586090567

GO:0044091,0.00270441135686226,4.74463007159905,1.4092977893368,6,51,membrane biogenesis,0.00436252586090567

GO:0043408,0.00272905702041606,1.96060638471541,12.9047464239272,24,467,regulation of MAPK cascade,0.00438789560145327

GO:0009611,0.00278380788046292,2.02276817002254,11.4678153446034,22,415,response to wounding,0.00445243140485762

GO:0007154,0.00282353601094038,1.33713579472913,131.147594278283,158,4746,cell communication,0.00445243140485762

GO:0003401,0.00283254274333422,7.88440221694378,0.607932379713914,4,22,axis elongation,0.00445243140485762

GO:0033119,0.00283254274333422,7.88440221694378,0.607932379713914,4,22,negative regulation of RNA splicing,0.00445243140485762

GO:1904036,0.00283254274333422,7.88440221694378,0.607932379713914,4,22,negative regulation of epithelial cell apoptotic process,0.00445243140485762

GO:0002053,0.00283254274333422,7.88440221694378,0.607932379713914,4,22,positive regulation of mesenchymal cell proliferation,0.00445243140485762

GO:0003422,0.00283254274333422,7.88440221694378,0.607932379713914,4,22,growth plate cartilage morphogenesis,0.00445243140485762

GO:0030240,0.00292768980530442,5.73118279569892,0.99479843953186,5,36,skeletal muscle thin filament assembly,0.00454769311878331

GO:0003417,0.00292768980530442,5.73118279569892,0.99479843953186,5,36,growth plate cartilage development,0.00454769311878331

GO:0007494,0.00293011934685835,13.2822867298578,0.303966189856957,3,11,midgut development,0.00454769311878331

GO:0071679,0.00293011934685835,13.2822867298578,0.303966189856957,3,11,commissural neuron axon guidance,0.00454769311878331

GO:0048660,0.00297325370119182,3.56714628297362,2.43172951885566,8,88,regulation of smooth muscle cell proliferation,0.00456450694035856

GO:0046580,0.00298733990812085,4.6411746394106,1.4369310793238,6,52,negative regulation of Ras protein signal transduction,0.00456450694035856

GO:0060675,0.00298733990812085,4.6411746394106,1.4369310793238,6,52,ureteric bud morphogenesis,0.00456450694035856

GO:0061005,0.00298733990812085,4.6411746394106,1.4369310793238,6,52,cell differentiation involved in kidney development,0.00456450694035856

GO:0072171,0.00298733990812085,4.6411746394106,1.4369310793238,6,52,mesonephric tubule morphogenesis,0.00456450694035856

GO:0000165,0.00300911800896068,1.84373728553992,15.9996749024707,28,579,MAPK cascade,0.0045835481746398

GO:0051098,0.00304597552474181,2.30102689486553,7.35045513654096,16,266,regulation of binding,0.00462140486611437

GO:0010810,0.00305275727944547,2.56119972260749,5.38849154746424,13,195,regulation of cell-substrate adhesion,0.00462140486611437

GO:0051897,0.00319007766880549,3.52287059241496,2.45936280884265,8,89,positive regulation of phosphatidylinositol 3-kinase/protein kinase B signal transduction,0.00481447304617271

GO:0051592,0.00326670276385138,2.9789156626506,3.59232769830949,10,130,response to calcium ion,0.00490005414577707

GO:0006937,0.00326670276385138,2.9789156626506,3.59232769830949,10,130,regulation of muscle contraction,0.00490005414577707

GO:0072132,0.00330966041086227,5.55171130952381,1.02243172951886,5,37,mesenchyme morphogenesis,0.00493440279437648

GO:0048546,0.00330966041086227,5.55171130952381,1.02243172951886,5,37,digestive tract morphogenesis,0.00493440279437648

GO:0003433,0.0033550041782375,7.46893361670209,0.63556566970091,4,23,chondrocyte development involved in endochondral bone morphogenesis,0.00494210196315224

GO:0055023,0.0033550041782375,7.46893361670209,0.63556566970091,4,23,positive regulation of cardiac muscle tissue growth,0.00494210196315224

GO:0021602,0.0033550041782375,7.46893361670209,0.63556566970091,4,23,cranial nerve morphogenesis,0.00494210196315224

GO:0003418,0.0033550041782375,7.46893361670209,0.63556566970091,4,23,growth plate cartilage chondrocyte differentiation,0.00494210196315224

GO:0048659,0.0034189932374976,3.47967479674797,2.48699609882965,8,90,smooth muscle cell proliferation,0.00501535291514919

GO:0001935,0.00342511906400433,3.18178789032749,3.03966189856957,9,110,endothelial cell proliferation,0.00501535291514919

GO:0014074,0.0034520187134219,2.95409738126058,3.61996098829649,10,131,response to purine-containing compound,0.00503974245401654

GO:0071840,0.00347094362344001,1.31982361814876,147.119635890767,174,5324,cellular component organization or biogenesis,0.0050523794755399

GO:0048708,0.00361880622956957,4.44719570405728,1.49219765929779,6,54,astrocyte differentiation,0.00525207275795938

GO:0001837,0.00363885548304803,3.15038178733032,3.06729518855657,9,111,epithelial to mesenchymal transition,0.00526563793429302

GO:0007189,0.00366044525573474,3.43751986362717,2.51462938881664,8,91,adenylate cyclase-activating G protein-coupled receptor signaling pathway,0.00528134623408063

GO:0071277,0.00367834116285353,3.83621641516378,1.98959687906372,7,72,cellular response to calcium ion,0.00529164869042086

GO:0010739,0.00382729624076263,11.8056872037915,0.331599479843953,3,12,positive regulation of protein kinase A signaling,0.00545805724769628

GO:0070885,0.00382729624076263,11.8056872037915,0.331599479843953,3,12,negative regulation of calcineurin-NFAT signaling cascade,0.00545805724769628

GO:0106057,0.00382729624076263,11.8056872037915,0.331599479843953,3,12,negative regulation of calcineurin-mediated signaling,0.00545805724769628

GO:1902075,0.00384767139705664,2.90567146635322,3.67522756827048,10,133,cellular response to salt,0.00547125528136377

GO:0050686,0.00393936353165762,7.09501187648456,0.663198959687906,4,24,negative regulation of mRNA processing,0.00556944499303318

GO:0061037,0.00393936353165762,7.09501187648456,0.663198959687906,4,24,negative regulation of cartilage development,0.00556944499303318

GO:0045165,0.00397193872486773,2.58858193062808,4.91872561768531,12,178,cell fate commitment,0.005586486228197

GO:0051781,0.00397412638184746,3.77783819051762,2.01723016905072,7,73,positive regulation of cell division,0.005586486228197

GO:0031399,0.0040887203152867,1.59280980085204,29.1531209362809,44,1055,regulation of protein modification process,0.00573119770689759

GO:0051050,0.0041600982886079,1.74277867705992,18.7077373211964,31,677,positive regulation of transport,0.00581468283521331

GO:0014866,0.00417901419277003,5.22443977591036,1.07769830949285,5,39,skeletal myofibril assembly,0.00582457502221772

GO:0009966,0.00427460905523336,1.40724734042553,64.6895318595579,85,2341,regulation of signal transduction,0.00592172499193302

GO:0030198,0.00435465574120803,2.15243544600939,8.31762028608583,17,301,extracellular matrix organization,0.00592172499193302

GO:0071493,0.00440518566473066,35.3498817966903,0.110533159947984,2,4,cellular response to UV-B,0.00592172499193302

GO:0090272,0.00440518566473066,35.3498817966903,0.110533159947984,2,4,negative regulation of fibroblast growth factor production,0.00592172499193302

GO:0030043,0.00440518566473066,35.3498817966903,0.110533159947984,2,4,actin filament fragmentation,0.00592172499193302

GO:1900238,0.00440518566473066,35.3498817966903,0.110533159947984,2,4,regulation of metanephric mesenchymal cell migration by platelet-derived growth factor receptor-beta signaling pathway,0.00592172499193302

GO:2000591,0.00440518566473066,35.3498817966903,0.110533159947984,2,4,positive regulation of metanephric mesenchymal cell migration,0.00592172499193302

GO:0014721,0.00440518566473066,35.3498817966903,0.110533159947984,2,4,twitch skeletal muscle contraction,0.00592172499193302

GO:0014724,0.00440518566473066,35.3498817966903,0.110533159947984,2,4,regulation of twitch skeletal muscle contraction,0.00592172499193302

GO:0035793,0.00440518566473066,35.3498817966903,0.110533159947984,2,4,positive regulation of metanephric mesenchymal cell migration by platelet-derived growth factor receptor-beta signaling pathway,0.00592172499193302

GO:0003010,0.00440518566473066,35.3498817966903,0.110533159947984,2,4,voluntary skeletal muscle contraction,0.00592172499193302

GO:0060484,0.00440518566473066,35.3498817966903,0.110533159947984,2,4,lung-associated mesenchyme development,0.00592172499193302

GO:0003415,0.00440518566473066,35.3498817966903,0.110533159947984,2,4,chondrocyte hypertrophy,0.00592172499193302

GO:0055006,0.0044645478697764,3.31693714795605,2.59752925877763,8,94,cardiac cell development,0.0059527304930352

GO:0150115,0.0044645478697764,3.31693714795605,2.59752925877763,8,94,cell-substrate junction organization,0.0059527304930352

GO:0007044,0.0044645478697764,3.31693714795605,2.59752925877763,8,94,cell-substrate junction assembly,0.0059527304930352

GO:0043062,0.00450070277046174,2.14473684210526,8.34525357607282,17,302,extracellular structure organization,0.00598471827855993

GO:0030336,0.00453911586634996,2.54211499061564,5.00162548764629,12,181,negative regulation of cell migration,0.00601952831871747

GO:0048566,0.00458889331570328,6.75670173057346,0.690832249674902,4,25,embryonic digestive tract development,0.00606812845311683

GO:0050806,0.00460043071750524,3.0306830551524,3.17782834850455,9,115,positive regulation of synaptic transmission,0.00606812845311683

GO:0050805,0.00466975722412542,5.07482993197279,1.10533159947984,5,40,negative regulation of synaptic transmission,0.00612672147805255

GO:0055021,0.00466975722412542,5.07482993197279,1.10533159947984,5,40,regulation of cardiac muscle tissue growth,0.00612672147805255

GO:0001649,0.00470430752392543,2.65826152239196,4.39369310793238,11,159,osteoblast differentiation,0.0061393084927621

GO:0035265,0.00470430752392543,2.65826152239196,4.39369310793238,11,159,organ growth,0.0061393084927621

GO:0050730,0.00474195586414194,2.52699045719983,5.02925877763329,12,182,regulation of peptidyl-tyrosine phosphorylation,0.00614767249976413

GO:0048008,0.00474486445786257,4.18475361504984,1.57509752925878,6,57,platelet-derived growth factor receptor signaling pathway,0.00614767249976413

GO:0001704,0.00476069760652466,3.27859092036715,2.62516254876463,8,95,formation of primary germ layer,0.00614767249976413

GO:0014902,0.00476069760652466,3.27859092036715,2.62516254876463,8,95,myotube differentiation,0.00614767249976413

GO:0060047,0.00485883773562726,2.41342622868354,5.6924577373212,13,206,heart contraction,0.00618483332225208

GO:0071313,0.00487434511033265,10.6244075829384,0.359232769830949,3,13,cellular response to caffeine,0.00618483332225208

GO:0010875,0.00487434511033265,10.6244075829384,0.359232769830949,3,13,positive regulation of cholesterol efflux,0.00618483332225208

GO:0006268,0.00487434511033265,10.6244075829384,0.359232769830949,3,13,DNA unwinding involved in DNA replication,0.00618483332225208

GO:0031581,0.00487434511033265,10.6244075829384,0.359232769830949,3,13,hemidesmosome assembly,0.00618483332225208

GO:0046716,0.00487434511033265,10.6244075829384,0.359232769830949,3,13,muscle cell cellular homeostasis,0.00618483332225208

GO:0042060,0.00487747018096302,2.07564732564733,9.11898569570871,18,330,wound healing,0.00618483332225208

GO:0043269,0.00495462773918503,1.9515349086455,11.3020156046814,21,409,regulation of monoatomic ion transport,0.00626652145932914

GO:0035107,0.00499485541613399,2.79122740963855,3.81339401820546,10,138,appendage morphogenesis,0.00630120221727673

GO:0035113,0.00514998715678351,2.97415865384615,3.23309492847854,9,117,embryonic appendage morphogenesis,0.00648029074459715

GO:0072078,0.00517189880999192,4.10400220304755,1.60273081924577,6,58,nephron tubule morphogenesis,0.00649126075131639

GO:0045785,0.00519673770587705,2.01769911504425,9.8927178153446,19,358,positive regulation of cell adhesion,0.00649352350628399

GO:0048741,0.00520009809243067,4.93353174603175,1.13296488946684,5,41,skeletal muscle fiber development,0.00649352350628399

GO:0006270,0.00530675822459716,6.44914705247247,0.718465539661899,4,26,DNA replication initiation,0.00660993682658684

GO:0055008,0.00534072085432028,3.561004784689,2.1277633289987,7,77,cardiac muscle tissue morphogenesis,0.00663544106142823

GO:0060349,0.00539797495225444,3.20448360412793,2.68042912873862,8,97,bone morphogenesis,0.00668968180480903

GO:0046683,0.00544317983783118,2.94667431192661,3.26072821846554,9,118,response to organophosphorus,0.0067287549754094

GO:0032879,0.00555677353165147,1.47322404371585,42.389466840052,59,1534,regulation of localization,0.00685196134729955

GO:0051058,0.00562643993237294,4.02629801413969,1.63036410923277,6,59,negative regulation of small GTPase mediated signal transduction,0.00692052111681871

GO:0003007,0.00572706235321339,2.20576769292283,7.15702210663199,15,259,heart morphogenesis,0.00701472155963588

GO:0006090,0.00573154078653176,3.51061392277108,2.1553966189857,7,78,pyruvate metabolic process,0.00701472155963588

GO:0055007,0.00606817491906497,2.89319126819127,3.31599479843953,9,120,cardiac muscle cell differentiation,0.00735724106839207

GO:0030325,0.00607778512420188,9.65790607496769,0.386866059817945,3,14,adrenal gland development,0.00735724106839207

GO:0051281,0.0060960067246264,6.16833625942373,0.746098829648895,4,27,positive regulation of release of sequestered calcium ion into cytosol,0.00735724106839207

GO:0001937,0.0060960067246264,6.16833625942373,0.746098829648895,4,27,negative regulation of endothelial cell proliferation,0.00735724106839207

GO:0060043,0.0060960067246264,6.16833625942373,0.746098829648895,4,27,regulation of cardiac muscle cell proliferation,0.00735724106839207

GO:0072088,0.00610948636408925,3.95147175815434,1.65799739921977,6,60,nephron epithelium morphogenesis,0.00735724106839207

GO:0090257,0.00611608048165113,2.439852520361,5.19505851755527,12,188,regulation of muscle system process,0.00735724106839207

GO:0003416,0.00638613315640836,4.67324561403509,1.18823146944083,5,43,endochondral bone growth,0.00765497690372241

GO:0050679,0.00639470631591445,2.68538816921823,3.95156046814044,10,143,positive regulation of epithelial cell proliferation,0.00765497690372241

GO:0016064,0.00657685955853453,3.41397391361342,2.21066319895969,7,80,immunoglobulin mediated immune response,0.00783490291234622

GO:0060415,0.00657685955853453,3.41397391361342,2.21066319895969,7,80,muscle tissue morphogenesis,0.00783490291234622

GO:0014909,0.00662203382758354,3.87936645693209,1.68563068920676,6,61,smooth muscle cell migration,0.00786966338930217

GO:0051017,0.00670804583487536,2.66516813522748,3.97919375812744,10,144,actin filament bundle assembly,0.00795267120664742

GO:0044093,0.0067720823376752,1.53516143299425,30.8387516254876,45,1116,positive regulation of molecular function,0.0080092896878274

GO:0030168,0.00686576922502148,3.06583120600294,2.79096228868661,8,101,platelet activation,0.00802333649331931

GO:0060536,0.00695956374460563,5.91092636579572,0.773732119635891,4,28,cartilage morphogenesis,0.00802333649331931

GO:0061098,0.00695956374460563,5.91092636579572,0.773732119635891,4,28,positive regulation of protein tyrosine kinase activity,0.00802333649331931

GO:0071312,0.00695956374460563,5.91092636579572,0.773732119635891,4,28,cellular response to alkaloid,0.00802333649331931

GO:0098801,0.00695956374460563,5.91092636579572,0.773732119635891,4,28,regulation of renal system process,0.00802333649331931

GO:0072210,0.00695956374460563,5.91092636579572,0.773732119635891,4,28,metanephric nephron development,0.00802333649331931

GO:0060421,0.00695956374460563,5.91092636579572,0.773732119635891,4,28,positive regulation of heart growth,0.00802333649331931

GO:0010595,0.00703270047964715,3.36761282813914,2.23829648894668,7,81,positive regulation of endothelial cell migration,0.00802333649331931

GO:0019724,0.00703270047964715,3.36761282813914,2.23829648894668,7,81,B cell mediated immunity,0.00802333649331931

GO:0098868,0.00704505268395727,4.55311355311355,1.21586475942783,5,44,bone growth,0.00802333649331931

GO:0014904,0.00704505268395727,4.55311355311355,1.21586475942783,5,44,myotube cell development,0.00802333649331931

GO:1903169,0.00710761122999638,2.81648532388664,3.39889466840052,9,123,regulation of calcium ion transmembrane transport,0.00802333649331931

GO:0010634,0.00710761122999638,2.81648532388664,3.39889466840052,9,123,positive regulation of epithelial cell migration,0.00802333649331931

GO:0003104,0.00720795676838848,23.565011820331,0.13816644993498,2,5,positive regulation of glomerular filtration,0.00802333649331931

GO:0071072,0.00720795676838848,23.565011820331,0.13816644993498,2,5,negative regulation of phospholipid biosynthetic process,0.00802333649331931

GO:0000395,0.00720795676838848,23.565011820331,0.13816644993498,2,5,mRNA 5'-splice site recognition,0.00802333649331931

GO:0061146,0.00720795676838848,23.565011820331,0.13816644993498,2,5,Peyer's patch morphogenesis,0.00802333649331931

GO:1903726,0.00720795676838848,23.565011820331,0.13816644993498,2,5,negative regulation of phospholipid metabolic process,0.00802333649331931

GO:1900020,0.00720795676838848,23.565011820331,0.13816644993498,2,5,positive regulation of protein kinase C activity,0.00802333649331931

GO:1900019,0.00720795676838848,23.565011820331,0.13816644993498,2,5,regulation of protein kinase C activity,0.00802333649331931

GO:0048865,0.00720795676838848,23.565011820331,0.13816644993498,2,5,stem cell fate commitment,0.00802333649331931

GO:0014034,0.00720795676838848,23.565011820331,0.13816644993498,2,5,neural crest cell fate commitment,0.00802333649331931

GO:2000561,0.00720795676838848,23.565011820331,0.13816644993498,2,5,regulation of CD4-positive, alpha-beta T cell proliferation,0.00802333649331931

GO:0009629,0.00720795676838848,23.565011820331,0.13816644993498,2,5,response to gravity,0.00802333649331931

GO:0002074,0.00720795676838848,23.565011820331,0.13816644993498,2,5,extraocular skeletal muscle development,0.00802333649331931

GO:0035739,0.00720795676838848,23.565011820331,0.13816644993498,2,5,CD4-positive, alpha-beta T cell proliferation,0.00802333649331931

GO:0043491,0.00737036911402577,2.6256201275691,4.03446033810143,10,146,phosphatidylinositol 3-kinase/protein kinase B signal transduction,0.00816716577500153

GO:0071241,0.00737036911402577,2.6256201275691,4.03446033810143,10,146,cellular response to inorganic substance,0.00816716577500153

GO:0010692,0.00744323973146067,8.85248815165877,0.414499349804941,3,15,regulation of alkaline phosphatase activity,0.00821092813425706

GO:0021554,0.00744323973146067,8.85248815165877,0.414499349804941,3,15,optic nerve development,0.00821092813425706

GO:0048589,0.00746511970507303,1.73098203784882,16.3312743823147,27,591,developmental growth,0.0082166418230334

GO:0072073,0.0074827352802127,2.79180602006689,3.42652795838752,9,124,kidney epithelium development,0.00821764678094787

GO:0090101,0.00770399261710631,3.00088350372334,2.8462288686606,8,103,negative regulation of transmembrane receptor protein serine/threonine kinase signaling pathway,0.00838023501015766

GO:0090497,0.00773959173480229,3.7427458861952,1.74089726918075,6,63,mesenchymal cell migration,0.00838023501015766

GO:0072028,0.00773959173480229,3.7427458861952,1.74089726918075,6,63,nephron morphogenesis,0.00838023501015766

GO:0001755,0.00773959173480229,3.7427458861952,1.74089726918075,6,63,neural crest cell migration,0.00838023501015766

GO:0048645,0.00775001408459702,4.4389880952381,1.24349804941482,5,45,animal organ formation,0.00838023501015766

GO:0010463,0.00775001408459702,4.4389880952381,1.24349804941482,5,45,mesenchymal cell proliferation,0.00838023501015766

GO:0001658,0.00775001408459702,4.4389880952381,1.24349804941482,5,45,branching involved in ureteric bud morphogenesis,0.00838023501015766

GO:0071248,0.00787279564778207,2.76755222148541,3.45416124837451,9,125,cellular response to metal ion,0.00839505486080927

GO:0007413,0.00790022439137132,5.67410926365796,0.801365409622887,4,29,axonal fasciculation,0.00839505486080927

GO:0032878,0.00790022439137132,5.67410926365796,0.801365409622887,4,29,regulation of establishment or maintenance of cell polarity,0.00839505486080927

GO:0051926,0.00790022439137132,5.67410926365796,0.801365409622887,4,29,negative regulation of calcium ion transport,0.00839505486080927

GO:0003009,0.00790022439137132,5.67410926365796,0.801365409622887,4,29,skeletal muscle contraction,0.00839505486080927

GO:0032371,0.00790022439137132,5.67410926365796,0.801365409622887,4,29,regulation of sterol transport,0.00839505486080927

GO:0032374,0.00790022439137132,5.67410926365796,0.801365409622887,4,29,regulation of cholesterol transport,0.00839505486080927

GO:0106030,0.00790022439137132,5.67410926365796,0.801365409622887,4,29,neuron projection fasciculation,0.00839505486080927

GO:0040013,0.00800700418351517,2.25913139786973,6.05169050715215,13,219,negative regulation of locomotion,0.00844312295256079

GO:0014032,0.00801410247732905,3.27855074288592,2.29356306892068,7,83,neural crest cell development,0.00844312295256079

GO:0048644,0.00801410247732905,3.27855074288592,2.29356306892068,7,83,muscle organ morphogenesis,0.00844312295256079

GO:0001823,0.00801410247732905,3.27855074288592,2.29356306892068,7,83,mesonephros development,0.00844312295256079

GO:0061572,0.00808214347565014,2.58721843897328,4.08972691807542,10,148,actin filament bundle organization,0.00849661237183732

GO:0030155,0.00820987818507671,1.69920375974169,17.2431729518856,28,624,regulation of cell adhesion,0.00861249481248985

GO:1901137,0.00840463372238915,1.75242718446602,14.9219765929779,25,540,carbohydrate derivative biosynthetic process,0.0087980421093946

GO:0031099,0.00845730230783357,2.56843200138684,4.11736020806242,10,149,regeneration,0.00882544287049728

GO:0003151,0.00850256081425957,4.33042973286876,1.27113133940182,5,46,outflow tract morphogenesis,0.00882544287049728

GO:0048146,0.00850256081425957,4.33042973286876,1.27113133940182,5,46,positive regulation of fibroblast proliferation,0.00882544287049728

GO:1905330,0.00850256081425957,4.33042973286876,1.27113133940182,5,46,regulation of morphogenesis of an epithelium,0.00882544287049728

GO:0061326,0.0085410040757113,3.23575467594606,2.32119635890767,7,84,renal tubule development,0.00882809664968478

GO:0051101,0.0085410040757113,3.23575467594606,2.32119635890767,7,84,regulation of DNA binding,0.00882809664968478

GO:0051896,0.00869920770255382,2.72027786831812,3.5094278283485,9,127,regulation of phosphatidylinositol 3-kinase/protein kinase B signal transduction,0.00897276769320016

GO:0002183,0.00892064861888381,5.45550886168463,0.828998699609883,4,30,cytoplasmic translational initiation,0.00917341865201773

GO:0019216,0.00893685383862681,2.15227175422658,6.82542262678804,14,247,regulation of lipid metabolic process,0.00917341865201773

GO:0010874,0.0089754924684204,8.17098067808968,0.442132639791938,3,16,regulation of cholesterol efflux,0.00917341865201773

GO:0010804,0.0089754924684204,8.17098067808968,0.442132639791938,3,16,negative regulation of tumor necrosis factor-mediated signaling pathway,0.00917341865201773

GO:0048010,0.00898696705339948,3.6153877270337,1.79616384915475,6,65,vascular endothelial growth factor receptor signaling pathway,0.00917341865201773

GO:0060993,0.00909293010831028,3.19405594405594,2.34882964889467,7,85,kidney morphogenesis,0.00926236358858935

GO:0010959,0.00911550706814094,2.08408961996597,7.54388816644993,15,273,regulation of metal ion transport,0.00926617660645731

GO:0043550,0.00930420324068301,4.22704081632653,1.29876462938882,5,47,regulation of lipid kinase activity,0.00938046720167221

GO:0044786,0.00930420324068301,4.22704081632653,1.29876462938882,5,47,cell cycle DNA replication,0.00938046720167221

GO:0035272,0.00930420324068301,4.22704081632653,1.29876462938882,5,47,exocrine system development,0.00938046720167221

GO:0060420,0.00930420324068301,4.22704081632653,1.29876462938882,5,47,regulation of heart growth,0.00938046720167221

GO:0045596,0.00935311326966843,1.80509504456892,12.7389466840052,22,461,negative regulation of cell differentiation,0.00941049433267253

GO:0070373,0.00966175395263067,3.55489260143198,1.82379713914174,6,66,negative regulation of ERK1 and ERK2 cascade,0.00969024089560216

GO:0048864,0.00967054528402574,3.15341287626431,2.37646293888166,7,86,stem cell development,0.00969024089560216

GO:0032970,0.0099510104188997,1.96195069667738,9.06371911573472,17,328,regulation of actin filament-based process,0.0099510104188997

Table S6. Supplementary data to accompany Figure 3D. Top gene ontology (Functional Enrichment Analysis) for each compartment (TTJ).

GOBPID,Pvalue,OddsRatio,ExpCount,Count,Size,Term,FDR

GO:0016266,1.20409797992351e-08,18.0226550079491,0.645838751625488,9,43,O-glycan processing,2.85371221241872e-06

GO:1901137,1.65456927934925e-07,3.61117965265256,8.11053315994798,26,540,carbohydrate derivative biosynthetic process,1.96066459602887e-05

GO:0006493,1.18178022784215e-06,8.3142037302726,1.38179453836151,10,92,protein O-linked glycosylation,8.96003856095736e-05

GO:1901135,1.51224279509829e-06,2.86311696658098,12.1507802340702,31,809,carbohydrate derivative metabolic process,8.96003856095736e-05

GO:0044281,4.70317188457938e-06,2.36431686952187,19.6455136540962,41,1308,small molecule metabolic process,0.000222930347329062

GO:0043207,1.07494789843438e-05,2.49947201689546,14.7191157347204,33,980,response to external biotic stimulus,0.000363946645612784

GO:0051707,1.07494789843438e-05,2.49947201689546,14.7191157347204,33,980,response to other organism,0.000363946645612784

GO:0019752,1.3981694171899e-05,2.84391637958033,9.68758127438231,25,645,carboxylic acid metabolic process,0.000401464772677023

GO:0070085,1.63948512321766e-05,4.26006280331145,3.60468140442133,14,240,glycosylation,0.000401464772677023

GO:0009607,1.69394418851065e-05,2.43894048847609,15.049544863459,33,1002,response to biotic stimulus,0.000401464772677023

GO:0043436,2.05790042247771e-05,2.77387049919731,9.91287386215865,25,660,oxoacid metabolic process,0.000438383089715794

GO:0006082,2.21966121375085e-05,2.7602565663329,9.95793237971391,25,663,organic acid metabolic process,0.000438383089715794

GO:0006955,2.73185475360698e-05,2.239959023333,18.9546163849155,38,1262,immune response,0.000498038135849888

GO:0006952,4.81532953910646e-05,2.27319996391143,16.5665149544863,34,1103,defense response,0.000740624273005612

GO:0043413,4.99999509202101e-05,4.06539315487411,3.48452535760728,13,232,macromolecule glycosylation,0.000740624273005612

GO:0006486,4.99999509202101e-05,4.06539315487411,3.48452535760728,13,232,protein glycosylation,0.000740624273005612

GO:0098542,6.10645822702442e-05,2.53062105724792,11.2496098829649,26,749,defense response to other organism,0.000790976037963095

GO:0010944,6.46936053983486e-05,66.4298245614035,0.0901170351105332,3,6,negative regulation of transcription by competitive promoter binding,0.000790976037963095

GO:0021603,6.46936053983486e-05,66.4298245614035,0.0901170351105332,3,6,cranial nerve formation,0.000790976037963095

GO:0009101,6.67490327395017e-05,3.52104664391354,4.6260078023407,15,308,glycoprotein biosynthetic process,0.000790976037963095

GO:0006665,7.19911207262663e-05,5.59385073146541,1.77230169050715,9,118,sphingolipid metabolic process,0.000812471219625006

GO:0002376,0.000114745098978968,1.92696563247979,27.9813394018205,48,1863,immune system process,0.00123611765718252

GO:0009100,0.000127409833572583,3.1651430098904,5.46710013003901,16,364,glycoprotein metabolic process,0.00131287524159575

GO:0044419,0.000166875651682106,2.14122486288848,16.9269830949285,33,1127,biological process involved in interspecies interaction between organisms,0.001565758624211

GO:0003404,0.000224623600097781,Inf,0.030039011703511,2,2,optic vesicle morphogenesis,0.001565758624211

GO:0003409,0.000224623600097781,Inf,0.030039011703511,2,2,optic cup structural organization,0.001565758624211

GO:0002265,0.000224623600097781,Inf,0.030039011703511,2,2,astrocyte activation involved in immune response,0.001565758624211

GO:0002282,0.000224623600097781,Inf,0.030039011703511,2,2,microglial cell activation involved in immune response,0.001565758624211

GO:1905673,0.000224623600097781,Inf,0.030039011703511,2,2,positive regulation of lysosome organization,0.001565758624211

GO:0021557,0.000224623600097781,Inf,0.030039011703511,2,2,oculomotor nerve development,0.001565758624211

GO:0021622,0.000224623600097781,Inf,0.030039011703511,2,2,oculomotor nerve morphogenesis,0.001565758624211

GO:0021623,0.000224623600097781,Inf,0.030039011703511,2,2,oculomotor nerve formation,0.001565758624211

GO:0032417,0.000224623600097781,Inf,0.030039011703511,2,2,positive regulation of sodium:proton antiporter activity,0.001565758624211

GO:0106016,0.000224623600097781,Inf,0.030039011703511,2,2,positive regulation of inflammatory response to wounding,0.001565758624211

GO:0048584,0.000256907674431866,1.92474489795918,24.1814044213264,42,1610,positive regulation of response to stimulus,0.0017396319668672

GO:0002072,0.000371234031846664,28.4624060150376,0.150195058517555,3,10,optic cup morphogenesis involved in camera-type eye development,0.00244395737632387

GO:0044255,0.000410203483152443,2.34202894470098,10.6037711313394,23,706,cellular lipid metabolic process,0.00262751960830079

GO:0008637,0.000424163541812979,6.82033898305085,0.976267880364109,6,65,apoptotic mitochondrial changes,0.00264544103709674

GO:0045087,0.000443823711967946,2.42904841402337,9.31209362808843,21,620,innate immune response,0.00269708255734367

GO:1901566,0.000500531539455671,1.92749115699731,21.0423276983095,37,1401,organonitrogen compound biosynthetic process,0.00284210372795495

GO:0034312,0.000504798342465511,24.9029605263158,0.165214564369311,3,11,diol biosynthetic process,0.00284210372795495

GO:0042339,0.000504798342465511,24.9029605263158,0.165214564369311,3,11,keratan sulfate metabolic process,0.00284210372795495

GO:0031349,0.000540482851325613,2.98971774193548,5.01651495448635,14,334,positive regulation of defense response,0.00284210372795495

GO:0006643,0.000587321648240587,4.13734142305571,2.34304291287386,9,156,membrane lipid metabolic process,0.00284210372795495

GO:0052547,0.000599008691646014,3.27886890025857,3.92009102730819,12,261,regulation of peptidase activity,0.00284210372795495

GO:0023052,0.000617233805855599,1.57379913081768,69.6154096228869,93,4635,signaling,0.00284210372795495

GO:0044282,0.000630200419363976,3.47302325581395,3.39440832249675,11,226,small molecule catabolic process,0.00284210372795495

GO:2000378,0.000663210837242219,11.5885845623444,0.405526657997399,4,27,negative regulation of reactive oxygen species metabolic process,0.00284210372795495

GO:1904220,0.000667180879052298,132.296943231441,0.0450585175552666,2,3,regulation of serine C-palmitoyltransferase activity,0.00284210372795495

GO:1904222,0.000667180879052298,132.296943231441,0.0450585175552666,2,3,positive regulation of serine C-palmitoyltransferase activity,0.00284210372795495

GO:1904217,0.000667180879052298,132.296943231441,0.0450585175552666,2,3,regulation of CDP-diacylglycerol-serine O-phosphatidyltransferase activity,0.00284210372795495

GO:1904219,0.000667180879052298,132.296943231441,0.0450585175552666,2,3,positive regulation of CDP-diacylglycerol-serine O-phosphatidyltransferase activity,0.00284210372795495

GO:0032415,0.000667180879052298,132.296943231441,0.0450585175552666,2,3,regulation of sodium:proton antiporter activity,0.00284210372795495

GO:1990858,0.000671551935719313,7.95775389801938,0.70591677503251,5,47,cellular response to lectin,0.00284210372795495

GO:1990840,0.000671551935719313,7.95775389801938,0.70591677503251,5,47,response to lectin,0.00284210372795495

GO:0002223,0.000671551935719313,7.95775389801938,0.70591677503251,5,47,stimulatory C-type lectin receptor signaling pathway,0.00284210372795495

GO:0031347,0.000795480954956952,2.49134242072151,7.7350455136541,18,515,regulation of defense response,0.00330752607587364

GO:0007154,0.000899003111528354,1.54822037142803,71.2825747724317,94,4746,cell communication,0.00367351271434862

GO:2000117,0.0011680213581531,6.9602691740413,0.796033810143043,5,53,negative regulation of cysteine-type endopeptidase activity,0.00444467503317289

GO:2000116,0.00116925169544987,3.43432023336472,3.10903771131339,10,207,regulation of cysteine-type endopeptidase activity,0.00444467503317289

GO:0050729,0.00119939889956155,5.50721461187215,1.18654096228869,6,79,positive regulation of inflammatory response,0.00444467503317289

GO:0002366,0.00120996432539744,2.73592753489232,5.45208062418726,14,363,leukocyte activation involved in immune response,0.00444467503317289

GO:0007165,0.00122041883883441,1.54260469004861,64.4787386215865,86,4293,signal transduction,0.00444467503317289

GO:0002263,0.0012419027544624,2.72792626728111,5.46710013003901,14,364,cell activation involved in immune response,0.00444467503317289

GO:0050728,0.00128191435859811,4.60998774509804,1.63712613784135,7,109,negative regulation of inflammatory response,0.00444467503317289

GO:0005975,0.00129585088152685,2.52320222912871,6.75877763328999,16,450,carbohydrate metabolic process,0.00444467503317289

GO:1903979,0.00132113070622304,66.1441048034934,0.0600780234070221,2,4,negative regulation of microglial cell activation,0.00444467503317289

GO:1905671,0.00132113070622304,66.1441048034934,0.0600780234070221,2,4,regulation of lysosome organization,0.00444467503317289

GO:0070172,0.00132113070622304,66.1441048034934,0.0600780234070221,2,4,positive regulation of tooth mineralization,0.00444467503317289

GO:0106014,0.00132113070622304,66.1441048034934,0.0600780234070221,2,4,regulation of inflammatory response to wounding,0.00444467503317289

GO:0046514,0.00133152711964251,16.5975877192982,0.225292587776333,3,15,ceramide catabolic process,0.00444467503317289

GO:0045071,0.00161430412297319,8.88046989720999,0.510663198959688,4,34,negative regulation of viral genome replication,0.00519072719931836

GO:0034311,0.00162073338712894,15.3198380566802,0.240312093628088,3,16,diol metabolic process,0.00519072719931836

GO:0035338,0.00162073338712894,15.3198380566802,0.240312093628088,3,16,long-chain fatty-acyl-CoA biosynthetic process,0.00519072719931836

GO:0008610,0.00180224932807647,2.3665469949092,7.64492847854356,17,509,lipid biosynthetic process,0.00545551392238653

GO:0002684,0.00182167753904753,2.11258326623234,11.1294538361508,22,741,positive regulation of immune system process,0.00545551392238653

GO:1901605,0.00186192037684393,3.81846515917692,2.23790637191157,8,149,alpha-amino acid metabolic process,0.00545551392238653

GO:0009913,0.0019419387355163,3.79132192256679,2.25292587776333,8,150,epidermal cell differentiation,0.00545551392238653

GO:0060900,0.00194634993525068,14.2246240601504,0.255331599479844,3,17,embryonic camera-type eye formation,0.00545551392238653

GO:0048268,0.00194634993525068,14.2246240601504,0.255331599479844,3,17,clathrin coat assembly,0.00545551392238653

GO:0051607,0.00196779990104629,3.1881243063263,3.33433029908973,10,222,defense response to virus,0.00545551392238653

GO:0140546,0.00196779990104629,3.1881243063263,3.33433029908973,10,222,defense response to symbiont,0.00545551392238653

GO:0002682,0.00209184791555471,1.91463992954407,15.7104031209363,28,1046,regulation of immune system process,0.00545551392238653

GO:1902564,0.0021800778127228,44.0931586608443,0.0750975292587776,2,5,negative regulation of neutrophil activation,0.00545551392238653

GO:0060671,0.0021800778127228,44.0931586608443,0.0750975292587776,2,5,epithelial cell differentiation involved in embryonic placenta development,0.00545551392238653

GO:0060672,0.0021800778127228,44.0931586608443,0.0750975292587776,2,5,epithelial cell morphogenesis involved in placental branching,0.00545551392238653

GO:0150079,0.0021800778127228,44.0931586608443,0.0750975292587776,2,5,negative regulation of neuroinflammatory response,0.00545551392238653

GO:0090594,0.0021800778127228,44.0931586608443,0.0750975292587776,2,5,inflammatory response to wounding,0.00545551392238653

GO:0051351,0.0021800778127228,44.0931586608443,0.0750975292587776,2,5,positive regulation of ligase activity,0.00545551392238653

GO:0002484,0.0021800778127228,44.0931586608443,0.0750975292587776,2,5,antigen processing and presentation of endogenous peptide antigen via MHC class I via ER pathway,0.00545551392238653

GO:0002486,0.0021800778127228,44.0931586608443,0.0750975292587776,2,5,antigen processing and presentation of endogenous peptide antigen via MHC class I via ER pathway, TAP-independent,0.00545551392238653

GO:0002536,0.0021800778127228,44.0931586608443,0.0750975292587776,2,5,respiratory burst involved in inflammatory response,0.00545551392238653

GO:0060264,0.0021800778127228,44.0931586608443,0.0750975292587776,2,5,regulation of respiratory burst involved in inflammatory response,0.00545551392238653

GO:0060266,0.0021800778127228,44.0931586608443,0.0750975292587776,2,5,negative regulation of respiratory burst involved in inflammatory response,0.00545551392238653

GO:1903900,0.00221523173218118,4.84048192771084,1.33673602080624,6,89,regulation of viral life cycle,0.00545551392238653

GO:0046949,0.00230990527274545,13.2754385964912,0.2703511053316,3,18,fatty-acyl-CoA biosynthetic process,0.00545551392238653

GO:0009084,0.00230990527274545,13.2754385964912,0.2703511053316,3,18,glutamine family amino acid biosynthetic process,0.00545551392238653

GO:1902307,0.00230990527274545,13.2754385964912,0.2703511053316,3,18,positive regulation of sodium ion transmembrane transport,0.00545551392238653

GO:0008544,0.00231541455969098,3.11362262026399,3.4094278283485,10,227,epidermis development,0.00545551392238653

GO:0043281,0.00233414592931016,3.35254591608183,2.85370611183355,9,190,regulation of cysteine-type endopeptidase activity involved in apoptotic process,0.00545551392238653

GO:0043065,0.00234443926775922,2.44733519404572,6.50344603381014,15,433,positive regulation of apoptotic process,0.00545551392238653

GO:0001817,0.00235733912147766,2.53483184625944,5.85760728218465,14,390,regulation of cytokine production,0.00545551392238653

GO:0009605,0.00237096174685997,1.68454597788926,27.8461638491547,43,1854,response to external stimulus,0.00545551392238653

GO:0032787,0.00241294679078558,2.52793702404381,5.87262678803641,14,391,monocarboxylic acid metabolic process,0.00549873451361714

GO:0050821,0.00244686034471926,4.08532608695652,1.83237971391417,7,122,protein stabilization,0.00551715299839825

GO:0006790,0.00246758741700513,3.08477447881155,3.43946684005202,10,229,sulfur compound metabolic process,0.00551715299839825

GO:0043525,0.00253450096608308,5.75640830027464,0.946228868660598,5,63,positive regulation of neuron apoptotic process,0.00558843673607602

GO:0050727,0.00254662939871819,3.07054709995886,3.45448634590377,10,230,regulation of inflammatory response,0.00558843673607602

GO:0044093,0.00269350593412458,1.85722353284086,16.7617685305592,29,1116,positive regulation of molecular function,0.00579226874297426

GO:0002833,0.00269495006649877,2.86326923076923,4.07028608582575,11,271,positive regulation of response to biotic stimulus,0.00579226874297426

GO:0035336,0.00271283472772212,12.4449013157895,0.285370611183355,3,19,long-chain fatty-acyl-CoA metabolic process,0.00579226874297426

GO:0050776,0.00279547922718361,2.11093396233283,10.078088426528,20,671,regulation of immune response,0.00591543372180818

GO:0043068,0.00291055746871953,2.38853193146417,6.6536410923277,15,443,positive regulation of programmed cell death,0.00599257208892111

GO:0002708,0.00291777510301517,4.56393939393939,1.41183355006502,6,94,positive regulation of lymphocyte mediated immunity,0.00599257208892111

GO:0016052,0.00293307325871244,3.9469537815126,1.8924577373212,7,126,carbohydrate catabolic process,0.00599257208892111

GO:0044242,0.00293307325871244,3.9469537815126,1.8924577373212,7,126,cellular lipid catabolic process,0.00599257208892111

GO:1904018,0.00306544282291285,3.91380208333333,1.90747724317295,7,127,positive regulation of vasculature development,0.00613881176146601

GO:0045766,0.00306544282291285,3.91380208333333,1.90747724317295,7,127,positive regulation of angiogenesis,0.00613881176146601

GO:0006979,0.0030997173335014,2.80830188679245,4.14538361508453,11,276,response to oxidative stress,0.00613881176146601

GO:0030149,0.00315648345387383,11.7120743034056,0.300390117035111,3,20,sphingolipid catabolic process,0.00613881176146601

GO:0046512,0.00323776991638503,33.0676855895196,0.0901170351105332,2,6,sphingosine biosynthetic process,0.00613881176146601

GO:0051340,0.00323776991638503,33.0676855895196,0.0901170351105332,2,6,regulation of ligase activity,0.00613881176146601

GO:0030259,0.00323776991638503,33.0676855895196,0.0901170351105332,2,6,lipid glycosylation,0.00613881176146601

GO:0001555,0.00323776991638503,33.0676855895196,0.0901170351105332,2,6,oocyte growth,0.00613881176146601

GO:0060268,0.00323776991638503,33.0676855895196,0.0901170351105332,2,6,negative regulation of respiratory burst,0.00613881176146601

GO:0009064,0.0033181047323096,5.38359977162432,1.00630689206762,5,67,glutamine family amino acid metabolic process,0.0061920537130502

GO:0048525,0.0033181047323096,5.38359977162432,1.00630689206762,5,67,negative regulation of viral process,0.0061920537130502

GO:0043066,0.00335136583323619,2.16540013366668,8.81644993498049,18,587,negative regulation of apoptotic process,0.00620526330060138

GO:0050896,0.00350943232421998,1.44268482490272,98.2876462938882,119,6544,response to stimulus,0.00640344333730518

GO:0090407,0.00351243727362732,2.33790999237223,6.7888166449935,15,452,organophosphate biosynthetic process,0.00640344333730518

GO:0031065,0.0036421093598437,11.0606725146199,0.315409622886866,3,21,positive regulation of histone deacetylation,0.00653924180517392

GO:0035337,0.0036421093598437,11.0606725146199,0.315409622886866,3,21,fatty-acyl-CoA metabolic process,0.00653924180517392

GO:0042981,0.00372697591892487,1.85084657008204,15.5902470741222,27,1038,regulation of apoptotic process,0.00664130295327214

GO:0006672,0.00376915869053156,5.21467090707965,1.03634590377113,5,69,ceramide metabolic process,0.00666634783325359

GO:0007155,0.00392844239548295,1.72613899855863,21.192522756827,34,1411,cell adhesion,0.00689659887207006

GO:0048679,0.00417088596347345,10.4778393351801,0.330429128738622,3,22,regulation of axon regeneration,0.00697731036801525

GO:0009226,0.00417088596347345,10.4778393351801,0.330429128738622,3,22,nucleotide-sugar biosynthetic process,0.00697731036801525

GO:0043388,0.00420412002831339,6.65594713656388,0.660858257477243,4,44,positive regulation of DNA binding,0.00697731036801525

GO:0090559,0.00420412002831339,6.65594713656388,0.660858257477243,4,44,regulation of membrane permeability,0.00697731036801525

GO:0072583,0.00420412002831339,6.65594713656388,0.660858257477243,4,44,clathrin-dependent endocytosis,0.00697731036801525

GO:0043069,0.0042091973825902,2.11514447509801,9.01170351105332,18,600,negative regulation of programmed cell death,0.00697731036801525

GO:0050731,0.00429568513969821,3.667236328125,2.027633289987,7,135,positive regulation of peptidyl-tyrosine phosphorylation,0.00697731036801525

GO:0080134,0.00445037794432308,1.86832786954146,14.253511053316,25,949,regulation of response to stress,0.00697731036801525

GO:0052548,0.00447554515575044,2.82284783884587,3.73985695708713,10,249,regulation of endopeptidase activity,0.00697731036801525

GO:0046520,0.00448809489960068,26.4524017467249,0.105136540962289,2,7,sphingoid biosynthetic process,0.00697731036801525

GO:0071281,0.00448809489960068,26.4524017467249,0.105136540962289,2,7,cellular response to iron ion,0.00697731036801525

GO:1900426,0.00448809489960068,26.4524017467249,0.105136540962289,2,7,positive regulation of defense response to bacterium,0.00697731036801525

GO:2000647,0.00448809489960068,26.4524017467249,0.105136540962289,2,7,negative regulation of stem cell proliferation,0.00697731036801525

GO:0002476,0.00448809489960068,26.4524017467249,0.105136540962289,2,7,antigen processing and presentation of endogenous peptide antigen via MHC class Ib,0.00697731036801525

GO:0002428,0.00448809489960068,26.4524017467249,0.105136540962289,2,7,antigen processing and presentation of peptide antigen via MHC class Ib,0.00697731036801525

GO:0007042,0.00448809489960068,26.4524017467249,0.105136540962289,2,7,lysosomal lumen acidification,0.00697731036801525

GO:0036446,0.00448809489960068,26.4524017467249,0.105136540962289,2,7,myofibroblast differentiation,0.00697731036801525

GO:0043085,0.00452420413436249,1.94702973650342,12.0005851755527,22,799,positive regulation of catalytic activity,0.00697731036801525

GO:1903036,0.00456080938533901,6.49317717846782,0.675877763328999,4,45,positive regulation of response to wounding,0.00697731036801525

GO:0044283,0.00456321986093824,2.34276179882409,6.30819245773732,14,420,small molecule biosynthetic process,0.00697731036801525

GO:0006629,0.00467413321040899,1.83772756745972,15.0795838751625,26,1004,lipid metabolic process,0.00710108699273673

GO:0007595,0.00474390517262892,9.95328947368421,0.345448634590377,3,23,lactation,0.00711585775894337

GO:0021602,0.00474390517262892,9.95328947368421,0.345448634590377,3,23,cranial nerve morphogenesis,0.00711585775894337

GO:0007586,0.00480034513856453,4.90662415408641,1.09642392717815,5,73,digestion,0.00715523143295467

GO:0043067,0.00499464298972059,1.80673284787803,15.9356957087126,27,1061,regulation of programmed cell death,0.00738679200310847

GO:0048518,0.00501803169831419,1.44209972327386,69.3750975292588,88,4619,positive regulation of biological process,0.00738679200310847

GO:1901564,0.00514918225804353,1.4361970338983,72.2588426527958,91,4811,organonitrogen compound metabolic process,0.00750685949735842

GO:0001816,0.00526577876259578,2.30196047801297,6.41332899869961,14,427,cytokine production,0.00750685949735842

GO:0090312,0.00536217999426199,9.47869674185464,0.360468140442133,3,24,positive regulation of protein deacetylation,0.00750685949735842

GO:0031063,0.00536217999426199,9.47869674185464,0.360468140442133,3,24,regulation of histone deacetylation,0.00750685949735842

GO:0045730,0.00536217999426199,9.47869674185464,0.360468140442133,3,24,respiratory burst,0.00750685949735842

GO:0046466,0.00536217999426199,9.47869674185464,0.360468140442133,3,24,membrane lipid catabolic process,0.00750685949735842

GO:0070570,0.00536217999426199,9.47869674185464,0.360468140442133,3,24,regulation of neuron projection regeneration,0.00750685949735842

GO:0032103,0.0053729116455017,2.29624435094281,6.42834850455137,14,428,positive regulation of response to external stimulus,0.00750685949735842

GO:0030148,0.00538466715000393,4.76580278128951,1.12646293888166,5,75,sphingolipid biosynthetic process,0.00750685949735842

GO:0043154,0.00575173092991675,6.04925911093312,0.720936280884265,4,48,negative regulation of cysteine-type endopeptidase activity involved in apoptotic process,0.00772062367610786

GO:0016477,0.00583421445444382,1.74761515281963,17.7079973992198,29,1179,cell migration,0.00772062367610786

GO:1903334,0.00592507803909551,22.042212518195,0.120156046814044,2,8,positive regulation of protein folding,0.00772062367610786

GO:0034625,0.00592507803909551,22.042212518195,0.120156046814044,2,8,fatty acid elongation, monounsaturated fatty acid,0.00772062367610786

GO:0034626,0.00592507803909551,22.042212518195,0.120156046814044,2,8,fatty acid elongation, polyunsaturated fatty acid,0.00772062367610786

GO:0018243,0.00592507803909551,22.042212518195,0.120156046814044,2,8,protein O-linked glycosylation via threonine,0.00772062367610786

GO:0002475,0.00592507803909551,22.042212518195,0.120156046814044,2,8,antigen processing and presentation via MHC class Ib,0.00772062367610786

GO:0019367,0.00592507803909551,22.042212518195,0.120156046814044,2,8,fatty acid elongation, saturated fatty acid,0.00772062367610786

GO:0019368,0.00592507803909551,22.042212518195,0.120156046814044,2,8,fatty acid elongation, unsaturated fatty acid,0.00772062367610786

GO:0019673,0.00592507803909551,22.042212518195,0.120156046814044,2,8,GDP-mannose metabolic process,0.00772062367610786

GO:0070170,0.00592507803909551,22.042212518195,0.120156046814044,2,8,regulation of tooth mineralization,0.00772062367610786

GO:0006954,0.00600143699965127,2.13610579115367,7.90026007802341,16,526,inflammatory response,0.00772062367610786

GO:0030212,0.00602664717333314,9.04724880382775,0.375487646293888,3,25,hyaluronan metabolic process,0.00772062367610786

GO:0018149,0.00602664717333314,9.04724880382775,0.375487646293888,3,25,peptide cross-linking,0.00772062367610786

GO:0010765,0.00602664717333314,9.04724880382775,0.375487646293888,3,25,positive regulation of sodium ion transport,0.00772062367610786

GO:0042119,0.00608649194320641,3.10552373052697,2.71853055916775,8,181,neutrophil activation,0.00775536876634365

GO:0046394,0.00628581230268127,2.85638704742478,3.31931079323797,9,221,carboxylic acid biosynthetic process,0.00796651077933402

GO:0006520,0.00646826353091938,2.84278644842025,3.33433029908973,9,222,amino acid metabolic process,0.00815414072780794

GO:1903034,0.00658002887017652,3.82069841269841,1.66716514954486,6,111,regulation of response to wounding,0.00825114731339596

GO:0021545,0.00665111084223045,5.78548170848496,0.750975292587776,4,50,cranial nerve development,0.00829638562951904

GO:0036230,0.00670073291683213,3.05197717081125,2.76358907672302,8,184,granulocyte activation,0.00831452199627861

GO:0016053,0.00684536943200557,2.81596480201131,3.36436931079324,9,224,organic acid biosynthetic process,0.00842988808151052

GO:0002705,0.00686753309609615,3.78440251572327,1.68218465539662,6,112,positive regulation of leukocyte mediated immunity,0.00842988808151052

GO:0002252,0.00690041471651072,2.10196641824549,8.02041612483745,16,534,immune effector process,0.00842988808151052

GO:0010951,0.00706243756236927,4.44660766961652,1.20156046814044,5,80,negative regulation of endopeptidase activity,0.00858357796041804

GO:0009617,0.0071343402066403,2.29292096712538,5.96274382314694,13,397,response to bacterium,0.00861866410261805

GO:0007173,0.00716403724985551,3.74878504672897,1.69720416124837,6,113,epidermal growth factor receptor signaling pathway,0.00861866410261805

GO:1903510,0.0074371989658155,4.38780857009781,1.2165799739922,5,81,mucopolysaccharide metabolic process,0.0086779727716316

GO:0002706,0.00746969636291214,3.71382716049383,1.71222366710013,6,114,regulation of lymphocyte mediated immunity,0.0086779727716316

GO:0060670,0.00754287928673464,18.8920773549595,0.1351755526658,2,9,branching involved in labyrinthine layer morphogenesis,0.0086779727716316

GO:0051764,0.00754287928673464,18.8920773549595,0.1351755526658,2,9,actin crosslink formation,0.0086779727716316

GO:0044090,0.00754287928673464,18.8920773549595,0.1351755526658,2,9,positive regulation of vacuole organization,0.0086779727716316

GO:2000679,0.00754287928673464,18.8920773549595,0.1351755526658,2,9,positive regulation of transcription regulatory region DNA binding,0.0086779727716316

GO:1905247,0.00754287928673464,18.8920773549595,0.1351755526658,2,9,positive regulation of aspartic-type peptidase activity,0.0086779727716316

GO:0002679,0.00754287928673464,18.8920773549595,0.1351755526658,2,9,respiratory burst involved in defense response,0.0086779727716316

GO:0006670,0.00754287928673464,18.8920773549595,0.1351755526658,2,9,sphingosine metabolic process,0.0086779727716316

GO:0009615,0.00759519854006955,2.60137318960848,4.04024707412224,10,269,response to virus,0.00869595195167383

GO:0002831,0.00786740925144484,2.35124081794719,5.36196358907672,12,357,regulation of response to biotic stimulus,0.00896430765669436

GO:0010822,0.00816729187598022,5.43018969702418,0.796033810143043,4,53,positive regulation of mitochondrion organization,0.00926147451965221

GO:1901342,0.0082994670197721,2.72589481373265,3.46950585175553,9,231,regulation of vasculature development,0.00934192515548299

GO:0048583,0.00831707260678022,1.46823252518703,44.788166449935,60,2982,regulation of response to stimulus,0.00934192515548299

GO:0050778,0.00844981330676317,2.05268977621764,8.20065019505852,16,546,positive regulation of immune response,0.00944625355520222

GO:0019637,0.00849480303454837,1.92475925200157,10.4085175552666,19,693,organophosphate metabolic process,0.0094519639398496

GO:2000377,0.00864431340090586,4.22034278032934,1.26163849154746,5,84,regulation of reactive oxygen species metabolic process,0.00948473275932726

GO:0010466,0.00864431340090586,4.22034278032934,1.26163849154746,5,84,negative regulation of peptidase activity,0.00948473275932726

GO:0002220,0.00864431340090586,4.22034278032934,1.26163849154746,5,84,innate immune response activating cell surface receptor signaling pathway,0.00948473275932726

GO:0045088,0.00877821009941033,2.41726384364821,4.77620286085826,11,318,regulation of innate immune response,0.00953699298408718

GO:0030216,0.0090750906555289,4.16731194690265,1.27665799739922,5,85,keratinocyte differentiation,0.00953699298408718

GO:0045069,0.00929438775083614,5.21655005614581,0.826072821846554,4,55,regulation of viral genome replication,0.00953699298408718

GO:0033148,0.00933579060045664,16.5294759825328,0.150195058517555,2,10,positive regulation of intracellular estrogen receptor signaling pathway,0.00953699298408718

GO:0048143,0.00933579060045664,16.5294759825328,0.150195058517555,2,10,astrocyte activation,0.00953699298408718

GO:1900424,0.00933579060045664,16.5294759825328,0.150195058517555,2,10,regulation of defense response to bacterium,0.00953699298408718

GO:1900125,0.00933579060045664,16.5294759825328,0.150195058517555,2,10,regulation of hyaluronan biosynthetic process,0.00953699298408718

GO:0048680,0.00933579060045664,16.5294759825328,0.150195058517555,2,10,positive regulation of axon regeneration,0.00953699298408718

GO:0018146,0.00933579060045664,16.5294759825328,0.150195058517555,2,10,keratan sulfate biosynthetic process,0.00953699298408718

GO:0045741,0.00933579060045664,16.5294759825328,0.150195058517555,2,10,positive regulation of epidermal growth factor-activated receptor activity,0.00953699298408718

GO:0002483,0.00933579060045664,16.5294759825328,0.150195058517555,2,10,antigen processing and presentation of endogenous peptide antigen,0.00953699298408718

GO:0035795,0.00933579060045664,16.5294759825328,0.150195058517555,2,10,negative regulation of mitochondrial membrane permeability,0.00953699298408718

GO:1905245,0.00933579060045664,16.5294759825328,0.150195058517555,2,10,regulation of aspartic-type peptidase activity,0.00953699298408718

GO:0019885,0.00933579060045664,16.5294759825328,0.150195058517555,2,10,antigen processing and presentation of endogenous peptide antigen via MHC class I,0.00953699298408718

GO:0046479,0.00933579060045664,16.5294759825328,0.150195058517555,2,10,glycosphingolipid catabolic process,0.00953699298408718

GO:0070572,0.00933579060045664,16.5294759825328,0.150195058517555,2,10,positive regulation of neuron projection regeneration,0.00953699298408718

GO:0050792,0.00950450577079902,3.51695906432749,1.80234070221066,6,120,regulation of viral process,0.00966767325184278

GO:0043523,0.00960166245430214,3.12479166666667,2.35806241872562,7,157,regulation of neuron apoptotic process,0.00972476069089575

GO:0030855,0.00984224838106891,2.12686243309706,6.90897269180754,14,460,epithelial cell differentiation,0.00992234026741076

GO:0045454,0.00989401835066537,5.11589291765503,0.84109232769831,4,56,cell redox homeostasis,0.00992234026741076

GO:0002285,0.00992234026741076,3.10389072847682,2.37308192457737,7,158,lymphocyte activation involved in immune response,0.00992234026741076

**Table S7. Supplement to Figure 4.** SL Versus ALL tissues. Complete list of 1390 transcript identities. Full Differential Expression Analysis (DEA) outputs from the RNEA (Regulatory Network Enrichment Analysis).

| gene | baseMean | log2 Fold Change | lfcSE | stat | pvalue | padj |
| --- | --- | --- | --- | --- | --- | --- |
| MYH4.1 | 447.568056 | 6.46395844 | 1.09992453 | 47.1185042 | 1.44E-09 | 1.06E-08 |
| PRVA.1 | 3535.83992 | 6.05140322 | 0.89102916 | 64.331453 | 3.56E-13 | 3.49E-12 |
| MYH4.2 | 31.4279496 | 5.82866905 | 1.29329323 | 29.0325882 | 7.70E-06 | 3.81E-05 |
| PRVA | 719.075562 | 5.3628963 | 0.93124715 | 43.1057048 | 9.84E-09 | 6.66E-08 |
| DUPD1 | 6.39349018 | 5.35836862 | 0.90698793 | 49.4328547 | 4.74E-10 | 3.68E-09 |
| ACTC.1 | 2758.62132 | 5.33771251 | 0.97777321 | 36.1254428 | 2.73E-07 | 1.59E-06 |
| MYOZ1 | 91.8899471 | 5.17539659 | 1.115815 | 27.4949336 | 1.58E-05 | 7.50E-05 |
| AT2A1 | 300.52685 | 5.15833996 | 0.75718347 | 62.1727106 | 1.01E-12 | 9.61E-12 |
| MLRS | 370.746976 | 5.11828253 | 0.91332775 | 45.873225 | 2.62E-09 | 1.88E-08 |
| TNNI2 | 227.383648 | 5.10566725 | 1.01694716 | 34.0647649 | 7.23E-07 | 4.05E-06 |
| RL3 | 17.5762678 | 4.98902807 | 0.74123119 | 68.1864726 | 5.48E-14 | 5.78E-13 |
| F16P2 | 5.54519634 | 4.97942598 | 1.09152421 | 24.1019903 | 7.62E-05 | 0.0003256 |
| MPSF | 17.0548493 | 4.91371704 | 0.74683884 | 62.3144792 | 9.46E-13 | 9.00E-12 |
| FOG2.3 | 4.82354161 | 4.87038547 | 1.01681905 | 36.5196023 | 2.26E-07 | 1.34E-06 |
| EPHB2 | 6.90048418 | 4.86945287 | 0.66301377 | 76.5371649 | 9.42E-16 | 1.15E-14 |
| AT1B4 | 4.60732331 | 4.86872151 | 0.90395505 | 39.644993 | 5.13E-08 | 3.24E-07 |
| DS13B | 5.54957671 | 4.80354472 | 0.77650499 | 57.6227106 | 9.16E-12 | 8.16E-11 |
| OBSCN | 5.36687892 | 4.72792077 | 0.79576937 | 50.8672655 | 2.38E-10 | 1.89E-09 |
| LDB3 | 24.722579 | 4.72560971 | 1.06359211 | 27.6276055 | 1.48E-05 | 7.07E-05 |
| SMYD1.1 | 4.13181493 | 4.71015908 | 1.13405027 | 23.9407741 | 8.21E-05 | 0.00034909 |
| SMYD1 | 5.4205426 | 4.69795243 | 0.96091609 | 38.4499119 | 9.05E-08 | 5.57E-07 |
| NDKA | 1002.14563 | 4.67035927 | 0.91656811 | 35.9194939 | 3.01E-07 | 1.75E-06 |
| DESM | 79.3812417 | 4.65646292 | 1.02966144 | 26.7592794 | 2.22E-05 | 0.00010363 |
| NEUV | 244.577101 | 4.64596719 | 0.82185306 | 65.2676573 | 2.26E-13 | 2.24E-12 |
| TECR | 8.54724464 | 4.54635188 | 0.56603386 | 92.0898847 | 4.74E-19 | 7.30E-18 |
| MYOM2 | 26.1166137 | 4.48839614 | 0.46978136 | 141.043524 | 1.69E-29 | 4.63E-28 |
| ADIPO | 17.5016309 | 4.35664763 | 0.66847496 | 64.1531799 | 3.88E-13 | 3.79E-12 |
| KCRM.2 | 636.551832 | 4.34799082 | 0.81761474 | 38.94925 | 7.14E-08 | 4.44E-07 |
| TNNT3.1 | 349.551916 | 4.22963751 | 0.82398986 | 38.5138382 | 8.78E-08 | 5.41E-07 |
| ACTN3 | 83.7029345 | 4.22928878 | 0.59580237 | 75.3304551 | 1.70E-15 | 2.03E-14 |
| FXI1C | 41.8609947 | 4.18307997 | 0.89670346 | 37.9492646 | 1.15E-07 | 6.98E-07 |
| CAV2.1 | 25.318338 | 4.17638935 | 0.7159501 | 50.7593368 | 2.51E-10 | 1.99E-09 |
| TITIN.1 | 104.168362 | 3.99841692 | 0.80607984 | 35.5073549 | 3.65E-07 | 2.11E-06 |
| SRCA | 13.6638068 | 3.91924862 | 0.65368956 | 57.2365574 | 1.10E-11 | 9.80E-11 |
| NEBU | 56.5269554 | 3.87452435 | 0.53481007 | 74.8380855 | 2.16E-15 | 2.55E-14 |
| NR0B1 | 4.61709448 | 3.82777969 | 0.70776922 | 40.2979737 | 3.76E-08 | 2.42E-07 |
| NTM1B | 5.03483889 | 3.79474383 | 0.94566752 | 25.0512023 | 4.91E-05 | 0.00021695 |
| MYCD.1 | 6.59531208 | 3.7879388 | 0.57319663 | 62.5175793 | 8.57E-13 | 8.20E-12 |
| ADT1 | 34.7029037 | 3.77232167 | 0.4894716 | 87.2835928 | 4.97E-18 | 7.10E-17 |
| DRS7C | 5.94211632 | 3.77056505 | 0.78923462 | 32.6723242 | 1.39E-06 | 7.50E-06 |
| NRX1A | 5.40300773 | 3.72522085 | 0.70843361 | 50.2684779 | 3.17E-10 | 2.50E-09 |
| ODF3A | 7.42163691 | 3.6981608 | 0.693736 | 41.7268289 | 1.90E-08 | 1.26E-07 |
| CYYR1 | 13.2132227 | 3.6896831 | 0.48266448 | 121.649754 | 2.37E-25 | 5.18E-24 |
| MYPC1 | 5.61655021 | 3.61125053 | 0.76885124 | 31.5230733 | 2.39E-06 | 1.25E-05 |
| FGF20 | 5.74655473 | 3.56553198 | 0.72313371 | 40.0513261 | 4.22E-08 | 2.69E-07 |
| UN45B | 5.84634545 | 3.55448373 | 0.67546939 | 37.6069732 | 1.35E-07 | 8.13E-07 |
| S47A1 | 6.36202048 | 3.51091143 | 0.71766596 | 35.4073086 | 3.83E-07 | 2.21E-06 |
| S26A2.1 | 833.541208 | 3.47353962 | 0.24071048 | 1263.24805 | 3.09E-272 | 2.37E-268 |
| MYOZ2.1 | 9.87489865 | 3.46521689 | 0.58656682 | 54.5795697 | 3.98E-11 | 3.38E-10 |
| TXLNB | 9.31047625 | 3.46277069 | 0.66144593 | 38.2143786 | 1.01E-07 | 6.20E-07 |
| ANK1.1 | 16.8050984 | 3.3803924 | 0.36635685 | 124.105218 | 7.09E-26 | 1.59E-24 |
| RSPO2 | 12.6049229 | 3.28774912 | 0.53862289 | 114.570407 | 7.71E-24 | 1.56E-22 |
| M2GD.2 | 4.27748749 | 3.27266069 | 0.83286973 | 26.6191156 | 2.37E-05 | 0.00011031 |
| PITX3 | 166.033482 | 3.26023991 | 0.26562797 | 298.029394 | 2.88E-63 | 4.60E-61 |
| MYL1 | 1423.88106 | 3.24988969 | 0.41238741 | 92.0107786 | 4.92E-19 | 7.57E-18 |
| ANGL3 | 58.5168394 | 3.24750032 | 0.41778593 | 180.848017 | 4.90E-38 | 2.05E-36 |
| PPR1C | 15.6982668 | 3.20370696 | 0.50491889 | 73.1323583 | 4.95E-15 | 5.68E-14 |
| TPM1.1 | 1320.89254 | 3.20245162 | 0.43687624 | 81.3320353 | 9.09E-17 | 1.19E-15 |
| WFKN2 | 8.58368415 | 3.18369322 | 0.52861772 | 66.7777113 | 1.09E-13 | 1.11E-12 |
| CCG1 | 5.96519206 | 3.17169629 | 0.78024351 | 28.6326828 | 9.28E-06 | 4.55E-05 |
| C1T9A | 5.94925762 | 3.16146393 | 0.60743941 | 33.5176045 | 9.36E-07 | 5.16E-06 |
| ASCL5 | 14.5437108 | 3.14868651 | 0.37149144 | 126.149172 | 2.59E-26 | 5.98E-25 |
| SOX12 | 10.1474676 | 3.12482151 | 0.54056427 | 65.6580679 | 1.87E-13 | 1.87E-12 |
| ACTM | 34.7805822 | 3.07253061 | 0.42287941 | 83.9247846 | 2.56E-17 | 3.50E-16 |
| SHH | 30.083863 | 3.06302295 | 0.27181883 | 222.516147 | 5.39E-47 | 3.77E-45 |
| EF1A2 | 69.6520728 | 3.04354268 | 0.33990731 | 110.810988 | 4.89E-23 | 9.43E-22 |
| RENI | 12.249974 | 3.03540182 | 0.37672401 | 76.3499458 | 1.03E-15 | 1.25E-14 |
| M3KCL | 4.87307785 | 3.01092273 | 0.70479315 | 26.4971691 | 2.51E-05 | 0.00011625 |
| GREM1.1 | 16.9057223 | 3.00320336 | 0.54138077 | 54.584512 | 3.97E-11 | 3.38E-10 |
| S13A4 | 163.680264 | 2.97810078 | 0.43553308 | 211.967788 | 1.00E-44 | 6.02E-43 |
| FREM2 | 24.7772491 | 2.96261648 | 0.33018761 | 154.240602 | 2.51E-32 | 7.83E-31 |
| SMAD1.1 | 12.5494128 | 2.94834635 | 0.59340338 | 78.2364275 | 4.12E-16 | 5.14E-15 |
| PPM1K.1 | 7.67398609 | 2.92020952 | 0.54612713 | 41.516518 | 2.10E-08 | 1.38E-07 |
| TMOD4 | 59.28135 | 2.91136613 | 0.3810475 | 82.3798782 | 5.45E-17 | 7.29E-16 |
| CALC | 26.8349619 | 2.90591334 | 0.52533537 | 74.0549568 | 3.16E-15 | 3.69E-14 |
| PITX3.1 | 46.4593855 | 2.90474005 | 0.34632232 | 173.917271 | 1.51E-36 | 5.85E-35 |
| GBX2 | 18.5388864 | 2.89658324 | 0.42758623 | 112.849324 | 1.80E-23 | 3.55E-22 |
| DAF2 | 8.9314787 | 2.89135453 | 0.59063954 | 28.2350197 | 1.12E-05 | 5.41E-05 |
| MBP.1 | 550.432599 | 2.88506172 | 0.29548021 | 175.253431 | 7.79E-37 | 3.06E-35 |
| LECA.2 | 164.021276 | 2.88304949 | 0.83769196 | 77.5303611 | 5.81E-16 | 7.19E-15 |
| ANXA3 | 933.960478 | 2.88004041 | 0.20885749 | 789.694331 | 1.31E-169 | 2.51E-166 |
| FA8.2 | 3.97463676 | 2.87252766 | 0.73903123 | 27.7297015 | 1.41E-05 | 6.76E-05 |
| CAV1.1 | 14.7326549 | 2.85394262 | 0.46559067 | 81.7399244 | 7.45E-17 | 9.83E-16 |
| TPM3.1 | 1247.02881 | 2.85020005 | 0.42838871 | 64.4711405 | 3.33E-13 | 3.27E-12 |
| CCDC3 | 24.9305395 | 2.83559651 | 0.33162573 | 108.850054 | 1.28E-22 | 2.41E-21 |
| MAG.3 | 17.7960673 | 2.82953963 | 0.36462789 | 100.478279 | 7.78E-21 | 1.32E-19 |
| FOXL1 | 8.76394742 | 2.82157854 | 0.57132429 | 35.9702345 | 2.93E-07 | 1.71E-06 |
| ADCY8 | 5.5962337 | 2.80725187 | 0.60550202 | 32.796237 | 1.31E-06 | 7.12E-06 |
| LMOD3 | 3.70781446 | 2.75050844 | 0.79664016 | 22.5777733 | 0.00015375 | 0.00062733 |
| TMM88 | 9.28331218 | 2.74044404 | 0.65054744 | 58.6717405 | 5.52E-12 | 4.99E-11 |
| TBX22 | 26.5708609 | 2.72523465 | 0.33430594 | 114.998467 | 6.25E-24 | 1.27E-22 |
| PDE9A | 4.97492374 | 2.70736582 | 0.67894649 | 23.2010888 | 0.00011543 | 0.00047944 |
| PA2G3.1 | 7.13788291 | 2.70724532 | 0.59281615 | 48.3317215 | 8.05E-10 | 6.10E-09 |
| KLH41 | 17.7306183 | 2.70630974 | 0.34486595 | 84.0164569 | 2.45E-17 | 3.35E-16 |
| MBPHL | 40.8233684 | 2.69593478 | 0.29314514 | 134.589403 | 4.06E-28 | 1.03E-26 |
| DLK1 | 3.32958231 | 2.68844204 | 0.78874063 | 21.7313568 | 0.00022667 | 0.00090033 |
| CAC1S.1 | 6.45879914 | 2.68783045 | 0.53829353 | 34.0784919 | 7.18E-07 | 4.03E-06 |
| ZIC4 | 19.9380184 | 2.67488175 | 0.56945479 | 105.471382 | 6.72E-22 | 1.21E-20 |
| CRYAB | 22.2403702 | 2.65380191 | 0.50609767 | 43.1330428 | 9.71E-09 | 6.58E-08 |
| CBLN1 | 29.4015304 | 2.65314525 | 0.26142183 | 182.292736 | 2.40E-38 | 1.03E-36 |
| CASQ1 | 93.2016272 | 2.6301387 | 0.25473453 | 154.18539 | 2.58E-32 | 8.03E-31 |
| PLS2 | 59.7739749 | 2.62863251 | 0.39580589 | 236.174457 | 6.18E-50 | 5.32E-48 |
| THS7A | 7.02763187 | 2.61995842 | 0.49883893 | 43.2980586 | 8.97E-09 | 6.10E-08 |
| BHMT1 | 33.5354029 | 2.55605271 | 0.35026902 | 70.2459906 | 2.01E-14 | 2.20E-13 |
| RBM24 | 6.21359787 | 2.53691531 | 0.63069737 | 23.9496675 | 8.18E-05 | 0.00034785 |
| KCNA2 | 7.89610235 | 2.52263743 | 0.54124406 | 30.5675405 | 3.75E-06 | 1.92E-05 |
| UD2A1 | 10.7597161 | 2.50273213 | 0.65116812 | 57.9300351 | 7.89E-12 | 7.07E-11 |
| FOXD2.1 | 15.6158181 | 2.49844599 | 0.31229565 | 93.7487558 | 2.10E-19 | 3.30E-18 |
| NAR5 | 24.4478486 | 2.49039 | 0.28178241 | 108.436915 | 1.57E-22 | 2.93E-21 |
| RASF9 | 8.63981261 | 2.48148657 | 0.41206351 | 43.5886302 | 7.81E-09 | 5.35E-08 |
| HSPB1 | 31.6277627 | 2.47760445 | 0.52274211 | 29.2105443 | 7.08E-06 | 3.52E-05 |
| LBX1 | 7.23673845 | 2.4657917 | 0.51155429 | 36.3082726 | 2.50E-07 | 1.47E-06 |
| BAT1 | 5.48545462 | 2.45476772 | 0.67548511 | 33.2826252 | 1.05E-06 | 5.75E-06 |
| CAVN4 | 7.19882321 | 2.45044936 | 0.61733331 | 24.8702091 | 5.34E-05 | 0.00023429 |
| PLVAP | 22.3996226 | 2.43147191 | 0.33006639 | 89.0456252 | 2.10E-18 | 3.08E-17 |
| CATL1 | 6.33391664 | 2.41835888 | 0.70476414 | 22.232762 | 0.00018013 | 0.00072627 |
| FXYD6 | 14.8588789 | 2.41375027 | 0.45707004 | 72.1434296 | 8.00E-15 | 9.08E-14 |
| RYR3.1 | 6.79078801 | 2.39054404 | 0.46508151 | 36.4945167 | 2.29E-07 | 1.35E-06 |
| STC1 | 126.635455 | 2.37679298 | 0.61489568 | 58.9764428 | 4.76E-12 | 4.32E-11 |
| LYAM3 | 14.0929826 | 2.36672099 | 0.36647513 | 49.4407639 | 4.72E-10 | 3.67E-09 |
| AMPE | 16.0947157 | 2.36612189 | 0.47378808 | 129.126326 | 5.99E-27 | 1.44E-25 |
| TBX15 | 6.95289258 | 2.35668264 | 0.53913452 | 34.1144399 | 7.06E-07 | 3.96E-06 |
| TR19L | 184.701014 | 2.34331016 | 0.17767334 | 615.453646 | 7.01E-132 | 8.25E-129 |
| FGF3 | 33.7218023 | 2.34101366 | 0.2341153 | 206.836409 | 1.27E-43 | 7.16E-42 |
| CO6 | 130.413129 | 2.33767494 | 0.34400282 | 159.782705 | 1.63E-33 | 5.49E-32 |
| MAF | 11.3230479 | 2.33379222 | 0.50051072 | 32.3580998 | 1.62E-06 | 8.65E-06 |
| ROBO1 | 40.354327 | 2.32937223 | 0.2026399 | 210.734907 | 1.85E-44 | 1.07E-42 |
| CO7 | 116.697275 | 2.32573488 | 0.23132608 | 278.412154 | 4.90E-59 | 6.64E-57 |
| EST2E | 12.5107124 | 2.29295145 | 0.48617734 | 31.8422191 | 2.06E-06 | 1.09E-05 |
| AGRE1.2 | 19.0865394 | 2.2836396 | 0.28092283 | 74.0736845 | 3.13E-15 | 3.66E-14 |
| NCKX2 | 5.13896553 | 2.27700782 | 0.63822659 | 30.9902203 | 3.08E-06 | 1.59E-05 |
| NCAM1.1 | 17.9737615 | 2.27388285 | 0.28801982 | 79.7992071 | 1.92E-16 | 2.47E-15 |
| BNIP3.1 | 22.0098163 | 2.2702815 | 0.44567459 | 67.3548533 | 8.21E-14 | 8.48E-13 |
| S38A4 | 5.11567438 | 2.25666893 | 0.56831956 | 21.7757156 | 0.00022211 | 0.00088319 |
| PTC1.1 | 34.6824331 | 2.2389812 | 0.23025322 | 215.453063 | 1.78E-45 | 1.14E-43 |
| BTBBA | 1157.85937 | 2.23707657 | 0.25156284 | 137.933032 | 7.82E-29 | 2.07E-27 |
| PDPN | 199.876591 | 2.2184543 | 0.31215469 | 196.886882 | 1.75E-41 | 9.13E-40 |
| DTNA | 19.5329697 | 2.21205797 | 0.28974702 | 99.8497478 | 1.06E-20 | 1.78E-19 |
| LORF2.65 | 9.23603093 | 2.20891413 | 0.70047966 | 22.3923164 | 0.00016741 | 0.00067857 |
| A2MG | 15.7997399 | 2.19568905 | 0.31214583 | 63.761109 | 4.69E-13 | 4.57E-12 |
| ALK | 5.46970093 | 2.19379714 | 0.7730657 | 43.3439707 | 8.78E-09 | 5.97E-08 |
| ANF1 | 11.2828717 | 2.1876333 | 0.43762026 | 66.4445379 | 1.28E-13 | 1.29E-12 |
| SOAT | 18.3477674 | 2.18675406 | 0.29641972 | 62.7304557 | 7.73E-13 | 7.42E-12 |
| THSD1 | 5.12470215 | 2.18394313 | 0.66338784 | 28.9047476 | 8.17E-06 | 4.02E-05 |
| SX17A | 14.4534908 | 2.17400652 | 0.32994462 | 49.8982804 | 3.79E-10 | 2.96E-09 |
| KREM2 | 18.8581914 | 2.16827684 | 0.31401407 | 116.155216 | 3.54E-24 | 7.32E-23 |
| MYOC | 52.8685862 | 2.1347729 | 0.22965973 | 195.901624 | 2.86E-41 | 1.46E-39 |
| ZP3 | 10.1656201 | 2.13149193 | 0.46273516 | 28.4225119 | 1.02E-05 | 4.99E-05 |
| HAAF.1 | 5.41203837 | 2.12468176 | 0.78435632 | 22.1149709 | 0.00019013 | 0.00076316 |
| TECTA.1 | 10.6654666 | 2.12386427 | 0.49340199 | 87.2509106 | 5.05E-18 | 7.20E-17 |
| DPEP1 | 13.5073988 | 2.12308415 | 0.32273004 | 47.6481837 | 1.12E-09 | 8.35E-09 |
| FOXL1.1 | 8.33448961 | 2.12267548 | 0.41138629 | 37.8676057 | 1.19E-07 | 7.23E-07 |
| KCJ12 | 90.2646552 | 2.11255441 | 0.3485459 | 210.633358 | 1.94E-44 | 1.13E-42 |
| FOXD2 | 5.96214054 | 2.08715328 | 0.6371027 | 38.130206 | 1.05E-07 | 6.43E-07 |
| S13A2.1 | 78.3521603 | 2.06677225 | 0.35205736 | 154.121182 | 2.66E-32 | 8.27E-31 |
| NCAM1 | 14.2319255 | 2.06158791 | 0.44668044 | 38.4093887 | 9.22E-08 | 5.67E-07 |
| CALGL | 58.193057 | 2.06078864 | 0.67239343 | 25.5574694 | 3.89E-05 | 0.0001744 |
| S17AB | 17.32374 | 2.05667789 | 0.39047712 | 33.7091947 | 8.55E-07 | 4.74E-06 |
| MMEL1 | 12.3640389 | 2.03577001 | 0.46994809 | 39.5952392 | 5.25E-08 | 3.31E-07 |
| SFRP5 | 80.9833088 | 2.03164581 | 0.19363249 | 176.197347 | 4.89E-37 | 1.94E-35 |
| SP7 | 46.3616967 | 2.02869567 | 0.41855069 | 216.213773 | 1.22E-45 | 8.00E-44 |
| SGCG | 14.1585807 | 2.02402103 | 0.31792997 | 55.9252644 | 2.08E-11 | 1.82E-10 |
| HSPB8 | 49.7242445 | 2.0235535 | 0.31943998 | 51.2921177 | 1.94E-10 | 1.55E-09 |
| CNMD.1 | 227.500892 | 2.02239128 | 0.7887039 | 42.8317225 | 1.12E-08 | 7.57E-08 |
| VWA2 | 42.6253624 | 2.02048525 | 0.22050851 | 249.89138 | 6.87E-53 | 6.96E-51 |
| RYR1.3 | 25.2201808 | 2.00653061 | 0.24227069 | 84.7042398 | 1.75E-17 | 2.41E-16 |
| DCK | 16.792277 | 2.00262341 | 0.51402504 | 26.0451049 | 3.10E-05 | 0.00014132 |
| TIE2.1 | 8.57480742 | 1.99994762 | 0.37055061 | 31.3218811 | 2.63E-06 | 1.37E-05 |
| ISL1 | 11.5607645 | 1.99970945 | 0.37086266 | 49.6766025 | 4.22E-10 | 3.29E-09 |
| TBX18 | 8.83524136 | 1.99161311 | 0.46515687 | 40.1959971 | 3.94E-08 | 2.53E-07 |
| ETS1A | 7.84529519 | 1.98957946 | 0.41595136 | 28.3221845 | 1.07E-05 | 5.22E-05 |
| S38A8.1 | 3.87806148 | 1.98538228 | 1.12479519 | 22.5383051 | 0.00015656 | 0.00063847 |
| CXA4 | 12.7271992 | 1.97421204 | 0.30451235 | 46.5540384 | 1.89E-09 | 1.38E-08 |
| DC1I1 | 16.6773026 | 1.97106984 | 0.39024018 | 95.5304006 | 8.79E-20 | 1.41E-18 |
| TCIM | 36.8291105 | 1.97059717 | 0.34674912 | 43.9110425 | 6.69E-09 | 4.63E-08 |
| SEM6A | 69.4191373 | 1.96906568 | 0.17453612 | 289.040233 | 2.50E-61 | 3.79E-59 |
| FZD9 | 8.34979399 | 1.94737921 | 0.50056279 | 29.2084799 | 7.09E-06 | 3.52E-05 |
| CGT | 13.2370856 | 1.94500741 | 0.37456008 | 43.1407992 | 9.67E-09 | 6.56E-08 |
| LYPD6 | 24.3676007 | 1.9371212 | 0.2742335 | 64.6319509 | 3.08E-13 | 3.03E-12 |
| TSP4 | 27.4159545 | 1.93636384 | 0.36354028 | 108.538221 | 1.49E-22 | 2.79E-21 |
| G3ST1 | 13.2956712 | 1.93482557 | 0.38304243 | 32.0637609 | 1.86E-06 | 9.86E-06 |
| FOXD1 | 17.5050843 | 1.93434383 | 0.36002593 | 53.8016494 | 5.79E-11 | 4.85E-10 |
| MYCT1 | 11.7793671 | 1.92849898 | 0.35239023 | 32.0251675 | 1.89E-06 | 1.00E-05 |
| LAMA5 | 148.898166 | 1.92721203 | 0.11228447 | 520.078065 | 3.04E-111 | 1.86E-108 |
| LMO3 | 14.9765242 | 1.92255335 | 0.47753448 | 126.620693 | 2.06E-26 | 4.79E-25 |
| APJA | 7.92070217 | 1.92079992 | 0.46547573 | 26.9188785 | 2.06E-05 | 9.66E-05 |
| PLTP | 67.7743081 | 1.91977828 | 0.29316931 | 79.4112084 | 2.32E-16 | 2.95E-15 |
| SIA4B | 15.5657923 | 1.91800855 | 0.37370942 | 43.8330074 | 6.95E-09 | 4.80E-08 |
| EVA1C | 36.6425987 | 1.9104034 | 0.22756366 | 152.237447 | 6.75E-32 | 2.07E-30 |
| CALRL | 7.18757264 | 1.90937126 | 0.46143149 | 33.8181317 | 8.12E-07 | 4.52E-06 |
| SP5 | 6.12516841 | 1.89881347 | 0.62908777 | 24.7932474 | 5.54E-05 | 0.00024202 |
| EFC10 | 12.1510533 | 1.89045929 | 0.40285974 | 39.4487651 | 5.63E-08 | 3.54E-07 |
| CRDL1 | 214.35114 | 1.88285687 | 0.40019297 | 33.5531482 | 9.20E-07 | 5.09E-06 |
| RTN4R | 4.67439937 | 1.87990193 | 0.76643931 | 36.4157103 | 2.38E-07 | 1.40E-06 |
| FOXF2 | 28.2494028 | 1.86513456 | 0.28507903 | 81.7659517 | 7.36E-17 | 9.72E-16 |
| TM182.1 | 10.3126566 | 1.85422903 | 0.43719364 | 33.6124179 | 8.95E-07 | 4.96E-06 |
| WNT11 | 73.6966588 | 1.85271402 | 0.20352828 | 132.5069 | 1.13E-27 | 2.82E-26 |
| AHNK.2 | 37.5076824 | 1.84229445 | 0.27559287 | 57.7065464 | 8.79E-12 | 7.85E-11 |
| ADA2C.1 | 9.20947391 | 1.83955102 | 0.44937975 | 45.1609913 | 3.68E-09 | 2.61E-08 |
| APCD1.1 | 148.333191 | 1.83854724 | 0.14817783 | 662.950387 | 3.66E-142 | 4.67E-139 |
| HMCN1.1 | 5.84020486 | 1.82577568 | 0.69354607 | 45.2169043 | 3.58E-09 | 2.54E-08 |
| INHBA | 17.3944992 | 1.82411959 | 0.45468697 | 88.732257 | 2.45E-18 | 3.57E-17 |
| MXRA5 | 7.73454115 | 1.8226407 | 0.59599513 | 59.3435279 | 3.99E-12 | 3.64E-11 |
| EPHA3 | 156.866442 | 1.81773792 | 0.18879863 | 191.10879 | 3.06E-40 | 1.44E-38 |
| RSPO3 | 45.2437001 | 1.80968843 | 0.30256117 | 198.081591 | 9.71E-42 | 5.11E-40 |
| IRX3 | 5.71044219 | 1.80777475 | 0.5590878 | 24.1340802 | 7.51E-05 | 0.00032143 |
| CHSTB.1 | 17.4689326 | 1.80085963 | 0.36964618 | 99.2270689 | 1.44E-20 | 2.39E-19 |
| HHATL | 11.2634304 | 1.79190703 | 0.33756427 | 32.2959375 | 1.66E-06 | 8.89E-06 |
| S13A2 | 147.67147 | 1.78843792 | 0.34287192 | 156.137572 | 9.84E-33 | 3.16E-31 |
| AGRL4 | 16.6745162 | 1.77325682 | 0.28800172 | 41.4937844 | 2.12E-08 | 1.40E-07 |
| IHH | 21.2838313 | 1.77045214 | 0.47763814 | 59.1213608 | 4.44E-12 | 4.04E-11 |
| YG31B.6 | 5.23195227 | 1.74911334 | 0.72673816 | 38.9635264 | 7.09E-08 | 4.41E-07 |
| NDNF.2 | 107.572184 | 1.74605337 | 0.2053469 | 295.236602 | 1.15E-62 | 1.82E-60 |
| SFRP3 | 333.221971 | 1.74380475 | 0.20045017 | 96.2482906 | 6.18E-20 | 9.99E-19 |
| MYCN.2 | 98.5991559 | 1.73465806 | 0.16583509 | 169.26158 | 1.51E-35 | 5.57E-34 |
| ABCA1.1 | 24.0258614 | 1.72616118 | 0.37364068 | 105.037324 | 8.32E-22 | 1.48E-20 |
| BPI | 16.6775391 | 1.72595475 | 0.53528402 | 60.3968291 | 2.39E-12 | 2.23E-11 |
| NUD15 | 21.8865441 | 1.72388728 | 0.34072242 | 95.3506768 | 9.60E-20 | 1.54E-18 |
| FGF8A | 4.6684041 | 1.721986 | 0.68092334 | 24.2266981 | 7.19E-05 | 0.00030901 |
| ETV5 | 62.4501672 | 1.72198175 | 0.20538342 | 129.081015 | 6.12E-27 | 1.47E-25 |
| LEF1 | 131.039093 | 1.71754387 | 0.19002895 | 242.815926 | 2.30E-51 | 2.18E-49 |
| LIPH | 15.2683624 | 1.71626057 | 0.40662495 | 53.9256141 | 5.45E-11 | 4.58E-10 |
| E41LA | 13.4946414 | 1.71377605 | 0.35633199 | 37.0211336 | 1.78E-07 | 1.06E-06 |
| MSD2 | 7.73664427 | 1.7134858 | 0.43655193 | 25.4816162 | 4.02E-05 | 0.00018007 |
| WFDC3 | 50.4486335 | 1.70659967 | 0.34273513 | 58.8111761 | 5.16E-12 | 4.67E-11 |
| TPH | 5.38513955 | 1.70522265 | 0.82044922 | 52.549154 | 1.06E-10 | 8.63E-10 |
| DMBT1.1 | 16.6581316 | 1.7026299 | 0.3926912 | 115.540605 | 4.78E-24 | 9.80E-23 |
| LEF1.1 | 156.837194 | 1.69724155 | 0.13511145 | 887.682577 | 7.77E-191 | 2.38E-187 |
| SBSPO | 10.230309 | 1.69668693 | 0.34828005 | 28.477679 | 9.98E-06 | 4.87E-05 |
| NELL1 | 11.5669454 | 1.69361027 | 0.36438012 | 29.0231599 | 7.73E-06 | 3.83E-05 |
| CLIC6 | 26.5092453 | 1.69215639 | 0.2261143 | 81.6250105 | 7.88E-17 | 1.04E-15 |
| KCMB4 | 4.9008207 | 1.69152752 | 0.66187744 | 24.7415441 | 5.67E-05 | 0.00024767 |
| NRAP | 6.86507337 | 1.69134195 | 0.45256892 | 23.1939835 | 0.00011581 | 0.00048075 |
| FHL1 | 126.880886 | 1.6903782 | 0.16232712 | 202.263992 | 1.22E-42 | 6.65E-41 |
| LRC4B | 22.5083641 | 1.68949879 | 0.30760896 | 93.1272547 | 2.85E-19 | 4.46E-18 |
| TNR19 | 593.483129 | 1.6890744 | 0.12407711 | 462.462557 | 8.78E-99 | 3.36E-96 |
| CAHM5 | 50.4269224 | 1.68829056 | 0.27449073 | 75.5313116 | 1.54E-15 | 1.84E-14 |
| LAMB1 | 11.0868695 | 1.6781405 | 0.53438436 | 25.2674645 | 4.44E-05 | 0.00019747 |
| KCNJ5 | 19.5565665 | 1.67573 | 0.30807506 | 62.8353077 | 7.35E-13 | 7.06E-12 |
| COF2.2 | 41.9690014 | 1.65417656 | 0.267162 | 54.7507004 | 3.66E-11 | 3.13E-10 |
| CNFN.1 | 11.9693743 | 1.65183283 | 0.5937457 | 40.7146036 | 3.08E-08 | 2.00E-07 |
| MYCN.1 | 50.155045 | 1.64682928 | 0.22424258 | 80.3317825 | 1.48E-16 | 1.91E-15 |
| FOXA2 | 58.1212446 | 1.64327011 | 0.41097384 | 344.511765 | 2.68E-73 | 6.74E-71 |
| DRC8 | 11.8237779 | 1.64030734 | 0.3893517 | 47.1820744 | 1.40E-09 | 1.03E-08 |
| PTC1 | 32.5229743 | 1.6364932 | 0.21028723 | 124.186334 | 6.81E-26 | 1.53E-24 |
| ITB1.1 | 11.883621 | 1.63585825 | 0.44507247 | 28.1012643 | 1.19E-05 | 5.73E-05 |
| IL8.2 | 66.9287111 | 1.63515208 | 0.40794984 | 53.0059969 | 8.50E-11 | 7.00E-10 |
| KOZA | 11.33261 | 1.63444525 | 0.69869498 | 66.7567461 | 1.10E-13 | 1.12E-12 |
| TFPI2 | 25.3254543 | 1.63143971 | 0.26566127 | 50.7061574 | 2.57E-10 | 2.04E-09 |
| MPZL2 | 5.62267464 | 1.62826652 | 0.7031437 | 25.8963947 | 3.32E-05 | 0.00015039 |
| COE3.1 | 19.2908634 | 1.62511488 | 0.25651567 | 43.3900374 | 8.59E-09 | 5.85E-08 |
| PAQR3 | 7.09946718 | 1.62347709 | 0.47402511 | 38.8590632 | 7.45E-08 | 4.63E-07 |
| GRM7 | 18.5669729 | 1.62126504 | 0.28091785 | 42.0683651 | 1.61E-08 | 1.08E-07 |
| TMEM9 | 48.9719165 | 1.61249148 | 0.15396007 | 117.036297 | 2.29E-24 | 4.79E-23 |
| IGS21 | 17.2598026 | 1.61031466 | 0.52635093 | 44.6621724 | 4.67E-09 | 3.28E-08 |
| KALM | 18.7374786 | 1.60683878 | 0.29455195 | 71.3995434 | 1.15E-14 | 1.29E-13 |
| CO4A2.1 | 520.15193 | 1.60669168 | 0.28986613 | 111.812012 | 2.99E-23 | 5.86E-22 |
| FGF9 | 11.8765284 | 1.60542221 | 0.37595809 | 39.0093876 | 6.94E-08 | 4.33E-07 |
| RBM38.1 | 13.0193471 | 1.60368263 | 0.39806923 | 47.0958186 | 1.46E-09 | 1.07E-08 |
| TKN1 | 34.7005047 | 1.59700092 | 0.40159395 | 201.90907 | 1.46E-42 | 7.87E-41 |
| CAH7 | 73.9750021 | 1.59683896 | 0.19577422 | 248.064923 | 1.70E-52 | 1.71E-50 |
| AAKB2 | 8.85541864 | 1.59452062 | 0.41016269 | 21.9637602 | 0.00020378 | 0.00081452 |
| XCT.1 | 15.5451131 | 1.59359505 | 0.30823612 | 55.5227391 | 2.52E-11 | 2.19E-10 |
| ATS18 | 42.9619532 | 1.59347905 | 0.21528446 | 153.278546 | 4.04E-32 | 1.24E-30 |
| CAC1H.1 | 7.65386836 | 1.59191395 | 0.43893466 | 27.9336128 | 1.29E-05 | 6.18E-05 |
| SYT17 | 5.93825565 | 1.5897235 | 0.5036277 | 23.7355754 | 9.02E-05 | 0.00038089 |
| ACE | 30.3732651 | 1.58929537 | 0.22669742 | 70.8780801 | 1.48E-14 | 1.64E-13 |
| NRSN1 | 18.5395995 | 1.5839929 | 0.41532114 | 49.5240445 | 4.54E-10 | 3.53E-09 |
| GLDN | 258.537565 | 1.58096893 | 0.21292045 | 78.2950364 | 4.00E-16 | 5.00E-15 |
| AT1A2 | 22.4410497 | 1.58091508 | 0.3850882 | 69.4143475 | 3.02E-14 | 3.25E-13 |
| BMP7.2 | 41.3812412 | 1.58073587 | 0.35133556 | 137.879847 | 8.03E-29 | 2.13E-27 |
| RAMP1.1 | 29.3405688 | 1.58051842 | 0.2517303 | 48.83221 | 6.33E-10 | 4.85E-09 |
| COL10 | 59.6161273 | 1.57472799 | 0.31634276 | 79.5172458 | 2.20E-16 | 2.81E-15 |
| EYA4 | 58.9774189 | 1.5696398 | 0.28834661 | 199.439496 | 4.96E-42 | 2.63E-40 |
| HS12A | 25.8747811 | 1.56938577 | 0.25678893 | 134.140348 | 5.07E-28 | 1.28E-26 |
| ATS9 | 50.6508912 | 1.5676641 | 0.24505661 | 123.209252 | 1.10E-25 | 2.45E-24 |
| FOXC2 | 31.2406555 | 1.56530037 | 0.27061689 | 62.9000919 | 7.12E-13 | 6.85E-12 |
| CHAD | 28.1527379 | 1.56460672 | 0.22069399 | 55.4893337 | 2.57E-11 | 2.23E-10 |
| EFNA3 | 6.19719472 | 1.55944523 | 0.58109561 | 28.8457295 | 8.40E-06 | 4.13E-05 |
| SMOC1.1 | 36.17042 | 1.55514819 | 0.18378475 | 98.0196376 | 2.60E-20 | 4.27E-19 |
| ESM1 | 205.384048 | 1.5533694 | 0.16121674 | 139.740643 | 3.21E-29 | 8.66E-28 |
| VGFR2 | 16.7627621 | 1.54381257 | 0.27550762 | 36.1233436 | 2.73E-07 | 1.60E-06 |
| ALKL2 | 8.53396981 | 1.54274036 | 0.52733332 | 31.0932089 | 2.93E-06 | 1.52E-05 |
| S4A4.1 | 44.4594919 | 1.53963581 | 0.31269304 | 98.4028777 | 2.15E-20 | 3.55E-19 |
| Z385D | 12.3311649 | 1.53751829 | 0.3114129 | 26.4576035 | 2.56E-05 | 0.00011829 |
| TICN3 | 20.7918658 | 1.53616277 | 0.29642855 | 31.8084594 | 2.09E-06 | 1.10E-05 |
| LPAR6 | 14.9605582 | 1.53143112 | 0.83732455 | 49.7641151 | 4.04E-10 | 3.15E-09 |
| VEGFA.1 | 29.3493051 | 1.52995444 | 0.28251367 | 47.8882752 | 9.96E-10 | 7.49E-09 |
| VWC2L | 21.5760635 | 1.52669907 | 0.27629166 | 64.3756003 | 3.48E-13 | 3.42E-12 |
| CADH7.1 | 10.2538588 | 1.52297742 | 0.34476067 | 21.8796097 | 0.00021179 | 0.00084478 |
| SGK1 | 349.466932 | 1.52285728 | 0.12670119 | 498.175676 | 1.66E-106 | 8.48E-104 |
| SIX2 | 33.9305226 | 1.51572896 | 0.453304 | 162.285506 | 4.73E-34 | 1.65E-32 |
| ADML | 132.460094 | 1.50951649 | 0.24147201 | 160.783281 | 9.93E-34 | 3.39E-32 |
| SIA4C | 9.5052951 | 1.50812512 | 0.39288939 | 31.4992689 | 2.42E-06 | 1.27E-05 |
| ZN227.15 | 11.40986 | 1.50791215 | 0.51506409 | 36.8805968 | 1.91E-07 | 1.13E-06 |
| TIE2 | 26.0709605 | 1.50585725 | 0.22925887 | 45.7763608 | 2.74E-09 | 1.97E-08 |
| HSPB7 | 9.58431584 | 1.50447969 | 0.64020495 | 27.4431626 | 1.62E-05 | 7.67E-05 |
| HHIP | 66.6565385 | 1.50256765 | 0.15383662 | 191.139529 | 3.02E-40 | 1.42E-38 |
| GLI2 | 82.1018521 | 1.50087668 | 0.1889893 | 228.308807 | 3.05E-48 | 2.34E-46 |
| PTH1R | 12.4339856 | 1.49967822 | 0.43685797 | 96.3420033 | 5.91E-20 | 9.58E-19 |
| SCUB1 | 40.9259174 | 1.4986456 | 0.18823121 | 156.179413 | 9.64E-33 | 3.11E-31 |
| GLYC | 9.34511626 | 1.49592537 | 0.41854171 | 27.6672289 | 1.46E-05 | 6.94E-05 |
| PTPRD.1 | 35.6526559 | 1.49583182 | 0.27471422 | 74.1855331 | 2.96E-15 | 3.48E-14 |
| FBLN1 | 288.959886 | 1.49051569 | 0.18834813 | 103.942546 | 1.42E-21 | 2.50E-20 |
| AT8A2 | 55.7257374 | 1.49024964 | 0.27711003 | 91.6159349 | 5.97E-19 | 9.12E-18 |
| T132A | 18.9970178 | 1.48891479 | 0.34906416 | 40.5067194 | 3.40E-08 | 2.20E-07 |
| TGM2.3 | 17.9050847 | 1.48811082 | 0.28434367 | 31.3905397 | 2.55E-06 | 1.33E-05 |
| MGP | 1052.71746 | 1.4839078 | 0.28689994 | 50.0078971 | 3.60E-10 | 2.82E-09 |
| HOX71 | 351.634921 | 1.48076065 | 0.1878111 | 379.803642 | 6.42E-81 | 2.01E-78 |
| DMA | 66.5546889 | 1.47951206 | 0.16682748 | 90.9320222 | 8.35E-19 | 1.27E-17 |
| DMTA2 | 22.5863884 | 1.47765015 | 0.49186657 | 260.890815 | 2.93E-55 | 3.27E-53 |
| ENPP2 | 537.870496 | 1.47754352 | 0.26290939 | 80.0370087 | 1.71E-16 | 2.20E-15 |
| LAMP5 | 6.13773389 | 1.47710745 | 0.64818377 | 24.1019062 | 7.62E-05 | 0.0003256 |
| SLIT3 | 135.40353 | 1.47636575 | 0.14943918 | 396.182283 | 1.86E-84 | 5.93E-82 |
| MET24 | 25.4321652 | 1.47615274 | 0.28153651 | 60.2369201 | 2.59E-12 | 2.39E-11 |
| EGFL6 | 55.6859927 | 1.46960592 | 0.16406477 | 134.436286 | 4.38E-28 | 1.11E-26 |
| CLD5 | 13.492664 | 1.46805903 | 0.33960507 | 22.7709626 | 0.00014069 | 0.00057805 |
| RSPO1 | 110.187553 | 1.46494311 | 0.25109488 | 160.764447 | 1.00E-33 | 3.42E-32 |
| TRI46 | 16.2827802 | 1.46482419 | 0.4404459 | 24.8131691 | 5.49E-05 | 0.00024001 |
| ENDD1.2 | 684.637315 | 1.4550412 | 0.25276647 | 185.02045 | 6.23E-39 | 2.80E-37 |
| FBLN7.1 | 24.5174331 | 1.45220963 | 0.37561368 | 47.7147223 | 1.08E-09 | 8.10E-09 |
| B3GA1 | 33.8728876 | 1.44982903 | 0.32009764 | 93.1279023 | 2.85E-19 | 4.46E-18 |
| S6A15 | 19.288755 | 1.44811527 | 0.30793698 | 59.6739403 | 3.40E-12 | 3.12E-11 |
| C1QC | 1381.23037 | 1.4471108 | 0.23041279 | 162.50761 | 4.24E-34 | 1.48E-32 |
| EGFL8 | 86.3725162 | 1.44379757 | 0.14833209 | 130.604456 | 2.89E-27 | 7.09E-26 |
| ANXA6 | 25.8087232 | 1.43904107 | 0.2926684 | 56.8705336 | 1.32E-11 | 1.16E-10 |
| WN10B | 18.5871197 | 1.43492912 | 0.38002982 | 68.3606997 | 5.03E-14 | 5.33E-13 |
| MAML3 | 49.0198766 | 1.43450273 | 0.1761849 | 98.7047224 | 1.86E-20 | 3.07E-19 |
| PGM5 | 25.2608296 | 1.43442017 | 0.26605865 | 30.8928603 | 3.22E-06 | 1.66E-05 |
| PODXL | 14.9854474 | 1.43364506 | 0.30346573 | 23.7435023 | 8.99E-05 | 0.00037992 |
| LMTK1 | 74.8703955 | 1.4323924 | 0.22939597 | 110.874713 | 4.74E-23 | 9.16E-22 |
| F180A.1 | 25.7542049 | 1.43177529 | 0.62458735 | 43.629787 | 7.66E-09 | 5.26E-08 |
| FLNC | 48.7195035 | 1.43096823 | 0.25079016 | 39.2433578 | 6.20E-08 | 3.88E-07 |
| CSPG4 | 22.3480844 | 1.4268192 | 0.27598114 | 61.473601 | 1.42E-12 | 1.34E-11 |
| FRAS1 | 138.607818 | 1.42389791 | 0.15485259 | 172.15369 | 3.61E-36 | 1.38E-34 |
| OBSL1.2 | 21.5832074 | 1.42319585 | 0.29267886 | 35.7125425 | 3.32E-07 | 1.92E-06 |
| NSG1.1 | 6.86980203 | 1.42232744 | 0.52451604 | 23.9161037 | 8.30E-05 | 0.00035289 |
| MAP9.1 | 12.3403339 | 1.41874797 | 0.46333797 | 29.6180042 | 5.85E-06 | 2.94E-05 |
| PKHH1 | 7.53638889 | 1.41317984 | 0.64746884 | 22.3100651 | 0.00017385 | 0.00070226 |
| LAMA2 | 251.884074 | 1.41077813 | 0.11887973 | 241.629245 | 4.14E-51 | 3.91E-49 |
| TMM79.2 | 5.82606358 | 1.40841005 | 0.50102729 | 21.6408696 | 0.00023625 | 0.00093506 |
| ANFD | 9.40957755 | 1.40826859 | 0.56026855 | 25.4089353 | 4.16E-05 | 0.00018602 |
| RASD1 | 10.7574491 | 1.4001254 | 0.36386904 | 22.1752794 | 0.00018494 | 0.0007441 |
| S4A11 | 9.19008597 | 1.39729917 | 0.41242251 | 27.0061127 | 1.98E-05 | 9.30E-05 |
| RAIN | 23.4848672 | 1.39632953 | 0.24410004 | 35.7532895 | 3.25E-07 | 1.89E-06 |
| FHL2 | 43.9630468 | 1.39383506 | 0.38150206 | 36.0208959 | 2.87E-07 | 1.67E-06 |
| FOXL2 | 3.11060349 | 1.39066711 | 0.99331448 | 25.5329118 | 3.93E-05 | 0.00017625 |
| AATC.1 | 28.1938575 | 1.38696111 | 0.20771515 | 76.9268693 | 7.79E-16 | 9.53E-15 |
| LIMA1 | 18.9137212 | 1.38278757 | 0.35939574 | 26.782099 | 2.20E-05 | 0.00010269 |
| ANGL7 | 11.0522536 | 1.38064973 | 0.42948031 | 22.7346335 | 0.00014306 | 0.00058716 |
| TIAM2 | 29.0779625 | 1.37952527 | 0.23335098 | 141.147368 | 1.60E-29 | 4.41E-28 |
| HXA3 | 7.36232529 | 1.37941297 | 0.91058571 | 96.3389354 | 5.92E-20 | 9.58E-19 |
| LIN1.40 | 11.3260325 | 1.37749683 | 0.49246157 | 35.156997 | 4.31E-07 | 2.47E-06 |
| DKK4 | 8.74529863 | 1.37653802 | 0.47042363 | 26.182746 | 2.91E-05 | 0.00013321 |
| ANKHB | 21.2759978 | 1.37572021 | 0.27911895 | 67.6452384 | 7.13E-14 | 7.40E-13 |
| SP6 | 244.624473 | 1.3746849 | 0.23414774 | 184.148258 | 9.58E-39 | 4.24E-37 |
| GPR54 | 6.18309868 | 1.37371285 | 0.64858052 | 25.7648121 | 3.53E-05 | 0.00015925 |
| LYL1 | 15.4170757 | 1.36515565 | 0.28858954 | 26.2670958 | 2.80E-05 | 0.00012851 |
| SIAT1 | 15.5409612 | 1.35749341 | 0.32314304 | 48.1154722 | 8.93E-10 | 6.74E-09 |
| CODA1 | 63.0884413 | 1.35301069 | 0.19470535 | 107.437673 | 2.56E-22 | 4.72E-21 |
| PDGFB | 19.2626288 | 1.3518894 | 0.25787031 | 29.7119361 | 5.60E-06 | 2.82E-05 |
| TINAL | 49.5313041 | 1.35091113 | 0.16904528 | 70.2276506 | 2.03E-14 | 2.22E-13 |
| NFIL3 | 34.5625374 | 1.34418057 | 0.18993738 | 95.7633475 | 7.84E-20 | 1.26E-18 |
| AQP1 | 46.8862667 | 1.33331873 | 0.205348 | 45.3771992 | 3.32E-09 | 2.36E-08 |
| CRIM1 | 76.2285676 | 1.33093406 | 0.15648073 | 145.400554 | 1.97E-30 | 5.64E-29 |
| VGFR1 | 28.7571268 | 1.32901057 | 0.30348086 | 38.1162746 | 1.06E-07 | 6.47E-07 |
| TM38A | 33.9466155 | 1.32610902 | 0.26722401 | 56.1859788 | 1.83E-11 | 1.60E-10 |
| C1QR1 | 77.0597954 | 1.32085346 | 0.26128205 | 92.9618447 | 3.09E-19 | 4.83E-18 |
| CYTL1 | 256.129031 | 1.31862523 | 0.41197389 | 134.435406 | 4.38E-28 | 1.11E-26 |
| HMGA1 | 6.71452027 | 1.31666419 | 0.55786791 | 26.7349759 | 2.25E-05 | 0.00010471 |
| SCUB3 | 87.3182386 | 1.3133306 | 0.21786574 | 223.176647 | 3.88E-47 | 2.75E-45 |
| CD151 | 958.557537 | 1.31303327 | 0.2024973 | 57.4245362 | 1.01E-11 | 8.97E-11 |
| PCD10.5 | 37.6954989 | 1.30958066 | 0.48875712 | 33.9203241 | 7.74E-07 | 4.32E-06 |
| GDAP1 | 19.779575 | 1.3046236 | 0.26672979 | 39.1261641 | 6.56E-08 | 4.10E-07 |
| ADML.1 | 96.2298556 | 1.30309011 | 0.28284213 | 38.072705 | 1.08E-07 | 6.60E-07 |
| EPHA7 | 31.6347777 | 1.30170336 | 0.30137827 | 69.3920401 | 3.05E-14 | 3.29E-13 |
| LFNG | 132.708917 | 1.29859713 | 0.1189356 | 261.667254 | 1.99E-55 | 2.26E-53 |
| PYGM | 54.4327031 | 1.29787588 | 0.2078019 | 49.4460876 | 4.71E-10 | 3.66E-09 |
| POPD1 | 10.0914037 | 1.29649144 | 0.41535031 | 23.782199 | 8.83E-05 | 0.00037402 |
| LORF2.167 | 15.9150078 | 1.29415193 | 0.37104642 | 33.5183063 | 9.35E-07 | 5.16E-06 |
| LGR6 | 89.516891 | 1.29381444 | 0.18907986 | 84.9728619 | 1.54E-17 | 2.13E-16 |
| INHBB | 28.8563127 | 1.29204563 | 0.40703785 | 103.629178 | 1.66E-21 | 2.91E-20 |
| FLNC.1 | 18.5074556 | 1.29129674 | 0.26871704 | 25.1774918 | 4.63E-05 | 0.00020534 |
| KANK1 | 78.5766626 | 1.2911021 | 0.2312753 | 42.4730909 | 1.33E-08 | 8.91E-08 |
| CCN1 | 9.74113074 | 1.29052662 | 0.66746414 | 22.3874141 | 0.00016779 | 0.00067974 |
| TUTLA | 223.564308 | 1.29037356 | 0.27489824 | 46.6027084 | 1.84E-09 | 1.35E-08 |
| C1QA | 676.581996 | 1.28644343 | 0.2206182 | 114.352695 | 8.58E-24 | 1.73E-22 |
| MEG10.1 | 39.022376 | 1.28520588 | 0.30195483 | 44.8477458 | 4.28E-09 | 3.01E-08 |
| RGCC | 68.7941069 | 1.2822818 | 0.24709875 | 42.8745002 | 1.10E-08 | 7.42E-08 |
| EPHA4 | 51.8721824 | 1.27864233 | 0.19752377 | 115.6929 | 4.44E-24 | 9.13E-23 |
| RAD | 12.0363521 | 1.27299631 | 0.6130293 | 24.6362634 | 5.95E-05 | 0.00025906 |
| CXG1 | 32.3005249 | 1.27276704 | 0.2637331 | 146.094651 | 1.40E-30 | 4.06E-29 |
| LGI1 | 82.0844158 | 1.26879518 | 0.21373789 | 50.1016757 | 3.44E-10 | 2.70E-09 |
| GDIR2 | 12.0510167 | 1.2679671 | 0.40827097 | 29.3194343 | 6.73E-06 | 3.36E-05 |
| VEGFC | 27.4410808 | 1.26694898 | 0.24932584 | 70.4504168 | 1.82E-14 | 2.00E-13 |
| CA216 | 21.8785214 | 1.26058664 | 0.30535603 | 105.11227 | 8.02E-22 | 1.43E-20 |
| FGF10 | 12.954586 | 1.25766206 | 0.33653982 | 45.3940283 | 3.29E-09 | 2.34E-08 |
| LAMA1 | 10.1584116 | 1.25749167 | 0.43971028 | 43.4954896 | 8.17E-09 | 5.58E-08 |
| IDHP | 280.573245 | 1.25656882 | 0.13734698 | 262.776639 | 1.15E-55 | 1.34E-53 |
| A4GAT | 6.48402001 | 1.25495615 | 0.55195656 | 22.4300591 | 0.00016454 | 0.00066815 |
| TEN2 | 16.6695644 | 1.25236855 | 0.32153129 | 47.1552724 | 1.42E-09 | 1.05E-08 |
| MSH4 | 8.10124589 | 1.24997523 | 0.4781774 | 34.4038089 | 6.16E-07 | 3.48E-06 |
| CUBN.3 | 11.895837 | 1.24516027 | 0.39879108 | 39.5786532 | 5.29E-08 | 3.34E-07 |
| VATB | 28.4065471 | 1.24442285 | 0.34396936 | 23.5434489 | 9.86E-05 | 0.00041399 |
| TRBM | 95.4363041 | 1.24246468 | 0.14095863 | 116.563623 | 2.89E-24 | 6.01E-23 |
| CALD1 | 52.3214203 | 1.240828 | 0.20788002 | 90.2819187 | 1.15E-18 | 1.72E-17 |
| NKD1 | 40.3816456 | 1.23653239 | 0.22793755 | 50.8854714 | 2.36E-10 | 1.88E-09 |
| OLM2B | 80.2523385 | 1.23597697 | 0.22474894 | 111.456566 | 3.56E-23 | 6.90E-22 |
| KCMB3 | 18.4016354 | 1.2358443 | 0.27212412 | 27.1625691 | 1.84E-05 | 8.68E-05 |
| SPP24.2 | 160.955001 | 1.23126391 | 0.55549801 | 104.672766 | 9.95E-22 | 1.76E-20 |
| SAM10 | 15.3940335 | 1.22165496 | 0.30941218 | 23.0757342 | 0.00012229 | 0.00050572 |
| EFCC1 | 30.5989486 | 1.22147353 | 0.22433233 | 129.876884 | 4.14E-27 | 1.01E-25 |
| CADH5 | 26.1526481 | 1.22034512 | 0.22846728 | 36.734057 | 2.04E-07 | 1.21E-06 |
| NID1 | 92.8779778 | 1.21900644 | 0.13276507 | 180.088662 | 7.14E-38 | 2.94E-36 |
| HIC1 | 26.4917227 | 1.21873967 | 0.22605192 | 34.1662075 | 6.89E-07 | 3.87E-06 |
| ID4 | 147.913986 | 1.21860188 | 0.15405877 | 126.377975 | 2.32E-26 | 5.37E-25 |
| VIPR2 | 6.56048085 | 1.21848765 | 0.7257428 | 40.1469031 | 4.04E-08 | 2.58E-07 |
| RBP | 6.5382775 | 1.21639923 | 0.73505694 | 29.572936 | 5.98E-06 | 3.00E-05 |
| TCF15.1 | 3.4770423 | 1.21539772 | 1.02064625 | 24.5334198 | 6.24E-05 | 0.00027053 |
| GRIP1.1 | 13.1212364 | 1.2137823 | 0.34629625 | 30.6438268 | 3.62E-06 | 1.86E-05 |
| CDHR3.1 | 3.83097947 | 1.2117336 | 0.98623778 | 35.0190081 | 4.60E-07 | 2.63E-06 |
| CD276.2 | 10.2262722 | 1.21084128 | 0.5242443 | 42.0177437 | 1.65E-08 | 1.10E-07 |
| TBX3 | 128.262976 | 1.2106111 | 0.13596395 | 311.575881 | 3.45E-66 | 6.43E-64 |
| HOME2 | 15.4656187 | 1.20212814 | 0.32426541 | 34.3407872 | 6.34E-07 | 3.58E-06 |
| LAMC3 | 304.549098 | 1.20021713 | 0.15553926 | 88.2985141 | 3.03E-18 | 4.38E-17 |
| LIN1.88 | 6.03589171 | 1.20009693 | 0.63664622 | 32.7921647 | 1.32E-06 | 7.12E-06 |
| BMP2.1 | 48.0008637 | 1.19958321 | 0.24487292 | 300.575367 | 8.14E-64 | 1.38E-61 |
| ASB17 | 17.4568484 | 1.19857208 | 0.39083895 | 22.129099 | 0.0001889 | 0.00075863 |
| PCD18.1 | 20.8933502 | 1.19662542 | 0.25418843 | 37.39374 | 1.49E-07 | 8.95E-07 |
| PTPRB | 31.2896985 | 1.19227479 | 0.22144384 | 45.7783104 | 2.74E-09 | 1.96E-08 |
| CA3A1 | 22.65209 | 1.187862 | 0.239316 | 32.8219391 | 1.30E-06 | 7.04E-06 |
| ACTA | 62.1866475 | 1.18667189 | 0.26626063 | 35.4334136 | 3.78E-07 | 2.18E-06 |
| THIO | 17.7664585 | 1.18530998 | 0.26650137 | 32.7522025 | 1.34E-06 | 7.25E-06 |
| AL1A2 | 29.7663494 | 1.18471075 | 0.33019458 | 40.8299692 | 2.91E-08 | 1.90E-07 |
| FXD3A | 36.4465908 | 1.18368058 | 0.1929076 | 41.0045179 | 2.68E-08 | 1.75E-07 |
| SESD1 | 104.37282 | 1.17737938 | 0.13042088 | 207.100465 | 1.12E-43 | 6.33E-42 |
| SEM3D | 79.0122307 | 1.17111634 | 0.15496959 | 121.969867 | 2.03E-25 | 4.45E-24 |
| IBP3 | 519.935074 | 1.17026203 | 0.10913489 | 161.150789 | 8.28E-34 | 2.85E-32 |
| RSLBB | 28.8264479 | 1.16976279 | 0.22943442 | 35.0994708 | 4.43E-07 | 2.54E-06 |
| KCNN3 | 18.4530581 | 1.16841829 | 0.29217719 | 24.7105845 | 5.75E-05 | 0.0002511 |
| MMRN1 | 67.2638991 | 1.16647961 | 0.15385337 | 77.2636828 | 6.61E-16 | 8.12E-15 |
| PA2A | 126.84492 | 1.16465799 | 0.52296104 | 38.9449091 | 7.15E-08 | 4.45E-07 |
| TENA.1 | 228.056834 | 1.1639118 | 0.19759929 | 212.340518 | 8.34E-45 | 5.04E-43 |
| HIPL1 | 12.3712684 | 1.16282007 | 0.50591762 | 88.3495575 | 2.95E-18 | 4.28E-17 |
| NICN1 | 18.4265892 | 1.16140756 | 0.27608448 | 44.9026742 | 4.17E-09 | 2.94E-08 |
| ISM1 | 126.485326 | 1.1609235 | 0.21070717 | 50.0448029 | 3.53E-10 | 2.77E-09 |
| N2F1A | 26.9105555 | 1.16063227 | 0.26037892 | 110.014121 | 7.23E-23 | 1.38E-21 |
| CP1A1 | 149.34593 | 1.15886742 | 0.24120355 | 100.503264 | 7.69E-21 | 1.30E-19 |
| PLPP3 | 233.391434 | 1.15865119 | 0.22614663 | 34.638028 | 5.51E-07 | 3.13E-06 |
| TOX | 14.2174706 | 1.15806375 | 0.31667112 | 46.4333698 | 2.00E-09 | 1.45E-08 |
| PPIC | 119.852538 | 1.15754065 | 0.20488552 | 109.752415 | 8.22E-23 | 1.57E-21 |
| ECE2 | 59.331568 | 1.15013494 | 0.17902381 | 118.615188 | 1.06E-24 | 2.23E-23 |
| NANO3 | 12.2935368 | 1.14641516 | 0.51796035 | 40.0343608 | 4.26E-08 | 2.71E-07 |
| SC6A1 | 119.555349 | 1.14608033 | 0.14556103 | 78.5286538 | 3.57E-16 | 4.48E-15 |
| PITX2 | 260.763042 | 1.14522679 | 0.12661562 | 472.989195 | 4.65E-101 | 1.98E-98 |
| NUA4L | 1000.23316 | 1.14370871 | 0.18964085 | 111.489297 | 3.50E-23 | 6.81E-22 |
| AS3MT | 30.2031556 | 1.14098698 | 0.22017745 | 30.6486662 | 3.61E-06 | 1.85E-05 |
| TEX9 | 23.0822876 | 1.13841749 | 0.26532252 | 32.7176337 | 1.36E-06 | 7.35E-06 |
| NET4 | 6.92286051 | 1.13831486 | 0.51039313 | 22.6830786 | 0.00014649 | 0.00060043 |
| SATB1 | 28.9781317 | 1.13822342 | 0.24679343 | 36.7407435 | 2.04E-07 | 1.21E-06 |
| PMP22 | 911.338499 | 1.13271686 | 0.18274627 | 47.5755934 | 1.16E-09 | 8.63E-09 |
| LARG1 | 38.5945476 | 1.13223443 | 0.21412925 | 33.9807343 | 7.52E-07 | 4.21E-06 |
| SPY4 | 12.7947571 | 1.13212687 | 0.32632978 | 28.3653376 | 1.05E-05 | 5.12E-05 |
| DLX4 | 10.7761342 | 1.13175813 | 0.44016886 | 40.7867181 | 2.98E-08 | 1.93E-07 |
| NOTUM | 113.031585 | 1.13052888 | 0.16189072 | 86.9449213 | 5.86E-18 | 8.33E-17 |
| CANX.1 | 25.5852961 | 1.12946796 | 0.33066138 | 26.4370706 | 2.58E-05 | 0.00011925 |
| LOXL4 | 158.907267 | 1.12877666 | 0.28794072 | 161.48342 | 7.03E-34 | 2.43E-32 |
| HS3S1 | 238.978864 | 1.1284344 | 0.10268317 | 187.44952 | 1.87E-39 | 8.50E-38 |
| Z585A | 24.5959633 | 1.12799523 | 0.25534106 | 28.3667917 | 1.05E-05 | 5.11E-05 |
| EGFLA | 38.6894245 | 1.12567804 | 0.19941126 | 94.3014181 | 1.60E-19 | 2.53E-18 |
| AGRL2 | 74.5395683 | 1.12135727 | 0.17465508 | 196.249842 | 2.40E-41 | 1.24E-39 |
| GFRA2 | 48.6648813 | 1.11835851 | 0.19058773 | 67.231798 | 8.71E-14 | 8.95E-13 |
| SMYD3 | 18.316745 | 1.11716673 | 0.3104037 | 34.9937584 | 4.66E-07 | 2.66E-06 |
| S6A20.1 | 28.5602957 | 1.11184528 | 0.24649093 | 52.4371935 | 1.12E-10 | 9.10E-10 |
| NEP | 14.2476714 | 1.11115792 | 0.33418569 | 30.1134172 | 4.64E-06 | 2.35E-05 |
| SULF1 | 54.7153641 | 1.10997484 | 0.18501146 | 218.971495 | 3.12E-46 | 2.08E-44 |
| MTAP2.1 | 39.9044857 | 1.1090914 | 0.1873699 | 104.916774 | 8.82E-22 | 1.57E-20 |
| EDNRB | 60.5130059 | 1.10679538 | 0.16385033 | 67.1014222 | 9.28E-14 | 9.51E-13 |
| SOX4 | 94.5852769 | 1.10508484 | 0.17183754 | 142.823455 | 7.02E-30 | 1.95E-28 |
| CO8A1 | 51.2970612 | 1.10289671 | 0.49862721 | 73.2795058 | 4.60E-15 | 5.30E-14 |
| DYN3 | 10.2815397 | 1.10242419 | 0.43594424 | 44.1990237 | 5.83E-09 | 4.06E-08 |
| FREM1.4 | 53.3834134 | 1.10209384 | 0.20401341 | 126.848664 | 1.84E-26 | 4.29E-25 |
| SDK2 | 16.5790377 | 1.09997381 | 0.28195127 | 27.5302895 | 1.55E-05 | 7.39E-05 |
| AF9 | 9.3869587 | 1.09632677 | 0.51637778 | 24.829475 | 5.44E-05 | 0.00023841 |
| WN10A | 80.423589 | 1.09443577 | 0.14913317 | 170.558412 | 7.94E-36 | 2.98E-34 |
| ENDOU | 232.001604 | 1.09270818 | 0.12729216 | 314.377796 | 8.57E-67 | 1.64E-64 |
| SPG16 | 3.09827564 | 1.09234294 | 1.04461582 | 23.7715285 | 8.88E-05 | 0.00037555 |
| WBP2.1 | 23.8943082 | 1.09208643 | 0.25570173 | 46.3230699 | 2.11E-09 | 1.53E-08 |
| CRLD1 | 63.6463437 | 1.09190622 | 0.42850175 | 57.2033999 | 1.12E-11 | 9.94E-11 |
| COAA1.3 | 3057.41382 | 1.08817315 | 0.4874718 | 183.386249 | 1.40E-38 | 6.09E-37 |
| BAMBI | 259.99377 | 1.08652165 | 0.17911597 | 323.857267 | 7.71E-69 | 1.62E-66 |
| SVEP1.2 | 18.7701928 | 1.08486411 | 0.2837306 | 35.3293382 | 3.98E-07 | 2.29E-06 |
| APBA2 | 15.1676621 | 1.08447452 | 0.31093985 | 23.3836858 | 0.00010613 | 0.00044294 |
| PTX3 | 16.2004303 | 1.07379512 | 0.31382884 | 29.8926073 | 5.15E-06 | 2.60E-05 |
| GAS6 | 222.602084 | 1.07236459 | 0.15521615 | 81.8328069 | 7.12E-17 | 9.42E-16 |
| LHPL6 | 110.470436 | 1.07073108 | 0.11479191 | 158.254204 | 3.46E-33 | 1.15E-31 |
| SPINZ | 23.5292955 | 1.0702632 | 0.23255467 | 57.8303997 | 8.28E-12 | 7.40E-11 |
| PSD13.1 | 20.9296927 | 1.06957233 | 0.25853318 | 42.7728028 | 1.15E-08 | 7.77E-08 |
| SC6A8.1 | 31.9908085 | 1.06538024 | 0.20818237 | 61.6563407 | 1.30E-12 | 1.23E-11 |
| SMAD6 | 35.0528969 | 1.0651471 | 0.23082574 | 28.3103771 | 1.08E-05 | 5.24E-05 |
| EDNRA | 94.3354423 | 1.062751 | 0.14510218 | 257.161334 | 1.86E-54 | 2.01E-52 |
| HARB1.53 | 6.28660061 | 1.06205496 | 0.61609568 | 23.3151641 | 0.00010953 | 0.00045627 |
| MNS1 | 13.882329 | 1.05562216 | 0.32232487 | 25.270532 | 4.44E-05 | 0.00019725 |
| GMPR1 | 24.566495 | 1.05476748 | 0.23835631 | 51.8918051 | 1.45E-10 | 1.17E-09 |
| NXNL2 | 58.5077705 | 1.04949344 | 0.21077759 | 57.5962609 | 9.28E-12 | 8.26E-11 |
| CRIP2 | 285.474342 | 1.04851732 | 0.19334661 | 164.961857 | 1.26E-34 | 4.50E-33 |
| LAMB1.1 | 190.326304 | 1.04586983 | 0.13093082 | 98.5253584 | 2.03E-20 | 3.35E-19 |
| ANR50.1 | 87.1807381 | 1.04512081 | 0.12880948 | 145.057889 | 2.33E-30 | 6.66E-29 |
| CBARP | 39.0235343 | 1.04507973 | 0.19082279 | 33.2473462 | 1.06E-06 | 5.83E-06 |
| EMIL3 | 50.1127656 | 1.04492808 | 0.21581452 | 43.8526546 | 6.88E-09 | 4.75E-08 |
| CNTFR | 38.6442822 | 1.04386518 | 0.21805904 | 38.3065175 | 9.69E-08 | 5.94E-07 |
| PROF2 | 174.844044 | 1.04343508 | 0.11366703 | 191.218738 | 2.90E-40 | 1.38E-38 |
| EXT1 | 26.5386993 | 1.0420769 | 0.22481143 | 31.1410344 | 2.87E-06 | 1.49E-05 |
| SO2A1 | 66.4008073 | 1.03953582 | 0.25499227 | 87.281813 | 4.97E-18 | 7.10E-17 |
| HEY1.1 | 64.5668337 | 1.03828576 | 0.19237755 | 169.243362 | 1.52E-35 | 5.61E-34 |
| OSR2B | 77.5076889 | 1.03597393 | 0.20387118 | 73.2981762 | 4.56E-15 | 5.26E-14 |
| SH3R3 | 15.1216796 | 1.03575731 | 0.30091668 | 35.4582027 | 3.74E-07 | 2.16E-06 |
| SUCB1 | 21.0744739 | 1.03465272 | 0.28609688 | 77.6629783 | 5.44E-16 | 6.76E-15 |
| SYT1 | 9.4038603 | 1.03377387 | 0.3970461 | 21.6993701 | 0.00023001 | 0.00091176 |
| FBLN5 | 214.511817 | 1.03316337 | 0.22877289 | 46.064121 | 2.39E-09 | 1.72E-08 |
| CO4A2 | 278.908321 | 1.03186138 | 0.09910198 | 164.345852 | 1.71E-34 | 6.07E-33 |
| RET4 | 671.709764 | 1.03009163 | 0.18372313 | 40.0594564 | 4.21E-08 | 2.69E-07 |
| CD81 | 736.089832 | 1.02988393 | 0.16184826 | 58.712374 | 5.41E-12 | 4.90E-11 |
| FOXL2.1 | 110.772981 | 1.0291279 | 0.16407346 | 193.529802 | 9.24E-41 | 4.53E-39 |
| FZD8 | 39.4976235 | 1.02817692 | 0.18053142 | 46.5507885 | 1.89E-09 | 1.38E-08 |
| TBKB1 | 25.9770614 | 1.02698255 | 0.22208753 | 37.2562076 | 1.60E-07 | 9.53E-07 |
| FGRL1 | 26.6598961 | 1.02501392 | 0.25510243 | 57.4224613 | 1.01E-11 | 8.98E-11 |
| FSCN1 | 133.625733 | 1.02225168 | 0.14230965 | 193.129292 | 1.13E-40 | 5.47E-39 |
| FA49A | 24.9045567 | 1.02044802 | 0.25270131 | 87.4941032 | 4.48E-18 | 6.43E-17 |
| CRYM | 14.9522107 | 1.01909947 | 0.32524674 | 24.2122441 | 7.24E-05 | 0.0003109 |
| ATS3 | 11.7713133 | 1.01808381 | 0.37993694 | 40.1436639 | 4.04E-08 | 2.58E-07 |
| UN5BA | 93.3274623 | 1.0137028 | 0.16897355 | 238.748601 | 1.73E-50 | 1.57E-48 |
| SELPB | 44.0340557 | 1.01231256 | 0.2271097 | 121.58457 | 2.45E-25 | 5.33E-24 |
| INAM2 | 21.6791964 | 1.01226775 | 0.29565514 | 48.0937614 | 9.02E-10 | 6.81E-09 |
| F167A.1 | 124.522022 | 1.00586214 | 0.11005193 | 105.107762 | 8.03E-22 | 1.43E-20 |
| HBEGF | 65.9242127 | 1.00385335 | 0.15156595 | 92.0205219 | 4.90E-19 | 7.54E-18 |
| PCD10.17 | 229.478625 | 1.0027554 | 0.11894625 | 324.90529 | 4.58E-69 | 9.74E-67 |
| T132C | 38.2556204 | 1.00191333 | 0.23956586 | 39.7085785 | 4.97E-08 | 3.15E-07 |
| RAMP1 | 27.8466424 | 1.00063356 | 0.2560096 | 55.1015942 | 3.09E-11 | 2.66E-10 |
| SPDLY | 39.1809983 | 0.9990069 | 0.25161288 | 82.1714673 | 6.04E-17 | 8.03E-16 |
| NUDT4 | 66.8161334 | 0.99846366 | 0.14108991 | 82.4928699 | 5.16E-17 | 6.92E-16 |
| NRCAM | 21.6338597 | 0.99732854 | 0.25027353 | 23.5481677 | 9.84E-05 | 0.00041332 |
| RGF1B | 27.7519157 | 0.99478793 | 0.27533434 | 150.349363 | 1.71E-31 | 5.20E-30 |
| SL9A5.1 | 4.28633538 | 0.99414367 | 0.86930028 | 23.4062266 | 0.00010503 | 0.00043886 |
| DREB | 27.0868391 | 0.99392885 | 0.25713462 | 95.8179066 | 7.63E-20 | 1.23E-18 |
| ACHA9 | 4.34092857 | 0.99007403 | 1.05819349 | 27.323674 | 1.71E-05 | 8.08E-05 |
| CALX | 26.9914504 | 0.98960005 | 0.23183368 | 72.6527451 | 6.25E-15 | 7.10E-14 |
| WDR47 | 27.7456538 | 0.98911175 | 0.23797926 | 52.8140249 | 9.32E-11 | 7.63E-10 |
| PLXB1 | 26.0576724 | 0.98890052 | 0.24568627 | 45.7931366 | 2.72E-09 | 1.95E-08 |
| GRAK.5 | 6.43199524 | 0.98736304 | 0.68126486 | 30.8685543 | 3.26E-06 | 1.68E-05 |
| OLFM | 14.5777118 | 0.98697948 | 0.3696332 | 32.8185374 | 1.30E-06 | 7.05E-06 |
| VCAM1 | 10.4761866 | 0.98660389 | 0.41707419 | 23.0765856 | 0.00012224 | 0.00050565 |
| ERG1 | 33.3408623 | 0.98635467 | 0.24748655 | 90.4319135 | 1.07E-18 | 1.60E-17 |
| ENOA | 209.228521 | 0.98530498 | 0.21318071 | 160.902777 | 9.36E-34 | 3.21E-32 |
| DPYL2 | 145.265047 | 0.98287898 | 0.12335438 | 220.171986 | 1.72E-46 | 1.17E-44 |
| PACN3 | 91.0953616 | 0.98242842 | 0.1694439 | 46.759231 | 1.71E-09 | 1.25E-08 |
| EM55 | 61.3737534 | 0.9806518 | 0.28264077 | 84.7260852 | 1.73E-17 | 2.39E-16 |
| DKK1 | 130.647983 | 0.97826886 | 0.20028292 | 153.138172 | 4.33E-32 | 1.33E-30 |
| F110B | 20.9252755 | 0.97814142 | 0.26557716 | 67.9328301 | 6.20E-14 | 6.49E-13 |
| CLUS | 832.232504 | 0.97796245 | 0.21824589 | 189.89856 | 5.57E-40 | 2.57E-38 |
| LYRM7 | 24.9034428 | 0.97704105 | 0.24523839 | 22.1482232 | 0.00018725 | 0.0007526 |
| PRM1A.1 | 14.4521144 | 0.97609712 | 0.30204695 | 21.6748694 | 0.00023261 | 0.00092133 |
| STMN1 | 1170.92992 | 0.9727591 | 0.15007223 | 81.0648403 | 1.04E-16 | 1.35E-15 |
| FGFR1 | 305.217536 | 0.97228704 | 0.1297372 | 269.389436 | 4.32E-57 | 5.42E-55 |
| SAPC2.2 | 128.97767 | 0.9715457 | 0.17765492 | 54.6227047 | 3.90E-11 | 3.32E-10 |
| RIMB2 | 27.8264738 | 0.97104538 | 0.28361644 | 143.132793 | 6.02E-30 | 1.68E-28 |
| RAI14.1 | 17.4420413 | 0.97104112 | 0.27511957 | 30.6221728 | 3.66E-06 | 1.87E-05 |
| NHSL2 | 120.533384 | 0.96863821 | 0.55242969 | 22.990493 | 0.00012718 | 0.00052438 |
| CNN3.1 | 471.27123 | 0.96645313 | 0.09968311 | 237.56835 | 3.10E-50 | 2.76E-48 |
| PP14C | 32.3964614 | 0.96628739 | 0.34291328 | 68.2047437 | 5.43E-14 | 5.74E-13 |
| CEIP2 | 165.941487 | 0.96598028 | 0.12201617 | 219.332229 | 2.61E-46 | 1.74E-44 |
| CD248 | 93.300731 | 0.96237291 | 0.25911323 | 81.623426 | 7.89E-17 | 1.04E-15 |
| PDGFA.1 | 86.2680316 | 0.96115826 | 0.12748311 | 113.760069 | 1.15E-23 | 2.31E-22 |
| CP26A | 27.4792946 | 0.95985339 | 0.26444216 | 78.9053434 | 2.97E-16 | 3.75E-15 |
| LRC20 | 25.0414796 | 0.95833189 | 0.25076083 | 24.0425836 | 7.83E-05 | 0.00033418 |
| DLX2A | 46.5425298 | 0.95783787 | 0.23553786 | 121.29638 | 2.82E-25 | 6.12E-24 |
| RB39B | 6.91309652 | 0.95685189 | 0.5433508 | 23.638781 | 9.44E-05 | 0.00039728 |
| MMP11 | 123.768295 | 0.95603789 | 0.12187947 | 86.1245132 | 8.76E-18 | 1.23E-16 |
| TBB | 61.8359459 | 0.95524237 | 0.1540337 | 122.525699 | 1.54E-25 | 3.41E-24 |
| CO4A1.1 | 330.620569 | 0.95521722 | 0.0858099 | 212.423026 | 8.00E-45 | 4.86E-43 |
| AKA12 | 22.5803265 | 0.95521076 | 0.24193718 | 25.3214366 | 4.34E-05 | 0.00019298 |
| RAB34 | 125.961375 | 0.95472865 | 0.12076052 | 154.407208 | 2.31E-32 | 7.27E-31 |
| CADH2 | 89.4562727 | 0.95414201 | 0.17239245 | 140.051091 | 2.75E-29 | 7.47E-28 |
| ID1 | 527.943734 | 0.95315254 | 0.08355976 | 469.280597 | 2.95E-100 | 1.22E-97 |
| SPY1 | 58.757726 | 0.9521839 | 0.16737757 | 59.7547159 | 3.27E-12 | 3.00E-11 |
| IGK | 27.3722829 | 0.95211271 | 0.27568898 | 34.2689549 | 6.56E-07 | 3.70E-06 |
| MXRA7 | 256.586692 | 0.94867843 | 0.18644601 | 74.0322756 | 3.19E-15 | 3.73E-14 |
| IGF2 | 31.5940111 | 0.94653048 | 0.23575016 | 25.3301031 | 4.32E-05 | 0.00019232 |
| KALM.1 | 61.4547151 | 0.94376819 | 0.19635816 | 46.4082668 | 2.03E-09 | 1.47E-08 |
| CO4A1 | 34.8644368 | 0.94298978 | 0.44356547 | 170.900011 | 6.70E-36 | 2.53E-34 |
| STAC | 9.02556538 | 0.94168114 | 0.54382423 | 26.7525603 | 2.23E-05 | 0.00010392 |
| HS12B | 13.2007596 | 0.93798555 | 0.40641342 | 34.1197445 | 7.04E-07 | 3.95E-06 |
| EMID1 | 67.6414211 | 0.93618715 | 0.13899147 | 103.796366 | 1.53E-21 | 2.68E-20 |
| ENOB | 1829.85445 | 0.93300242 | 0.12610758 | 66.7453132 | 1.10E-13 | 1.12E-12 |
| CPXM2.1 | 24.3551504 | 0.93081823 | 0.44994354 | 131.907634 | 1.52E-27 | 3.78E-26 |
| ANGL1 | 132.570983 | 0.92803573 | 0.28924349 | 91.6624124 | 5.84E-19 | 8.93E-18 |
| ANGP2.1 | 35.6526522 | 0.92708277 | 0.28558393 | 34.3545669 | 6.30E-07 | 3.56E-06 |
| VWA1 | 71.379808 | 0.92595757 | 0.17345222 | 46.7461656 | 1.72E-09 | 1.26E-08 |
| LDHA.1 | 240.499025 | 0.92582813 | 0.10743394 | 141.444525 | 1.38E-29 | 3.82E-28 |
| TSP2.1 | 99.1337401 | 0.92540963 | 0.21195305 | 281.577503 | 1.02E-59 | 1.43E-57 |
| NKD2 | 86.4134763 | 0.92172823 | 0.20464512 | 53.2193284 | 7.67E-11 | 6.34E-10 |
| LRC71 | 28.1097562 | 0.91887819 | 0.29652597 | 22.6781148 | 0.00014682 | 0.00060164 |
| MEG10 | 28.5480058 | 0.91756567 | 0.25701644 | 24.5154666 | 6.30E-05 | 0.00027247 |
| TFR2 | 7.07290172 | 0.91719566 | 0.63910757 | 29.2353511 | 7.00E-06 | 3.49E-05 |
| LRRN3 | 8.66579031 | 0.91712996 | 0.40593598 | 25.2750474 | 4.43E-05 | 0.00019689 |
| 1-Mar | 69.757738 | 0.9162327 | 0.17185914 | 84.0689461 | 2.39E-17 | 3.27E-16 |
| FYN.1 | 43.8001239 | 0.91592984 | 0.17768793 | 41.7009584 | 1.92E-08 | 1.27E-07 |
| SYNP2 | 15.8262109 | 0.91474605 | 0.3042097 | 32.7301514 | 1.36E-06 | 7.32E-06 |
| KCNJ8.1 | 9.17523875 | 0.91352088 | 0.47139131 | 31.7375227 | 2.16E-06 | 1.14E-05 |
| KCNS2 | 21.8846962 | 0.91244 | 0.30673683 | 43.126148 | 9.74E-09 | 6.60E-08 |
| DYL2 | 142.776708 | 0.91166197 | 0.11849777 | 146.275953 | 1.28E-30 | 3.73E-29 |
| 41 | 84.9583659 | 0.91162911 | 0.1397939 | 98.8133661 | 1.76E-20 | 2.92E-19 |
| PHTF2 | 102.172845 | 0.91060196 | 0.21066351 | 174.924613 | 9.17E-37 | 3.59E-35 |
| GALA | 6.48909578 | 0.91045254 | 0.90340032 | 25.5331352 | 3.93E-05 | 0.00017625 |
| RAB15 | 18.9002851 | 0.91039382 | 0.26838768 | 23.6665383 | 9.32E-05 | 0.00039266 |
| DCX | 60.4533996 | 0.91030009 | 0.17423182 | 101.997435 | 3.69E-21 | 6.37E-20 |
| COMTB | 64.1768912 | 0.91025729 | 0.18330651 | 43.9758177 | 6.49E-09 | 4.50E-08 |
| EPHB3 | 109.280949 | 0.90938849 | 0.1992123 | 69.6312455 | 2.72E-14 | 2.95E-13 |
| RFOX1 | 28.8642758 | 0.90764344 | 0.2727203 | 63.1746342 | 6.24E-13 | 6.02E-12 |
| SEM3A | 64.8347798 | 0.90081771 | 0.1503987 | 81.9682392 | 6.67E-17 | 8.84E-16 |
| PPIA | 392.14456 | 0.90032355 | 0.07368537 | 313.273707 | 1.48E-66 | 2.80E-64 |
| TTC28 | 7.85796048 | 0.89697457 | 0.45539868 | 27.0831744 | 1.91E-05 | 8.98E-05 |
| PGAM2 | 466.522159 | 0.89692162 | 0.1135423 | 82.4099113 | 5.37E-17 | 7.19E-16 |
| ST2B1 | 143.983992 | 0.89215168 | 0.23912344 | 27.1713868 | 1.84E-05 | 8.65E-05 |
| TM145 | 21.7640863 | 0.88955581 | 0.26551799 | 32.9191192 | 1.24E-06 | 6.75E-06 |
| EPO | 5.58111037 | 0.8877177 | 0.58722814 | 25.590174 | 3.83E-05 | 0.00017198 |
| VAV | 35.6530653 | 0.88662676 | 0.21416086 | 76.2986182 | 1.06E-15 | 1.28E-14 |
| VAFNB | 7.01227068 | 0.88654098 | 0.63411861 | 34.2204928 | 6.71E-07 | 3.78E-06 |
| HEY2 | 45.0760579 | 0.88620263 | 0.18000099 | 30.6659937 | 3.58E-06 | 1.84E-05 |
| TSP2 | 147.808263 | 0.8850893 | 0.22609634 | 245.477391 | 6.13E-52 | 5.98E-50 |
| FGRL1.1 | 6.17450207 | 0.88499604 | 0.61211417 | 23.6488804 | 9.39E-05 | 0.00039555 |
| RALY | 16.1517736 | 0.88464766 | 0.34181823 | 51.4132608 | 1.83E-10 | 1.47E-09 |
| COT2 | 106.788094 | 0.88333756 | 0.11675023 | 243.081626 | 2.01E-51 | 1.93E-49 |
| SPTB2 | 132.826071 | 0.88285158 | 0.16209505 | 39.9328574 | 4.47E-08 | 2.85E-07 |
| S27A6 | 31.1247401 | 0.88159961 | 0.22514075 | 59.2817802 | 4.11E-12 | 3.75E-11 |
| AUTS2.1 | 40.1886169 | 0.88146527 | 0.22254115 | 39.6915568 | 5.01E-08 | 3.18E-07 |
| SX18A | 103.566229 | 0.88036135 | 0.12505384 | 89.4426353 | 1.73E-18 | 2.56E-17 |
| EFHC1 | 27.9508894 | 0.87980407 | 0.2350282 | 29.0435509 | 7.66E-06 | 3.79E-05 |
| NEXMI | 31.619048 | 0.8792928 | 0.22227301 | 46.4198986 | 2.01E-09 | 1.46E-08 |
| LRRN1 | 12.2811481 | 0.87838796 | 0.33626576 | 23.0081319 | 0.00012615 | 0.00052071 |
| COEA1 | 324.493448 | 0.87797177 | 0.14610423 | 80.7870488 | 1.19E-16 | 1.54E-15 |
| ANDR | 8.967406 | 0.87743825 | 0.45245795 | 33.7704094 | 8.31E-07 | 4.62E-06 |
| CO1A2 | 458.148012 | 0.87572125 | 0.14927311 | 153.537901 | 3.55E-32 | 1.10E-30 |
| SVIL.1 | 80.193799 | 0.87304114 | 0.14679334 | 43.3848754 | 8.61E-09 | 5.86E-08 |
| CXA1 | 44.9222546 | 0.87177541 | 0.24964895 | 23.3291602 | 0.00010882 | 0.00045358 |
| FBN2.4 | 101.778814 | 0.87044358 | 0.16117147 | 89.3357081 | 1.82E-18 | 2.69E-17 |
| IBP2 | 66.7311842 | 0.87039275 | 0.22530215 | 100.18556 | 8.98E-21 | 1.52E-19 |
| TWIST | 56.0842076 | 0.86955009 | 0.16335891 | 140.522663 | 2.18E-29 | 5.94E-28 |
| LX15B | 362.966745 | 0.86929762 | 0.19057089 | 532.259943 | 7.05E-114 | 4.90E-111 |
| SCMH1.1 | 20.595292 | 0.86913323 | 0.24998321 | 37.6001703 | 1.35E-07 | 8.15E-07 |
| CHADL | 28.2248589 | 0.86786016 | 0.41114913 | 25.3867947 | 4.21E-05 | 0.00018777 |
| CF157 | 38.8307418 | 0.86777669 | 0.20449649 | 44.3860912 | 5.33E-09 | 3.73E-08 |
| LIPO | 1552.55487 | 0.86643387 | 0.6948078 | 29.2901401 | 6.83E-06 | 3.40E-05 |
| RT07 | 183.9721 | 0.86460972 | 0.15234569 | 90.5720006 | 9.95E-19 | 1.50E-17 |
| SDC3 | 35.6443569 | 0.86408305 | 0.20335795 | 61.0813696 | 1.72E-12 | 1.61E-11 |
| PP14A | 195.967053 | 0.86358192 | 0.26056193 | 63.3116654 | 5.83E-13 | 5.64E-12 |
| PZRN4 | 30.9902971 | 0.86153208 | 0.22703711 | 62.2422841 | 9.80E-13 | 9.31E-12 |
| STRA6 | 37.1755431 | 0.86130721 | 0.27975263 | 70.6087353 | 1.69E-14 | 1.86E-13 |
| LYPA1 | 38.7035891 | 0.85736541 | 0.2028523 | 65.8825161 | 1.68E-13 | 1.69E-12 |
| TWF2 | 14.0983823 | 0.85709373 | 0.33961256 | 27.2419244 | 1.78E-05 | 8.38E-05 |
| EYA1 | 12.344126 | 0.85662349 | 0.36863039 | 27.8536919 | 1.34E-05 | 6.41E-05 |
| CHSS2 | 70.5399919 | 0.8549087 | 0.15581384 | 88.9619833 | 2.19E-18 | 3.21E-17 |
| AGRE1.5 | 8.18466611 | 0.85408771 | 0.45141896 | 27.8111479 | 1.36E-05 | 6.53E-05 |
| PDLI4 | 40.1515174 | 0.85374151 | 0.26592954 | 66.644144 | 1.16E-13 | 1.17E-12 |
| FBLN4 | 358.363839 | 0.85215195 | 0.20056533 | 201.96714 | 1.42E-42 | 7.67E-41 |
| SOBP | 26.6725042 | 0.85201567 | 0.27609339 | 66.5722459 | 1.20E-13 | 1.22E-12 |
| EMIL1 | 38.1732997 | 0.85131784 | 0.21485004 | 67.8173018 | 6.56E-14 | 6.84E-13 |
| GTR10 | 16.1885402 | 0.85074183 | 0.32621544 | 76.5082792 | 9.56E-16 | 1.16E-14 |
| TMM47.1 | 93.7016671 | 0.84912543 | 0.15515856 | 114.707984 | 7.20E-24 | 1.46E-22 |
| LIPE.1 | 56.6719163 | 0.84644435 | 0.18062387 | 46.007911 | 2.45E-09 | 1.77E-08 |
| 5NT1A | 33.3315068 | 0.8429029 | 0.33184217 | 90.3057047 | 1.13E-18 | 1.70E-17 |
| SARDH | 22.356595 | 0.84103664 | 0.25620734 | 41.5847314 | 2.03E-08 | 1.34E-07 |
| SOX9 | 188.145221 | 0.84014333 | 0.10386764 | 113.681345 | 1.19E-23 | 2.39E-22 |
| RAP1B | 80.8602618 | 0.84011924 | 0.13739732 | 85.3865116 | 1.26E-17 | 1.75E-16 |
| S12A4 | 85.6517727 | 0.83940583 | 0.1344922 | 145.477354 | 1.90E-30 | 5.44E-29 |
| HAAF | 11.3645666 | 0.83625454 | 0.41683396 | 33.437733 | 9.72E-07 | 5.36E-06 |
| FREM1 | 12.39592 | 0.83547503 | 0.38944707 | 28.6560768 | 9.18E-06 | 4.50E-05 |
| RBPS2 | 72.1027667 | 0.8332282 | 0.1581942 | 36.0672068 | 2.80E-07 | 1.64E-06 |
| PSIP1 | 146.065416 | 0.83265647 | 0.10623679 | 90.7276597 | 9.22E-19 | 1.40E-17 |
| SOX10 | 60.6215303 | 0.83264502 | 0.15254788 | 35.5590634 | 3.57E-07 | 2.06E-06 |
| PGFRL | 50.121373 | 0.83148352 | 0.32400263 | 54.9917455 | 3.26E-11 | 2.80E-10 |
| NPNT | 162.562934 | 0.82988359 | 0.14783582 | 62.5011887 | 8.64E-13 | 8.25E-12 |
| CC85A | 19.7212322 | 0.82766179 | 0.26734741 | 26.0801585 | 3.05E-05 | 0.00013917 |
| FBN2.1 | 38.5391067 | 0.82616443 | 0.23010293 | 64.1740507 | 3.84E-13 | 3.76E-12 |
| CALCB | 24.3970131 | 0.82570747 | 0.32480091 | 47.8710965 | 1.00E-09 | 7.54E-09 |
| STON1 | 17.3405321 | 0.8200231 | 0.35273264 | 39.6442723 | 5.13E-08 | 3.24E-07 |
| ONCO | 52.9513825 | 0.81737824 | 0.74198913 | 31.6634456 | 2.24E-06 | 1.18E-05 |
| FXC1A | 47.1278219 | 0.81718028 | 0.18410447 | 32.094065 | 1.83E-06 | 9.73E-06 |
| PGBM.1 | 331.464679 | 0.8127576 | 0.14396968 | 75.2311477 | 1.78E-15 | 2.12E-14 |
| NPNT.1 | 224.499025 | 0.81268828 | 0.11600016 | 109.404278 | 9.75E-23 | 1.84E-21 |
| APLP2 | 1044.71467 | 0.81230395 | 0.12275984 | 238.276739 | 2.18E-50 | 1.96E-48 |
| PREX1 | 13.7511952 | 0.81170415 | 0.31354354 | 23.4883129 | 0.00010113 | 0.00042406 |
| RAB38 | 25.2802276 | 0.80584853 | 0.34123345 | 73.6644743 | 3.82E-15 | 4.44E-14 |
| NXPE3 | 45.0224328 | 0.80584248 | 0.23530876 | 79.4346459 | 2.29E-16 | 2.92E-15 |
| BGH3 | 312.023701 | 0.8043454 | 0.24931509 | 57.0912031 | 1.18E-11 | 1.05E-10 |
| CPMD8 | 159.321544 | 0.8031601 | 0.14994011 | 72.6760616 | 6.18E-15 | 7.03E-14 |
| C4BPA.1 | 33.348323 | 0.80256743 | 0.26317737 | 107.651631 | 2.31E-22 | 4.28E-21 |
| ITB1B | 21.1557533 | 0.80159997 | 0.41103167 | 29.5817809 | 5.95E-06 | 2.99E-05 |
| DLX1 | 19.7590758 | 0.80137955 | 0.29651886 | 34.4246374 | 6.10E-07 | 3.45E-06 |
| AGRA3 | 88.6244345 | 0.80088124 | 0.21796866 | 67.2741353 | 8.53E-14 | 8.78E-13 |
| CADM1 | 265.444969 | 0.7988213 | 0.14136145 | 214.896756 | 2.35E-45 | 1.48E-43 |
| PCDGI | 17.0705276 | 0.79763099 | 0.31892519 | 53.7726435 | 5.87E-11 | 4.91E-10 |
| MEX3B | 37.4876407 | 0.79621624 | 0.20453508 | 86.6219481 | 6.87E-18 | 9.74E-17 |
| CATF | 29.5397017 | 0.79470667 | 0.51268023 | 24.0984594 | 7.63E-05 | 0.00032603 |
| GLCNE | 23.9366867 | 0.79406102 | 0.23417478 | 47.7735805 | 1.05E-09 | 7.89E-09 |
| FBLN3 | 108.29851 | 0.79090903 | 0.36314504 | 34.1710964 | 6.87E-07 | 3.87E-06 |
| BAMBI.1 | 45.6063259 | 0.79061716 | 0.30164099 | 97.0003151 | 4.28E-20 | 6.96E-19 |
| TRIL | 22.7754712 | 0.78918619 | 0.2807771 | 26.8948051 | 2.09E-05 | 9.76E-05 |
| MSRB3 | 75.6477722 | 0.78760048 | 0.25180093 | 22.2036673 | 0.00018255 | 0.00073544 |
| MB211 | 11.520244 | 0.78678627 | 0.47698175 | 56.2478489 | 1.78E-11 | 1.56E-10 |
| CYT.2 | 75.9942853 | 0.78644651 | 0.22830612 | 85.9337201 | 9.61E-18 | 1.35E-16 |
| SSUH2.1 | 33.3732735 | 0.7853361 | 0.20259584 | 23.1669682 | 0.00011726 | 0.00048637 |
| CCDC3.1 | 110.700444 | 0.78195332 | 0.23146834 | 118.507087 | 1.11E-24 | 2.35E-23 |
| LAMC3.1 | 303.849424 | 0.78015075 | 0.1556043 | 71.7992344 | 9.46E-15 | 1.07E-13 |
| GFRA1 | 173.344075 | 0.78004607 | 0.15687463 | 57.9459628 | 7.83E-12 | 7.02E-11 |
| F181B | 11.0733795 | 0.77951166 | 0.37877481 | 23.0459177 | 0.00012398 | 0.00051242 |
| PITX1 | 605.076466 | 0.77876654 | 0.09395989 | 171.66065 | 4.60E-36 | 1.75E-34 |
| UD11 | 30.4655177 | 0.77794794 | 0.22742733 | 25.511315 | 3.97E-05 | 0.00017781 |
| VGP.1 | 699.190532 | 0.77633872 | 0.1251518 | 158.349745 | 3.30E-33 | 1.10E-31 |
| ATS8 | 29.4217996 | 0.77595177 | 0.22782084 | 54.6043236 | 3.93E-11 | 3.35E-10 |
| CYBR1 | 63.6741377 | 0.77591674 | 0.17589071 | 58.6395654 | 5.60E-12 | 5.06E-11 |
| ACKR4 | 69.4214405 | 0.77510257 | 0.16803103 | 116.873016 | 2.49E-24 | 5.18E-23 |
| PZRN3 | 25.1925809 | 0.7750467 | 0.30330703 | 75.8970145 | 1.29E-15 | 1.55E-14 |
| DLX4B | 462.103146 | 0.77252786 | 0.12156298 | 567.969266 | 1.32E-121 | 1.35E-118 |
| GHC2 | 1352.04624 | 0.77116733 | 0.0857064 | 216.80448 | 9.13E-46 | 6.00E-44 |
| DAG1 | 375.014553 | 0.7704235 | 0.10200491 | 101.773553 | 4.12E-21 | 7.07E-20 |
| GP161 | 27.1476121 | 0.76759622 | 0.22163653 | 51.6845388 | 1.61E-10 | 1.29E-09 |
| CD276.1 | 17.0294626 | 0.76549016 | 0.40481604 | 74.9019528 | 2.09E-15 | 2.47E-14 |
| RGAP1 | 70.4626109 | 0.76420489 | 0.17742431 | 36.0344678 | 2.85E-07 | 1.66E-06 |
| CBPA1 | 171.870608 | 0.76341721 | 0.29132336 | 232.407598 | 4.00E-49 | 3.22E-47 |
| ZFP2.8 | 26.2710948 | 0.75964869 | 0.25314419 | 26.7479724 | 2.24E-05 | 0.00010411 |
| MSI1H | 48.4994483 | 0.75888681 | 0.21247955 | 43.4335684 | 8.41E-09 | 5.74E-08 |
| TCEA1 | 31.280255 | 0.75818129 | 0.20329446 | 22.6212947 | 0.00015071 | 0.0006159 |
| RGS5 | 1028.12466 | 0.75785943 | 0.14233385 | 32.8567249 | 1.28E-06 | 6.93E-06 |
| FAT1.4 | 154.112659 | 0.75737731 | 0.11111164 | 165.707214 | 8.72E-35 | 3.14E-33 |
| MARH3 | 3.00789077 | 0.75680222 | 1.12557997 | 24.5517169 | 6.19E-05 | 0.0002684 |
| COIA1 | 249.382947 | 0.75643683 | 0.12144506 | 42.8198411 | 1.13E-08 | 7.61E-08 |
| COEA1.1 | 107.319285 | 0.75536632 | 0.11439304 | 81.6376969 | 7.83E-17 | 1.03E-15 |
| BIVM | 44.0861191 | 0.75523974 | 0.18144794 | 65.7938069 | 1.75E-13 | 1.76E-12 |
| HBB | 1383.46033 | 0.75473355 | 0.5608907 | 37.6957523 | 1.29E-07 | 7.81E-07 |
| HBA | 1306.58409 | 0.75377056 | 0.48335129 | 50.8595113 | 2.39E-10 | 1.90E-09 |
| SNF5.1 | 328.166358 | 0.75335752 | 0.11423503 | 77.0686973 | 7.27E-16 | 8.92E-15 |
| SEPT9.1 | 42.2914586 | 0.75289609 | 0.19329262 | 34.78742 | 5.14E-07 | 2.93E-06 |
| PAX9 | 102.450311 | 0.75243573 | 0.14434083 | 59.3722467 | 3.93E-12 | 3.59E-11 |
| G3P | 2086.71122 | 0.75035859 | 0.13170294 | 43.3951176 | 8.57E-09 | 5.84E-08 |
| CE170 | 27.6729809 | 0.75027376 | 0.2354175 | 69.9512752 | 2.32E-14 | 2.53E-13 |
| CBY1 | 34.9846232 | 0.75024355 | 0.22686771 | 34.6419801 | 5.50E-07 | 3.12E-06 |
| EGR2B | 35.8458833 | 0.75003655 | 0.26758814 | 46.9844718 | 1.54E-09 | 1.13E-08 |
| RIPL1 | 68.656827 | -0.7515455 | 0.22611067 | 22.8747111 | 0.00013414 | 0.00055217 |
| BIRC1 | 40.8790522 | -0.7523491 | 0.24266379 | 77.4531591 | 6.03E-16 | 7.45E-15 |
| SOX7 | 9.24339236 | -0.7528532 | 0.84602683 | 91.1153405 | 7.63E-19 | 1.16E-17 |
| AT2L1 | 268.643596 | -0.754453 | 0.13751056 | 45.3505141 | 3.36E-09 | 2.39E-08 |
| BTK | 17.2817667 | -0.7554466 | 0.47402731 | 104.33037 | 1.18E-21 | 2.08E-20 |
| LGMN.1 | 232.080694 | -0.7563354 | 0.12089628 | 196.033301 | 2.68E-41 | 1.37E-39 |
| ZN208.1 | 26.4965206 | -0.7565071 | 0.27374195 | 32.6752814 | 1.39E-06 | 7.50E-06 |
| PSB9 | 92.3158042 | -0.7571436 | 0.18387798 | 184.600055 | 7.66E-39 | 3.42E-37 |
| RND2 | 144.539739 | -0.7597706 | 0.13435127 | 170.172046 | 9.61E-36 | 3.59E-34 |
| RN223 | 63.9869757 | -0.7602528 | 0.21514778 | 166.61127 | 5.58E-35 | 2.02E-33 |
| ABCAD | 13.231528 | -0.7609005 | 0.51773094 | 32.9460034 | 1.23E-06 | 6.66E-06 |
| ZN214.81 | 52.9722235 | -0.7629625 | 0.22377462 | 32.1635742 | 1.77E-06 | 9.43E-06 |
| GOLM1 | 383.762922 | -0.7629858 | 0.11513069 | 121.838285 | 2.16E-25 | 4.74E-24 |
| NADK | 34.6160579 | -0.7642735 | 0.26456465 | 31.3344505 | 2.62E-06 | 1.36E-05 |
| OTOP3 | 45.670099 | -0.7645462 | 0.27464624 | 23.6593261 | 9.35E-05 | 0.00039375 |
| GSH1 | 322.71405 | -0.764959 | 0.16902188 | 328.173308 | 9.03E-70 | 1.97E-67 |
| IF44L.1 | 40.8480707 | -0.7660002 | 0.26492013 | 44.1149797 | 6.07E-09 | 4.22E-08 |
| S2543 | 68.7715984 | -0.7683924 | 0.17227574 | 73.9023975 | 3.40E-15 | 3.97E-14 |
| STRP2 | 34.0598146 | -0.7697643 | 0.24716309 | 73.8627264 | 3.47E-15 | 4.04E-14 |
| ITB3 | 318.800584 | -0.7711167 | 0.1478964 | 157.07245 | 6.20E-33 | 2.04E-31 |
| MOT12.1 | 16.69579 | -0.7712904 | 0.38529206 | 45.529956 | 3.08E-09 | 2.20E-08 |
| MUC16.3 | 469.642725 | -0.772759 | 0.14653882 | 247.063533 | 2.79E-52 | 2.74E-50 |
| GRT1A | 52.9834786 | -0.7728065 | 0.20665555 | 47.0121252 | 1.52E-09 | 1.12E-08 |
| RNF38.1 | 79.744368 | -0.7731927 | 0.17297042 | 77.6776436 | 5.41E-16 | 6.71E-15 |
| UAP1 | 77.8846641 | -0.7736539 | 0.19251691 | 52.3356075 | 1.17E-10 | 9.54E-10 |
| TNF14 | 8.60321136 | -0.7740491 | 0.63836036 | 26.6941243 | 2.29E-05 | 0.00010659 |
| STK40 | 39.2885713 | -0.7743611 | 0.25928932 | 30.0195236 | 4.85E-06 | 2.45E-05 |
| SYTL1 | 66.1396585 | -0.7745368 | 0.22211829 | 40.1530466 | 4.02E-08 | 2.58E-07 |
| STBD1 | 58.7618028 | -0.7752865 | 0.2063081 | 180.274689 | 6.51E-38 | 2.69E-36 |
| SOX14 | 113.207928 | -0.780557 | 0.18228095 | 214.434014 | 2.95E-45 | 1.85E-43 |
| RHOG.1 | 82.1903637 | -0.7809833 | 0.18418862 | 180.209093 | 6.72E-38 | 2.77E-36 |
| SDCB1 | 753.314366 | -0.7810805 | 0.10280913 | 225.179009 | 1.44E-47 | 1.05E-45 |
| CATL1.4 | 162.405481 | -0.7822142 | 0.19135541 | 146.493757 | 1.15E-30 | 3.35E-29 |
| S6A14 | 23.1360363 | -0.7828685 | 0.42023997 | 128.082754 | 1.00E-26 | 2.37E-25 |
| PAR14.1 | 8.31671362 | -0.7836663 | 0.52504392 | 24.8666986 | 5.35E-05 | 0.00023454 |
| NLS1 | 18.3242931 | -0.7841077 | 0.35282617 | 44.7687602 | 4.44E-09 | 3.13E-08 |
| FGD3 | 18.221527 | -0.7857444 | 0.33733584 | 33.5211143 | 9.34E-07 | 5.16E-06 |
| CC038.1 | 78.3516188 | -0.7860054 | 0.20161358 | 128.317367 | 8.92E-27 | 2.13E-25 |
| MB12A | 262.206523 | -0.7862069 | 0.09706565 | 298.549573 | 2.23E-63 | 3.63E-61 |
| CD22 | 6.17561346 | -0.7864705 | 0.72412454 | 47.7608279 | 1.06E-09 | 7.93E-09 |
| KLF7 | 14.3417943 | -0.7871257 | 0.38948926 | 22.434502 | 0.0001642 | 0.00066697 |
| LFG3 | 111.917141 | -0.7881861 | 0.1868445 | 29.0209145 | 7.74E-06 | 3.83E-05 |
| M4K3 | 37.866022 | -0.7882799 | 0.29419822 | 139.510406 | 3.59E-29 | 9.66E-28 |
| CBL | 85.5437567 | -0.7887071 | 0.17384194 | 180.695425 | 5.29E-38 | 2.20E-36 |
| RPH3L | 27.2347169 | -0.7888622 | 0.29328146 | 46.6820562 | 1.78E-09 | 1.30E-08 |
| PTN18 | 74.962601 | -0.7891198 | 0.18978864 | 74.9501265 | 2.04E-15 | 2.42E-14 |
| MACC1 | 43.1955861 | -0.7909765 | 0.23163508 | 57.9341982 | 7.88E-12 | 7.06E-11 |
| CD2AP.1 | 160.449643 | -0.7918556 | 0.13206727 | 199.670907 | 4.42E-42 | 2.36E-40 |
| TNF15 | 17.9890117 | -0.7924094 | 0.35220161 | 21.6785988 | 0.00023221 | 0.00092 |
| TM165 | 129.855555 | -0.7950124 | 0.12793202 | 65.7220787 | 1.81E-13 | 1.82E-12 |
| ASPN | 922.5829 | -0.7963331 | 0.13781278 | 42.7599297 | 1.16E-08 | 7.81E-08 |
| VILI | 1151.08944 | -0.7969481 | 0.0928417 | 257.827879 | 1.34E-54 | 1.45E-52 |
| STXB | 6.07720188 | -0.7979399 | 0.69970504 | 32.5865726 | 1.45E-06 | 7.80E-06 |
| NBEL2 | 42.6443712 | -0.7979408 | 0.21387539 | 46.3024884 | 2.13E-09 | 1.54E-08 |
| XCT | 16.5868723 | -0.8001204 | 0.40036435 | 34.6740259 | 5.42E-07 | 3.08E-06 |
| ASSY | 2103.35738 | -0.8008162 | 0.15884149 | 175.417831 | 7.19E-37 | 2.84E-35 |
| ES8L1 | 265.937541 | -0.8009154 | 0.11698624 | 237.633542 | 3.00E-50 | 2.68E-48 |
| PKHB2 | 60.273056 | -0.8022999 | 0.19173241 | 113.613779 | 1.23E-23 | 2.47E-22 |
| GNMT | 256.868004 | -0.8043327 | 0.13931113 | 499.800686 | 7.40E-107 | 3.90E-104 |
| PINLY.1 | 56.5516939 | -0.8054737 | 0.39025512 | 73.5193566 | 4.10E-15 | 4.74E-14 |
| C2C4C | 71.5010277 | -0.806358 | 0.17644314 | 139.402353 | 3.79E-29 | 1.02E-27 |
| CCL20.1 | 19.0106099 | -0.807224 | 0.70185033 | 24.8705411 | 5.34E-05 | 0.00023429 |
| GALT4 | 130.786888 | -0.8092072 | 0.13784784 | 111.71191 | 3.14E-23 | 6.15E-22 |
| DCBD1 | 43.9226674 | -0.8092072 | 0.25028028 | 178.736211 | 1.39E-37 | 5.70E-36 |
| PRDX6 | 1962.63602 | -0.809371 | 0.11604166 | 223.207371 | 3.83E-47 | 2.72E-45 |
| A33.43 | 60.5168966 | -0.8098779 | 0.44818531 | 67.6417058 | 7.14E-14 | 7.41E-13 |
| SPIT1 | 227.068494 | -0.8104007 | 0.1378849 | 120.767754 | 3.66E-25 | 7.90E-24 |
| DDX6 | 75.6231296 | -0.8106436 | 0.27833551 | 26.8931415 | 2.09E-05 | 9.77E-05 |
| PNCB | 23.3010722 | -0.8124616 | 0.30819765 | 40.2966195 | 3.76E-08 | 2.42E-07 |
| FA83B | 137.354896 | -0.8131017 | 0.13714762 | 107.895492 | 2.05E-22 | 3.80E-21 |
| PDCD4.2 | 258.825571 | -0.8134817 | 0.11304509 | 154.31734 | 2.42E-32 | 7.55E-31 |
| ZO2.1 | 65.5710403 | -0.8155432 | 0.21866595 | 117.768837 | 1.60E-24 | 3.37E-23 |
| METL4 | 90.711716 | -0.8160053 | 0.15902831 | 88.7036176 | 2.48E-18 | 3.62E-17 |
| NCF1 | 40.7837533 | -0.8166441 | 0.23720112 | 53.3556467 | 7.18E-11 | 5.95E-10 |
| UPK3A | 396.436618 | -0.818427 | 0.27531649 | 204.80902 | 3.47E-43 | 1.90E-41 |
| CO039 | 313.726441 | -0.8188891 | 0.11242366 | 233.833814 | 1.97E-49 | 1.64E-47 |
| HPSE2 | 22.2108708 | -0.8195147 | 0.33887015 | 94.135649 | 1.74E-19 | 2.74E-18 |
| FRY | 124.011136 | -0.8196193 | 0.23803587 | 102.4087 | 3.02E-21 | 5.23E-20 |
| CPNS2 | 57.0484378 | -0.8203073 | 0.21444241 | 24.5775285 | 6.12E-05 | 0.00026567 |
| AGR2 | 3482.99732 | -0.8210536 | 0.14742136 | 340.33395 | 2.14E-72 | 5.12E-70 |
| HEXD.1 | 117.02525 | -0.8219727 | 0.14224224 | 82.1737148 | 6.03E-17 | 8.03E-16 |
| FUCO | 677.621058 | -0.8239631 | 0.13219302 | 69.4962981 | 2.90E-14 | 3.14E-13 |
| RBGP1 | 114.519707 | -0.8242906 | 0.21422383 | 56.2069267 | 1.81E-11 | 1.59E-10 |
| FXA1A | 24.9569721 | -0.8256983 | 0.44807181 | 318.80805 | 9.48E-68 | 1.86E-65 |
| ERBB4 | 8.2668026 | -0.8261029 | 0.61491109 | 37.9260764 | 1.16E-07 | 7.05E-07 |
| PCKGM | 163.485306 | -0.8280455 | 0.1454598 | 59.2339498 | 4.20E-12 | 3.83E-11 |
| ITLN1 | 4009.69766 | -0.8280596 | 0.40813788 | 252.859319 | 1.58E-53 | 1.63E-51 |
| CD9 | 87.3197661 | -0.8281046 | 0.18102478 | 229.258625 | 1.91E-48 | 1.47E-46 |
| CEBPB | 419.25517 | -0.8300044 | 0.11832353 | 109.624142 | 8.75E-23 | 1.66E-21 |
| A33.2 | 30.1986955 | -0.8300271 | 0.26406566 | 44.4028744 | 5.29E-09 | 3.70E-08 |
| ST17A | 63.975626 | -0.8301099 | 0.19376071 | 213.766581 | 4.11E-45 | 2.54E-43 |
| MOT7 | 71.5876687 | -0.8316318 | 0.23728995 | 28.1165787 | 1.18E-05 | 5.70E-05 |
| AHNK | 549.929489 | -0.831971 | 0.19878933 | 53.085194 | 8.18E-11 | 6.74E-10 |
| ARLY | 640.935433 | -0.833217 | 0.16140773 | 274.808891 | 2.93E-58 | 3.84E-56 |
| LECA.1 | 269.479749 | -0.8338958 | 0.51226129 | 142.342103 | 8.90E-30 | 2.47E-28 |
| GGCT | 285.455601 | -0.8354326 | 0.2066022 | 105.956398 | 5.30E-22 | 9.63E-21 |
| B3GT5.1 | 621.874972 | -0.8354494 | 0.12151277 | 52.7435544 | 9.64E-11 | 7.89E-10 |
| HPS6 | 84.9814523 | -0.8367393 | 0.26205395 | 26.5972883 | 2.40E-05 | 0.00011132 |
| HEBP1 | 85.5373252 | -0.8368861 | 0.20201996 | 102.737267 | 2.57E-21 | 4.47E-20 |
| ELN | 1829.62375 | -0.8373126 | 0.27884341 | 73.658104 | 3.83E-15 | 4.45E-14 |
| K2012 | 87.3884207 | -0.8376859 | 0.20338431 | 85.0621795 | 1.47E-17 | 2.04E-16 |
| OSGI1 | 136.098114 | -0.8379792 | 0.19558404 | 215.934246 | 1.41E-45 | 9.04E-44 |
| RAI3 | 113.971892 | -0.8382728 | 0.16356222 | 71.3470515 | 1.18E-14 | 1.32E-13 |
| NBEL1 | 18.2801121 | -0.8404151 | 0.35190688 | 36.3775049 | 2.42E-07 | 1.42E-06 |
| GCNT2 | 29.1566518 | -0.8415108 | 0.30886175 | 107.539341 | 2.44E-22 | 4.50E-21 |
| GP2 | 28.9789737 | -0.8418179 | 0.40896165 | 163.695231 | 2.36E-34 | 8.33E-33 |
| KCT2 | 112.539289 | -0.8423444 | 0.23700012 | 61.5422705 | 1.37E-12 | 1.30E-11 |
| CXB6 | 59.7372432 | -0.8424934 | 0.27498052 | 145.59087 | 1.79E-30 | 5.17E-29 |
| NR3L1.3 | 45.8468982 | -0.8433306 | 0.23761351 | 27.4749503 | 1.59E-05 | 7.56E-05 |
| K1324 | 13.9865159 | -0.8433501 | 0.58753638 | 156.16751 | 9.70E-33 | 3.12E-31 |
| CD276 | 153.614291 | -0.8451531 | 0.13739246 | 211.955659 | 1.01E-44 | 6.03E-43 |
| G6PC | 3.613516 | -0.8452995 | 1.24346384 | 22.1857274 | 0.00018406 | 0.00074094 |
| CXAR | 1639.75014 | -0.845829 | 0.22047805 | 135.825113 | 2.21E-28 | 5.73E-27 |
| RGF1A | 58.5554071 | -0.8463593 | 0.21311457 | 196.235958 | 2.42E-41 | 1.25E-39 |
| PIR | 36.8017791 | -0.8465923 | 0.24945847 | 35.3368582 | 3.96E-07 | 2.28E-06 |
| VTCN1.1 | 274.366409 | -0.8469782 | 0.24952566 | 183.69486 | 1.20E-38 | 5.26E-37 |
| S17A9 | 166.911815 | -0.8471752 | 0.14901849 | 95.2216494 | 1.02E-19 | 1.63E-18 |
| IMPA3.1 | 47.7384517 | -0.849032 | 0.22346327 | 65.7444226 | 1.79E-13 | 1.80E-12 |
| ALAT2.1 | 229.795266 | -0.8497127 | 0.1369078 | 102.864474 | 2.41E-21 | 4.20E-20 |
| ANXA2.1 | 775.281597 | -0.8502175 | 0.12297028 | 326.244888 | 2.35E-69 | 5.08E-67 |
| LDLR1 | 268.237977 | -0.8503441 | 0.14720082 | 47.5170541 | 1.19E-09 | 8.86E-09 |
| TM238.1 | 15.0805784 | -0.8511576 | 0.39858757 | 38.777654 | 7.74E-08 | 4.80E-07 |
| AVIL | 9.37567223 | -0.8520442 | 0.63225542 | 73.9574637 | 3.31E-15 | 3.87E-14 |
| LBH | 244.343597 | -0.8523473 | 0.13239268 | 359.318013 | 1.71E-76 | 4.75E-74 |
| PARM1 | 85.1266413 | -0.8530515 | 0.20551182 | 134.88451 | 3.51E-28 | 8.99E-27 |
| NLRP3.5 | 65.4279109 | -0.8532364 | 0.2945121 | 24.9826775 | 5.07E-05 | 0.00022349 |
| ERN1 | 60.3978603 | -0.8558049 | 0.19136862 | 127.289727 | 1.48E-26 | 3.47E-25 |
| ERO1B | 743.529581 | -0.856052 | 0.10244375 | 167.043718 | 4.51E-35 | 1.64E-33 |
| ADPRH | 910.802801 | -0.8564258 | 0.13080994 | 188.112052 | 1.35E-39 | 6.14E-38 |
| CSAD | 667.018158 | -0.8568423 | 0.17749446 | 71.1944762 | 1.27E-14 | 1.41E-13 |
| LPAR3 | 44.6082314 | -0.8570622 | 0.23124511 | 50.9019641 | 2.34E-10 | 1.86E-09 |
| GALT3.1 | 75.2858733 | -0.8573172 | 0.17848794 | 77.2946667 | 6.51E-16 | 8.01E-15 |
| MX3.4 | 95.3762468 | -0.8585024 | 0.16562588 | 224.058783 | 2.51E-47 | 1.80E-45 |
| K1522 | 43.8018302 | -0.8595743 | 0.2263344 | 26.0898829 | 3.04E-05 | 0.00013858 |
| FEL | 5918.5322 | -0.8622162 | 0.46693817 | 155.341593 | 1.46E-32 | 4.64E-31 |
| HUMMR | 53.1352565 | -0.8627124 | 0.23106917 | 71.4564048 | 1.12E-14 | 1.25E-13 |
| FGF13 | 86.2815217 | -0.863377 | 0.19407684 | 40.2670968 | 3.81E-08 | 2.45E-07 |
| CD63 | 1549.75238 | -0.8636583 | 0.06783112 | 193.592212 | 8.96E-41 | 4.41E-39 |
| ST3A1 | 11.7306861 | -0.8650379 | 0.50204219 | 41.7896372 | 1.84E-08 | 1.22E-07 |
| GRDN.1 | 18.9632052 | -0.8651196 | 0.35851394 | 44.7581588 | 4.46E-09 | 3.14E-08 |
| GOLP3.1 | 249.409117 | -0.8653408 | 0.13608844 | 122.589221 | 1.49E-25 | 3.31E-24 |
| GALT7 | 183.858391 | -0.8655146 | 0.15350353 | 141.031602 | 1.70E-29 | 4.65E-28 |
| HHLA2.1 | 32.3275109 | -0.8667807 | 0.30999619 | 225.681268 | 1.12E-47 | 8.30E-46 |
| TMCC3 | 69.2401876 | -0.8690358 | 0.17650028 | 79.0708988 | 2.74E-16 | 3.46E-15 |
| I20RA.1 | 24.8631547 | -0.8700669 | 0.35793533 | 69.0632457 | 3.58E-14 | 3.83E-13 |
| KLF4.1 | 520.243732 | -0.8706769 | 0.08264921 | 238.460694 | 1.99E-50 | 1.80E-48 |
| LOX12 | 10.2350637 | -0.8707262 | 0.82120449 | 26.0325362 | 3.12E-05 | 0.000142 |
| ELF3.1 | 239.463215 | -0.8707844 | 0.19823252 | 147.43358 | 7.22E-31 | 2.14E-29 |
| PAR12 | 33.5320406 | -0.8732222 | 0.26060294 | 33.1462804 | 1.11E-06 | 6.09E-06 |
| GPR39 | 7.45543148 | -0.8737985 | 0.66864914 | 50.0160797 | 3.58E-10 | 2.81E-09 |
| EMX1 | 48.2829453 | -0.8748216 | 0.22866284 | 92.3918119 | 4.09E-19 | 6.34E-18 |
| DSG2 | 486.343084 | -0.8750498 | 0.25061872 | 28.9884052 | 7.86E-06 | 3.88E-05 |
| OVOL1.1 | 162.349063 | -0.877073 | 0.12238991 | 95.3627904 | 9.54E-20 | 1.53E-18 |
| CMPK2 | 17.344299 | -0.8787191 | 0.42154317 | 106.627075 | 3.81E-22 | 6.98E-21 |
| ABLM1 | 39.7337219 | -0.8789854 | 0.24938501 | 25.0126991 | 5.00E-05 | 0.0002206 |
| HECA2.6 | 17.6589214 | -0.8800267 | 0.48366271 | 192.071975 | 1.90E-40 | 9.15E-39 |
| GCNT3 | 52.5526327 | -0.8801837 | 0.23256163 | 235.785761 | 7.50E-50 | 6.41E-48 |
| SVIL | 53.1539123 | -0.8812183 | 0.22139644 | 90.9706358 | 8.19E-19 | 1.25E-17 |
| AHNK.1 | 958.547409 | -0.8818011 | 0.20075477 | 48.360795 | 7.94E-10 | 6.02E-09 |
| IRF1 | 108.171047 | -0.8818324 | 0.20002317 | 239.940304 | 9.56E-51 | 8.81E-49 |
| LBH.1 | 11.0012838 | -0.8843184 | 0.58225997 | 25.6575182 | 3.71E-05 | 0.00016684 |
| ARHG4 | 11.8648365 | -0.8853395 | 0.42899529 | 24.6239329 | 5.99E-05 | 0.0002604 |
| RHG27 | 55.5562793 | -0.8866844 | 0.2513797 | 136.654077 | 1.47E-28 | 3.85E-27 |
| STA10.1 | 13.2891281 | -0.8870872 | 0.40295341 | 26.1356157 | 2.97E-05 | 0.00013599 |
| OCLN.2 | 826.62258 | -0.8873428 | 0.13969953 | 169.02278 | 1.70E-35 | 6.24E-34 |
| IF | 175.116048 | -0.8894747 | 0.2037344 | 285.373888 | 1.55E-60 | 2.30E-58 |
| DIAP3 | 28.9945139 | -0.8897562 | 0.2991986 | 43.7643726 | 7.18E-09 | 4.95E-08 |
| UPAR.5 | 53.9131193 | -0.8907182 | 0.31784108 | 168.225926 | 2.51E-35 | 9.20E-34 |
| ADPGK | 47.9295521 | -0.8915085 | 0.21662237 | 23.1068478 | 0.00012055 | 0.00049893 |
| TMPS6.2 | 9.78127961 | -0.8934522 | 0.77204411 | 53.2484299 | 7.56E-11 | 6.25E-10 |
| ICA1L | 48.5494206 | -0.893962 | 0.24295364 | 49.9236404 | 3.75E-10 | 2.93E-09 |
| TMM52 | 234.402307 | -0.8953607 | 0.21277873 | 332.909495 | 8.58E-71 | 1.96E-68 |
| GA45A | 86.4216819 | -0.8985661 | 0.24084809 | 68.0392891 | 5.89E-14 | 6.19E-13 |
| STX11 | 24.8548267 | -0.8988814 | 0.30628686 | 30.0909454 | 4.69E-06 | 2.38E-05 |
| C1GLT | 51.8247241 | -0.8990309 | 0.20792276 | 77.613841 | 5.58E-16 | 6.92E-15 |
| CO6A6.1 | 73.5603099 | -0.8995315 | 0.21713124 | 76.2772267 | 1.07E-15 | 1.29E-14 |
| TNAP2.1 | 17.9944103 | -0.9002739 | 0.41600619 | 91.8814881 | 5.24E-19 | 8.05E-18 |
| SUSD6 | 92.6128878 | -0.9014138 | 0.17203801 | 85.0123101 | 1.51E-17 | 2.09E-16 |
| DOXA2 | 81.9071649 | -0.9016173 | 0.2772379 | 194.7817 | 4.97E-41 | 2.52E-39 |
| A33.15 | 10.9359143 | -0.9042795 | 0.66614166 | 32.4712879 | 1.53E-06 | 8.22E-06 |
| ANKK1.2 | 56.8149884 | -0.9052017 | 0.21470596 | 48.8957806 | 6.14E-10 | 4.72E-09 |
| PINLY | 161.634883 | -0.9065559 | 0.1481334 | 250.04174 | 6.38E-53 | 6.51E-51 |
| TYRO3 | 89.366375 | -0.907718 | 0.15680134 | 51.8346201 | 1.49E-10 | 1.20E-09 |
| UCP2 | 101.862915 | -0.9103583 | 0.23232318 | 67.9443026 | 6.16E-14 | 6.47E-13 |
| STAT1 | 64.8883763 | -0.9104226 | 0.23625272 | 46.5437235 | 1.90E-09 | 1.38E-08 |
| TDR15.1 | 16.8014992 | -0.9156602 | 0.37735312 | 45.041619 | 3.90E-09 | 2.76E-08 |
| PRDM8.1 | 8.13423637 | -0.9159584 | 0.73852111 | 51.7657226 | 1.54E-10 | 1.24E-09 |
| RB27A | 324.967421 | -0.9166348 | 0.11010984 | 107.418446 | 2.58E-22 | 4.76E-21 |
| TM117 | 12.9076669 | -0.917366 | 0.42272487 | 36.246655 | 2.57E-07 | 1.51E-06 |
| CY561 | 32.3511986 | -0.9176012 | 0.29194457 | 124.890417 | 4.82E-26 | 1.09E-24 |
| CP8B1 | 79.7578295 | -0.918399 | 0.20915183 | 65.551304 | 1.97E-13 | 1.97E-12 |
| RBM47 | 66.6923476 | -0.9201888 | 0.20797488 | 303.422366 | 1.98E-64 | 3.40E-62 |
| RN225 | 96.7932177 | -0.9210517 | 0.23935482 | 208.685932 | 5.09E-44 | 2.93E-42 |
| RDH12.2 | 158.8161 | -0.9229412 | 0.17732074 | 217.986404 | 5.08E-46 | 3.37E-44 |
| DYR.1 | 15.8879437 | -0.9230632 | 0.40197875 | 34.3844804 | 6.21E-07 | 3.51E-06 |
| BCAS3 | 904.128153 | -0.9245651 | 0.26260397 | 36.7192996 | 2.06E-07 | 1.22E-06 |
| CPLX2 | 22.4809683 | -0.9246256 | 0.37423796 | 36.2510084 | 2.57E-07 | 1.50E-06 |
| SYNE2 | 97.3433758 | -0.9277931 | 0.21126016 | 49.5958414 | 4.39E-10 | 3.41E-09 |
| PEPL | 892.057029 | -0.9288823 | 0.11373438 | 241.494326 | 4.42E-51 | 4.15E-49 |
| TRI66 | 41.6087807 | -0.9314604 | 0.23686479 | 78.6312819 | 3.40E-16 | 4.27E-15 |
| PGDH | 226.631229 | -0.9318003 | 0.16723113 | 375.014551 | 6.95E-80 | 2.05E-77 |
| GLNA | 654.183365 | -0.9336373 | 0.14964985 | 138.216417 | 6.80E-29 | 1.81E-27 |
| ES8L2 | 32.9752359 | -0.9367781 | 0.30310963 | 46.6732981 | 1.78E-09 | 1.30E-08 |
| GNA14 | 27.5449767 | -0.9381278 | 0.30702861 | 55.3308817 | 2.77E-11 | 2.40E-10 |
| GBP1.3 | 18.6999182 | -0.9382759 | 0.38741417 | 87.6748024 | 4.10E-18 | 5.90E-17 |
| ERRFI | 46.6971021 | -0.9384214 | 0.22697735 | 85.1723407 | 1.39E-17 | 1.94E-16 |
| UBP47 | 48.1938618 | -0.9393674 | 0.24055847 | 222.364185 | 5.81E-47 | 4.04E-45 |
| CCKAR.1 | 31.0082509 | -0.9407691 | 0.35890247 | 191.411806 | 2.64E-40 | 1.26E-38 |
| PK3CG.2 | 5.09140715 | -0.943043 | 0.84387674 | 28.0840477 | 1.20E-05 | 5.77E-05 |
| OTOAN | 409.500506 | -0.9433286 | 0.1357734 | 136.306914 | 1.74E-28 | 4.54E-27 |
| TTPA | 39.4874124 | -0.9440446 | 0.3228172 | 207.564332 | 8.88E-44 | 5.07E-42 |
| MSLNL | 555.645198 | -0.9475202 | 0.15297553 | 217.560627 | 6.28E-46 | 4.14E-44 |
| UROM | 17676.9668 | -0.9482733 | 0.26058433 | 149.293291 | 2.89E-31 | 8.66E-30 |
| ABHEB | 321.410827 | -0.951679 | 0.15781495 | 86.1893801 | 8.48E-18 | 1.19E-16 |
| SCLY | 191.847655 | -0.9533228 | 0.12863837 | 281.417976 | 1.10E-59 | 1.53E-57 |
| MALD3.1 | 95.934702 | -0.9545847 | 0.20486948 | 165.171198 | 1.14E-34 | 4.06E-33 |
| GIPC1 | 239.611358 | -0.9562541 | 0.14303215 | 173.760338 | 1.63E-36 | 6.30E-35 |
| S35A2 | 255.034375 | -0.9562977 | 0.11387766 | 226.745885 | 6.62E-48 | 4.97E-46 |
| EFC4A | 22.345621 | -0.9563658 | 0.37679838 | 139.244385 | 4.10E-29 | 1.09E-27 |
| CLD23 | 139.982465 | -0.9569812 | 0.16597251 | 286.495102 | 8.86E-61 | 1.33E-58 |
| ABCA3 | 8.26445477 | -0.9582549 | 0.58998238 | 27.1439726 | 1.86E-05 | 8.75E-05 |
| EPDR1 | 963.392363 | -0.9590243 | 0.14959803 | 465.35322 | 2.08E-99 | 8.17E-97 |
| RASEF.1 | 133.90589 | -0.9598592 | 0.15775032 | 133.898129 | 5.71E-28 | 1.44E-26 |
| EPGN | 54.1662123 | -0.9614624 | 0.30184077 | 77.7808905 | 5.14E-16 | 6.40E-15 |
| S39A4 | 157.911934 | -0.9631723 | 0.23578441 | 184.827025 | 6.85E-39 | 3.07E-37 |
| GSTO1 | 328.440061 | -0.9648204 | 0.12812111 | 253.587366 | 1.10E-53 | 1.15E-51 |
| AP1M1.1 | 96.637462 | -0.9659972 | 0.1730099 | 56.4413205 | 1.62E-11 | 1.42E-10 |
| RAC1 | 24.6909639 | -0.9661221 | 0.38855545 | 82.0744596 | 6.33E-17 | 8.41E-16 |
| PERT.1 | 334.989596 | -0.9677437 | 0.22448454 | 542.269713 | 4.81E-116 | 3.88E-113 |
| LIN1.13 | 104.832125 | -0.9680256 | 0.22735882 | 77.0031322 | 7.51E-16 | 9.20E-15 |
| PLEK2 | 35.0768624 | -0.971377 | 0.30586598 | 69.1397101 | 3.45E-14 | 3.69E-13 |
| P4K2B | 51.2339429 | -0.9724512 | 0.23805879 | 79.5000939 | 2.22E-16 | 2.83E-15 |
| GLSK.1 | 43.0066184 | -0.9755107 | 0.30850922 | 81.2379411 | 9.52E-17 | 1.24E-15 |
| ST1A1 | 853.711598 | -0.9769963 | 0.17972271 | 148.308953 | 4.69E-31 | 1.39E-29 |
| PDLI2 | 352.69935 | -0.9775016 | 0.11305572 | 128.21236 | 9.39E-27 | 2.24E-25 |
| MEDAG | 146.168342 | -0.9775254 | 0.16161746 | 197.603471 | 1.23E-41 | 6.43E-40 |
| STYK1 | 93.2130447 | -0.9789662 | 0.23089853 | 180.480194 | 5.88E-38 | 2.44E-36 |
| FLVC2 | 151.855548 | -0.9803683 | 0.13937504 | 220.645716 | 1.36E-46 | 9.39E-45 |
| NAL12.4 | 98.687327 | -0.9816011 | 0.24003473 | 30.0374289 | 4.81E-06 | 2.43E-05 |
| UPK1B | 573.205916 | -0.9820004 | 0.16379148 | 231.755488 | 5.53E-49 | 4.38E-47 |
| NAL12.6 | 30.1966363 | -0.9820778 | 0.34584482 | 40.6538852 | 3.17E-08 | 2.06E-07 |
| ITA11 | 11.066336 | -0.9829369 | 0.51915511 | 22.8732821 | 0.00013422 | 0.00055223 |
| AKD1B | 50.5277599 | -0.9848185 | 0.27358297 | 73.6774418 | 3.79E-15 | 4.41E-14 |
| FCGBP | 2465.51237 | -0.9859757 | 0.15146773 | 927.023012 | 2.33E-199 | 1.19E-195 |
| IFI6 | 50.0917226 | -0.9886759 | 0.37630745 | 89.3721613 | 1.79E-18 | 2.65E-17 |
| HCK | 187.046149 | -0.9911765 | 0.14684856 | 143.712965 | 4.52E-30 | 1.28E-28 |
| MANS1 | 204.681301 | -0.9913288 | 0.20872644 | 78.1287359 | 4.34E-16 | 5.41E-15 |
| PTPRH | 20.8448266 | -0.9915474 | 0.33090972 | 29.2191621 | 7.06E-06 | 3.51E-05 |
| CXB7 | 27.9238465 | -0.9968488 | 0.3286174 | 71.3083136 | 1.20E-14 | 1.34E-13 |
| TRPV3.1 | 17.8587062 | -0.9982401 | 0.43473757 | 106.642297 | 3.78E-22 | 6.93E-21 |
| MMP20 | 81.4637824 | -0.9993502 | 0.48473648 | 534.951003 | 1.84E-114 | 1.41E-111 |
| NALP5 | 43.5033196 | -1.0004052 | 0.28399097 | 49.2476037 | 5.18E-10 | 4.01E-09 |
| TRI36 | 58.2922628 | -1.0032853 | 0.24566762 | 41.8461017 | 1.80E-08 | 1.19E-07 |
| PRSR2 | 45.956611 | -1.0044608 | 0.24156191 | 34.725758 | 5.29E-07 | 3.01E-06 |
| AAAD | 31.9678363 | -1.0052406 | 0.30723218 | 27.499897 | 1.57E-05 | 7.49E-05 |
| ABCAD.1 | 275.439185 | -1.0067133 | 0.20739555 | 183.731041 | 1.18E-38 | 5.18E-37 |
| NRAM1 | 22.423047 | -1.0077182 | 0.33961988 | 27.8615314 | 1.33E-05 | 6.38E-05 |
| SORL | 37.6380905 | -1.0084006 | 0.24589884 | 66.8900822 | 1.03E-13 | 1.05E-12 |
| SORT.1 | 174.129322 | -1.0087411 | 0.18732008 | 269.477273 | 4.13E-57 | 5.23E-55 |
| K132L | 174.569159 | -1.0089301 | 0.19093691 | 309.275638 | 1.08E-65 | 1.97E-63 |
| ABCG2 | 51.0077541 | -1.0094244 | 0.2149788 | 49.3850026 | 4.85E-10 | 3.77E-09 |
| PI2R | 26.408477 | -1.0095272 | 0.31669316 | 58.2286917 | 6.83E-12 | 6.15E-11 |
| RAB7A.1 | 39.618148 | -1.0107562 | 0.26057276 | 32.5442562 | 1.48E-06 | 7.95E-06 |
| CNFN | 588.706528 | -1.0111099 | 0.19643491 | 278.983492 | 3.69E-59 | 5.04E-57 |
| FPS | 89.3219501 | -1.0124582 | 0.17050954 | 175.399738 | 7.25E-37 | 2.86E-35 |
| GLT17 | 16.8968724 | -1.0126441 | 0.41613066 | 90.6860875 | 9.41E-19 | 1.43E-17 |
| RN213.1 | 16.7832486 | -1.0130449 | 0.37596625 | 24.4235963 | 6.57E-05 | 0.00028317 |
| RGPA2 | 15.006422 | -1.0145881 | 0.39086042 | 24.1914135 | 7.31E-05 | 0.00031382 |
| S45A3 | 49.4922833 | -1.0161346 | 0.24260734 | 43.5579577 | 7.93E-09 | 5.42E-08 |
| NUCKS | 1868.91792 | -1.0166504 | 0.18867237 | 401.716197 | 1.18E-85 | 3.86E-83 |
| S14L2 | 23.3527468 | -1.0182505 | 0.32438484 | 46.8382879 | 1.65E-09 | 1.21E-08 |
| RELN | 7.61475035 | -1.0185208 | 0.9213724 | 78.5386647 | 3.55E-16 | 4.46E-15 |
| TSN1 | 576.027484 | -1.0200681 | 0.24447625 | 83.3236028 | 3.44E-17 | 4.66E-16 |
| I22R1 | 69.2617359 | -1.0233342 | 0.22456255 | 206.521512 | 1.49E-43 | 8.31E-42 |
| IL8.3 | 89.4019032 | -1.0264816 | 0.2166856 | 51.8980803 | 1.45E-10 | 1.17E-09 |
| F200A.3 | 87.3076317 | -1.0271661 | 0.20285424 | 40.6491251 | 3.18E-08 | 2.06E-07 |
| UROM.2 | 47.1769017 | -1.0273506 | 0.37835548 | 83.3230426 | 3.44E-17 | 4.66E-16 |
| MYOME | 156.61659 | -1.0282427 | 0.12966944 | 247.217601 | 2.59E-52 | 2.56E-50 |
| CRUM3 | 21.5597144 | -1.0287689 | 0.39448999 | 36.4628597 | 2.32E-07 | 1.37E-06 |
| MSLNL.3 | 251.615917 | -1.0300534 | 0.23811317 | 177.499212 | 2.57E-37 | 1.03E-35 |
| MUC16.2 | 494.246555 | -1.0308833 | 0.15256795 | 222.518509 | 5.38E-47 | 3.77E-45 |
| SPTC3 | 145.88797 | -1.0314083 | 0.13304714 | 125.556087 | 3.47E-26 | 7.94E-25 |
| TSEAR | 9.21669291 | -1.0325995 | 0.52795248 | 26.3641626 | 2.67E-05 | 0.0001231 |
| AGRE2 | 469.799491 | -1.0328107 | 0.19815695 | 192.337568 | 1.67E-40 | 8.05E-39 |
| F102A.1 | 67.3539854 | -1.0331427 | 0.19723123 | 221.644833 | 8.30E-47 | 5.75E-45 |
| PKHF1 | 41.1123552 | -1.0334866 | 0.2863755 | 135.244434 | 2.94E-28 | 7.54E-27 |
| COQ8A.1 | 14.7449853 | -1.033813 | 0.46451359 | 35.2802159 | 4.07E-07 | 2.34E-06 |
| COAA1 | 532.082019 | -1.0346991 | 0.2893492 | 1529.11399 | 0 | 0 |
| CRBG3 | 32.6928879 | -1.0348433 | 0.29268328 | 137.475182 | 9.80E-29 | 2.59E-27 |
| RN128 | 175.10261 | -1.0355144 | 0.23858345 | 140.133836 | 2.64E-29 | 7.18E-28 |
| BICL2 | 34.3654422 | -1.0358306 | 0.30575274 | 87.0435749 | 5.59E-18 | 7.94E-17 |
| TM125.1 | 34.5581167 | -1.0370162 | 0.28752441 | 126.910756 | 1.78E-26 | 4.17E-25 |
| CHSS1 | 89.1149429 | -1.0382945 | 0.24997818 | 26.4071558 | 2.62E-05 | 0.0001207 |
| T53I1 | 32.5107316 | -1.0391952 | 0.29415643 | 33.1980662 | 1.09E-06 | 5.96E-06 |
| NPDC1 | 394.225988 | -1.0392364 | 0.1229363 | 243.320486 | 1.79E-51 | 1.72E-49 |
| F16P1 | 161.637155 | -1.0394447 | 0.1653387 | 99.4790294 | 1.27E-20 | 2.12E-19 |
| GDPD5 | 20.9607144 | -1.0403551 | 0.36727487 | 48.3201348 | 8.09E-10 | 6.13E-09 |
| ITPK1 | 58.008064 | -1.0405103 | 0.22524654 | 53.7193151 | 6.03E-11 | 5.04E-10 |
| GARE1.1 | 30.0653228 | -1.0417767 | 0.36586348 | 158.145495 | 3.65E-33 | 1.21E-31 |
| MX3.1 | 43.9490188 | -1.0421378 | 0.24396979 | 110.307409 | 6.26E-23 | 1.20E-21 |
| GMDS | 682.486948 | -1.0428511 | 0.136585 | 157.639645 | 4.69E-33 | 1.55E-31 |
| S26A4.1 | 106.83691 | -1.0431853 | 0.23856257 | 168.683477 | 2.00E-35 | 7.36E-34 |
| DUOX2 | 191.075074 | -1.045454 | 0.18747433 | 478.562035 | 2.90E-102 | 1.31E-99 |
| BCAT1 | 176.952906 | -1.0461913 | 0.21416992 | 89.5411406 | 1.65E-18 | 2.44E-17 |
| STX2.1 | 18.0187692 | -1.0467392 | 0.42408151 | 81.1858398 | 9.77E-17 | 1.27E-15 |
| KALRN.1 | 47.2986245 | -1.0499624 | 0.21982871 | 27.3375276 | 1.70E-05 | 8.03E-05 |
| GTR2 | 26.6996131 | -1.05293 | 0.31514274 | 74.2902948 | 2.82E-15 | 3.32E-14 |
| TM45B.1 | 102.211496 | -1.0532754 | 0.22903485 | 211.139091 | 1.51E-44 | 8.86E-43 |
| S39AE | 105.599474 | -1.0543161 | 0.16694756 | 60.3120361 | 2.49E-12 | 2.32E-11 |
| PGDH.3 | 797.377664 | -1.055151 | 0.17634085 | 171.335361 | 5.41E-36 | 2.05E-34 |
| GCH1 | 8.70443574 | -1.0551643 | 0.53801168 | 24.1718297 | 7.38E-05 | 0.00031632 |
| F166B | 21.5087845 | -1.0555145 | 0.41075711 | 34.8510597 | 4.98E-07 | 2.84E-06 |
| STA10 | 98.5422789 | -1.0555844 | 0.19407677 | 153.983202 | 2.85E-32 | 8.82E-31 |
| RELL1 | 28.7027449 | -1.0560682 | 0.27872553 | 29.1810934 | 7.18E-06 | 3.57E-05 |
| OLFM4.1 | 1668.03591 | -1.0568171 | 0.23472255 | 155.543283 | 1.32E-32 | 4.22E-31 |
| SCEL | 1168.20949 | -1.0598606 | 0.11811188 | 452.476247 | 1.27E-96 | 4.61E-94 |
| SAM9L.1 | 26.9924674 | -1.0606629 | 0.29500747 | 56.4543805 | 1.61E-11 | 1.41E-10 |
| HA2D.1 | 821.67334 | -1.0627202 | 0.23208981 | 232.211839 | 4.41E-49 | 3.52E-47 |
| FOXQ1 | 205.793213 | -1.0628055 | 0.20859038 | 156.876577 | 6.83E-33 | 2.24E-31 |
| A4GCT.2 | 1185.51194 | -1.0633971 | 0.1714151 | 198.710941 | 7.11E-42 | 3.75E-40 |
| CD276.3 | 27.530248 | -1.064061 | 0.29643348 | 33.8080373 | 8.16E-07 | 4.54E-06 |
| BCAP.1 | 22.5815605 | -1.066443 | 0.480863 | 75.1032674 | 1.89E-15 | 2.25E-14 |
| CANT1 | 69.6301297 | -1.0668104 | 0.20548713 | 93.8751636 | 1.98E-19 | 3.11E-18 |
| UPK2 | 2470.37399 | -1.0732213 | 0.18103609 | 206.679251 | 1.38E-43 | 7.71E-42 |
| CAN2 | 1181.67427 | -1.0747751 | 0.11733062 | 203.360926 | 7.12E-43 | 3.88E-41 |
| CD3G | 5.17527082 | -1.0756502 | 1.14399446 | 23.9073927 | 8.34E-05 | 0.00035411 |
| ACBG2 | 77.7150696 | -1.0759377 | 0.18902319 | 92.0893684 | 4.74E-19 | 7.30E-18 |
| MUC1 | 22.249641 | -1.0770841 | 0.4761049 | 161.912624 | 5.68E-34 | 1.97E-32 |
| CD276.10 | 102.182984 | -1.0781021 | 0.31672105 | 67.3005895 | 8.42E-14 | 8.68E-13 |
| CLD4.1 | 4400.95151 | -1.0793807 | 0.12960426 | 198.0396 | 9.92E-42 | 5.20E-40 |
| MLP | 9.45501133 | -1.0794882 | 0.55067705 | 36.5019196 | 2.28E-07 | 1.35E-06 |
| ZA2G | 447.840536 | -1.079647 | 0.22598896 | 254.613143 | 6.60E-54 | 6.97E-52 |
| MUC19.3 | 5122.2664 | -1.0813675 | 0.22069332 | 306.577462 | 4.13E-65 | 7.35E-63 |
| NLRX1 | 26.8339232 | -1.082104 | 0.35233903 | 58.8960792 | 4.95E-12 | 4.49E-11 |
| DHX58 | 11.5422748 | -1.0834592 | 0.51674082 | 36.2919009 | 2.52E-07 | 1.48E-06 |
| SHRM2 | 87.4655705 | -1.0860375 | 0.16281676 | 64.1836253 | 3.82E-13 | 3.74E-12 |
| TMPS9 | 104.541565 | -1.0866211 | 0.20873732 | 273.91331 | 4.57E-58 | 5.93E-56 |
| ESYT3 | 65.4413756 | -1.0880085 | 0.20764244 | 117.296648 | 2.02E-24 | 4.24E-23 |
| TBA3 | 42.1008362 | -1.0881719 | 0.31171458 | 101.918989 | 3.84E-21 | 6.61E-20 |
| FGL1L.1 | 3878.14852 | -1.0886012 | 0.30028057 | 121.587753 | 2.45E-25 | 5.33E-24 |
| CASC4 | 1365.9192 | -1.0886775 | 0.10875727 | 377.985927 | 1.59E-80 | 4.85E-78 |
| RGS2.1 | 204.308621 | -1.0920779 | 0.23367879 | 211.045673 | 1.58E-44 | 9.25E-43 |
| TGM2.1 | 1818.50677 | -1.0927944 | 0.12664833 | 158.683872 | 2.80E-33 | 9.36E-32 |
| GBB3 | 34.0081304 | -1.0957956 | 0.48926182 | 309.655778 | 8.95E-66 | 1.65E-63 |
| K1C19 | 1879.89367 | -1.0963727 | 0.22655243 | 184.187504 | 9.40E-39 | 4.17E-37 |
| CLTR1.1 | 28.0393479 | -1.0982352 | 0.33650109 | 66.7077628 | 1.12E-13 | 1.14E-12 |
| CEAM6 | 149.935035 | -1.0983878 | 0.18322425 | 105.799529 | 5.72E-22 | 1.04E-20 |
| HS3S4 | 8.3630912 | -1.0984419 | 0.64125214 | 25.129447 | 4.74E-05 | 0.00020965 |
| EMX2 | 54.2315769 | -1.0985244 | 0.22994515 | 82.5278429 | 5.07E-17 | 6.81E-16 |
| HEXB | 732.815732 | -1.0998379 | 0.0961999 | 224.837854 | 1.71E-47 | 1.23E-45 |
| LRRF2.3 | 138.459609 | -1.1004697 | 0.23814487 | 53.4751317 | 6.78E-11 | 5.63E-10 |
| STXA | 15.2163194 | -1.1009037 | 0.41066202 | 45.3260854 | 3.40E-09 | 2.41E-08 |
| HBEGF.1 | 58.8773733 | -1.1061007 | 0.21330868 | 102.213338 | 3.32E-21 | 5.74E-20 |
| LORF2.398 | 42.6601859 | -1.1062804 | 0.34955763 | 31.6615731 | 2.24E-06 | 1.18E-05 |
| LRC19 | 27.8919969 | -1.1067642 | 0.37074093 | 65.558878 | 1.96E-13 | 1.96E-12 |
| OXDD | 20.7013491 | -1.1068163 | 0.41349972 | 39.8429236 | 4.66E-08 | 2.96E-07 |
| S6A14.3 | 265.891536 | -1.1085296 | 0.20803835 | 148.384602 | 4.52E-31 | 1.35E-29 |
| PARPT | 15.2606343 | -1.1096874 | 0.41175028 | 23.5416984 | 9.87E-05 | 0.00041421 |
| FGL1L | 2311.53329 | -1.1113388 | 0.29937562 | 120.879082 | 3.47E-25 | 7.49E-24 |
| UROM.1 | 19.3262959 | -1.1120892 | 0.53147565 | 59.1172419 | 4.45E-12 | 4.04E-11 |
| NPL | 144.437393 | -1.112574 | 0.19598622 | 58.5387408 | 5.88E-12 | 5.31E-11 |
| ZN239.40 | 254.656749 | -1.11383 | 0.2275926 | 32.0350609 | 1.88E-06 | 9.99E-06 |
| RAB7B | 44.3478141 | -1.1140907 | 0.23837163 | 72.1414018 | 8.01E-15 | 9.08E-14 |
| ACT2 | 13.5362288 | -1.1145337 | 0.49595765 | 62.6489076 | 8.04E-13 | 7.70E-12 |
| CK052 | 48.7976785 | -1.1145835 | 0.23299825 | 89.2285328 | 1.92E-18 | 2.83E-17 |
| MUC18.2 | 1381.84826 | -1.1160146 | 0.11304955 | 263.039642 | 1.01E-55 | 1.19E-53 |
| GPC6A.4 | 16.232866 | -1.1160273 | 0.53206429 | 139.713331 | 3.25E-29 | 8.76E-28 |
| B4GT4 | 19.8906319 | -1.1193447 | 0.38107397 | 55.0171818 | 3.22E-11 | 2.77E-10 |
| MUC17 | 12.8799948 | -1.1204991 | 0.56881304 | 81.2475366 | 9.48E-17 | 1.24E-15 |
| SYTL2.2 | 450.351208 | -1.1212145 | 0.18690026 | 259.663557 | 5.39E-55 | 5.89E-53 |
| LPAR5.1 | 87.1940894 | -1.1223495 | 0.33043476 | 29.0188029 | 7.75E-06 | 3.83E-05 |
| TMPS4 | 134.893771 | -1.123466 | 0.17503173 | 167.404681 | 3.77E-35 | 1.37E-33 |
| IGL1.2 | 10.8865817 | -1.1281857 | 0.96413939 | 28.3866966 | 1.04E-05 | 5.07E-05 |
| SAM9L.2 | 73.7658714 | -1.1334151 | 0.24507887 | 146.657071 | 1.06E-30 | 3.11E-29 |
| LORF2.417 | 9.41138449 | -1.1350178 | 0.64572471 | 46.3285149 | 2.10E-09 | 1.52E-08 |
| VWA3B | 13.4296301 | -1.137588 | 0.4959844 | 25.4184412 | 4.14E-05 | 0.00018525 |
| RFIP1 | 141.978439 | -1.1401767 | 0.16330835 | 177.412869 | 2.68E-37 | 1.08E-35 |
| SIM14 | 94.9324001 | -1.1421845 | 0.2140321 | 74.9476571 | 2.04E-15 | 2.42E-14 |
| AA1R | 4.42653905 | -1.142816 | 1.05900174 | 27.3298302 | 1.70E-05 | 8.06E-05 |
| EDEM3.1 | 14.0616526 | -1.1435379 | 0.44162261 | 25.4526973 | 4.08E-05 | 0.00018244 |
| ANKK1 | 74.0229113 | -1.1436396 | 0.19982316 | 141.444324 | 1.38E-29 | 3.82E-28 |
| F177A | 65.9866631 | -1.1472219 | 0.2586973 | 120.201252 | 4.84E-25 | 1.04E-23 |
| RSAD2 | 26.8437975 | -1.1476359 | 0.4314178 | 70.8972718 | 1.47E-14 | 1.62E-13 |
| FUCL5.1 | 4481.59159 | -1.1480186 | 0.59801751 | 90.383976 | 1.09E-18 | 1.64E-17 |
| AGRG6.1 | 14.3841759 | -1.1480908 | 0.5767666 | 135.459623 | 2.65E-28 | 6.82E-27 |
| CD83.1 | 44.6257607 | -1.1493958 | 0.30376139 | 29.3255611 | 6.71E-06 | 3.35E-05 |
| A33.28 | 21.7069478 | -1.1496068 | 0.48037193 | 113.468727 | 1.32E-23 | 2.65E-22 |
| MYO15 | 7.28294333 | -1.1501333 | 0.74192421 | 37.5215052 | 1.41E-07 | 8.44E-07 |
| KEAP1 | 35.7532337 | -1.1522863 | 0.27254457 | 32.7431424 | 1.35E-06 | 7.28E-06 |
| GRHL1 | 162.44209 | -1.1526177 | 0.16728603 | 291.943688 | 5.92E-62 | 9.25E-60 |
| AT2A3 | 458.480788 | -1.1544637 | 0.13812139 | 232.428728 | 3.96E-49 | 3.21E-47 |
| PR15L | 95.5014825 | -1.1589942 | 0.2140965 | 193.701697 | 8.49E-41 | 4.19E-39 |
| FGFR3 | 47.9963919 | -1.1603024 | 0.26468781 | 54.3976622 | 4.34E-11 | 3.68E-10 |
| FOXR1 | 17.1580153 | -1.1605188 | 0.45588382 | 47.1484828 | 1.42E-09 | 1.05E-08 |
| PITC1.1 | 258.151974 | -1.1616182 | 0.22456038 | 59.9455186 | 2.98E-12 | 2.75E-11 |
| UPK3B | 966.465441 | -1.1620626 | 0.14878074 | 194.325839 | 6.23E-41 | 3.12E-39 |
| TRI25.6 | 56.64021 | -1.1622852 | 0.53951947 | 216.130936 | 1.27E-45 | 8.23E-44 |
| PAR6B.1 | 23.141128 | -1.1626263 | 0.34258872 | 81.024769 | 1.06E-16 | 1.37E-15 |
| NLRP3.18 | 63.1634489 | -1.1669075 | 0.26622182 | 30.7034803 | 3.52E-06 | 1.81E-05 |
| PRSS8 | 152.447422 | -1.1678581 | 0.1633077 | 160.420612 | 1.19E-33 | 4.02E-32 |
| MA7D2 | 13.4397588 | -1.173883 | 0.58828021 | 118.749154 | 9.88E-25 | 2.09E-23 |
| PPM1H.2 | 47.1778746 | -1.17484 | 0.23548635 | 75.5372589 | 1.53E-15 | 1.84E-14 |
| VGLL1 | 28.2085305 | -1.1766332 | 0.37983809 | 106.359195 | 4.35E-22 | 7.94E-21 |
| KCNK1.1 | 143.529756 | -1.1811612 | 0.20212203 | 87.5510234 | 4.36E-18 | 6.26E-17 |
| ILK1 | 37.8442704 | -1.1822273 | 0.38133822 | 50.4298629 | 2.94E-10 | 2.32E-09 |
| AVID | 154.765021 | -1.1830427 | 0.32465802 | 65.4704216 | 2.05E-13 | 2.04E-12 |
| TRPM5 | 58.8977441 | -1.1875006 | 0.24478737 | 266.395459 | 1.91E-56 | 2.34E-54 |
| MUC2 | 216.114388 | -1.1883925 | 0.29860923 | 135.734776 | 2.31E-28 | 5.98E-27 |
| ATP9A | 28.6105336 | -1.1886311 | 0.30086236 | 47.2513422 | 1.35E-09 | 1.00E-08 |
| MCF2L | 17.9215843 | -1.1896003 | 0.42617107 | 47.3156122 | 1.31E-09 | 9.71E-09 |
| ARH37 | 9.545908 | -1.1966592 | 0.57031159 | 39.6811255 | 5.04E-08 | 3.19E-07 |
| CLD4 | 202.637351 | -1.1993269 | 0.29237845 | 136.46902 | 1.61E-28 | 4.20E-27 |
| GALT6 | 40.0715563 | -1.1998362 | 0.27786297 | 150.717861 | 1.43E-31 | 4.35E-30 |
| STAT2 | 1424.97124 | -1.2020132 | 0.20230381 | 236.947463 | 4.21E-50 | 3.69E-48 |
| ELOV7 | 115.570634 | -1.2027981 | 0.22274364 | 88.7526607 | 2.42E-18 | 3.55E-17 |
| GTR11 | 58.193674 | -1.2032676 | 0.26508233 | 358.046618 | 3.21E-76 | 8.77E-74 |
| SCUB3.1 | 307.865156 | -1.2052547 | 0.15936457 | 193.751636 | 8.28E-41 | 4.10E-39 |
| F167A | 22.9661434 | -1.2084824 | 0.44904151 | 86.4758991 | 7.38E-18 | 1.04E-16 |
| AGRF5 | 10.7708433 | -1.2092153 | 0.70636778 | 96.9767697 | 4.33E-20 | 7.03E-19 |
| CAN9 | 762.076023 | -1.2096957 | 0.12481944 | 362.092961 | 4.29E-77 | 1.22E-74 |
| SC5A8.2 | 97.1152685 | -1.210321 | 0.1991686 | 82.2113081 | 5.92E-17 | 7.89E-16 |
| GGT1 | 156.453786 | -1.2123004 | 0.18534281 | 177.892398 | 2.11E-37 | 8.61E-36 |
| LORF2.342 | 127.3846 | -1.2151642 | 0.26345567 | 271.50158 | 1.51E-57 | 1.93E-55 |
| HXK2 | 62.6457596 | -1.215296 | 0.21852525 | 146.514163 | 1.14E-30 | 3.33E-29 |
| G6PE | 15.4429848 | -1.2190236 | 0.40031431 | 23.4723882 | 0.00010188 | 0.00042669 |
| SARG | 105.048482 | -1.2199863 | 0.17291705 | 220.280144 | 1.63E-46 | 1.11E-44 |
| HMX2 | 4.49679107 | -1.2220172 | 1.06980747 | 29.3713316 | 6.57E-06 | 3.28E-05 |
| PAR14.2 | 6.3607732 | -1.2222351 | 0.71808754 | 33.1531115 | 1.11E-06 | 6.07E-06 |
| I20RA | 33.3645266 | -1.2270315 | 0.34318466 | 134.751719 | 3.75E-28 | 9.58E-27 |
| IFIT5.4 | 172.228541 | -1.2271153 | 0.35413135 | 38.4356011 | 9.11E-08 | 5.61E-07 |
| EHF | 38.8636308 | -1.2306697 | 0.3385293 | 119.907635 | 5.59E-25 | 1.20E-23 |
| TMPSD | 153.469084 | -1.2313697 | 0.23121441 | 131.213546 | 2.14E-27 | 5.28E-26 |
| PDE2A | 74.4673936 | -1.2326226 | 0.22990761 | 89.3347872 | 1.82E-18 | 2.69E-17 |
| KV13 | 8.35691503 | -1.2380476 | 0.69535428 | 42.3656565 | 1.40E-08 | 9.37E-08 |
| S6A14.1 | 65.8125976 | -1.2389288 | 0.39957591 | 87.4017343 | 4.69E-18 | 6.72E-17 |
| ILDR1 | 82.1482193 | -1.2390325 | 0.24279035 | 178.324654 | 1.71E-37 | 6.97E-36 |
| HMR1.5 | 238.857867 | -1.2453681 | 0.18576713 | 230.957714 | 8.21E-49 | 6.44E-47 |
| CPEB3 | 25.5773607 | -1.2491401 | 0.3048341 | 28.3705164 | 1.05E-05 | 5.11E-05 |
| EVPL | 669.665288 | -1.2492294 | 0.13356557 | 259.990768 | 4.58E-55 | 5.04E-53 |
| MSLNL.2 | 21.9075221 | -1.251066 | 0.402063 | 79.6888828 | 2.03E-16 | 2.60E-15 |
| LAMP3 | 425.969812 | -1.2513544 | 0.13576293 | 235.641085 | 8.06E-50 | 6.85E-48 |
| TAC2N | 210.25557 | -1.2514218 | 0.24531796 | 131.291857 | 2.06E-27 | 5.10E-26 |
| KCRU | 390.954079 | -1.2516818 | 0.19504697 | 79.7140963 | 2.00E-16 | 2.57E-15 |
| HMX3 | 5.65231203 | -1.2518748 | 0.95752328 | 35.0277311 | 4.58E-07 | 2.62E-06 |
| CYTB | 88.6110099 | -1.2520434 | 0.24444616 | 79.6469262 | 2.07E-16 | 2.65E-15 |
| PERE.1 | 985.315511 | -1.2537516 | 0.25215148 | 199.647017 | 4.47E-42 | 2.38E-40 |
| FRRS1.1 | 70.3395697 | -1.2558537 | 0.43186037 | 55.2049455 | 2.94E-11 | 2.54E-10 |
| PKHS1 | 12.3971015 | -1.2584264 | 0.52231066 | 39.545764 | 5.37E-08 | 3.38E-07 |
| DHI1L | 9.2093493 | -1.2602007 | 0.63573866 | 22.3718637 | 0.00016899 | 0.00068406 |
| MUC4 | 812.441153 | -1.2645195 | 0.2034728 | 211.175636 | 1.48E-44 | 8.74E-43 |
| ST1C1 | 82.9683925 | -1.2699947 | 0.27527756 | 61.9040413 | 1.15E-12 | 1.09E-11 |
| DGAT2 | 21.9066444 | -1.2709047 | 0.46009026 | 41.2683347 | 2.37E-08 | 1.55E-07 |
| CAR14 | 32.376259 | -1.2761532 | 0.31908163 | 129.65699 | 4.61E-27 | 1.12E-25 |
| S43A3.2 | 20.7075128 | -1.2780017 | 0.39897 | 125.134176 | 4.27E-26 | 9.75E-25 |
| GPC6A.3 | 96.5365738 | -1.2817602 | 0.24078622 | 143.495232 | 5.04E-30 | 1.41E-28 |
| AGRF5.1 | 7.42242996 | -1.282733 | 0.75630509 | 58.0626096 | 7.40E-12 | 6.65E-11 |
| CAN5.1 | 6.86320069 | -1.283146 | 0.75397492 | 27.4235077 | 1.63E-05 | 7.74E-05 |
| AL1A3 | 32.126456 | -1.2864263 | 0.38972249 | 46.9599158 | 1.55E-09 | 1.14E-08 |
| MSLN | 42.4096288 | -1.2868475 | 0.56920097 | 50.9048768 | 2.34E-10 | 1.86E-09 |
| FUT2.1 | 249.465317 | -1.2868916 | 0.16437911 | 196.408572 | 2.22E-41 | 1.15E-39 |
| EDSP | 216.320958 | -1.2877486 | 0.26937 | 41.9058097 | 1.74E-08 | 1.16E-07 |
| PGDH.2 | 13.0570599 | -1.2912833 | 0.52032949 | 57.9527151 | 7.81E-12 | 7.00E-11 |
| DEGS2 | 125.684585 | -1.2931003 | 0.18838662 | 266.035331 | 2.28E-56 | 2.75E-54 |
| DNER | 8.52283157 | -1.2945164 | 0.87560945 | 30.6219713 | 3.66E-06 | 1.87E-05 |
| NUAK2 | 75.6772877 | -1.2957088 | 0.25213022 | 59.2515002 | 4.17E-12 | 3.80E-11 |
| GP160 | 52.9516608 | -1.2971975 | 0.22941752 | 55.5394419 | 2.50E-11 | 2.18E-10 |
| CS067 | 16.6162012 | -1.3003654 | 0.5198498 | 26.1216392 | 2.99E-05 | 0.00013676 |
| MAL.1 | 330.325067 | -1.3022759 | 0.14154531 | 368.099223 | 2.17E-78 | 6.25E-76 |
| AGRG2 | 35.4317131 | -1.3034684 | 0.29408993 | 141.737715 | 1.20E-29 | 3.32E-28 |
| BEST2.1 | 5.35092252 | -1.3038011 | 0.83712831 | 23.1631668 | 0.00011747 | 0.00048696 |
| MUC5B | 3579.22261 | -1.3043285 | 0.38200014 | 177.647843 | 2.39E-37 | 9.66E-36 |
| GRB7 | 33.5142869 | -1.3048248 | 0.37542258 | 28.2599095 | 1.10E-05 | 5.35E-05 |
| FRK | 25.2126305 | -1.3062253 | 0.32839475 | 54.7125624 | 3.73E-11 | 3.18E-10 |
| P2RY4 | 5.05931067 | -1.3103971 | 1.22400298 | 49.0363252 | 5.74E-10 | 4.42E-09 |
| SL9A2 | 24.1534946 | -1.3126711 | 0.33526784 | 69.1736836 | 3.39E-14 | 3.64E-13 |
| NOS1 | 108.897456 | -1.3142052 | 0.27578504 | 137.497284 | 9.69E-29 | 2.56E-27 |
| SLAF7 | 12.5358901 | -1.3144603 | 0.80243493 | 68.9124299 | 3.85E-14 | 4.11E-13 |
| AT2C2 | 28.5222113 | -1.3159421 | 0.30142387 | 59.9359186 | 2.99E-12 | 2.76E-11 |
| RFT2.1 | 36.053592 | -1.3211196 | 0.27975075 | 82.693225 | 4.68E-17 | 6.30E-16 |
| LORF2.221 | 45.193465 | -1.3238379 | 0.31986099 | 22.7023346 | 0.0001452 | 0.00059562 |
| LORF2.310 | 42.8443504 | -1.3293139 | 0.32359856 | 25.3277604 | 4.32E-05 | 0.00019248 |
| CCD68 | 102.425942 | -1.332989 | 0.21731959 | 81.2216681 | 9.60E-17 | 1.25E-15 |
| ARAP2 | 28.8525017 | -1.3365531 | 0.31093338 | 49.6085703 | 4.36E-10 | 3.39E-09 |
| TS1R1.1 | 19.970462 | -1.3372416 | 0.6418381 | 232.275876 | 4.27E-49 | 3.42E-47 |
| SYTL5 | 81.9703857 | -1.3466573 | 0.18922597 | 90.6153543 | 9.74E-19 | 1.47E-17 |
| STAP1 | 39.8216177 | -1.3473278 | 0.30046222 | 66.9959307 | 9.77E-14 | 1.00E-12 |
| FAM3C | 264.607779 | -1.3531235 | 0.17158514 | 212.992385 | 6.04E-45 | 3.68E-43 |
| GGH.1 | 27.9387783 | -1.3542886 | 0.35711239 | 81.5363137 | 8.23E-17 | 1.08E-15 |
| TT39A | 139.758855 | -1.3583746 | 0.14271155 | 182.943377 | 1.74E-38 | 7.48E-37 |
| CMTR2 | 349.13097 | -1.3619508 | 0.2125847 | 108.736258 | 1.35E-22 | 2.54E-21 |
| LITAF.1 | 5.57349582 | -1.3638742 | 1.28609249 | 29.658141 | 5.74E-06 | 2.89E-05 |
| RPGP2 | 10.4417834 | -1.3640279 | 0.56617787 | 44.2850702 | 5.60E-09 | 3.91E-08 |
| HARB1.14 | 2106.32243 | -1.3666012 | 0.36343033 | 205.224716 | 2.83E-43 | 1.56E-41 |
| AGRF5.6 | 48.6639791 | -1.3675969 | 0.28838334 | 108.352373 | 1.63E-22 | 3.05E-21 |
| CK053 | 7.4173579 | -1.3693261 | 0.87539201 | 60.5352078 | 2.24E-12 | 2.08E-11 |
| TRPV3 | 6.88121145 | -1.3715742 | 0.88670298 | 50.1978576 | 3.28E-10 | 2.58E-09 |
| LORF2.78 | 39.6236058 | -1.3743792 | 0.38270391 | 42.7467732 | 1.17E-08 | 7.86E-08 |
| NEUL1 | 34.5735198 | -1.3746087 | 0.31658966 | 96.5548309 | 5.32E-20 | 8.64E-19 |
| B3GN3 | 383.925987 | -1.3775631 | 0.21481789 | 142.147886 | 9.79E-30 | 2.71E-28 |
| HIP1R | 245.549807 | -1.3784111 | 0.1656872 | 168.112866 | 2.66E-35 | 9.71E-34 |
| ADAM9.2 | 84.679307 | -1.3809235 | 0.24007586 | 237.325653 | 3.49E-50 | 3.07E-48 |
| A33.39 | 51.1361464 | -1.3821908 | 0.26055336 | 71.2466377 | 1.24E-14 | 1.38E-13 |
| ANO10.1 | 49.5583035 | -1.3875543 | 0.29989209 | 156.852257 | 6.92E-33 | 2.27E-31 |
| PIRT | 12.7909517 | -1.3906214 | 0.49901841 | 37.7466685 | 1.26E-07 | 7.64E-07 |
| TCO1.1 | 77.5360844 | -1.3942723 | 0.34083588 | 117.130407 | 2.19E-24 | 4.59E-23 |
| ERBB4.1 | 15.3482462 | -1.3959669 | 0.49105097 | 55.9081619 | 2.10E-11 | 1.83E-10 |
| TRI39.1 | 45.0880518 | -1.3981708 | 0.43299316 | 78.3476377 | 3.90E-16 | 4.88E-15 |
| PEG3 | 625.0257 | -1.3999757 | 0.22433805 | 176.326494 | 4.59E-37 | 1.83E-35 |
| LORF2.195 | 100.061394 | -1.4009628 | 0.29037787 | 66.9607392 | 9.94E-14 | 1.02E-12 |
| G6PT2 | 19.9371441 | -1.4010189 | 0.42200332 | 31.9795979 | 1.93E-06 | 1.02E-05 |
| A33.1 | 14.444658 | -1.4039658 | 0.4699121 | 52.8240631 | 9.28E-11 | 7.60E-10 |
| RASL2 | 13.2921049 | -1.4043267 | 0.53091993 | 32.8599191 | 1.28E-06 | 6.92E-06 |
| IFT1B | 20.7667647 | -1.40683 | 0.50691219 | 26.9832018 | 2.00E-05 | 9.39E-05 |
| LITD1.10 | 18.0708765 | -1.4081224 | 0.47057623 | 122.261991 | 1.76E-25 | 3.87E-24 |
| MUC16.4 | 23.1683369 | -1.4108838 | 0.40079196 | 65.4434507 | 2.08E-13 | 2.07E-12 |
| DOCK8 | 14.6430674 | -1.4143164 | 0.4546286 | 39.2463253 | 6.20E-08 | 3.88E-07 |
| CO6A3.6 | 7.66622066 | -1.4226625 | 0.76307008 | 29.5729295 | 5.98E-06 | 3.00E-05 |
| STX19 | 7.43019832 | -1.4231814 | 0.70175102 | 37.1303487 | 1.69E-07 | 1.01E-06 |
| PLEC.1 | 24.1410236 | -1.4252935 | 0.49120291 | 22.2064903 | 0.00018232 | 0.00073469 |
| FA83E | 35.516281 | -1.4273341 | 0.34072044 | 105.351404 | 7.13E-22 | 1.28E-20 |
| MUC15 | 212.246 | -1.4300561 | 0.14925023 | 315.733229 | 4.37E-67 | 8.47E-65 |
| NIPA2.1 | 36.7138858 | -1.4306445 | 0.28085893 | 70.182575 | 2.08E-14 | 2.27E-13 |
| A33.22 | 4039.02307 | -1.4314808 | 0.24149731 | 260.463592 | 3.62E-55 | 4.02E-53 |
| SGPP2 | 85.413968 | -1.4318043 | 0.19843054 | 116.308776 | 3.28E-24 | 6.79E-23 |
| GPC6A.2 | 3.53761229 | -1.4386656 | 1.20264216 | 27.1435579 | 1.86E-05 | 8.75E-05 |
| IIGP5.4 | 26.8526281 | -1.4401178 | 0.42151303 | 38.6402131 | 8.27E-08 | 5.11E-07 |
| CD83 | 31.4204851 | -1.4402862 | 0.39267664 | 68.8909689 | 3.89E-14 | 4.15E-13 |
| S12A8 | 38.5878283 | -1.4410221 | 0.27740902 | 92.1577297 | 4.58E-19 | 7.08E-18 |
| IRX4 | 16.7088463 | -1.443257 | 0.81074762 | 87.0485124 | 5.58E-18 | 7.93E-17 |
| TRI32 | 5.85874791 | -1.4450853 | 0.76781076 | 22.3177591 | 0.00017324 | 0.00070015 |
| MUC19.6 | 7.00795045 | -1.4491125 | 1.34147594 | 44.8686144 | 4.23E-09 | 2.99E-08 |
| KLF3 | 82.852036 | -1.4571985 | 0.24514915 | 92.8333553 | 3.29E-19 | 5.13E-18 |
| GRHL3 | 177.47579 | -1.4733143 | 0.17723444 | 224.902688 | 1.65E-47 | 1.20E-45 |
| POL.7 | 336.350888 | -1.4745399 | 0.15162886 | 189.791096 | 5.88E-40 | 2.70E-38 |
| HYAL.2 | 4825.22677 | -1.4746972 | 0.39722738 | 227.301814 | 5.03E-48 | 3.83E-46 |
| BTD | 63.3688614 | -1.4804463 | 0.23827448 | 146.93727 | 9.22E-31 | 2.72E-29 |
| BICL1 | 24.9757261 | -1.4822327 | 0.40413894 | 69.9092897 | 2.37E-14 | 2.58E-13 |
| R144B | 52.2225626 | -1.4885686 | 0.24440596 | 104.216453 | 1.24E-21 | 2.19E-20 |
| HA2B | 10.6242042 | -1.4905126 | 0.80592451 | 28.6150029 | 9.36E-06 | 4.59E-05 |
| PK3CG.3 | 31.1517118 | -1.4927661 | 0.36866263 | 60.0291098 | 2.86E-12 | 2.64E-11 |
| UPAR | 31.3350967 | -1.4953496 | 0.45075832 | 66.8902066 | 1.03E-13 | 1.05E-12 |
| CLIP2.2 | 10.4367491 | -1.4980864 | 0.61752821 | 56.0679639 | 1.94E-11 | 1.70E-10 |
| CAH4.1 | 79.4168339 | -1.4992622 | 0.48444339 | 43.7001391 | 7.40E-09 | 5.10E-08 |
| KCNK1 | 41.768966 | -1.4995335 | 0.29264234 | 99.662618 | 1.16E-20 | 1.95E-19 |
| TECTA.4 | 10.6178137 | -1.5047011 | 0.72537339 | 62.0836478 | 1.06E-12 | 1.00E-11 |
| MGT4B | 24.729739 | -1.5090727 | 0.37531374 | 81.938742 | 6.76E-17 | 8.96E-16 |
| MF6LB | 23.0435712 | -1.5105516 | 0.37036844 | 45.0303881 | 3.92E-09 | 2.77E-08 |
| CAN5.2 | 118.922287 | -1.5190672 | 0.17928055 | 346.126345 | 1.20E-73 | 3.07E-71 |
| CSDC2 | 22.6165311 | -1.5197245 | 0.51671754 | 27.9517863 | 1.28E-05 | 6.13E-05 |
| BI2L1 | 21.6255478 | -1.5202521 | 0.38598472 | 49.1065321 | 5.55E-10 | 4.28E-09 |
| YM012.2 | 322.035528 | -1.5208127 | 0.42996157 | 50.2502013 | 3.20E-10 | 2.52E-09 |
| MAP1B.2 | 5.91525569 | -1.5230806 | 1.02439779 | 52.0904307 | 1.32E-10 | 1.07E-09 |
| TFR1 | 22.0073859 | -1.5269182 | 0.37015988 | 36.4232848 | 2.37E-07 | 1.39E-06 |
| KCD14 | 59.7477003 | -1.5349863 | 0.25380576 | 124.033841 | 7.34E-26 | 1.64E-24 |
| CHIA | 18.1719201 | -1.5498956 | 0.42718987 | 29.2271857 | 7.03E-06 | 3.50E-05 |
| TLC3B | 59.3554958 | -1.5617549 | 0.29586198 | 69.4890689 | 2.91E-14 | 3.14E-13 |
| ELF3 | 111.109095 | -1.5639592 | 0.36424277 | 116.793026 | 2.59E-24 | 5.38E-23 |
| FR1L6 | 30.4532021 | -1.5711635 | 0.59071751 | 155.306521 | 1.48E-32 | 4.71E-31 |
| BORG1.2 | 10.1891416 | -1.5733952 | 0.5681555 | 28.7274443 | 8.88E-06 | 4.36E-05 |
| HB2D | 9.56672652 | -1.5737918 | 0.72412927 | 32.5646072 | 1.47E-06 | 7.88E-06 |
| SPIR2 | 38.0942198 | -1.5804324 | 0.32676179 | 54.8121235 | 3.56E-11 | 3.04E-10 |
| POLY.3 | 25.1823284 | -1.5817046 | 0.52045719 | 21.7198689 | 0.00022786 | 0.00090418 |
| YM012.3 | 117.108536 | -1.5903111 | 0.28768391 | 148.83617 | 3.62E-31 | 1.08E-29 |
| F111A | 205.277135 | -1.5949697 | 0.42420207 | 54.4994981 | 4.14E-11 | 3.51E-10 |
| STYK1.1 | 25.1084355 | -1.5961446 | 0.6937922 | 174.295533 | 1.25E-36 | 4.87E-35 |
| DAF2.1 | 10.8293789 | -1.598363 | 0.76589357 | 90.9690573 | 8.20E-19 | 1.25E-17 |
| PERT | 76.8944591 | -1.6024562 | 0.38349288 | 150.27311 | 1.78E-31 | 5.38E-30 |
| FR1L4 | 166.593706 | -1.6029505 | 0.15736001 | 273.63219 | 5.26E-58 | 6.76E-56 |
| MUC16.1 | 85.8851686 | -1.6125949 | 0.33675091 | 109.406196 | 9.74E-23 | 1.84E-21 |
| PLLP | 11.1332027 | -1.6347272 | 0.57750843 | 32.6877376 | 1.38E-06 | 7.46E-06 |
| CTL4 | 94.9301776 | -1.6389341 | 0.27835332 | 185.682557 | 4.49E-39 | 2.03E-37 |
| HEBP2.2 | 298.78416 | -1.6409274 | 0.13378525 | 877.042721 | 1.57E-188 | 4.00E-185 |
| BPI.1 | 151.995955 | -1.6426445 | 0.26046095 | 103.950731 | 1.42E-21 | 2.50E-20 |
| OTX1B | 10.8546258 | -1.6435041 | 0.58082243 | 56.1781296 | 1.84E-11 | 1.61E-10 |
| SGPP2.1 | 18.4519313 | -1.649363 | 0.47830245 | 85.536453 | 1.17E-17 | 1.63E-16 |
| SAMH1.1 | 14.8231235 | -1.6501928 | 0.46749462 | 33.891672 | 7.84E-07 | 4.37E-06 |
| AHNK.3 | 334.478141 | -1.6614431 | 0.22700763 | 132.440842 | 1.17E-27 | 2.91E-26 |
| S15A1 | 80.2469975 | -1.6632128 | 0.24200966 | 139.765231 | 3.17E-29 | 8.57E-28 |
| ACER1 | 99.050017 | -1.6834553 | 0.25469629 | 93.5955777 | 2.27E-19 | 3.55E-18 |
| ZN214.28 | 91.1341405 | -1.702614 | 0.26378958 | 47.5783566 | 1.16E-09 | 8.63E-09 |
| RHCG | 26.8296277 | -1.7074169 | 0.47033557 | 43.3671928 | 8.68E-09 | 5.91E-08 |
| S38AA.1 | 48.6785397 | -1.7151831 | 0.29592379 | 149.404705 | 2.73E-31 | 8.21E-30 |
| CEBPA | 173.296279 | -1.717653 | 0.16176569 | 903.432429 | 3.01E-194 | 1.15E-190 |
| CPAS1.1 | 186.852475 | -1.7258052 | 0.23343968 | 97.8797746 | 2.78E-20 | 4.57E-19 |
| GLSK.2 | 56.3359645 | -1.745097 | 0.35565415 | 28.3687333 | 1.05E-05 | 5.11E-05 |
| TGM2.4 | 13.2225807 | -1.7700428 | 0.54102983 | 40.9861414 | 2.71E-08 | 1.77E-07 |
| FA83C | 20.2182755 | -1.7757358 | 0.44426451 | 32.8364039 | 1.29E-06 | 7.00E-06 |
| FBN2 | 46.2797823 | -1.7829342 | 0.33183971 | 75.5502115 | 1.52E-15 | 1.83E-14 |
| CASZ1 | 8.94868944 | -1.7921237 | 0.6666641 | 23.355776 | 0.0001075 | 0.0004483 |
| GCNT3.2 | 17.1931145 | -1.7972515 | 0.47101537 | 38.691455 | 8.07E-08 | 4.99E-07 |
| TSCOT.2 | 15.1691504 | -1.8006717 | 0.55122031 | 25.4935151 | 4.00E-05 | 0.00017913 |
| ELF5 | 40.9449237 | -1.8041204 | 0.37132052 | 119.751306 | 6.04E-25 | 1.29E-23 |
| SIM29 | 72.6675053 | -1.8145055 | 0.25647727 | 85.8558627 | 9.99E-18 | 1.40E-16 |
| SYT1.2 | 106.621799 | -1.8158913 | 0.2522152 | 88.7030533 | 2.48E-18 | 3.62E-17 |
| HEYL | 111.673037 | -1.8183638 | 0.19405681 | 173.291123 | 2.06E-36 | 7.93E-35 |
| FYCO1.1 | 32.7305246 | -1.8281039 | 0.35958445 | 94.7547177 | 1.29E-19 | 2.04E-18 |
| CASPE | 88.5814513 | -1.8339311 | 0.43360096 | 215.347156 | 1.88E-45 | 1.19E-43 |
| UPAR.6 | 18.50606 | -1.846752 | 0.70381181 | 36.687345 | 2.09E-07 | 1.24E-06 |
| LIAT1 | 92.6722177 | -1.8483695 | 0.34227217 | 36.1374029 | 2.71E-07 | 1.59E-06 |
| YRD6.50 | 52.0990022 | -1.8856823 | 0.40160806 | 23.4236339 | 0.00010419 | 0.00043581 |
| NIBA1.1 | 61.0530651 | -1.984735 | 0.28604535 | 130.679516 | 2.79E-27 | 6.85E-26 |
| RN208 | 12.8781799 | -1.9902388 | 0.54307541 | 37.9350638 | 1.16E-07 | 7.02E-07 |
| MUC19 | 12.2434092 | -1.9933838 | 0.69311854 | 48.8402995 | 6.30E-10 | 4.84E-09 |
| CDAC1 | 53.8222042 | -2.0024153 | 0.32692673 | 89.5452251 | 1.64E-18 | 2.44E-17 |
| IGL1 | 3.86135076 | -2.0116482 | 1.22784297 | 21.7613949 | 0.00022357 | 0.00088854 |
| HISAT.1 | 13.6085379 | -2.0685235 | 0.61430994 | 36.7951979 | 1.99E-07 | 1.18E-06 |
| AGRF3 | 7.76174426 | -2.0912587 | 0.92730628 | 72.6551256 | 6.24E-15 | 7.10E-14 |
| TSTD1 | 24.8182724 | -2.1242241 | 0.67502132 | 101.651867 | 4.38E-21 | 7.50E-20 |
| TS1R1 | 4.75958729 | -2.1310421 | 1.36252681 | 41.3460192 | 2.28E-08 | 1.50E-07 |
| UROL1 | 15.4688478 | -2.1478854 | 0.69783139 | 92.2735586 | 4.33E-19 | 6.71E-18 |
| CY24B.1 | 58.1295041 | -2.1594014 | 0.3539931 | 74.9642014 | 2.03E-15 | 2.41E-14 |
| S43A3 | 6.06324057 | -2.1942229 | 1.11066486 | 25.5015968 | 3.99E-05 | 0.00017851 |
| AHNK.7 | 6.04598932 | -2.2233347 | 0.93904763 | 31.3314268 | 2.62E-06 | 1.37E-05 |
| TRCG1 | 69.636551 | -2.251852 | 0.32064836 | 129.312434 | 5.46E-27 | 1.31E-25 |
| LORF2.311 | 10.0870532 | -2.2656025 | 0.78416142 | 77.3872981 | 6.23E-16 | 7.67E-15 |
| TRAR1 | 4.68225339 | -2.281148 | 1.1113376 | 32.9091337 | 1.25E-06 | 6.78E-06 |
| CERS2 | 14.6962712 | -2.2922706 | 0.5874498 | 37.8041194 | 1.23E-07 | 7.45E-07 |
| FUT9 | 30.3365172 | -2.2998413 | 0.43444141 | 97.7170934 | 3.01E-20 | 4.94E-19 |
| KMT2D.4 | 100.594802 | -2.3030654 | 0.42973792 | 63.5295045 | 5.25E-13 | 5.10E-12 |
| MALD3.2 | 4.44723173 | -2.3263335 | 1.11106334 | 24.8994554 | 5.27E-05 | 0.00023134 |
| NOS | 63.6195494 | -2.3381387 | 0.57939188 | 56.5649302 | 1.53E-11 | 1.34E-10 |
| BSPRY | 9.84939071 | -2.344972 | 0.74432991 | 30.554071 | 3.77E-06 | 1.93E-05 |
| UPAR.1 | 119.037359 | -2.352827 | 0.56221865 | 29.0953898 | 7.48E-06 | 3.71E-05 |
| KMT2D.2 | 233.813284 | -2.4351353 | 0.53717996 | 37.5919469 | 1.36E-07 | 8.18E-07 |
| YM012.1 | 317.484418 | -2.4409269 | 0.38969759 | 66.2808629 | 1.38E-13 | 1.39E-12 |
| OTX5 | 4.78497949 | -2.4476631 | 1.37197668 | 38.8357877 | 7.53E-08 | 4.67E-07 |
| K118B | 1019.58295 | -2.4507069 | 0.18267685 | 223.079826 | 4.08E-47 | 2.87E-45 |
| NOD1.4 | 7.14258349 | -2.5542945 | 0.87043092 | 22.4673841 | 0.00016174 | 0.00065837 |
| K2C8 | 769.867238 | -2.6079683 | 0.19226183 | 284.04732 | 2.99E-60 | 4.35E-58 |
| LORF2.243 | 18.0798287 | -2.6545193 | 0.66522335 | 22.154126 | 0.00018675 | 0.00075077 |
| DCE2 | 6.08432705 | -2.6563404 | 1.54314844 | 42.484325 | 1.32E-08 | 8.87E-08 |
| YV023 | 232.396445 | -2.6605426 | 0.47492688 | 52.9180873 | 8.86E-11 | 7.28E-10 |
| USPL1.1 | 15.8951622 | -2.6677766 | 0.64711599 | 22.626098 | 0.00015037 | 0.00061471 |
| CSPG5 | 11.6663278 | -2.7564756 | 1.14067627 | 135.953821 | 2.07E-28 | 5.40E-27 |
| ICEF1 | 4.20306092 | -2.8528861 | 1.34573257 | 26.131253 | 2.98E-05 | 0.00013623 |
| LORF2.135 | 4.37268726 | -3.085662 | 1.47664673 | 35.2544669 | 4.12E-07 | 2.37E-06 |
| IRX6 | 7.22860701 | -3.1394412 | 1.34293634 | 84.6942024 | 1.76E-17 | 2.42E-16 |
| TS1R3 | 6.80755903 | -3.3273167 | 1.40655802 | 27.3211075 | 1.71E-05 | 8.09E-05 |
| PDIA1.1 | 5.08034465 | -3.4349461 | 1.59473119 | 23.7937705 | 8.78E-05 | 0.00037213 |
| UD12.1 | 5.7820951 | -3.6764831 | 1.49151286 | 22.5857456 | 0.00015319 | 0.00062521 |
| PA2GX | 19.0444364 | -4.5990595 | 1.32076512 | 114.913315 | 6.51E-24 | 1.33E-22 |
| IGG2A | 9.38389065 | -4.813957 | 1.24699872 | 89.983512 | 1.33E-18 | 1.98E-17 |

Movie S1. Contrast-enhanced Micro-Computed Tomography (CT) of the catshark lower jaw with soft tissue segmentation of the compartments sequenced (RNAseq) in this study. Segmentation movie shows slices through the jaw revealing the various compartments (BHTB, blue; TTJ, purple; LT, yellow; ET, Orange; SL, red) and teeth, white.
